# Immunosuppressive tocilizumab prevents astrocyte induced neurotoxicity in hiPSC-LRRK2 Parkinson’s disease by targeting receptor interleukin-6

**DOI:** 10.1101/2022.08.23.504918

**Authors:** Meritxell Pons-Espinal, Lucas Blasco-Agell, Irene Fernandez-Carasa, Angelique di Domenico, Yvonne Richaud, Jose Luis Mosquera, Laura Marruecos, Lluís Espinosa, Alicia Garrido, Eduardo Tolosa, Michael J. Edel, Manel Juan Otero, Isidre Ferrer, Angel Raya, Antonella Consiglio

**Author notes:** Correspondence and requests for materials should be addressed to: A.C. and /or M.P.E.

## Abstract

Parkinson’s disease (PD) is associated with premature death of dopamine-producing neurons in the brain. Previous studies have shown that astrocytes of PD patients may contribute to neuronal degeneration by mechanisms involving both direct cell-to-cell contact and transfer of soluble molecules. Since it has been proposed that PD patients exhibit an overall pro-inflammatory state, and since astrocytes are key mediators of the inflammation response in the brain, here we sought to address whether astrocyte-mediated inflammatory signaling could contribute to PD neuropathology. For this purpose, we generated astrocytes from induced pluripotent stem cells (iPSCs) representing PD patients and healthy controls. Transcriptomic analyses identified a unique inflammatory gene expression signature in PD astrocytes compared to controls. In particular, the pro-inflammatory cytokine IL-6 was found to be highly expressed and released by PD astrocytes, and to induce toxicity in dopamine neurons. Mechanistically, neuronal cell death was mediated by IL-6 signaling via IL-6 receptor (IL-6R) expressed in human PD neurons, leading to downstream activation of STAT3. Importantly, astrocyte-induced cell death in PD disease midbrain neurons could be prevented by blocking IL6R-mediated signaling using clinically available antibodies. Moreover, examination of postmortem tissue brain of early-stage PD patients uncovered increased numbers of dopamine neurons overexpressing IL-6R and of reactive astrocytes overexpressing IL-6, compared to healthy brains. Our findings highlight the potential role of astrocyte-mediated inflammatory signaling in neuronal loss in PD, and open the way for new therapies based on IL-6 immunomodulation for preventing PD pathogenesis.

## Introduction

Parkinson’s disease (PD) is a common, chronic and incurable neurodegenerative disease associated with a selective loss of dopamine-producing neurons in the midbrain responsible for controlling body movements (1, 2). Although most cases of PD are of unknown cause, the so-called idiopathic PD, approximately 5% has been shown to have a genetic basis, including mutations in the LRRK2 gene encoding for leucine rich repetitive kinase 2 (LRRK2), which are found in the largest number of patients with familial PD (3). Furthermore, it was found that mutations in the LRRK2 gene are also associated with ID-PD and typically occur with late onset disease that closely resembles ID-PD in its neuropathological changes and response to therapy. These changes include typical α-synuclein aggregates in the form of Lewy bodies and neurites and the loss of cells in vulnerable areas. The close similarities between LRRK2-PD (L2-PD) and ID-PD raise the possibility that unraveling the mechanisms underlying neurodegeneration in L2-PD may provide new insights into the pathophysiology and treatment of ID-PD.

Associated with neurodegeneration, several neuroinflammatory signs have been described in PD. A link between imbalanced inflammatory signaling and PD, and its correlation with disease progression and motor symptoms, has been suggested by several lines of evidence. At the histopathological level, activated glial cells and lymphocytes infiltration have been observed in PD post-mortem brains (4–6), together with higher pro-inflammatory cytokines such as interleukin (IL)-6 in serum, cerebrospinal fluid (CSF) and substantia nigra pars compacta (SNc) (7–9) already at Braak stage 1 and 2. Moreover, higher IL-6 levels have been correlated with a worst PD prognosis (10, 11). Astrocytes are emerging as critical immunocompetent cells that mediate inflammatory responses. Following different pathological insults, astrocytes undergo a pronounced transformation called ‘astrocyte reactivity’, characterized by both structural and biochemical changes such as high levels of glial fibrillary acidic protein (GFAP) and alterations in their immunocompetent capacity (6, 12–14). Recent observations found A1 “neurotoxic” reactive astrocytes in PD post-mortem brains (6, 15). However, the mechanisms driving astrocyte reactivity in PD and how this impacts their function and pathogenesis remain unclear.

We have recently provided the first direct experimental evidence that astrocytes from PD patients, carrying the LRRK2 G2019S mutation, present dysfunctional protein degradation pathways, leading to α-syn accumulation and propagation that affects healthy DAn in an induced pluripotent stem cell (iPSC) based model (16). In this line, it has been recently confirmed that L2-PD astrocytes fail to provide full neurotrophic support to DAn (17). However, the exact mechanisms involved remain unknown. L2-PD astrocytes display atrophic morphology and altered mitochondrial function (18), metabolic alterations (19), decreased capacity to internalize α-syn (20) and downregulated genes involved in extracellular matrix (21). Yet, whether those physio-pathological alterations in L2-PD astrocytes are the cause or a consequence of the disease and whether astrocyte play a role in idiopathic forms of PD impacting on neuronal survival is unknown.

In this study, we used an experimental platform based on astrocyte cultures derived from ID-PD patients, L2-PD and their gene-edited isogenic counterparts, or from healthy individuals. PD astrocytes showed changes in transcriptome with a unique inflammatory gene expression signature and increased secretion of the pro-inflammatory cytokine IL-6. Conditioned medium from PD astrocytes induced neurotoxicity to surrounding dopaminergic neurons, and this effect was partially blocked by the IL-6 inactivating antibody. We confirmed higher IL-6R in PD patients, highlighting the potential role of astrocyte-mediated inflammatory signaling in PD neuropathology.

## Results

### hiPSC-derived L2-PD astrocytes are morphologically reactive and display an increase in pro-inflammatory markers

We have previously described the generation of astrocytes from human iPSCs following a protocol that efficiently allows propagating, expanding, freezing and thawing proliferating glial intermediates (16, 22). Using this method, astrocytes from three L2-PD patients carrying the G2019S mutation (L2-PD), three healthy donors (CTL), and one isogenic iPSC patient line with the LRRK2 G2019S missense mutation corrected (L2-PD^corr^) (**Supplemental Table 1**), were generated and cultured them for two weeks. Interestingly, we observed that L2-PD astrocytes adopted a hypertrophic morphology with thin and retracted processes as shown by GFAP staining (**Figure 1, A-C** and **Supplemental Figure 1A**; GFAP intensity p<0.05; Form Factor p<0.001), similarly to CTL astrocytes when stimulated with a cocktail of cytokines (C1q, TNFα and IL-1α) (6) (**Figure 1, A-C**; GFAP intensity p<0.01; Form Factor p<0.05). Immunocytochemistry (ICC) and quantitative real time PCR (qRT-PCR) of canonical markers of reactive astrocytes (“pan reactive”), such as Vimentin and AQP4 (**Figure 1,** D and F p<0.05), as well as A1-specific proteins such as C3 (**Figure 1, D-E** p<0.05) confirmed an overexpression of those markers indicating that L2-PD astrocytes are reactive.

**Figure 1.**
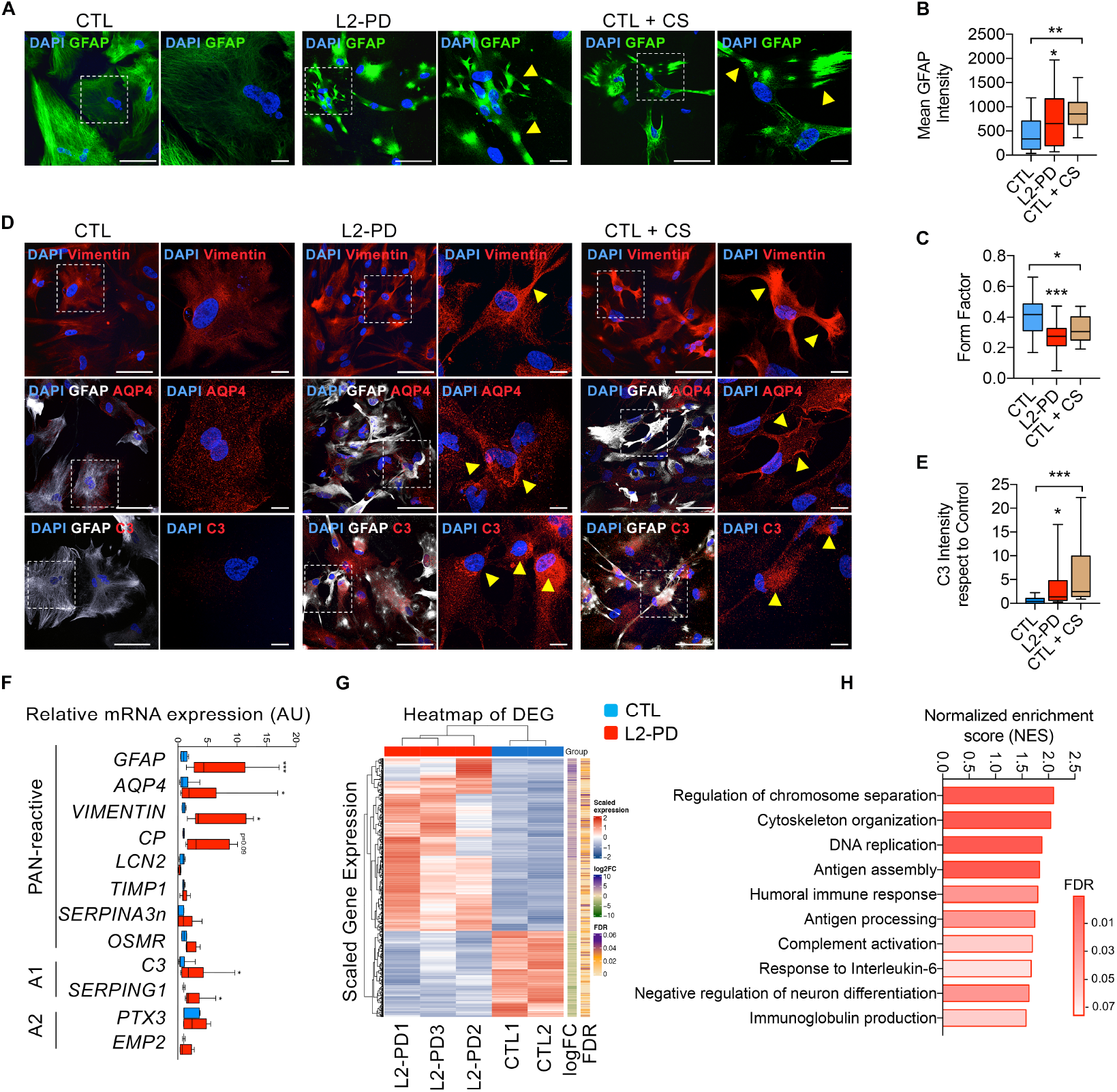
iPSC-derived L2-PD astrocytes are reactive. (**A**) Representative immunocytochemistry (ICC) images of CTL (SP09) and L2-PD astrocytes (SP13) expressing DAPI (blue) and GFAP (green) after 14 days in culture. Scale bar: 100 μm. CTL astrocytes (SP09) treated for 48h with C1q, TNFα and IL-1α were used as positive control. Images on the right show a magnification of the area boxed in the left images; scale bar: 10 μm. Yellow arrows indicate high GFAP staining in hypertrophic astrocytes. (**B**) Mean intensity of GFAP staining. (**C**) Form factor of GFAP positive cells calculated as: FF = 4pi(area/perimeter^2). (**D**) Representative ICC images of CTL, L2-PD and activated CTL astrocytes expressing Vimentin (CTL: SP09; L2-PD: SP13), AQP4 (CTL: SP17; L2-PD: SP06) and C3 (CTL: SP09; L2-PD: SP13). Scale bar: 100 μm. Images on the right show a magnification of the area boxed in the left images; scale bar: 10 μm. Yellow arrows indicate high expression of the specific marker shown in the images. (**E**) Mean intensity of C3 staining respect to CTL. (**F**) Relative mRNA expression of panreactive, A1-specific and A2-specific transcripts in L2-PD astrocytes respect to CTL. Box-and-whisker plots show median, 25th and 75th percentiles, minimum, and maximum values (n=3 experiments; Form factor, Mean GFAP and C3 intensity was performed from 30 astrocytes per experiment per condition). One-way ANOVA Bonferroni as post-hoc. *p<0.05; **p<0.01; ***p<0.001. (**G**) Heat map of differentially expressed genes (DEG) of CTL and L2-PD astrocytes. (**H**) Selected upregulated gene sets derived from the Gene Ontology (GO) biological processes between L2-PD and CTL astrocytes.

To evaluate if L2-PD astrocytes have an altered inflammation-driven signaling that may play a role in early stage of L2-PD, we pursued an unbiased transcriptional analysis using next generation deep RNA sequencing (RNA-seq) to identify altered signaling pathways in PD-astrocytes. We detected 656 differentially expressed genes (DEG) out of 19383 genes with False Discovery Rate (FDR) <0.05 and |Fold change| > 2 (**Figure 1G and Supplemental Figure 1, B-C**). Gene Set Enrichment Analysis (GSEA), conducted based on gene sets derived from the Gene Ontology (GO) Biological Process ontology, revealed a total of 2874 out of 4721 gene sets upregulated (259 FDR <25%; 212 p<0.01) and 1847 out of 4721 gene sets downregulated (19 FDR <25%; 57 p<0.01) in L2-PD astrocytes compared to CTL astrocytes (**Supplemental Table 2**). This functional enrichment analysis showed significant upregulation of gene sets in L2-PD astrocytes related with the regulation of chromosome separation (GO:1905818; 2.17 NES; p=0; FDR<0.001), DNA replication (GO:0006260; 1.87 NES; p=0; FDR<0.001) and inflammation-related GO terms such as antigen assembly with MHC protein complex (GO:0002501; 1.83 NES; p=0; FDR<0.01), humoral immune response (GO:0002455; 1.80 NES; p=0; FDR<0.02), complement activation (GO:0006958; 1.69 NES; p=0; FDR<0.05) and response to IL-6 (GO:0070741; 1.67 NES; p=0; FDR <0.07) (**Figure 1H**). Given the relevance of altered cytokine and immune response-related genes in L2-PD, we next examined their secretome cytokine profile. We found that while most secreted cytokines were at comparable levels between CTL and L2-PD, IL-6 was significantly higher in L2-PD astrocytes similar to CTL cytokine-activated astrocytes (**Figure 2A** and **Supplemental Figure 1D**; p<0.01), suggesting that L2-PD astrocyte-specific effects might lie downstream of IL-6. Importantly, IL-6 was significantly reduced to CTL levels in L2-PD^corr^ astrocytes (**Figure 2A**; p<0.05), indicating that specific neuroinflammatory alterations associated with the *LRRK2 G2019S* mutation is directly related to astrocyte immune function. In line with our cytokine profiling data, we found that L2-PD^corr^ astrocytes did not show an altered morphology, nor and increase in GFAP, AQP4 or C3 overstaining through ICC analysis (**Figure 2, B-E** and **Supplemental Figure 1E**).

**Figure 2.**
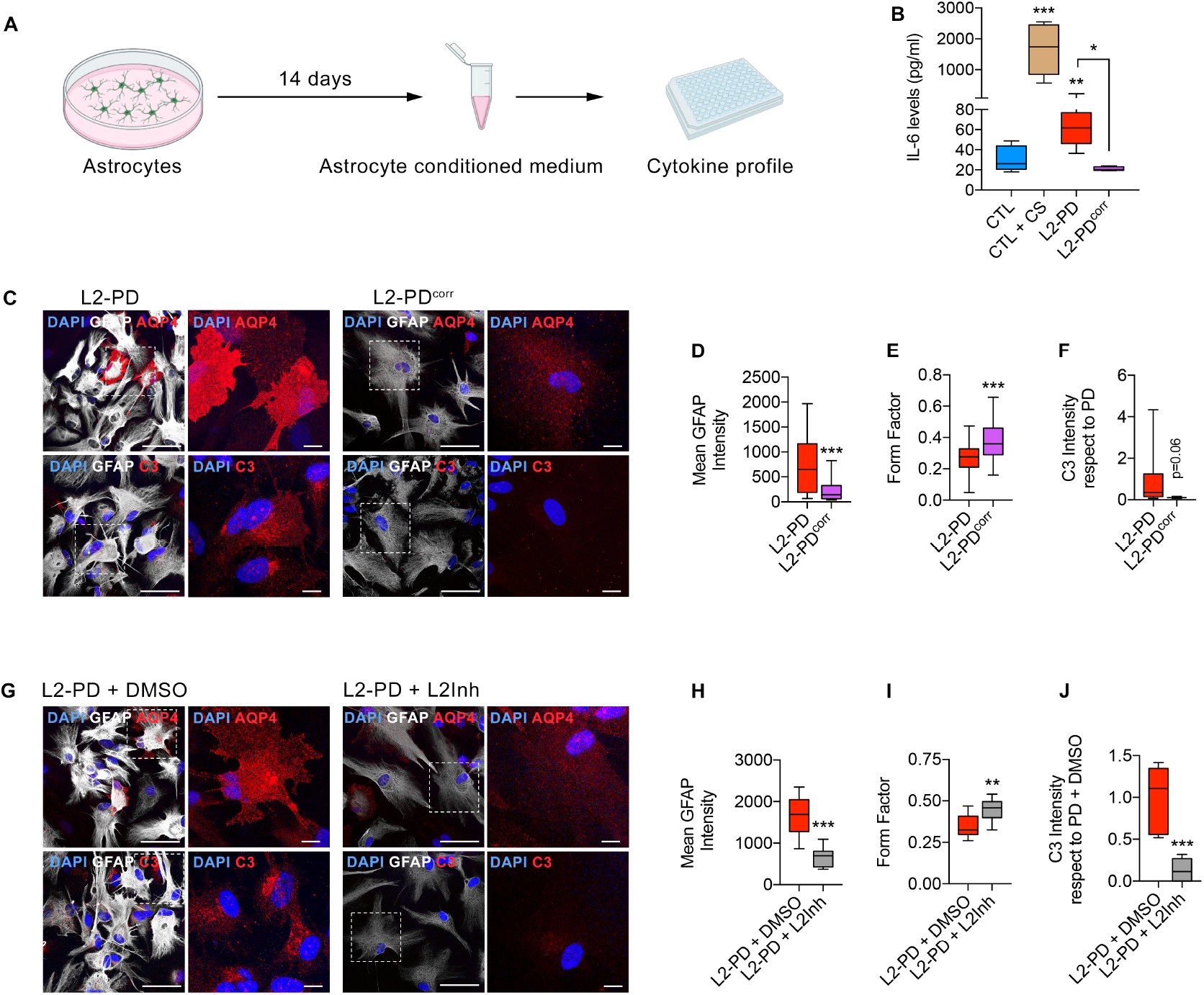
L2-PD astrocyte reactivity is mediated by LRRK2 G2019S-increased kinase activity. Schematic experimental set up to perform a cytokine array from astrocyte’s conditioned medium. (**B**) Cytokine IL-6 levels released by CTL, L2-PD and isogenic (L2-PD^corr^) astrocyte conditioned medium (ACM) after 2 weeks in culture. CTL astrocytes were treated for 48h with C1q, TNFα and IL-1α as positive control. (**C**) Representative ICC images of L2-PD (SP13) and L2-PD^corr^ (SP13wt/wt) astrocytes staining positive for: top panels DAPI (blue), GFAP (white) and AQP4 (red); bottom panels DAPI (blue), GFAP (white) and C3 (red). Scale bar: 100 μm. Images on the right show a magnification of the area boxed in the left images; scale bar: 10 μm. (**D**) Mean intensity of GFAP staining. (**E**) Form factor of GFAP positive cells calculated as: FF = 4pi(area/perimeter^2). (**F**) Mean intensity of C3 staining respect to L2-PD astrocytes. (**G**) Representative ICC images of L2-PD (SP13) treated with either DMSO or LRRK2-kinase inhibitor (1uM) astrocytes staining positive for: top panels DAPI (blue), GFAP (white); middle panels DAPI (blue) GFAP (white) and AQP4 (red); bottom panels DAPI (blue), GFAP (white) and C3 (red). Scale bar: 100 μm. Images on the right show a magnification of the area boxed in the left images; scale bar: 10 μm. (**H**) Mean intensity of GFAP staining. (**I**) Form factor of GFAP positive cells calculated as: FF = 4pi(area/perimeter^2). (**J**) Mean intensity of C3 staining respect to L2-PD astrocytes treated with DMSO. Box- and-whisker plots show median, 25th and 75th percentiles, minimum, and maximum values (n=3 experiments; Form factor, Mean GFAP and C3 intensity was performed from 30 astrocytes per experiment per condition). One-way ANOVA Bonferroni as post-hoc. Student t-test or Mann-Whitney test for non-parametric conditions were used when only two groups were compared. *p<0.05; **p<0.01; ***p<0.001.

To test whether LRRK2-kinase activity is involved in astrocyte’s reactivity, we treated L2-PD astrocytes with a LRRK2 kinase inhibitor for one week. We found that inhibition of LRRK2 kinase activity in L2-PD astrocytes prevented astrocyte’s reactivity, as shown by reduced GFAP and C3 intensity (**Figure 2, F-G and I**; p<0.001), and increased form factor (**Figure 2H**; p<0.01). Importantly, inhibition of LRRK2 kinase activity significantly downregulated *IL-6* expression (**Supplemental Figure 1G**; p<0.01) along with Toll like receptors *(TLR2* and *TLR4)* and inflammasome *(NLRP1)* as compared to non-treated conditions (**Supplemental Figure 1F**). In contrast, overexpression of LRRK2 G2019S (pDEST51-*LRRK2* G2019S) in CTL astrocytes significantly increased *IL-6* mRNA levels (**Supplemental Figure 1G**; p<0.01). These results suggested that the induction of IL-6 expression and neuroinflammatory alterations in human L2-PD astrocytes is mediated through the LRRK2 G2019S kinase activity.

### L2-PD astrocytes induce DA neurodegeneration by IL-6/IL-6R signaling

We have recently described that astrocyte-induced neurotoxicity in healthy DAn was not only observed through direct cell contact, but also through indirect mechanisms using L2-PD astrocyte conditioned medium (ACM) (16). This opened the possibility that some factors might be secreted from astrocytes in addition to α-syn contributing to DAn neurodegeneration. Recently, it was shown that A1 reactive astrocytes in rodents are highly neurotoxic and rapidly kill neurons through the release of saturated lipids. Yet, reducing these lipids does not eliminate neurotoxicity, thus suggesting the presence of other unknown neurotoxins that may contribute to neurodegeneration (6, 14, 23).

To test whether IL-6 released by L2-PD astrocytes could be involved in DAn neurodegeneration, we added Tocilizumab (a humanized monoclonal antibody against IL-6R FDA-approved, clinically used in rheumatoid arthritis) to L2-PD-ACM and treated hiPSC-derived CTL or L2-PD neurons for one week (**Figure 3A**). In concordance with a putative IL-6 signaling, LRRK2-PD-ACM significantly increased phosphorylated Signal Transducer and Activator of Transcription 3 (p-STAT3) nuclear levels in neurons (**Supplemental Figure 2, A-C;** p<0.05) and Suppressor Of Cytokine Signaling 3 *(SOCS3,* a STAT-induced STAT inhibitor) mRNA levels (Figure S2D; p<0.001), whereas treatment with Tocilizumab reduced nuclear P-STAT3 and *SOCS3* mRNA levels to CTL-ACM levels (**Supplemental Figure 2, A-D**). Moreover, the chronic addition of IL-6 in CTL-ACM to CTL DAn induced neurodegeneration in a concentration dependent manner as shown by reduced survival of Tyrosine Hydroxilase (TH) positive neurons (**Supplemental Figure 2, F-H**; p<0.05) and neurite arborization (**Supplemental Figure 2, I-J**; p<0.001), whereas blockage of IL-6 signaling prevented these alterations in DAn (**Supplemental Figure 2, G-J**). These results highlight the role of IL-6/IL-6R signaling in astrocyte-neuron communication in PD.

**Figure 3.**
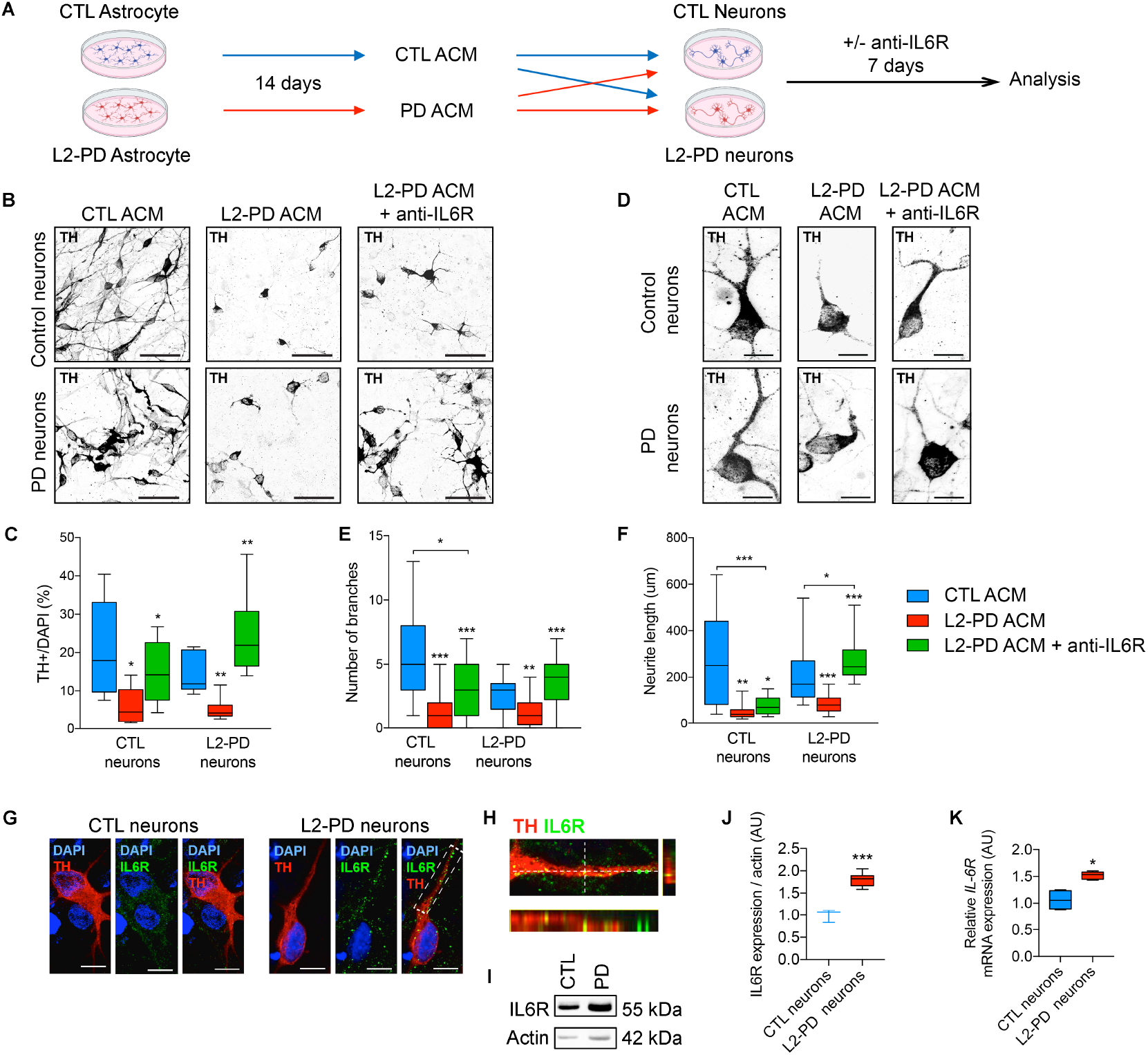
L2-PD astrocytes induce iPSC-derived DA neuronal degeneration through IL-6/IL-6R signaling. (**A**) Schematic representation of the experimental procedure to analyze the effect of astrocyte conditioned medium (ACM) and IL-6 involvement on neuronal survival and degeneration. ACM was collected after 14 days in culture and added to iPSC-derived dopaminergic (DA) neurons from both CTL and L2-PD patients for one week. Anti-IL6R antibody Tocilizumab (10ug/ml) was added to ACM. (**B**) Representative ICC images of iPSC-derived CTL (SP11) and L2-PD neurons (SP12) expressing tyrosine hydroxylase (TH, black) treated for one week with CTL ACM, L2-PD ACM and L2-PD ACM + anti-IL6R antibody. Scale bars: 100 μm. CTL neurons are derived from SP11 iPSC line. LRRK2-PD neurons are derived from SP12, SP06 and SP13 iPSC lines. (**C**) Percentage of TH+ cells respect to DAPI. (**D**) Representative ICC images of iPSC-derived CTL and LK2-PD neurons staining for TH (black). Scale bars: 20 μm. (**E**) Number of branches and (**F**) neurite length of CTL and L2-PD TH+ neurons treated for one week with ACM. (**G**) Representative ICC images of DAn (TH+, red) expressing IL-6R (green) from CTL (SP11) and L2-PD (SP13) iPSC-derived DAn after 35 days of differentiation (Scale bar: 20 μm). (**H**) Orthogonal views show colocalization between IL6R and TH in L2-PD (SP13) DAn. (**I**) Representative western blot for IL6R of iPSC-derived CTL neurons (SP11) and L2-PD neurons (SP12). (**J**) Quantification of protein IL6R respect to Actin. (**K**) Relative *IL6R* mRNA expression from iPSC-derived CTL neurons (SP11 and SP17) and L2-PD neurons (SP12 and SP13). Box-and-whisker plots show median, 25th and 75th percentiles, minimum, and maximum values (n=3 experiments; 30 neurons per experiment per condition for each line). One-way ANOVA Bonferroni as post-hoc. Student t-test or Mann-Whitney test for non-parametric conditions were when only two groups were compared. *p<0.05; **p<0.01; ***p<0.001.

Addition of L2-PD-ACM to DAn significantly affected DAn survival as revealed by a reduced number of TH positive cells (**Figure 3, B-C**; CTL neurons p<0.05; L2-PD neurons p<0.01) and increased expression of *Caspase 3* mRNA levels (Supplemental Figure 2E, p<0.05). Moreover, surviving TH+ cells displayed morphological alterations, including fewer and shortened neurites (**Figure 3, D-F**). Remarkably, Tocilizumab treatment for one week promoted DAn survival and prevented neurodegeneration induced by L2-PD-ACM (**Figure 3, B-F and Supplemental Figure 2E**) to both CTL and L2-PD neurons. However, the effect was significantly more pronounced in L2-PD neurons as compared to CTL ones as shown by higher percentage of TH+ neurons (**Figure 3, B-C**; CTL neurons p<0.05; L2-PD neurons p<0.01) and increased neurite complexity (**Figure 3, D-F**; CTL neurons p<0.05; L2-PD neurons p<0.001). These results shed light on the putative use of specific IL-6 blockers to significantly prevent DAn neurodegeneration in PD.

### L2-PD neurons are vulnerable to IL-6 signaling

We have consistently observed that L2-PD neurons are more sensitive to anti-IL6R treatment compared to CTL neurons, even completely reversing neuronal abnormalities. In order to investigate the molecular mechanisms responsible, we first examined the expression of IL-6 receptor (IL-6R) in both CTL and L2-PD iPSC-derived DAn by ICC and qRT-PCR expression analysis. We found that TH+ neurons express IL-6R (**Figure 3, G-H**), however that L2-PD DAn harbor an increased expression of IL-6R both at an mRNA and protein levels compared to CTL DAn (**Figure 3, I-K and Supplemental Figure 2K**; p<0.05). Moreover, L2-PD neurons significantly induced higher IL-6R downstream signaling effectors as shown by the nuclear translocation of p-STAT3 along with increased STAT3 protein expression after the addition of L2-PD ACM, compared to CTL DAn (**Supplemental Figure 2**, L-N; p<0.05). These results suggest that increased expression of IL6R in PD DAn may explain specific neuronal vulnerability to IL-6 and therefore, to IL-6 secreted by PD-derived astrocytes, who are the main producers of IL-6. Notably, prevention of PD astrocyte-mediated neuronal degeneration through specific blockage of IL-6R points toward new potential immunotherapeutic approaches for PD.

### IL-6/IL-6R mediates neuronal degeneration in idiopathic PD patients both in vitro and in vivo

To understand the impact of IL-6/IL-6R signaling in PD pathophysiology, we examined whether idiopathic PD (ID-PD) astrocytes harbored similar pathology to familial L2-PD astrocytes. To investigate this, we generated astrocytes from three different ID-PD patients with no familiar history of PD (**Supplemental Table 1**). We obtained ~95% of astrocyte purity as shown by the expression of GFAP, S100b and CD44, and the absence of TUJ1, MAP2 and NG2 (neuronal and oligodendrocyte markers, respectively) (**Supplemental Figure 3, A-B**). Similar to familial L2-PD, ID-PD-derived astrocytes display a hypertrophic and ramified morphology (**Figure 4, A-B**), overexpress GFAP and C3 at protein levels (Figure 4, C-D) comparable to CTL astrocytes stimulated with cytokines, and upregulated inflammatory-related genes (**Supplemental Figure 3C**), suggesting a reactive phenotype. Moreover, we found that ID-PD-astrocytes consistently release higher levels of IL-6 compared to unstimulated CTL astrocytes (**Figure 4E**; p<0.01). To examine whether IL-6 could also be involved in astrocyte-dependent neurotoxicity in ID-PD, we added Tocilizumab to idiopathic ACM and treated CTL and PD neurons for one week. Addition of ID-PD ACM significantly reduced the number of TH+ cells (Figure 4, F-G; p<0.05), as well as their neurite complexity (**Figure 4**, H-I; p<0.001). Importantly, blocking IL-6 signaling prevented signs of neurodegeneration (**Figure 4, F-I**), thus reinforcing the idea that IL-6 is one of the main astrocyte-dependent neurotoxic factors mediators found in the pathogenesis of PD.

**Figure 4.**
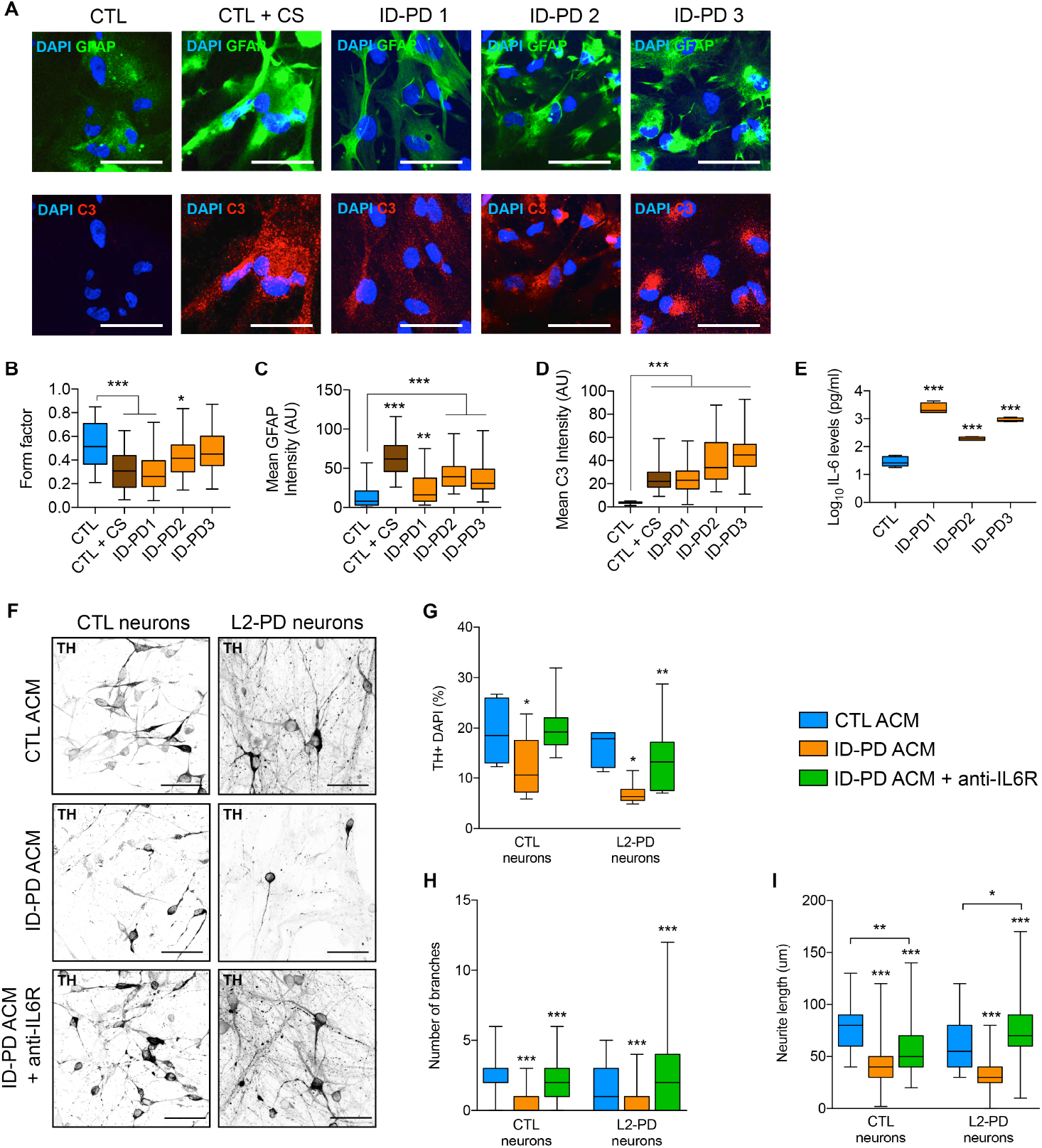
IL-6 signaling mediates neuronal degeneration in idiopathic PD patients. (**A**) Representative ICC images of CTL (SP09) or idiopathic PD (ID-PD1: SP04; ID-PD2: SP08; ID-PD3: SP16) astrocytes staining positive for: top panels DAPI (blue) and GFAP (green); bottom panels DAPI (blue) and C3 (red). CTL astrocytes treated for 48h with cytokines were used as positive control. Scale bar: 100 μm. (**B**) Form factor of GFAP positive cells calculated as: FF = 4pi(area/perimeter^2). (**C**) Mean intensity of GFAP staining. (**D**) Mean intensity of C3 staining respect to CTL. (**E**) IL-6 protein levels released by CTL and ID-PD ACM after 2 weeks in culture. Box-and-whisker plots show median, 25th and 75th percentiles, minimum, and maximum values (n=3 experiments; Form factor, Mean GFAP and C3 intensity was performed from 30 astrocytes per experiment per condition). (**F**) Representative ICC images of iPSC-derived CTL (SP11) and L2-PD (SP12) neurons expressing tyrosine hydroxylase (TH, black) treated for one week with CTL astrocyte conditioned medium (ACM), idiopathic (ID-PD) ACM and ID-PD ACM + anti-IL6R Tocilizumab (10ug/ml). Scale bar: 100 μm. (**G**) Percentage of TH+ cells respect to DAPI. (**H**) Number of branches and (**I**) neurite length of TH+ neurons cultured with ACM for one week. Box-and-whisker plots show median, 25th and 75th percentiles, minimum, and maximum values (n=3 experiments; 30 neurons per experiment per condition for each line). One-way ANOVA Bonferroni as post-hoc: *p<0.05; **p<0.01; ***p<0.001.

In accordance, we evaluated the presence of IL-6 and IL-6R by ICC in the SNc from CTL and ID-PD post-mortem brain tissue at stages 2 (early) and 4 (late) (**Figure 5A**). We found that IL-6R is mainly localized in larger shaped neurons in CTL brains, and interestingly, expression was found to be highly elevated in PD tissue already in early stages of the disease (**Figure 5B**), confirming our findings in our hiPSC-derived neuronal model. In contrast, IL-6 expression was higher in astrocytes from PD post-mortem brain tissue compared to CTL (**Figure 5C**). Importantly, although IL-6 immunoreactivity in astrocytes was high at early stages, it significantly increased at later stages of PD, suggesting a progressive increase throughout the pathogenesis of the disease. Taking into account that IL-6R neuronal immunoreactivity was already present at high levels at early stages of PD, our results suggest that PD neurons may be vulnerable to IL-6, which is being massively produced by PD-astrocytes, hence resulting in a loss of DAn. Our findings shed light on the discovery of new immunomodulators targeting IL-6 signaling in the pathophysiology of PD.

**Figure 5.**
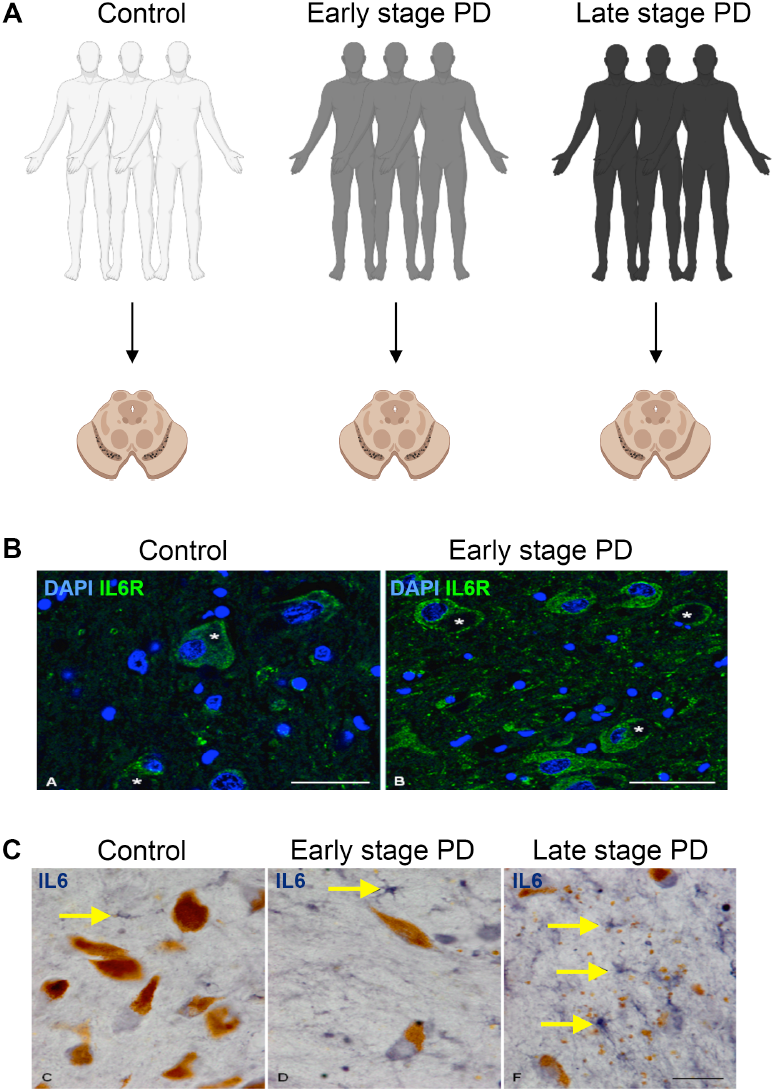
IL-6/IL-6R signaling is increased in postmortem brains during early PD stages. (**A**) Postmortem samples from the substantia nigra were obtained from healthy individuals and PD patients both at early and late stages of the disease. (**B-C**) Immunoreactivity of IL-6R (B) and IL-6 (C) in the substantia nigra pars compacta in control and sporadic PD postmortem brain samples at early (2) and late (4) Braak stages. IL-6R is localized in large neurons (asterisk) whereas IL-6 is present in astrocytes (yellow arrows). Scale bars (B) images = 40μm; (C) images: 25μm.

## Discussion

In this study, we identified pro-inflammatory cytokine IL-6 as an essential modulator by which PD astrocytes induce neurotoxicity in DAn in humans both in vitro and in vivo. We found that the LRRK2 G2019S mutation induces astrocyte reactivity and alters inflammatory-dependent pathways leading to a specific overproduction and release of IL-6. This is mediated by hyperactive LRRK2 kinase activity. Once released into the extracellular space, IL-6 induces STAT3 pathway activation in neurons, impairing neuronal survival and neurite complexity. Thus, our study provides evidence of a novel mechanism for the emerging notion of reactive astrocytes as neurotoxic promoters of neurodegeneration in L2-PD and ID-PD.

Studies investigating PD pathogenesis have been mostly focused on intrinsic mechanisms underlying DAn degeneration and death. However, there is growing evidence that indeed other cell types, apart from DAn, especially glial cells, may exhibit their own specific regulatory features and therefore, could be contributing to PD pathology (24–26). Our previously published results demonstrated a novel, non-cell autonomous astrocytic-role of neurodegeneration in PD, elicited by dysfunctional protein degradation pathways in astrocytes that lead to the accumulation and propagation of endogenously produced α-syn, which was then transferred to healthy DAn in a co-culture model causing neurodegeneration (16). After rescuing Chaperone Mediated Autophagy with an activator compound, lysosomal function was restored and α-syn was cleared from the PD-astrocytes. However, under these conditions, healthy neurons only partially rescued their neurite arborization, therefore the notion that astrocytes are releasing other neurotoxic factors contributing to PD pathology remained to be discovered.

It is well established that innate and adaptive immune cells play a crucial role on inflammation-dependent neurodegeneration in PD (5, 27, 28). Associated with the immune system, the presence of reactive astrocytes (A1) in post-mortem brain tissue from PD patients has been described (6). Reactive astrocytes have also been found in the α-syn preformed fibril (α-syn PFF) mouse model of ID-PD, in which the authors identified the microglial glucagon-like peptide-1 receptor (GLP-1R) necessary to prevent the microglial-mediated conversion of astrocytes into A1 neurotoxic phenotype (15). However, it remains unknown whether they hinder or support CNS recovery, since recently the presence of a subtype of A1 astrocytes overexpressing C3 has been key to prevent prion like disease progression (29). The fact that this reaction is observed not only in PD, but present in other neurodegenerative diseases such as Alzheimer disease and amyotrophic lateral sclerosis, stroke and traumatic brain injury, suggests that glial involvement is secondary to neuronal death. However, recent hypotheses suggest that astrocyte-mediated reactivity could be neuroprotective and have a positive effect locally at the initial phases of disease (e.g. removal of aberrantly firing neurons), while over extended periods of time, become neurotoxic and detrimental (e.g. glia hyper-clearing and killing of too many neurons). Moreover, the presence of specific mutations in certain genes may promote the switch of glial cells to a hyper-reactive phenotype that never turns off, which accelerates neurodegeneration.

We found that IL-6 is present in reactive astrocytes both at early and late stages of PD through the analysis of post-mortem brain tissue. Although we could not rule out whether reactive astrocytes appear as a secondary response to DAn degeneration in vivo, using an iPSC-based model we found that familial L2-PD astrocytes are reactive, defined by the transcriptomic signature, morphology and C3 upregulation (typical marker of A1 neurotoxic astrocytes) (6, 30), as well as the increased release of cytokine IL-6 without being in contact with PD neurons. These results suggest that these altered reactive cell responses might be influenced by the presence of specific genetic mutations significantly linked to PD, such as LRRK2 G2019S. Indeed, L2-PD^corr^ astrocytes with the LRRK2 G2019S missense mutation corrected, as well as the use of a LRRK2 kinase inhibitor, significantly restored inflammatory-dependent phenotypes to unstimulated CTL levels. Thus, suggesting that the presence of reactive astrocytes depends on LRRK2 kinase hyperactivation. Importantly, increased levels of IL-6 were also detected in ID-PD astrocytes, thus reinforcing the notion that neurotoxic levels of IL-6 could be a common mechanism in PD pathology. Although the mechanisms by which IL-6 is increased in ID-PD astrocytes remains unknown, recent studies point to the presence of hyperactivated kinase LRRK2 in patient cells (31).

Importantly, we found that IL-6/IL-6R signaling might be particularly susceptible in PD, since human PD neurons overexpress IL-6R both in vitro and in vivo in post-mortem brain tissue. This results in the downstream activation of STAT3 upon exposure to PD-ACM. In accordance, previous studies found increased levels of nuclear pSTAT3 in the SNc of PD mouse models (32). Moreover, blocking IL-6 signaling in a purely PD context (PD-neurons/PD-astrocyte) with a specific antibody FDA-approved (Tocilizumab) prevented neuronal degeneration significantly more compared to CTL neurons treated with PD-ACM. These results elucidate a new potential immunotherapeutic approach to treat PD. The mechanisms by which IL-6R is upregulated in PD remain unknown.

Preliminary results from our lab suggest very mild associations between IL-6R genetic variants present in the genome of PD patients that could confer increased IL-6R expression (data not shown). However, LRRK2 has been widely involved in vesicle trafficking and endocytosis thus it is possible to speculate that IL-6R membrane trafficking could be altered in PD leading to its accumulation.

Altogether, our results demonstrate that astrocytes are key contributors to PD-related neurodegeneration, both in L2-PD and ID-PD, by releasing high amounts of pro-inflammatory IL-6 that induce a hyperactivation of the STAT3 pathway in PD neurons leading to neurodegeneration. Thus, we propose the inhibition of astrocyte-dependent neurotoxic factor IL-6 to prevent, slow down or possibly halt PD pathogenesis.

## Methods

### iPSC-derived astrocyte generation and culture

The parental iPSC lines used in our studies were previously generated and fully characterized (33). Specifically, we used iPSC generated from three patients harboring the G2019S mutation on the *LRRK2* gene (L2-PD1: SP06, L2-PD 2: SP12, and L2-PD 3: SP13), and from three healthy age-matched controls (CTL 1: SP09, CTL 2: SP17 and CTL 3: SP11-FLAG). The isogenic line (L2-PD^corr^: SP13 wt/wt) was generated previously (16). The generation and/or use of human iPSCs in this work were approved by the Spanish competent authorities (Commission on Guarantees concerning the Donation and Use of Human Tissues and Cells of the Carlos III National Institute of Health). All procedures were done in accordance with institutional guidelines and the human iPSC lines have been (or are in the process of being) deposited at the Spanish National Stem Cell Bank, according to the Spanish legislation. Astrocytes were generated from patient-derived iPSC as previously described (16, 22). Briefly, iPSCs were cultured and differentiated into spherical neural masses (SNMs) that were pushed towards an astrocytic lineage. First, SNMs were grown in suspension for 28 days with Induction Medium (DMEM/F12, 1% N2 supplement, 0.1% B27 supplement (Life, 17504-044), 1% nonessential amino acids (NEAA), 1% penicillin/streptomycin (PenStrep), 1% Glutamax) supplemented with 20 ng/mL LIF (Sigma) and 20 ng/mL EGF (R&D Systems), and then for further 21 days with Propagation Medium (DMEM/F12, 1% N2 supplement, 0.1% B27 supplement, 1% NEAA, 1% PenStrep, 1% Glutamax) containing 20 ng/mL FGF-2 (PeproTech) and 20 ng/mL EGF (R&D Systems). Finally, SNMs were dissociated into a monolayer, plated on matrigel-coated plates and cultured for 14 more days in Propagation Medium and then for 14 more days in CNTF medium (Neurobasal, 1% Glutamax, 1% PenStrep, 1% NEAA, 0.2% B27 supplement, 10 ng/mL CNTF (Prospec Cyt-272). Experiments were performed with astrocytes growing on Thermanox™ plastic coverslips (Thermofisher) coated with matrigel in 24-well plates, or in 6-well plates for RNA analysis.

Astrocyte conditioned medium (ACM) was obtained as follows: 2,5×10^5^ astrocytes were plated per well on a matrigel coated 6-well plate in 2mL of CNTF medium. Each line was cultured for 14 days without changing the initial medium. At day 6, 1mL of fresh CNTF medium was added to each well. After the 14-day time-point, the medium was collected and frozen at −80°C.

### iPSC-derived vmDAn generation

Four different iPSC lines, three L2-PD (SP06, SP12 and SP13) and one CTL (SP11) were differentiated into dopaminergic neurons using the three-step system PSC Dopaminergic Neuron Differentiation Kit (GIBCO) according to the manufacturer’s protocol. Briefly, iPSCs were maintained in conditioned HES medium until they reached 70% of confluence and then passaged with EDTA (Invitrogen™) to Vitronectin-coated plates (10 μg/mL) with 3.0 x 10^4^ viable cells/cm^2^ in conditioned HES medium with 10 μM ROCK inhibitor (Y27632; RI). The day after, step 1 was started with changing the medium to Complete Floor Plate Specification Medium (Neurobasal, 20x Floor Plate Specification Supplement, 1% PenStrep) for up to 10 days, with a complete medium change every two days to move iPSC into midbrain-specified floor plate progenitor (FP) cells. On day 10, the second step of vmDAn differentiation started with the expansion of FP cells as adherent cultures in Complete Floor Plate Cell Expansion Medium (Floor Plate Cell Expansion Base Medium, 50x Floor Plate Cell Expansion Supplement, 1% PenStrep) on laminin-coated plates (3 μg/mL) for two passages on day 12 and 16 (FPp1 and FPp2 respectively) with Accutase (Invitrogen™) to a 1:4-6 split ratio onto laminin-coated plates with 5 μM of RI. Medium was fully changed every two days. On day 16, FPp2 can be either frozen with FP freezing media (90% Floor Plate Cell Expansion Medium and 10% DMSO) or expand the FP progenitors in sphere formation transferring cells to low-attachment plastic culture plates in Floor Plate Cell Expansion Medium, with a complete medium change every two days. Once the spheres are formed by day 21, they can be either frozen with FP freezing media or dissociated, following the Step 3 for vmDAn maturation. Dissociated cells were plated into Poly-D-lysine and Laminin double-coated plates (100 μg/mL-3 μg/mL) in Complete Maturation Medium (Neurobasal, 50x Dopaminergic Neuron Maturation Supplement, 1% NEAA, 1% PenStrep, 1% Glutamax), with 5 μM of RI to a density of 1.0 x 10^5^ cells/cm^2^. Medium was fully changed every two days until day 35, in which vmDAn were used for experiments.

### Cytokine Array

The HCYTOMAG-60K-15 Human Cytokine/Chemokine Multiplex Assay (Merck Millipore) was employed to simultaneously analyze TNF-α, IL-1β, IL-1a, IL-2, IL-4, IL-6, IL-10, Eotaxin, FLT-3L, GM-CSF, Rantes, MCP1, Fractalkine, and INFγ with Bead-Based Multiplex Assay using the Luminex^®^ technology. C3 and C1 were analyzed with HCMP2MAG-19K-02 Human Complement Magnetic Bead Panel 2 (Merck Millipore). Supernatants from non-stimulated CTL and L2-PD astrocytes cultured for 14 days were collected and stored at −80°C for long storage. CTL astrocytes stimulated with C1q, TNFα and IL-1α served as positive controls. Quality Controls (QC) and Human Cytokine Standard (STD) were prepared with deionized water by serial dilutions according to manufacturer’s instructions. Briefly, wells were filled with 200 μL of Wash Buffer (WB), sealed and mixed at RT for 10 minutes on a Thermomixer Comfort plate shaker at 500-800 rpm. The plate was firmly tap upside down on absorbent paper and 25 μL of each STD and QC with 25 μL of CNTF medium in duplicates. 25 μL of sample with 25 μL of Assay Buffer were added in duplicates. Finally, 25 μL of the mixed antibody-bead preparation was added to every well and incubated at 2-8°C O/N in a plate shaker. The day after, well contents were gently removed by firmly attaching plate to a handheld magnet and washed with WB for 3 times. Then, 25 μL of Detection Antibodies were added into each well and incubated at RT for 1 hour on a plate shaker. After that, 25 μL of Streptavidin-Phycoerythrin were added into each well and incubated at RT for 30 minutes on a plate shaker. Well contents were gently removed with a handheld magnet and washed with WB for 3 times. Antibody-beads were resuspended by adding 150 μL of Sheath Fluid to all wells and incubated at RT for 5 minutes on a plate shaker. Finally, plate was run on a multiplex system Luminex^®^ 200™ (Invitrogen™). Probe heights of every antibody were adjusted according to Luminex^®^ recommended protocols employing the xPONENT^®^ software (Luminex^®^). Mean Fluorescent Intensity of every analyte was analyzed using a 5-parameter logistic or spline curve-fitting method for calculating analyte concentrations in samples.

### Cell culture treatments

LRRK2-kinase inhibitor (1uM; Merck-Millipore, 438193) was added to astrocytes every 48h and fixed after one week. Anti-IL6R antibody (Tocilizumab) was kindly provided by Dr. Manel Juan Otero (Hospital Clinic, Barcelona). Tocilizumab (10ug/ml) and IL-6 (10ng/ml) was added to neurons together with ACM every 48h. Neurons were fixed after one week of treatment.

### Immunocytochemistry and Imaging analysis

Samples were fixed using 4% PFA and then washed with PBS. Samples were blocked and permeabilized with blocking solution (1xTBS, 3% Normal Donkey Serum, 0.01% Triton X-100) for 2 hours and subsequently incubated with the primary antibody for 48 hours at 4°C. Primary antibodies used include rabbit anti-AQP4 (Sigma, B104662), rabbit anti-C3 (Dako, A0063), guinea pig anti-GFAP (Synaptic Systems, 173 004), rabbit anti-GFAP (Dako, Z0334), mouse anti-IL6 (R&D systems, MAB206-SP), mouse anti-IL-6R (R&D systems, MAB2271), sheep anti-TH (Pel-Freez, P60101-0), rabbit anti-TH (Santa Cruz, sc-14007), mouse anti-Vimentin (Iowa, 40E-C). Samples were then washed three times with TBS, and incubated with secondary antibodies (1:250) for 2 hours at RT: Alexa Fluor 488 anti-Mouse IgG (Jackson 715-545-150), Cy3 anti-rabbit IgG (Jackson 711-165-152), DyLight 649 anti-Guinea pig IgG (Jackson 706-495-148), Alexa Fluor 647 anti-Sheep (Jackson 713-605-147), Cy™2 AffiniPure Donkey Anti-Rabbit IgG (H+L) (Jackson 711-225-152), Cy™3 AffiniPure Donkey Anti-Mouse IgG (H+L) (Jackson 715-165-151). Samples were washed with TBS followed with incubation with nuclear staining DAPI (Invitrogen, 1:5000) for 10 minutes, mounted with PVA:DABCO and stored at 4°C until imaged. Samples were imaged using an SPE or SP5 confocal microscope (Leica) and analyzed with FIJI is Just ImageJ™ using either the Cell Counter Plugin, Colocalization Plugin or Simple Neurite Tracer Plugin. Neurite length and number of terminals were analyzed for TH+ cells from an average of 5 images and ten neurons per image.

### Cellular transfection

2×10^4^ astrocytes seeded in 24-well plates (or 2,5×10^5^ in 6-well plates) were cotransfected after 7 days in culture with 1 ug pDEST51-LRRK2-G2019S, which was a gift from Mark Cookson (Addgene plasmid # 29401) or 1 ug of GFP expression plasmid as transfection control and collected after 48h. Transfection was done using Lipofectamine Stem Reagent (Invitrogen) following manufacturer’s instructions.

### RNA extraction and gene expression analysis

The isolation of total mRNA was performed with the RNeasy Micro Kit and treated with Rnase free Dnase I (Qiagen). 500ng were used to synthesize cDNA with the SuperScript III Reverse Transcriptase Synthesis Kit (Invitrogen). Quantitative RT-PCR analyzes were done in triplicate using 2ng/ul cDNA with Platinum SYBR Green qPCR Super Mix (Invitrogen) in an ABI Prism 7000 thermocycler (Applied Biosystems). All results were normalized to β-actin. Primers used are listed in **Supplemental Table 3**.

### Stranded mRNA library preparation and sequencing

Total RNA was assayed for quantity and quality using Qubit^®^ RNA HS Assay (Life Technologies) and RNA 6000 Nano Assay on a Bioanalyzer 2100. The RNASeq libraries were prepared from total RNA using the TruSeq^®^Stranded mRNA LT Sample Prep Kit (Illumina Inc., Rev.E, October 2013). Briefly, 500ng of total RNA was used as the input material and was enriched for the mRNA fraction using oligo-dT magnetic beads. The mRNA was fragmented in the presence of divalent metal cations and at high temperature (resulting RNA fragment size was 80-250 nt, with the major peak at 130nt). The second strand cDNA synthesis was performed in the presence of dUTP instead of dTTP, this allowed to achieve the strand specificity. The blunt-ended double stranded cDNA was 3’adenylated and Illumina indexed adapters were ligated. The ligation product was enriched with 15 PCR cycles and the final library was validated on an Agilent 2100 Bioanalyzer with the DNA 7500 assay. The libraries were sequenced on HiSeq2000 (Illumina, Inc) in paired-end mode with a read length of 2×76 bp using TruSeq SBS Kit v4. We generated over 30 million paired-end reads for each sample in a fraction of a sequencing v4 flow cell lane, following the manufacturer’s protocol. Image analysis, base calling and quality scoring of the run were processed using the manufacturer’s software Real Time Analysis (RTA 1.18.66.3) and followed by generation of FASTQ sequence files by CASAVA. The RNA-seq data have been deposited in Gene Expression Omnibus (GEO) of the National Center for Biotechnology Information and are accessible through GEO Series accession number GSE207713.

### Bioinformatics of RNA sequencing

The FASTQ files were then examined using FastQC (v0.11.8). Illumina adapters were removed with Trimmomatic (v0.36), considering default parameters but requiring a minimum average quality of 20 (34). Trimmed paired-end reads were aligned to the human reference genome (GRCh38_primary) using STAR v2.7.1a (35) with ENCODE parameters for long RNA. Annotated gene (gencode version 30) were quantified using the feature Counts function from the Bioconductor package Rsubread (v1.34.7) (36). Differential expression analysis was conducted using with DESeq2 package v1.24.0 (37). The method fits a generalized linear model (GLM) of the negative binomial distribution to estimate log2 Fold Change between groups. Then, it performs a Wald test to test the null hypothesis that there is no differential expression between L2-PD and CTL for each gene. Raw p-values were adjusted for multiple testing using the Benjamini and Hochberg False Discovery Rate (FDR). Any gene with an adjusted p-value lower than 0.05 and an absolute log2 Fold Change higher than 1 were considered to be differentially expressed. The Variance Stabilizing Transformation (VST) featured in DESeq2 package was applied to normalize the counts matrix. A Principal Component Analysis (PCA) was performed to explore characteristic patterns of samples and identify potential undesired effects.

Functional enrichment analysis was conducted using the Gene Set Enrichment Analysis (GSEA) approach. Genes considered for the analysis were pre-ranked by the -log10 (p-value) multiplied by the sign of the log2-fold change and gene sets derived from the Gene Ontology Biological Processes were downloaded from the Broad institute’s Molecular Signatures Database (MsigDB v3.1).

### Protein extraction

Cytoplasm and nucleus fractions were separated using Buffer A (10mM HEPES, 1.5mM MgCl2, 10mM KCl, 0.5mM DTT, 0.05% NP40 pH 7.9) followed by centrifugation at 3000 rpm for 10 min at 4°C. Pellets were homogenized in Laemmli buffer (4% SDS, 10% beta-mercaeptoethanol, 20% glycerol, 0.125M Tris-HCl pH 6.8, and 0.005% of bromophenol blue) and sonicated for 10 sec. Samples were then centrifuged at 15000 g for 20 min at 4°C. The resulting supernatant was normalized for protein using BCA kit (Pierce).

### Western blot (WB)

Cell extracts were boiled at 95°C for 5 minutes, followed by 10% SDS-PAGE, transferred to Nitrocellulose or PVDF membranes for 1.5 hours at 4°C and blocked with 5% not-fat milk in 0.1M Tris-buffered saline (pH= 7.4) for 1 hour. Membranes were incubated O/N at 4°C with primary antibodies diluted in TBS/ 3% BSA/ 0.1% TWEEN or TBS/ 5% milk/ 0.1% TWEEN. After incubation with peroxidase-tagged secondary antibodies (1:2500), membranes were revealed with ECL-plus chemiluminescence western blot kit (Amershan-Pharmacia Biotech). The following antibodies were used: mouse anti-beta actin (Affinity, T0022), mouse anti-IL6R (R&D Systems, MAB2271), rabbit anti-P-STAT3 (Tyr705) (Cell Signaling #9145, D3A7), rabbit anti-STAT3 (Cell Signaling #12640, D3Z2G), rabbit anti-a-tubulin (Sigma, T3526) and goat anti-LaminB (Santa Cruz, sc-6217). Films were scanned at 2,400 x 2,400 dpi (i800 MICROTEK high quality film scanner), and the densitometric analysis was performed using FIJI is Just ImageJ™. Other membranes were imaged using the ChemiTouch machine under the ‘Optimal exposure’ setting.

### Statistical Analysis

Box-and-whisker plots show median, 25^th^ and 75^th^ percentiles, minimum, and maximum values and were analyzed using GraphPad Prism 7.0 (Mac OS X). Statistical significance was assessed with a two-tailed unpaired t test or Mann-Whitney test for two experimental groups. For experiments with three or more groups, one-way ANOVA with Bonferroni’s multiple comparison test as post hoc was used. Results were considered significant when p < 0.05.

## Acknowledgments

The authors are indebted to the patients with PD who have participated in this study. The authors thank Giulia Carola for helping with some culture experiments. Research from the authors’ laboratories is supported by the Spanish Ministry of Economy and Competitiveness-MINECO (PID2019-108792GB-I00 supported by MCIN/AEI/10.13039/501100011033 and PDC2021-121051-I00), Instituto de Salud Carlos III-ISCIII/FEDER (Red de Terapia Celular - TerCel RD16/0011/0024 and RD16/0011/0025), AGAUR (2017-SGR-899 and 2017-SGR-1061), the ERC-2020-PoC and the Marató de TV3 Foundation (202012–32) and CERCA Program / Generalitat de Catalunya. MPE was partially supported by a Beatriu de Pinós fellowship from the Agency for Management of University and Research Grants (AGAUR) of the Government of Catalonia (2017 BP 00133). LB was the recipient of a pre-doctoral fellowship FPI (BES-2017-080579) from the Spanish Ministry of Economy and Competitiveness (MINECO). The authors declare that they have no competing interests.

## Author contributions

MPE and AC conceived and designed the experiments; MPE, LBA, IFC, ADD, YR, JLM, LM and LE performed the experiments, analyzed the data and performed statistical analysis; ET, MJE, IF, AR and MJO provided discussions; MPE and AC wrote the paper, while ET, IF and AR reviewed the manuscript; ADD edited the paper.

## References

1. W. Poewe et al., Parkinson disease. Nat Rev Dis Primers 3, 17013 (2017).

2. R. Balestrino, A. H. V. Schapira, Parkinson disease. Eur J Neurol 27, 27–42 (2020).

3. E. Tolosa, M. Vila, C. Klein, O. Rascol, LRRK2 in Parkinson disease: challenges of clinical trials. Nat Rev Neurol 16, 97–107 (2020).

4. P. Yang et al., String Vessel Formation is Increased in the Brain of Parkinson Disease. J Parkinsons Dis 5, 821–836 (2015).

5. A. Sommer et al., Th17 Lymphocytes Induce Neuronal Cell Death in a Human iPSC-Based Model of Parkinson’s Disease. Cell Stem Cell 23, 123–131.e126 (2018).

6. S. A. Liddelow et al., Neurotoxic reactive astrocytes are induced by activated microglia. Nature 541, 481–487 (2017).

7. X. Y. Qin, S. P. Zhang, C. Cao, Y. P. Loh, Y. Cheng, Aberrations in Peripheral Inflammatory Cytokine Levels in Parkinson Disease: A Systematic Review and Meta-analysis. JAMA Neurol 73, 1316–1324 (2016).

8. P. Garcia-Esparcia, F. Llorens, M. Carmona, I. Ferrer, Complex deregulation and expression of cytokines and mediators of the immune response in Parkinson’s disease brain is region dependent. Brain Pathol 24, 584–598 (2014).

9. H. Chen, E. J. O’Reilly, M. A. Schwarzschild, A. Ascherio, Peripheral inflammatory biomarkers and risk of Parkinson’s disease. Am J Epidemiol 167, 90–95 (2008).

10. J. R. Pereira et al., IL-6 serum levels are elevated in Parkinson’s disease patients with fatigue compared to patients without fatigue. J Neurol Sci 370, 153–156 (2016).

11. M. Dufek, I. Rektorova, V. Thon, J. Lokaj, I. Rektor, Interleukin-6 May Contribute to Mortality in Parkinson’s Disease Patients: A 4-Year Prospective Study. Parkinsons Dis 2015, 898192 (2015).

12. C. Escartin et al., Reactive astrocyte nomenclature, definitions, and future directions. Nat Neurosci 24, 312–325 (2021).

13. Y. Zhang et al., Purification and Characterization of Progenitor and Mature Human Astrocytes Reveals Transcriptional and Functional Differences with Mouse. Neuron 89, 37–53 (2016).

14. L. E. Clarke et al., Normal aging induces A1-like astrocyte reactivity. Proc Natl Acad Sci U S A 115, E1896–E1905 (2018).

15. S. P. Yun et al., Block of A1 astrocyte conversion by microglia is neuroprotective in models of Parkinson’s disease. Nat Med 24, 931–938 (2018).

16. A. di Domenico et al., Patient-Specific iPSC-Derived Astrocytes Contribute to Non-Cell-Autonomous Neurodegeneration in Parkinson’s Disease. Stem Cell Reports 12, 213–229 (2019).

17. A. de Rus Jacquet et al., The LRRK2 G2019S mutation alters astrocyte-to-neuron communication via extracellular vesicles and induces neuron atrophy in a human iPSC-derived model of Parkinson’s disease. Elife 10, (2021).

18. P. Ramos-Gonzalez et al., Astrocytic atrophy as a pathological feature of Parkinson’s disease with LRRK2 mutation. NPJ Parkinsons Dis 7, 31 (2021).

19. T. M. Sonninen et al., Metabolic alterations in Parkinson’s disease astrocytes. Sci Rep 10, 14474 (2020).

20. L. Streubel-Gallasch et al., Parkinson’s Disease-Associated LRRK2 Interferes with Astrocyte-Mediated Alpha-Synuclein Clearance. Mol Neurobiol 58, 3119–3140 (2021).

21. H. D. E. Booth et al., RNA sequencing reveals MMP2 and TGFB1 downregulation in LRRK2 G2019S Parkinson’s iPSC-derived astrocytes. Neurobiol Dis 129, 56–66 (2019).

22. A. Serio et al., Astrocyte pathology and the absence of non-cell autonomy in an induced pluripotent stem cell model of TDP-43 proteinopathy. Proc Natl Acad Sci U S A 110, 4697–4702 (2013).

23. K. A. Guttenplan et al., Neurotoxic reactive astrocytes induce cell death via saturated lipids. Nature 599, 102–107 (2021).

24. R. H. Reynolds et al., Moving beyond neurons: the role of cell type-specific gene regulation in Parkinson’s disease heritability. NPJ Parkinsons Dis 5, 6 (2019).

25. G. S. Nido et al., Common gene expression signatures in Parkinson’s disease are driven by changes in cell composition. Acta Neuropathol Commun 8, 55 (2020).

26. J. Bryois et al., Genetic identification of cell types underlying brain complex traits yields insights into the etiology of Parkinson’s disease. Nat Genet 52, 482–493 (2020).

27. E. C. Hirsch, S. Vyas, S. Hunot, Neuroinflammation in Parkinson’s disease. Parkinsonism Relat Disord 18 Suppl 1, S210–212 (2012).

28. R. M. Ransohoff, How neuroinflammation contributes to neurodegeneration. Science 353, 777–783 (2016).

29. K. Hartmann et al., Complement 3+-astrocytes are highly abundant in prion diseases, but their abolishment led to an accelerated disease course and early dysregulation of microglia. Acta Neuropathol Commun 7, 83 (2019).

30. L. Barbar et al., CD49f Is a Novel Marker of Functional and Reactive Human iPSC-Derived Astrocytes. Neuron, (2020).

31. R. Di Maio et al., LRRK2 activation in idiopathic Parkinson’s disease. Sci Transl Med 10, (2018).

32. S. Tanaka et al., Activation of microglia induces symptoms of Parkinson’s disease in wild-type, but not in IL-1 knockout mice. J Neuroinflammation 10, 143 (2013).

33. A. Sánchez-Danés et al., Disease-specific phenotypes in dopamine neurons from human iPS-based models of genetic and sporadic Parkinson’s disease. EMBO Mol Med 4, 380–395 (2012).

34. A. M. Bolger, M. Lohse, B. Usadel, Trimmomatic: a flexible trimmer for Illumina sequence data. Bioinformatics 30, 2114–2120 (2014).

35. A. Dobin et al., STAR: ultrafast universal RNA-seq aligner. Bioinformatics 29, 15–21 (2013).

36. B. Li, C. N. Dewey, RSEM: accurate transcript quantification from RNA-Seq data with or without a reference genome. BMC Bioinformatics 12, 323 (2011).

37. M. I. Love, W. Huber, S. Anders, Moderated estimation of fold change and dispersion for RNA-seq data with DESeq2. Genome Biol 15, 550 (2014).

