## Supplemental file for "Immunosuppressive tocilizumab prevents astrocyte induced neurotoxicity in hiPSC-LRRK2 Parkinson’s disease by targeting receptor interleukin-6"

## 1

**Supplemental Information contains:**

### Supplementary Figures:

#### **Figure S1.** L2-PD astrocytes overexpress genes related to reactivity**. Figure S2.** IL6/IL6R signaling is involved in DA neuronal degeneration **Figure S3.** iPSC-derived astrocyte characterization from idiopathic PD patients

**Supplementary Table:**

**Table S1.** List of iPSC lines used antibodies used in our studies.

**Table S2.** Results from GSEA analysis.

#### **Table S3.** List of primers used qRT-PCR used in our studies.

2

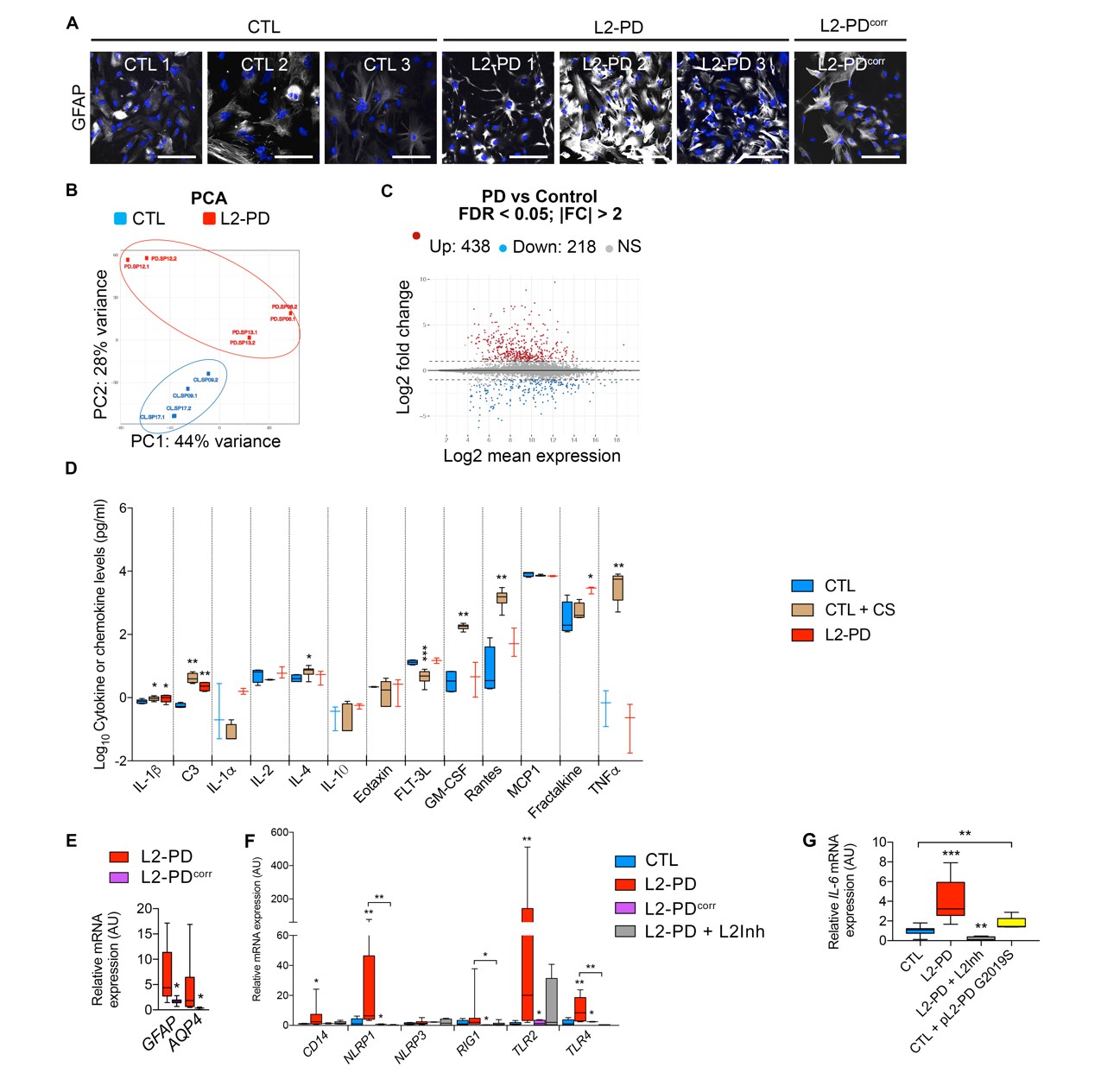

##### Supplemental Figure 1. L2-PD astrocytes overexpress genes related to reactivity.

(A) Representative ICC images of astrocytes from three CTL iPSC lines (CTL 1: SP09, CTL 2: SP17, CTL 3: SP11 FLAG), three L2-PD iPSC lines (L2-PD 1: SP06, L2-PD 2: SP12 and L2-PD 3: SP13) and one isogenic iPSC line (L2-PDcorr: SP13wt/wt) staining positive for DAPI (blue) and GFAP (white) after 2 weeks in culture. Scale bar: 100 μm.
(B) Representation of the first two dimensions from the Principal Component Analysis (PCA) showing the clustering patterns of CTL iPSC lines (blue dots; CTL 1: SP09 and CTL 2: SP17) and three L2-PD iPSC lines (red dots; L2-PD 1: SP06, L2-PD 2: SP12 and L2-PD 3: SP13). (**C**) RNA-seq MA plot showing expression changes (FDR <0.05; Fold change >2) of transcripts between CTL and L2-PD iPSC-derived astrocytes. Red and blue dots represent upregulated and downregulated genes, respectively. (**D**) Cytokine and chemokine levels released by CTL and L2-PD astrocyte conditioned medium (ACM) after 2 weeks of cell culture. CTL astrocytes were treated for 48h with C1q, TNFα and IL-1α as positive control. (**E**) Relative mRNA expression of *GFAP* and *AQP4* in L2-PD and L2-PDcorr. (**F**) Gene expression analysis using qRT-PCR showing several inflamed- dependent receptors in CTL, L2-PD, L2-PDcorr or L2-PD-treated astrocytes with LRRK2- kinase inhibitor (1uM). (**G**) Relative *IL-6* mRNA expression in CTL, L2-PD, L2-PD-treated astrocytes with LRRK2-kinase inhibitor (1uM) or CTL astrocytes after 48h transfection with a plasmid pLRRK2-G2019S to overexpress LRRK2G2019S.Box-and-whisker plots show median, 25th and 75th percentiles, minimum, and maximum values (n=3 experiments). One-way ANOVA Bonferroni as post-hoc. Student t-test or Mann-Whitney test for non-parametric conditions were when only two groups were compared. *p<0.05;

**p<0.01; ***p<0.001.

## 1

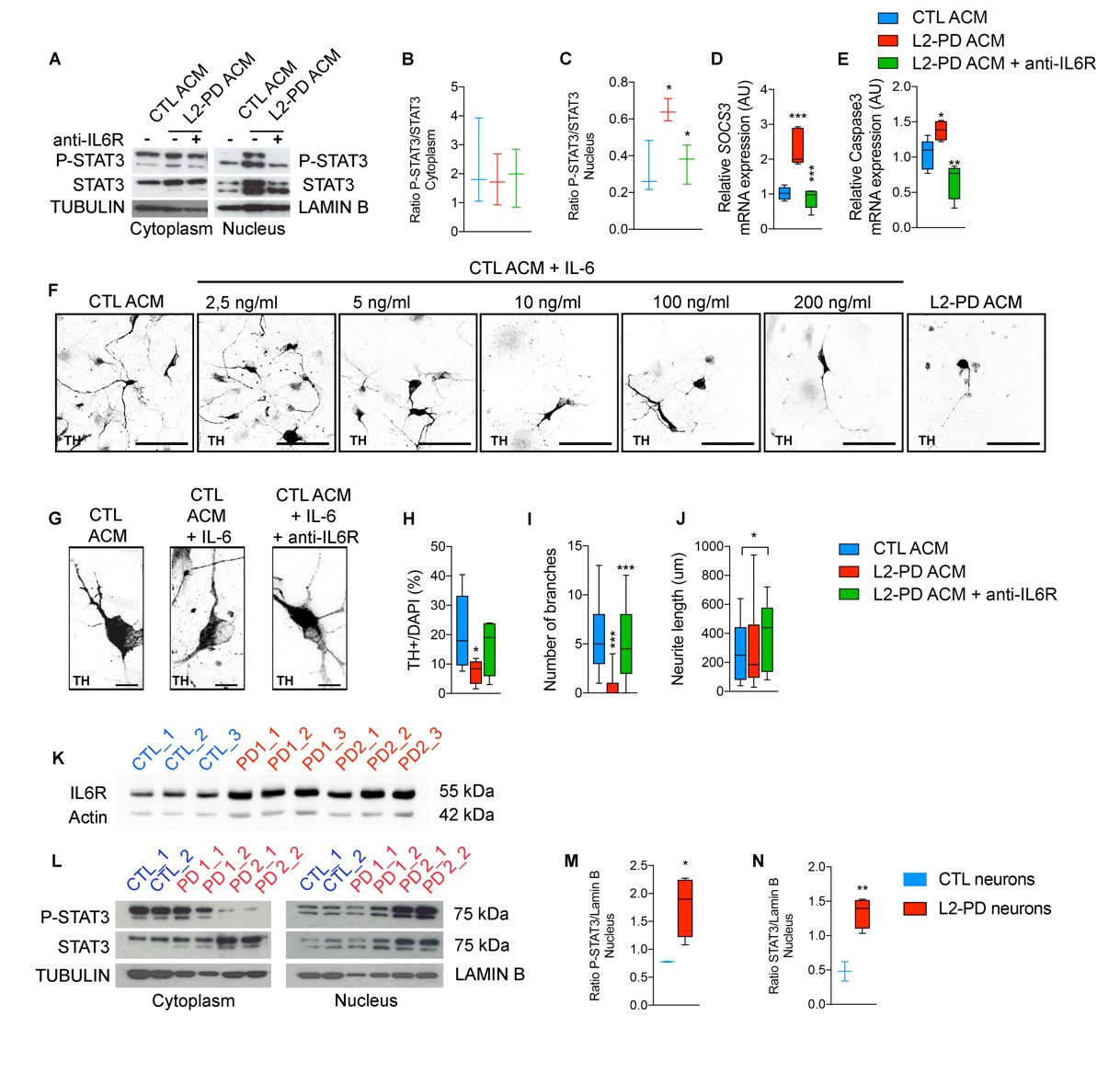

**Supplemental Figure 2. IL-6/IL-6R signaling is involved in DA neuronal degeneration.** (**A**) Western blot for STAT3 and p-STAT3 in the cytoplasm and nuclear fractions of iPSC-derived CTL neurons treated with CTL ACM or L2-PD ACM with anti- IL6R antibody Tocilizumab. (**B-C**) Quantification of protein p-STAT3/STAT ratio in the cytoplasm (**B**) or nucleus (**C**). (**D**) Relative *SOCS3* mRNA levels. (**E**) Relative *Caspase 3* mRNA levels. (**F**) Representative ICC images of DAn (TH+, black) from CTL SP11 neurons treated with either CTL ACM alone or with increasing IL-6 concentrations (2.5 ng/ml to 200ng/ml), or with L2-PD ACM during one week. (**G**) Representative ICC images of tyrosine hydroxylase (TH, black) from CTL SP11 neurons with CTL ACM treated with IL-6 (10ng/ml) and anti-IL6R Tocilizumab (10ug/ml) after one week in culture. Scale bar: 20 μm. (**H**) Percentage of TH+ cells respect to DAPI in CTL SP11 neurons when cultured with ACM plus IL-6 and anti-IL6R antibody. (**I**) Number of branches and (**J**) neurite length of CTL SP11 TH+ neurons. Box-and-whisker plots show median, 25th and 75th percentiles, minimum, and maximum values (n=3 experiments; 30 neurons per experiment per condition). (**K**) Western blot for IL-6R of iPSC-derived CTL neurons (SP11) and L2-PD neurons (SP12 and SP13). (**L**) Western blot for STAT3 and p-STAT3 in the cytoplasm and nuclear fractions of iPSC-derived CTL neurons (SP11) and L2-PD neurons (SP06 and SP13) treated with L2-PD ACM. (**M**) Quantification of protein p- STAT3/ LaminB ratio in the nucleus. (**N**) Quantification of protein STAT3/ LaminB ratio in the nucleus. Box-and-whisker plots show median, 25th and 75th percentiles, minimum, and maximum values (n=3 experiments). One-way ANOVA Bonferroni as post-hoc. Student t-test or Mann-Whitney test for non-parametric conditions were when only two groups were compared. *p<0.05; **p<0.01; ***p<0.001.

## 2

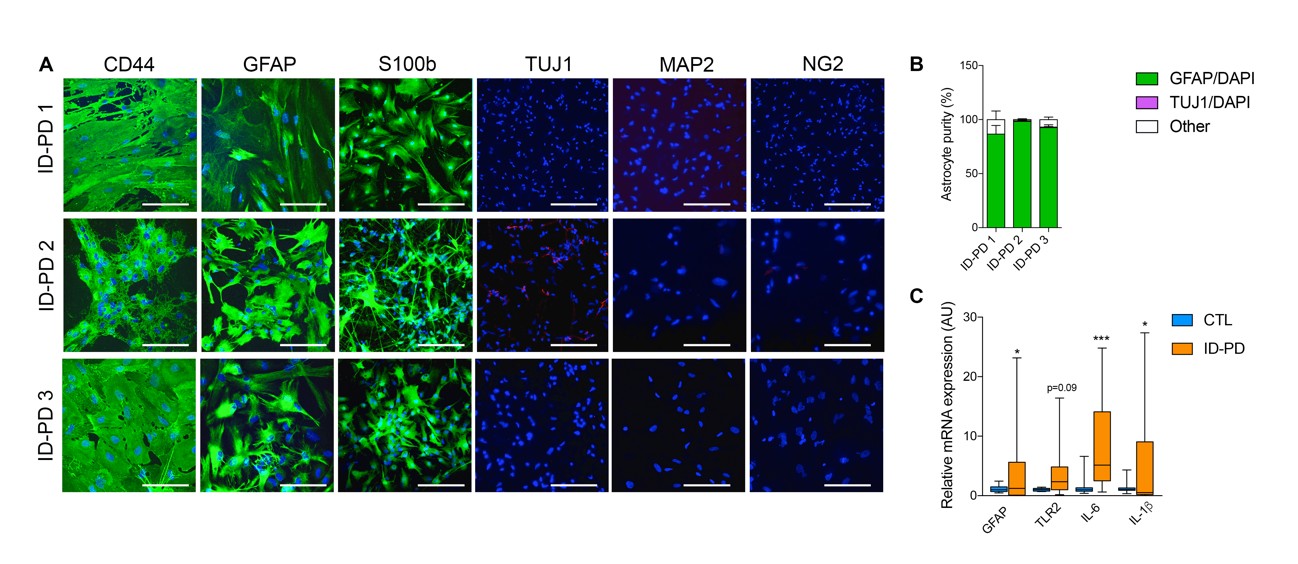

**Supplemental Figure 3. iPSC-derived astrocyte characterization from idiopathic PD patients.** (**A**) Representative ICC images of astrocytes derived from idiopathic PD iPSC lines (ID-PD 1: SP04, ID-PD 2: SP08 and ID-PD 3: SP16) staining positive for CD44 (astrocytic precursor marker), GFAP (general astrocytes), S100β (mature astrocytes), and negative for TUJ1 (immature neurons), MAP2 (mature neurons), and NG2 (oligodendrocytes) expression. Number of independent experiments per astrocyte line generated = 3. Scale bar, 100 μm. (**B**) Astrocyte cultures are composed of approximately 95% astrocytes, 4% neurons, and 1% other. (**C**) Relative *GFAP*, *TLR2*, *IL-6* and IL*-1ß* mRNA expression analysis using qRT-PCR. Box-and-whisker plots show median, 25th and 75th percentiles, minimum, and maximum values (n=3 experiments; Form factor, Mean GFAP and C3 intensity was performed from 30 astrocytes per experiment per condition). Student t-test or Mann-Whitney test for non-parametric conditions were when only two groups were compared. *p<0.05; **p<0.01; ***p<0.001.

## 3

##### Supplemental Table 1. Summary of iPSC lines used.

**Table S1. Summary of iPSC Lines Used**

Patient	Disease

**Code	Status	Sex Age at Biopsy Age of Onset Family history**

**Mutation**

| SP09 | Control | M | 66 | N/A | No | No |
| --- | --- | --- | --- | --- | --- | --- |
| SP17 | Control | M | 52 | N/A | No | No |
| SP11-FLAG | Control | F | 48 | N/A | No | No |
| SP11 | Control | F | 48 | N/A | No | No |
| SP13wtwt | PD LRRK2 Mutant Corrected | F | 68 | 57 | Yes | No |
| SP06 | PD LRRK2 Mutant | M | 44 | 33 | Yes | LRRK2G2019S |
| SP12 | PD LRRK2 Mutant | F | 63 | 49 | Yes | LRRK2G2019S |
| SP13 | PD LRRK2 Mutant | F | 68 | 57 | Yes | LRRK2G2019S |
| SP04 | Idiopathic PD | M | 46 | 40 | No | No |
| SP08 | Idiopathic PD | F | 66 | 60 | No | No |
| SP16 | Idiopathic PD | F | 51 | 48 | No | No |

##### Supplemental Table 3. Primers used for qRT-PCR.

| Primer ID | Forward (5’ – 3’) | Revers (5’ – 3’) |
| --- | --- | --- |
| *GFAP* | CCTCTCCCTGGCTCGAATG | GGAAGCGAACCTTCTCGATGTA |
| *AQP4* | GGTAAGTGTGGACCTTTGTGT | CAAAGCAAAGGGAGATGAGAAC |
| *VIMENTIN* | GCCCTAGACGAACTGGGTC | GGCTGCAACTGCCTAATGAG |
| *ACTIN* | AGGCCAACCGCGAGAAG | ACAGCCTGGATAGCAACGTACA |
| *TLR2* | ATCCTCCAATCAGGCTTCTCT | ACACCTCTGTAGGTCACTGTTG |
| *TLR4* | ATATTGACAGGAAACCCCATCCA | AGAGAGATTGAGTAGGGGCATTT |
| *CD14* | ACTTGCACTTTCCAGCTTGC | GCCCAGTCCAGGATTGTCAG |
| *NLRP1* | GGACTGACGATGACTTCTGG | ATCACAAAGCAGAGACCCG |
| *NLRP3* | GGAGAGACCTTTATGAGAAAGCAA | GCTGTCTTCCTGGCATATCACA |
| *RIG1* | GCAGGATTTGTAAAGCCCTGTT | CACTGATAATGAGGGCATCATTATATTT |
| *IL-6* | AATTCGGTACATCCTCGACGG | GGTTGTTTTCTGCCAGTGCC |
| *IL-6R* | CCCATCCCTGACGACAA | ACTGCTAACTGGCAGGAGAA |
| *IL-1b* | GCTGAGGAAGATGCTGGTTC | TCCATATCCTGTCCCTGGAG |
| *C3* | AAAAGGGGCGCAACAAGTTC | GATGCCTTCCGGGTTCTCAA |
| *SOCS3* | AGCAGCGATGGAATTACCTGGAAC | TCCAGCCCAATACCTGACACAGAA |
| *CASPASE3* | CAAACTTTTTCAGAGGGGATCG | GCATACTGTTTCAGCATGGCAC |
| *LCN* | CCCAGCCCCACCTCTGA | CTTCCCCTGGAATTGGTTGTC |
| *TIMP1* | TTGTGGGACCTGTGGAAGTA | CTGTTGTTGCTGTGGCTGAT |
| *SERPIN3N* | AGCAGTGGGGCTCTCAGTAA | ATAAGCAGACAGGGCCACAC |
| *PTX3* | CATCCAGTGAGACCAATGAG | GTAGCCGCCAGTTCACCATT |
| *CP* | ACGGCCATAGCTTCCAATACAA | AGTTGTATGCTTCCAGTCTTCT |
| *SERPING1* | AACCTGTGGCCCATTTCATT | TCTGGGGTACCAGGATCAC |
| *OSMR* | ACCTGCCACAGAGTACATGG | GCTCCAAGCTCACAATTCTCCA |
| *EMP2* | AGGGAATACATGGTTTACTCCA | AGAGAGATTGGCCAGCAAAA |

4

**Supplemental Table 2. GSEA analysis results.**

5

pathway	pval	padj	log2err	ES	NES	size

GOBP_MITOTIC_SISTER_CHROMATID_SEGREGATION	3,078E-22	3,855E-19	1,2210538	0,7671737	2,2396425	166

GOBP_REGULATION_OF_CHROMOSOME_SEGREGATION	2,136E-15	8,918E-13	1,007318	0,8102861	2,2193791	82

GOBP_MITOTIC_METAPHASE_PLATE_CONGRESSION	1,532E-13	4,112E-11	0,9436322	0,845586	2,1911043	53

GOBP_REGULATION_OF_CHROMOSOME_SEPARATION	4,779E-13	1,132E-10	0,921426	0,8120968	2,1748719	68

GOBP_CHROMOSOME_LOCALIZATION	6,367E-14	1,993E-11	0,9545416	0,7955293	2,1719211	79

GOBP_SISTER_CHROMATID_SEGREGATION	1,084E-21	1,018E-18	1,2039752	0,7360291	2,1717182	195

GOBP_METAPHASE_ANAPHASE_TRANSITION_OF_CELL_CYCLE	4,819E-13	1,132E-10	0,921426	0,8202851	2,1694632	63

GOBP_METAPHASE_PLATE_CONGRESSION	2,189E-12	4,059E-10	0,8986712	0,8148319	2,1681328	66

GOBP_NUCLEAR_CHROMOSOME_SEGREGATION	6,065E-25	1,139E-21	1,2951231	0,7155131	2,1509794	257

GOBP_CHROMOSOME_SEPARATION	1,179E-12	2,46E-10	0,9101197	0,7669668	2,1226403	90

GOBP_POSITIVE_REGULATION_OF_MITOTIC_CELL_CYCLE_PHASE_TRANSITION	1,181E-11	1,929E-09	0,8753251	0,765858	2,1056883	83

GOBP_CHROMOSOME_SEGREGATION	2,33E-25	8,754E-22	1,3030932	0,6936052	2,1000348	315

GOBP_POSITIVE_REGULATION_OF_CELL_CYCLE_PHASE_TRANSITION	4,571E-12	7,807E-10	0,887075	0,7397227	2,0672493	102

GOBP_ATTACHMENT_OF_SPINDLE_MICROTUBULES_TO_KINETOCHORE	1,198E-08	1,217E-06	0,7477397	0,8317321	2,0493964	37

GOBP_MICROTUBULE_CYTOSKELETON_ORGANIZATION_INVOLVED_IN_MITOSIS	1,078E-13	3,115E-11	0,9545416	0,7061935	2,0456864	149

GOBP_MEIOSIS_I_CELL_CYCLE_PROCESS	1,733E-10	2,412E-08	0,8266573	0,7276492	2,0177651	96

GOBP_NEGATIVE_REGULATION_OF_CHROMOSOME_ORGANIZATION	3,098E-09	3,423E-07	0,774939	0,7283235	2,0077995	84

GOBP_REGULATION_OF_MITOTIC_SISTER_CHROMATID_SEGREGATION	6,932E-08	5,788E-06	0,7049757	0,7926272	2,0039627	44

GOBP_MEIOTIC_CHROMOSOME_SEGREGATION	2,176E-08	2,096E-06	0,733762	0,7436346	1,9914852	70

GOBP_NEGATIVE_REGULATION_OF_METAPHASE_ANAPHASE_TRANSITION_OF_CELL_CYCLE	4,051E-08	3,623E-06	0,7195128	0,7977107	1,9897391	41

GOBP_MITOTIC_NUCLEAR_DIVISION	2,642E-17	1,241E-14	1,0768682	0,6592622	1,9860701	273

GOBP_CELL_CYCLE_DNA_REPLICATION	5,499E-08	4,805E-06	0,7195128	0,7953427	1,9838325	41

GOBP_DNA_UNWINDING_INVOLVED_IN_DNA_REPLICATION	4,15E-07	2,784E-05	0,6749629	0,8757032	1,9821406	22

GOBP_KINETOCHORE_ORGANIZATION	4,236E-06	0,000218	0,6105269	0,8592222	1,9784964	23

GOBP_POSITIVE_REGULATION_OF_MITOTIC_CELL_CYCLE	2,765E-10	3,463E-08	0,8140358	0,6995943	1,9705608	109

GOBP_DNA_DEPENDENT_DNA_REPLICATION	3,551E-11	5,558E-09	0,8513391	0,6782784	1,9692603	152

GOBP_ESTABLISHMENT_OF_MITOTIC_SPINDLE_LOCALIZATION	1,905E-07	1,379E-05	0,6901325	0,8089683	1,9680908	34

GOBP_MITOTIC_SPINDLE_ORGANIZATION	2,077E-10	2,786E-08	0,8266573	0,690184	1,9670603	123

GOBP_REGULATION_OF_DNA_DEPENDENT_DNA_REPLICATION	9,784E-08	7,821E-06	0,7049757	0,7603877	1,9485668	49

GOBP_CELL_CYCLE_G2_M_PHASE_TRANSITION	2,343E-10	3,035E-08	0,8140358	0,6716928	1,9420372	145

GOBP_NEGATIVE_REGULATION_OF_CELL_DIVISION	4,568E-07	3,011E-05	0,6749629	0,9197416	1,9399954	15

GOBP_POSITIVE_REGULATION_OF_CELL_CYCLE_G2_M_PHASE_TRANSITION	1,192E-06	6,892E-05	0,6435518	0,8140803	1,9387704	30

GOBP_REGULATION_OF_MITOTIC_CELL_CYCLE_PHASE_TRANSITION	6,569E-15	2,244E-12	0,9969862	0,6417351	1,9332688	273

GOBP_MITOTIC_CELL_CYCLE_PHASE_TRANSITION	1,902E-19	1,429E-16	1,1421912	0,6299161	1,9314866	386

GOBP_PROTEIN_LOCALIZATION_TO_CHROMOSOME_CENTROMERIC_REGION	1,16E-05	0,0005662	0,5933255	0,8263298	1,925652	25

GOBP_MEIOTIC_CELL_CYCLE_PROCESS	2,877E-10	3,486E-08	0,8140358	0,6631075	1,9245258	151

GOBP_FEMALE_MEIOTIC_NUCLEAR_DIVISION	1,909E-06	0,0001055	0,6272567	0,8105481	1,9239528	29

GOBP_DNA_REPLICATION_INITIATION	3,408E-06	0,0001778	0,6272567	0,7864125	1,9214879	35

GOBP_REGULATION_OF_CELL_CYCLE_G2_M_PHASE_TRANSITION	1,552E-08	1,535E-06	0,733762	0,6887587	1,9191471	100

GOBP_DNA_GEOMETRIC_CHANGE	4,043E-08	3,623E-06	0,7195128	0,6929967	1,9146392	94

GOBP_POSITIVE_REGULATION_OF_METAPHASE_ANAPHASE_TRANSITION_OF_CELL_CYCLE	3,208E-06	0,0001697	0,6272567	0,8878014	1,9061523	16

GOBP_ORGANELLE_FISSION	7,406E-19	4,637E-16	1,123915	0,6181279	1,9023164	424

GOBP_SPINDLE_LOCALIZATION	9,207E-07	5,579E-05	0,6594444	0,7378197	1,890734	49

GOBP_CYTOSKELETON_DEPENDENT_CYTOKINESIS	1,27E-07	9,545E-06	0,6901325	0,6836174	1,8887256	94

GOBP_MITOTIC_CYTOKINESIS	9,16E-07	5,579E-05	0,6594444	0,7045775	1,8868887	70

GOBP_MEIOTIC_CELL_CYCLE	1,047E-10	1,512E-08	0,8390889	0,6383454	1,8816716	192

GOBP_DNA_REPLICATION	5,722E-13	1,265E-10	0,921426	0,6233651	1,8757579	268

GOBP_REGULATION_OF_CELL_CYCLE_PHASE_TRANSITION	3,417E-15	1,284E-12	1,007318	0,6126978	1,8671586	353

GOBP_PROTEIN_LOCALIZATION_TO_CONDENSED_CHROMOSOME	0,0001078	0,0043083	0,5384341	0,8394284	1,8668369	19

GOBP_CELL_CYCLE_PHASE_TRANSITION	5,57E-18	2,99E-15	1,0959293	0,6041328	1,8664574	470

GOBP_REGULATION_OF_CHROMOSOME_ORGANIZATION	6,613E-11	9,938E-09	0,8390889	0,6307888	1,8642413	204

GOBP_POSITIVE_REGULATION_OF_CHROMOSOME_SEPARATION	4,278E-05	0,0018689	0,5573322	0,8436787	1,8612361	18

GOBP_REGULATION_OF_MITOTIC_NUCLEAR_DIVISION	1,908E-07	1,379E-05	0,6901325	0,6667008	1,8576851	100

GOBP_CENTROMERE_COMPLEX_ASSEMBLY	2,149E-05	0,0009967	0,5756103	0,7758878	1,8557266	31

GOBP_POSITIVE_REGULATION_OF_CELL_CYCLE_PROCESS	3,929E-10	4,474E-08	0,8140358	0,6275052	1,8518394	198

GOBP_SPINDLE_ORGANIZATION	3,665E-09	3,934E-07	0,774939	0,6270761	1,8461333	188

GOBP_ESTABLISHMENT_OF_SPINDLE_ORIENTATION	1,666E-05	0,0007925	0,5756103	0,7548816	1,8364343	33

GOBP_PEPTIDE_ANTIGEN_ASSEMBLY_WITH_MHC_PROTEIN_COMPLEX	4,782E-05	0,0020417	0,5573322	0,8525782	1,8305263	16

GOBP_KINETOCHORE_ASSEMBLY	0,0001423	0,0054549	0,5188481	0,8234157	1,8165339	18

GOBP_NEGATIVE_REGULATION_OF_NUCLEAR_DIVISION	3,949E-05	0,0017456	0,5573322	0,7010847	1,8118715	52

GOBP_CHROMOSOME_ORGANIZATION_INVOLVED_IN_MEIOTIC_CELL_CYCLE	3,49E-05	0,0015797	0,5573322	0,7027766	1,8102144	51

GOBP_HOMOLOGOUS_CHROMOSOME_SEGREGATION	0,0001298	0,0050293	0,5188481	0,7177696	1,8050174	43

GOBP_REGULATION_OF_NUCLEAR_DIVISION	1,013E-06	5,945E-05	0,6435518	0,6333404	1,8007878	117

GOBP_ATTACHMENT_OF_MITOTIC_SPINDLE_MICROTUBULES_TO_KINETOCHORE	0,0001676	0,0062968	0,5188481	0,8537293	1,8007569	15

GOBP_HUMORAL_IMMUNE_RESPONSE_MEDIATED_BY_CIRCULATING_IMMUNOGLOBULIN	0,0003393	0,011589	0,4984931	0,772572	1,8003764	25

GOBP_MITOTIC_CELL_CYCLE_CHECKPOINT_SIGNALING	6E-07	3,887E-05	0,6594444	0,6296926	1,7994967	127

GOBP_CELL_CYCLE_CHECKPOINT_SIGNALING	7,336E-08	5,991E-06	0,7049757	0,6155674	1,7989484	164

GOBP_SPINDLE_ASSEMBLY	8,665E-07	5,426E-05	0,6594444	0,6310831	1,7960689	122

GOBP_MEIOTIC_CHROMOSOME_SEPARATION	0,0002618	0,0093658	0,4984931	0,7936123	1,7944832	21

GOBP_DNA_SYNTHESIS_INVOLVED_IN_DNA_REPAIR	5,285E-05	0,0022309	0,5573322	0,7363058	1,7913145	34

GOBP_MICROTUBULE_BUNDLE_FORMATION	1,555E-06	8,85E-05	0,6435518	0,6364301	1,7893036	107

GOBP_AXONEME_ASSEMBLY	8,124E-06	0,0004016	0,5933255	0,6555436	1,7853156	78

GOBP_PROTEIN_LOCALIZATION_TO_CHROMOSOME	5,022E-06	0,0002516	0,6105269	0,6449239	1,781579	89

GOBP_DNA_CONFORMATION_CHANGE	3,63E-10	4,262E-08	0,8140358	0,5902561	1,7772419	275

GOBP_MITOTIC_SPINDLE_ASSEMBLY	3,645E-05	0,0016304	0,5573322	0,6628913	1,7752855	68

GOBP_MIDBODY_ABSCISSION	0,0003214	0,0110797	0,4984931	0,8143857	1,7746045	17

GOBP_HOMOLOGOUS_RECOMBINATION	9,673E-05	0,00395	0,5384341	0,6982192	1,7718253	47

GOBP_SISTER_CHROMATID_COHESION	6,648E-05	0,0027751	0,5384341	0,6745781	1,7712255	57

GOBP_RECOMBINATIONAL_REPAIR	6,864E-07	4,371E-05	0,6594444	0,6069692	1,7635138	154

GOBP_REGULATION_OF_SPINDLE_ORGANIZATION	0,0002338	0,0085295	0,5188481	0,704142	1,7631475	42

GOBP_CHROMOSOME_CONDENSATION	0,0001703	0,0063344	0,5188481	0,7157325	1,7585167	36

GOBP_CYTOKINESIS	4,103E-07	2,784E-05	0,6749629	0,6049032	1,7583324	155

GOBP_MITOTIC_CHROMOSOME_CONDENSATION	0,0012424	0,0356311	0,4550599	0,7898306	1,7565344	19

GOBP_REGULATION_OF_MITOTIC_CELL_CYCLE	2,269E-12	4,059E-10	0,8986712	0,5695634	1,7527803	415

GOBP_ANTIGEN_PROCESSING_AND_PRESENTATION_OF_EXOGENOUS_PEPTIDE_ANTIGEN_VIA_MHC_CLASS_II	0,0007541	0,0236099	0,4772708	0,7492466	1,7403669	24

GOBP_OVULATION_CYCLE_PROCESS	0,0003051	0,010711	0,4984931	0,7012406	1,73889	39

GOBP_CILIUM_MOVEMENT	2,879E-06	0,0001548	0,6272567 0,6065296	1,7347201	133

GOBP_CELL_PROLIFERATION_INVOLVED_IN_KIDNEY_DEVELOPMENT	0,0009946	0,0296578	0,4550599	0,7841458	1,7299008	18

GOBP_EXTRACELLULAR_TRANSPORT	0,0002955	0,0104734	0,4984931	0,6838614	1,7289751	44

GOBP_SYNAPSE_MATURATION	0,0029915	0,069808	0,4317077	0,7725232	1,7180437	19

GOBP_POSITIVE_REGULATION_OF_CELL_CYCLE	2,742E-08	2,575E-06	0,733762	0,5693643	1,7159581	271

GOBP_REGULATION_OF_UBIQUITIN_PROTEIN_LIGASE_ACTIVITY	0,0025799	0,0645328	0,4317077	0,7444755	1,7142736	23

GOBP_NEGATIVE_REGULATION_OF_MITOTIC_CELL_CYCLE_PHASE_TRANSITION	2,885E-06	0,0001548	0,6272567 0,5887641	1,7126926	156

GOBP_DNA_REPAIR	1,459E-12	2,886E-10	0,9101197	0,5518282	1,7115373	497

GOBP_REGULATION_OF_CYCLIN_DEPENDENT_PROTEIN_KINASE_ACTIVITY	7,549E-05	0,0031169	0,5384341	0,6166537	1,7070629	91

GOBP_MITOTIC_G2_DNA_DAMAGE_CHECKPOINT_SIGNALING	0,0004348	0,0145865	0,4984931	0,7010455	1,705532	34

GOBP_DOUBLE_STRAND_BREAK_REPAIR	1,135E-07	8,705E-06	0,7049757	0,5676685	1,7055026	256

GOBP_REGULATION_OF_DNA_REPLICATION	1,941E-05	0,0009116	0,5756103	0,5977792	1,7012856	122

GOBP_NEGATIVE_REGULATION_OF_CELL_CYCLE_G2_M_PHASE_TRANSITION	0,0003172	0,0110329	0,4984931	0,6466592	1,6995121	59

GOBP_POSTREPLICATION_REPAIR	0,00055	0,0178142	0,4772708	0,6978615	1,6977856	34

GOBP_COMPLEMENT_ACTIVATION	0,0010684	0,0316064	0,4550599	0,7192265	1,695435	28

GOBP_POSITIVE_REGULATION_OF_UBIQUITIN_PROTEIN_TRANSFERASE_ACTIVITY	0,0010803	0,0317098	0,4550599	0,7190282	1,6949673	28

GOBP_REGULATION_OF_CYTOKINESIS	0,0001207	0,0047734	0,5384341	0,6205462	1,6900033	78

GOBP_DNA_INTEGRITY_CHECKPOINT_SIGNALING	4,449E-05	0,0019211	0,5573322	0,5941503	1,6898863	118

GOBP_DNA_STRAND_ELONGATION_INVOLVED_IN_DNA_REPLICATION	0,0026346	0,0651207	0,4317077	0,7988466	1,6849937	15

GOBP_CYTOKINETIC_PROCESS	0,0008767	0,0267783	0,4772708	0,6793909	1,6847086	39

GOBP_MITOTIC_CYTOKINETIC_PROCESS	0,0040266	0,0845138	0,4070179	0,7313006	1,6839364	23

GOBP_NEGATIVE_REGULATION_OF_UBIQUITIN_PROTEIN_TRANSFERASE_ACTIVITY	0,0016071	0,0438636	0,4550599	0,7434926	1,682884	22

GOBP_REGULATION_OF_UBIQUITIN_PROTEIN_TRANSFERASE_ACTIVITY	0,000891	0,0269966	0,4772708	0,6493343	1,6781284	52

GOBP_INTERSTRAND_CROSS_LINK_REPAIR	0,00144	0,0403723	0,4550599	0,7008903	1,6763515	31

GOBP_EXIT_FROM_MITOSIS	0,0029389	0,0694439	0,4317077	0,71401	1,6760232	26

GOBP_NEGATIVE_REGULATION_OF_CELL_CYCLE_PHASE_TRANSITION	9,698E-07	5,783E-05	0,6435518	0,5644841	1,6755054	219

GOBP_MITOTIC_G2_M_TRANSITION_CHECKPOINT	0,0008668	0,0266924	0,4772708	0,6627191	1,6743787	46

GOBP_MICROTUBULE_BASED_MOVEMENT	1,077E-08	1,124E-06	0,7477397	0,5522648	1,6740052	332

GOBP_OVULATION_CYCLE	0,0006731	0,0216153	0,4772708	0,6368879	1,6722634	57

GOBP_RESPONSE_TO_INTERLEUKIN_6	0,0030345	0,0699439	0,4317077	0,7123962	1,6722351	26

GOBP_PROTEIN_DNA_COMPLEX_ASSEMBLY	1,47E-05	0,000708	0,5933255	0,5813554	1,6711181	140

GOBP_TRANSLESION_SYNTHESIS	0,0031422	0,0706911	0,4317077	0,7190667	1,6702644	24

GOBP_RENAL_SYSTEM_VASCULATURE_DEVELOPMENT	0,0034256	0,0757067	0,4317077	0,7166507	1,6700593	25

GOBP_NEGATIVE_REGULATION_OF_CELL_CYCLE_PROCESS	3,877E-07	2,697E-05	0,6749629	0,5550639	1,6686361	257

GOBP_NEGATIVE_REGULATION_OF_MITOTIC_CELL_CYCLE	1,669E-06	9,36E-05	0,6435518	0,5642298	1,6678048	202

GOBP_NEGATIVE_REGULATION_OF_DNA_DEPENDENT_DNA_REPLICATION	0,0033372	0,0741883	0,4317077	0,7894678	1,6652111	15

GOBP_DNA_REPLICATION_CHECKPOINT_SIGNALING	0,0030532	0,0699439	0,4317077	0,7739065	1,6616145	16

GOBP_DNA_RECOMBINATION	3,413E-07	2,419E-05	0,6749629	0,5467728	1,6482698	279

GOBP_ANAPHASE_PROMOTING_COMPLEX_DEPENDENT_CATABOLIC_PROCESS	0,0038971	0,0831905	0,4317077	0,7647806	1,6420207	16

GOBP_POSITIVE_REGULATION_OF_ALCOHOL_BIOSYNTHETIC_PROCESS	0,0110858	0,1819611	0,3807304	0,7361009	1,6370429	19

GOBP_DNA_PACKAGING	2,789E-05	0,001278	0,5756103 0,5555069	1,6351719	179

GOBP_OLIGODENDROCYTE_DEVELOPMENT	0,0027321	0,0666531	0,4317077	0,6624378	1,6322535	37

GOBP_NEGATIVE_REGULATION_OF_NEURON_DIFFERENTIATION	0,0007125	0,0224933	0,4772708	0,6181346	1,6306356	60

GOBP_NEGATIVE_REGULATION_OF_CELL_CYCLE	1,091E-07	8,539E-06	0,7049757	0,5372338	1,6288756	324

GOBP_MICROTUBULE_DEPOLYMERIZATION	0,0021671	0,0565415	0,4317077	0,6437692	1,6276119	44

GOBP_B_CELL_MEDIATED_IMMUNITY	0,0005291	0,0174381	0,4772708	0,586452	1,6259934	97

GOBP_SUBPALLIUM_DEVELOPMENT	0,0037232	0,0811002	0,4317077	0,7181865	1,6256041	22

GOBP_REGULATION_OF_SPINDLE_ASSEMBLY	0,0037855	0,081715	0,4317077	0,689353	1,625014	28

GOBP_HISTONE_PHOSPHORYLATION	0,0029731	0,069808	0,4317077	0,6482627	1,6232277	42

GOBP_REGULATION_OF_MITOTIC_CELL_CYCLE_SPINDLE_ASSEMBLY_CHECKPOINT	0,0053455	0,1056528	0,4070179	0,7447211	1,6228004	17

GOBP_DNA_STRAND_ELONGATION	0,0031057	0,0702906	0,4317077	0,6604909	1,6227908	36

GOBP_VASCULAR_ENDOTHELIAL_GROWTH_FACTOR_SIGNALING_PATHWAY	0,0031057	0,0702906	0,4317077	0,6603668	1,6224861	36

GOBP_POSITIVE_REGULATION_OF_CELL_CYCLE_G1_S_PHASE_TRANSITION	0,0016948	0,0455167	0,4550599	0,6330343	1,6222113	49

GOBP_CELLULAR_RESPONSE_TO_VASCULAR_ENDOTHELIAL_GROWTH_FACTOR_STIMULUS	0,0024163	0,0617564	0,4317077	0,6200823	1,6169822	55

GOBP_LIGAND_GATED_ION_CHANNEL_SIGNALING_PATHWAY	0,0130832	0,203114	0,3807304	0,7264939	1,6156774	19

GOBP_ANTIGEN_PROCESSING_AND_PRESENTATION_OF_EXOGENOUS_PEPTIDE_ANTIGEN	0,004419	0,0913887	0,4070179	0,6671749	1,610317	32

GOBP_CELLULAR_RESPONSE_TO_OXYGEN_RADICAL	0,0073734	0,1349072	0,4070179	0,7281109	1,6062824	18

GOBP_CILIUM_ORGANIZATION	6,337E-08	5,411E-06	0,7049757	0,5260341	1,6061549	373

GOBP_DNA_BIOSYNTHETIC_PROCESS	0,0001069	0,0043083	0,5384341	0,5451255	1,6016997	175

GOBP_DNA_DEPENDENT_DNA_REPLICATION_MAINTENANCE_OF_FIDELITY	0,0027025	0,0663603	0,4317077	0,6338774	1,6015093	46

GOBP_ANTIGEN_PROCESSING_AND_PRESENTATION_OF_EXOGENOUS_ANTIGEN	0,0043667	0,0911425	0,4070179	0,6498393	1,5966204	36

GOBP_AMINE_CATABOLIC_PROCESS	0,0088827	0,153084	0,3807304	0,7047657	1,593587	21

GOBP_REGULATION_OF_LIPID_KINASE_ACTIVITY	0,0040069	0,0845138	0,4070179	0,6157408	1,59131	52

GOBP_POSITIVE_REGULATION_OF_CYTOKINESIS	0,0078177	0,1412074	0,3807304	0,6687239	1,5873127	29

GOBP_REGULATION_OF_PHOSPHATIDYLINOSITOL_3_KINASE_ACTIVITY	0,0038063	0,081715	0,4317077	0,6354684	1,5850561	41

GOBP_DEVELOPMENT_OF_PRIMARY_FEMALE_SEXUAL_CHARACTERISTICS	0,0016972	0,0455167	0,4550599	0,5734909	1,5839744	92

GOBP_NEGATIVE_REGULATION_OF_DOUBLE_STRAND_BREAK_REPAIR_VIA_HOMOLOGOUS_RECOMBINATION	0,0111904	0,1827926	0,3807304	0,7179196	1,5837995	18

GOBP_FEMALE_GAMETE_GENERATION	0,0007041	0,0224167	0,4772708	0,556456	1,582181	117

GOBP_REGULATION_OF_MEIOTIC_CELL_CYCLE	0,0054912	0,1074506	0,4070179	0,6467729	1,5802981	35

GOBP_SIALYLATION	0,0214782	0,2849045	0,3524879	0,7102553	1,5795637	19

GOBP_IMMUNOGLOBULIN_PRODUCTION	0,0013809	0,0390091	0,4550599	0,5716415	1,5777513	93

GOBP_IMMUNOGLOBULIN_PRODUCTION_INVOLVED_IN_IMMUNOGLOBULIN_MEDIATED_IMMUNE_RESPONSE	0,0044271	0,0913887	0,4070179	0,6005204	1,576774	57

GOBP_ANTIGEN_PROCESSING_AND_PRESENTATION_OF_PEPTIDE_OR_POLYSACCHARIDE_ANTIGEN_VIA_MHC_CLASS_II	0,0092565	0,1587982	0,3807304	0,6636309	1,5752236	29

GOBP_MITOTIC_INTRA_S_DNA_DAMAGE_CHECKPOINT_SIGNALING	0,0101277	0,1721706	0,3807304	0,7317351	1,5710704	16

GOBP_PROTEIN_DNA_COMPLEX_SUBUNIT_ORGANIZATION	0,0001486	0,005641	0,5188481	0,5331752	1,5696859	188

GOBP_NEGATIVE_REGULATION_OF_DNA_RECOMBINATION	0,0086826	0,152433	0,3807304	0,636688	1,5688058	37

GOBP_EMBRYONIC_DIGESTIVE_TRACT_DEVELOPMENT	0,0150971	0,2250787	0,3807304	0,6810169	1,5681502	23

GOBP_REGULATION_OF_WNT_SIGNALING_PATHWAY_PLANAR_CELL_POLARITY_PATHWAY	0,0130755	0,203114	0,3807304	0,7431918	1,567602	15

GOBP_POSITIVE_REGULATION_OF_CHROMOSOME_ORGANIZATION	0,0028051	0,0671403	0,4317077	0,5667563	1,5673843	86

GOBP_AXONEMAL_DYNEIN_COMPLEX_ASSEMBLY	0,0070869	0,1305176	0,4070179	0,6414561	1,5673074	35

GOBP_PROTEIN_LOCALIZATION_TO_CILIUM	0,0037345	0,0811002	0,4317077	0,5853889	1,564786	67

GOBP_MITOTIC_DNA_INTEGRITY_CHECKPOINT_SIGNALING	0,0023443	0,0603253	0,4317077	0,5712038	1,564531	82

GOBP_NEGATIVE_REGULATION_OF_DNA_REPLICATION	0,0102272	0,1730785	0,3807304	0,6669482	1,5637297	27

GOBP_REGULATION_OF_GLIAL_CELL_PROLIFERATION	0,0158763	0,2339109	0,3524879	0,6788079	1,5630635	23

GOBP_MOTILE_CILIUM_ASSEMBLY	0,0070602	0,1305176	0,4070179	0,6135787	1,56262	48

GOBP_HOMOLOGOUS_CHROMOSOME_PAIRING_AT_MEIOSIS	0,00625	0,1198027	0,4070179	0,6415391	1,5607624	34

GOBP_MALE_GENITALIA_DEVELOPMENT	0,0137559	0,2100848	0,3807304	0,7142964	1,5565028	17

GOBP_POSITIVE_REGULATION_OF_G1_S_TRANSITION_OF_MITOTIC_CELL_CYCLE	0,0067968	0,1270419	0,4070179	0,6299467	1,5564186	38

GOBP_PROTEIN_LOCALIZATION_TO_CHROMATIN	0,0053712	0,1056528	0,4070179	0,6532161	1,5556647	30

GOBP_MITOTIC_SISTER_CHROMATID_COHESION	0,0141886	0,2140825	0,3807304	0,6625172	1,5551523	26

GOBP_INNER_DYNEIN_ARM_ASSEMBLY	0,0125342	0,2003162	0,3807304	0,7242353	1,554968	16

GOBP_REGULATION_OF_NEURON_MIGRATION	0,0073971	0,1349072	0,4070179	0,6257207	1,5516207	39

GOBP_MICROTUBULE_POLYMERIZATION_OR_DEPOLYMERIZATION	0,0020781	0,0545975	0,4317077	0,5476632	1,5513251	115

GOBP_INTRACILIARY_TRANSPORT	0,0075832	0,1376329	0,4070179	0,6252723	1,5505089	39

GOBP_NEURON_CELL_CELL_ADHESION	0,0159648	0,2342962	0,3524879	0,7101268	1,5474168	17

GOBP_POSITIVE_REGULATION_OF_DNA_BIOSYNTHETIC_PROCESS	0,005627	0,1095377	0,4070179	0,5770847	1,5454874	68

GOBP_REGULATION_OF_CELLULAR_RESPONSE_TO_VASCULAR_ENDOTHELIAL_GROWTH_FACTOR_STIMULUS	0,0168109	0,2421563	0,3524879	0,6850911	1,5393438	20

GOBP_CELL_CYCLE_G1_S_PHASE_TRANSITION	0,0002187	0,0080568	0,5188481	0,5201617	1,5386264	209

GOBP_INOSITOL_PHOSPHATE_BIOSYNTHETIC_PROCESS	0,0209468	0,2805513	0,3524879	0,6666682	1,5351099	23

GOBP_PROTEIN_MONOUBIQUITINATION	0,0033098	0,0740175	0,4317077	0,5609525	1,527705	78

GOBP_FEMALE_SEX_DIFFERENTIATION	0,0028057	0,0671403	0,4317077	0,5438555	1,5270214	104

GOBP_PIGMENT_CELL_DIFFERENTIATION	0,0140729	0,2140567	0,3807304	0,6312011	1,5234893	32

GOBP_NEGATIVE_REGULATION_OF_GLIOGENESIS	0,0079379	0,1422617	0,3807304	0,6395172	1,52304	30

GOBP_RESPONSE_TO_RADIATION	4,677E-06	0,0002375	0,6105269 0,4985182	1,5221399	373

GOBP_NUCLEOBASE_METABOLIC_PROCESS	0,011414	0,1856379	0,3807304	0,6221316	1,5200906	35

GOBP_MICROTUBULE_ORGANIZING_CENTER_ORGANIZATION	0,001378	0,0390091	0,4550599	0,5285453	1,5196233	139

GOBP_MELANOCYTE_DIFFERENTIATION	0,0236875	0,301674	0,3524879	0,6597559	1,5191933	23

GOBP_VIRAL_RNA_GENOME_REPLICATION	0,0082231	0,1451453	0,3807304	0,636118	1,5149447	30

GOBP_NEUROTRANSMITTER_METABOLIC_PROCESS	0,0217924	0,2850425	0,3524879	0,6741582	1,5147784	20

GOBP_CILIUM_OR_FLAGELLUM_DEPENDENT_CELL_MOTILITY	0,0025041	0,0634035	0,4317077	0,5433856	1,5121448	99

GOBP_CEREBRAL_CORTEX_NEURON_DIFFERENTIATION	0,0247012	0,3093416	0,3524879	0,6488207	1,5119904	25

GOBP_POSITIVE_REGULATION_OF_MUSCLE_CELL_APOPTOTIC_PROCESS	0,0218505	0,2850425	0,3524879	0,6680795	1,5106336	21

GOBP_REGULATION_OF_SISTER_CHROMATID_COHESION	0,0222514	0,28728	0,3524879	0,6678843	1,5101922	21

GOBP_OLFACTORY_LOBE_DEVELOPMENT	0,0129396	0,2025581	0,3807304	0,6201167	1,5086451	34

GOBP_NEGATIVE_REGULATION_OF_VASCULAR_PERMEABILITY	0,0209835	0,2805513	0,3524879	0,7146332	1,5073637	15

GOBP_REGULATION_OF_NEURAL_PRECURSOR_CELL_PROLIFERATION	0,0082073	0,1451453	0,3807304	0,5596079	1,5056653	72

GOBP_CEREBELLAR_PURKINJE_CELL_LAYER_MORPHOGENESIS	0,0335121	0,3758353	0,2878571	0,7006948	1,5044253	16

GOBP_POSITIVE_REGULATION_OF_DNA_REPLICATION	0,0166754	0,2421563	0,3524879	0,6081445	1,5025517	38

GOBP_REGULATION_OF_CELL_CYCLE_CHECKPOINT	0,0134215	0,2058148	0,3807304	0,6171284	1,5013159	33

GOBP_REGULATION_OF_MICROTUBULE_BASED_PROCESS	0,0002483	0,0089709	0,4984931	0,5047704	1,5011508	224

GOBP_REGULATION_OF_CELL_DIVISION	0,0016088	0,0438636	0,4550599	0,5177529	1,4998158	149

GOBP_CATECHOL_CONTAINING_COMPOUND_METABOLIC_PROCESS	0,0171113	0,2453711	0,3524879	0,6044543	1,4988858	39

GOBP_NON_MOTILE_CILIUM_ASSEMBLY	0,0133449	0,2058148	0,3807304	0,5698459	1,4962325	57

GOBP_CHONDROITIN_SULFATE_BIOSYNTHETIC_PROCESS	0,0375839	0,3973034	0,2712886	0,6865652	1,4960745	17

GOBP_GAMMA_AMINOBUTYRIC_ACID_TRANSPORT	0,036193	0,3918652	0,2765006	0,6967851	1,4960311	16

GOBP_ADRENAL_GLAND_DEVELOPMENT	0,0286281	0,3469545	0,3524879	0,643135	1,4938883	24

GOBP_NEGATIVE_REGULATION_OF_PROTEIN_ACETYLATION	0,0336927	0,3767368	0,2878571	0,6765725	1,4925838	18

GOBP_NEURON_MATURATION	0,0191248	0,2661185	0,3524879	0,603113	1,4901202	38

GOBP_GLOMERULUS_DEVELOPMENT	0,01341	0,2058148	0,3807304	0,5708671	1,4886442	55

GOBP_SPERM_MOTILITY	0,012831	0,2016989	0,3807304	0,5481666	1,4873569	77

GOBP_BRAIN_MORPHOGENESIS	0,0164249	0,2401101	0,3524879	0,610549	1,4853097	33

GOBP_REGULATION_OF_NEURON_DIFFERENTIATION	0,0025937	0,0645328	0,4317077	0,5100289	1,4843414	158

GOBP_EMBRYONIC_CAMERA_TYPE_EYE_MORPHOGENESIS	0,0332447	0,3750759	0,2878571	0,659171	1,4811034	20

GOBP_DOPAMINE_METABOLIC_PROCESS	0,0115598	0,1871984	0,3807304	0,6218346	1,4809282	30

GOBP_AXON_ENSHEATHMENT_IN_CENTRAL_NERVOUS_SYSTEM	0,0332447	0,3750759	0,2878571	0,6587629	1,4801865	20

GOBP_INOSITOL_PHOSPHATE_METABOLIC_PROCESS	0,0220988	0,2866357	0,3524879	0,5811688	1,4800808	48

GOBP_GLIAL_CELL_PROLIFERATION	0,0166964	0,2421563	0,3524879	0,6056981	1,4799377	35

GOBP_POSITIVE_REGULATION_OF_CYCLIN_DEPENDENT_PROTEIN_KINASE_ACTIVITY	0,0267662	0,3286298	0,3524879	0,6129969	1,4795511	32

GOBP_PYRIMIDINE_NUCLEOSIDE_METABOLIC_PROCESS	0,0442359	0,4412775	0,2489111	0,6890143	1,4793467	16

GOBP_REGULATION_OF_PLATELET_ACTIVATION	0,0186064	0,2618142	0,3524879	0,6079762	1,479109	34

GOBP_NEGATIVE_REGULATION_OF_LYMPHOCYTE_DIFFERENTIATION	0,0216124	0,2849045	0,3524879	0,5964464	1,4790284	39

GOBP_NEURAL_PRECURSOR_CELL_PROLIFERATION	0,0048223	0,0979318	0,4070179	0,5188366	1,4788821	125

GOBP_PYRIMIDINE_NUCLEOBASE_METABOLIC_PROCESS	0,0363881	0,3928455	0,2765006	0,6701454	1,4784049	18

GOBP_NEGATIVE_REGULATION_OF_PEPTIDYL_LYSINE_ACETYLATION	0,0455764	0,4458081	0,2450418	0,6874551	1,4759991	16

GOBP_POSITIVE_REGULATION_OF_MEIOTIC_CELL_CYCLE	0,042953	0,4326394	0,2529611	0,6771615	1,4755831	17

GOBP_SIGNAL_TRANSDUCTION_IN_RESPONSE_TO_DNA_DAMAGE	0,0036313	0,0797822	0,4317077	0,5048335	1,4753369	164

GOBP_CELL_DIFFERENTIATION_IN_SPINAL_CORD	0,0212053	0,2825113	0,3524879	0,6163611	1,4741791	31

GOBP_RESPONSE_TO_OXYGEN_RADICAL	0,0314306	0,367865	0,3217759	0,6511952	1,4739703	22

GOBP_RESPONSE_TO_LIGHT_STIMULUS	0,0005413	0,0176831	0,4772708	0,4894187	1,4726175	260

GOBP_POSITIVE_REGULATION_OF_TELOMERE_CAPPING	0,0456376	0,4458081	0,2450418	0,6755668	1,4721081	17

GOBP_POSITIVE_REGULATION_OF_VASCULAR_PERMEABILITY	0,0482574	0,4476616	0,2377938	0,6853135	1,471401	16

GOBP_PROTEIN_K11_LINKED_UBIQUITINATION	0,0331304	0,3750759	0,3217759	0,6194267	1,4702986	29

GOBP_NEGATIVE_REGULATION_OF_DNA_REPAIR	0,0209392	0,2805513	0,3524879	0,6034288	1,4679882	33

GOBP_ESTABLISHMENT_OF_ORGANELLE_LOCALIZATION	0,0001248	0,0048837	0,5188481	0,4802233	1,4673929	367

GOBP_SINGLE_STRANDED_VIRAL_RNA_REPLICATION_VIA_DOUBLE_STRANDED_DNA_INTERMEDIATE	0,0482574	0,4476616	0,2377938	0,6833738	1,4672364	16

GOBP_NEGATIVE_REGULATION_OF_AMYLOID_PRECURSOR_PROTEIN_CATABOLIC_PROCESS	0,0372836	0,3973034	0,2712886	0,6473243	1,4652086	22

GOBP_HUMORAL_IMMUNE_RESPONSE	0,0082289	0,1451453	0,3807304	0,5218099	1,4651225	104

GOBP_AMINE_METABOLIC_PROCESS	0,0106863	0,1776477	0,3807304	0,5301035	1,4643918	89

GOBP_REGULATION_OF_DNA_BIOSYNTHETIC_PROCESS	0,0050149	0,1012962	0,4070179	0,5184478	1,4640284	112

GOBP_BASE_EXCISION_REPAIR	0,029461	0,3536264	0,3524879	0,58461	1,4638434	42

GOBP_FACULTATIVE_HETEROCHROMATIN_ASSEMBLY	0,0148079	0,2216464	0,3807304	0,6146547	1,4638288	30

GOBP_REGULATION_OF_ALCOHOL_BIOSYNTHETIC_PROCESS	0,0225615	0,2884173	0,3524879	0,5865959	1,463153	41

GOBP_POSITIVE_REGULATION_OF_NEURAL_PRECURSOR_CELL_PROLIFERATION	0,0302213	0,3581749	0,3524879	0,5832362	1,4604036	42

GOBP_REGULATION_OF_PHOSPHOLIPASE_C_ACTIVITY	0,0332056	0,3750759	0,2820134	0,6129801	1,4598409	30

GOBP_CENTROSOME_DUPLICATION	0,0225698	0,2884173	0,3524879	0,5426001	1,4571826	71

GOBP_GLOMERULAR_MESANGIUM_DEVELOPMENT	0,0474255	0,4476616	0,24134	0,6904918	1,4564427	15

GOBP_CELLULAR_RESPONSE_TO_RADIATION	0,0065726	0,1247141	0,4070179	0,4994784	1,4546553	159

GOBP_NEURON_CELLULAR_HOMEOSTASIS	0,0347938	0,3856058	0,2765006	0,6126273	1,4541592	29

GOBP_CEREBELLAR_PURKINJE_CELL_LAYER_DEVELOPMENT	0,0401554	0,4179058	0,2572065	0,6242611	1,4500477	24

GOBP_REPLICATION_FORK_PROCESSING	0,0304858	0,3597322	0,3524879	0,5857007	1,4470995	38

GOBP_RESPONSE_TO_ARSENIC_CONTAINING_SUBSTANCE	0,0366162	0,3930483	0,2663507	0,5994612	1,4468807	32

GOBP_MITOTIC_RECOMBINATION	0,0375162	0,3973034	0,2663507	0,6203466	1,4456353	25

GOBP_EMBRYONIC_CAMERA_TYPE_EYE_DEVELOPMENT	0,0344828	0,3832891	0,2765006	0,606678	1,4448319	30

GOBP_DICARBOXYLIC_ACID_CATABOLIC_PROCESS	0,0493992	0,4544318	0,2343926	0,6496063	1,4446841	19

GOBP_REGULATION_OF_CAMP_MEDIATED_SIGNALING	0,045273	0,4458081	0,2450418	0,63781	1,4436732	22

GOBP_HINDLIMB_MORPHOGENESIS	0,0401035	0,4179058	0,2572065	0,6193653	1,4433486	25

GOBP_CELLULAR_RESPONSE_TO_IONIZING_RADIATION	0,0254132	0,31615	0,3524879	0,5371566	1,4425638	71

GOBP_NEGATIVE_REGULATION_OF_GLIAL_CELL_DIFFERENTIATION	0,0492021	0,4544318	0,2343926	0,641919	1,4423396	20

GOBP_INTERLEUKIN_1_MEDIATED_SIGNALING_PATHWAY	0,0466321	0,4458081	0,2377938	0,6208973	1,4422341	24

GOBP_DNA_LIGATION	0,0563003	0,4731993	0,2192503	0,6716153	1,4419904	16

GOBP_REGULATION_OF_SMOOTHENED_SIGNALING_PATHWAY	0,0154007	0,2286981	0,3807304	0,533313	1,4401828	75

GOBP_REGULATION_OF_MICROTUBULE_CYTOSKELETON_ORGANIZATION	0,0066301	0,1251723	0,4070179	0,4984022	1,4400559	146

GOBP_NEGATIVE_REGULATION_OF_REACTIVE_OXYGEN_SPECIES_METABOLIC_PROCESS	0,0353535	0,3895109	0,2712886	0,5917457	1,4396229	34

GOBP_RETINA_DEVELOPMENT_IN_CAMERA_TYPE_EYE	0,0126884	0,2011413	0,3807304	0,5050829	1,4395125	123

GOBP_POSITIVE_REGULATION_OF_POTASSIUM_ION_TRANSMEMBRANE_TRANSPORTER_ACTIVITY	0,0505319	0,4574017	0,2311267	0,6402327	1,4385505	20

GOBP_ZINC_ION_TRANSPORT	0,0546667	0,4717694	0,222056	0,6353345	1,436592	21

GOBP_SYNAPTONEMAL_COMPLEX_ORGANIZATION	0,0577181	0,4816976	0,2165428	0,6585385	1,4350022	17

GOBP_DNA_METHYLATION_DEPENDENT_HETEROCHROMATIN_ASSEMBLY	0,0479361	0,4476616	0,2377938	0,6332411	1,4333316	22

GOBP_RESPONSE_TO_PROSTAGLANDIN_E	0,0590604	0,4845135	0,2139279	0,6575955	1,4329474	17

GOBP_OOGENESIS	0,0263859	0,3250221	0,3524879	0,5309047	1,432685	74

GOBP_POSITIVE_REGULATION_OF_PROTEIN_MATURATION	0,0492676	0,4544318	0,2343926	0,6329279	1,4326224	22

GOBP_CELLULAR_RESPONSE_TO_ALKALOID	0,0481771	0,4476616	0,2343926	0,6109881	1,4325254	27

GOBP_DNA_REPLICATION_INDEPENDENT_CHROMATIN_ORGANIZATION	0,0573333	0,4797357	0,2165428	0,6333932	1,4322024	21

GOBP_REGULATION_OF_TELOMERE_CAPPING	0,0464516	0,4458081	0,2377938	0,6095454	1,4308094	26

GOBP_NEUROTRANSMITTER_UPTAKE	0,035309	0,3895109	0,2712886	0,5805059	1,4303726	37

GOBP_CHROMATIN_ASSEMBLY_OR_DISASSEMBLY	0,0067266	0,1263599	0,4070179	0,4895054	1,4300989	165

GOBP_RESPONSE_TO_X_RAY	0,0494792	0,4544318	0,2311267	0,6097035	1,4295135	27

GOBP_RESPONSE_TO_UV	0,0064495	0,1229987	0,4070179	0,4990499	1,4291112	135

GOBP_SPECIFICATION_OF_SYMMETRY	0,0106623	0,1776477	0,3807304	0,5073222	1,4289845	109

GOBP_POSITIVE_REGULATION_OF_CARDIAC_MUSCLE_CELL_PROLIFERATION	0,0545213	0,4717694	0,222056	0,6357764	1,4285377	20

GOBP_REGULATION_OF_CILIUM_MOVEMENT	0,0533854	0,4664396	0,222056	0,6201499	1,4279942	23

GOBP_INORGANIC_ION_IMPORT_ACROSS_PLASMA_MEMBRANE	0,0221252	0,2866357	0,3524879	0,5222162	1,42751	80

GOBP_RESPONSE_TO_MANGANESE_ION	0,0604027	0,4901358	0,2114002	0,654803	1,4268624	17

GOBP_DIGESTIVE_SYSTEM_DEVELOPMENT	0,0179596	0,2546197	0,3524879	0,5116137	1,4255528	100

GOBP_GANGLIOSIDE_BIOSYNTHETIC_PROCESS	0,0582656	0,4821669	0,2165428	0,6755374	1,4248997	15

GOBP_SYNAPTIC_MEMBRANE_ADHESION	0,0546875	0,4717694	0,2192503	0,6187925	1,4248686	23

GOBP_NEGATIVE_REGULATION_OF_DNA_METABOLIC_PROCESS	0,0145357	0,2184426	0,3807304	0,5118319	1,4243365	99

GOBP_POSITIVE_REGULATION_OF_LIPID_KINASE_ACTIVITY	0,0478655	0,4476616	0,2343926	0,6111133	1,4241185	25

GOBP_FEEDING_BEHAVIOR	0,024454	0,3072694	0,3524879	0,5398234	1,4240511	60

GOBP_PROTEIN_HETEROOLIGOMERIZATION	0,0559896	0,4725528	0,2165428	0,6179849	1,423009	23

GOBP_ENDOCARDIAL_CUSHION_FORMATION	0,0507813	0,4575184	0,2279872	0,6033978	1,4223915	28

GOBP_CELLULAR_RESPONSE_TO_REACTIVE_NITROGEN_SPECIES	0,0630872	0,5000396	0,2065879	0,6525684	1,4219931	17

GOBP_ESTROGEN_METABOLIC_PROCESS	0,0630872	0,5000396	0,2065879	0,6522091	1,4212102	17

GOBP_RESPONSE_TO_COPPER_ION	0,045977	0,4458081	0,2377938	0,5966323	1,4209077	30

GOBP_DNA_DAMAGE_RESPONSE_SIGNAL_TRANSDUCTION_RESULTING_IN_TRANSCRIPTION	0,0644295	0,501163	0,2042948	0,651407	1,4194622	17

GOBP_FIBROBLAST_PROLIFERATION	0,0318108	0,3710617	0,3217759	0,5283236	1,4188422	71

GOBP_ASPARTATE_FAMILY_AMINO_ACID_CATABOLIC_PROCESS	0,0670241	0,5170629	0,1999152	0,6607054	1,4185663	16

GOBP_NEGATIVE_REGULATION_OF_PLATELET_ACTIVATION	0,0623306	0,4982471	0,208955	0,6720393	1,4175212	15

GOBP_MUSCLE_HYPERTROPHY_IN_RESPONSE_TO_STRESS	0,0599201	0,4890772	0,2114002	0,6259894	1,4169174	22

GOBP_DEVELOPMENT_OF_PRIMARY_SEXUAL_CHARACTERISTICS	0,0046573	0,0956138	0,4070179	0,4819476	1,4165931	182

GOBP_REGULATION_OF_RESPONSE_TO_EXTRACELLULAR_STIMULUS	0,0600801	0,4890772	0,2114002	0,6355601	1,4134462	19

GOBP_APPENDAGE_MORPHOGENESIS	0,0128117	0,2016989	0,3807304	0,49695	1,4129867	117

GOBP_CHROMATIN_REMODELING	0,0070155	0,1304821	0,4070179	0,4784458	1,4122255	199

GOBP_POSITIVE_REGULATION_OF_AMYLOID_PRECURSOR_PROTEIN_CATABOLIC_PROCESS	0,064	0,5002749	0,2042948	0,6243296	1,4117082	21

GOBP_REGULATION_OF_DNA_REPAIR	0,0118542	0,1911419	0,3807304	0,4880319	1,4100924	146

GOBP_EMBRYONIC_APPENDAGE_MORPHOGENESIS	0,0207434	0,2805513	0,3524879	0,5084168	1,4098353	96

GOBP_DEOXYRIBONUCLEOTIDE_BIOSYNTHETIC_PROCESS	0,0677507	0,5171182	0,1999152	0,6683577	1,4097557	15

GOBP_REGULATION_OF_NON_CANONICAL_WNT_SIGNALING_PATHWAY	0,0620957	0,4981249	0,2042948	0,6045668	1,4088627	25

GOBP_NEGATIVE_REGULATION_OF_CELL_AGING	0,0714286	0,5367143	0,1938133	0,6375989	1,4066044	18

GOBP_GLIOGENESIS	0,0028801	0,0684841	0,4317077	0,4699428	1,4059532	244

GOBP_POSITIVE_REGULATION_OF_CILIUM_ASSEMBLY	0,0686528	0,5210682	0,1938133	0,6051529	1,4056626	24

GOBP_POSITIVE_REGULATION_OF_FIBROBLAST_PROLIFERATION	0,0404551	0,4184363	0,2529611	0,5612682	1,4053965	42

GOBP_GANGLIOSIDE_METABOLIC_PROCESS	0,0664063	0,5144088	0,197822	0,609522	1,4035217	23

GOBP_REGULATION_OF_DOUBLE_STRAND_BREAK_REPAIR	0,025227	0,314877	0,3524879	0,499481	1,4022452	105

GOBP_ANTIGEN_PROCESSING_AND_PRESENTATION_OF_PEPTIDE_ANTIGEN	0,0418972	0,427739	0,3217759	0,540926	1,4016614	53

GOBP_GLIAL_CELL_DIFFERENTIATION	0,0088427	0,153084	0,3807304	0,4757635	1,400664	188

GOBP_POSITIVE_REGULATION_OF_DNA_METABOLIC_PROCESS	0,0058018	0,1123568	0,4070179	0,4724208	1,3996835	213

GOBP_RESPONSE_TO_IONIZING_RADIATION	0,0124535	0,1999487	0,3807304	0,4888309	1,3995287	136

GOBP_NUCLEOSOME_ASSEMBLY	0,0357995	0,3909849	0,2616635	0,5234781	1,399294	67

GOBP_HISTONE_H3_K9_METHYLATION	0,0505051	0,4574017	0,2249661	0,5741913	1,396916	34

GOBP_GLUTAMATE_RECEPTOR_SIGNALING_PATHWAY	0,0393401	0,4117013	0,2572065	0,5526987	1,396409	46

GOBP_EMBRYONIC_HINDLIMB_MORPHOGENESIS	0,0638298	0,5002749	0,2042948	0,6214527	1,3963534	20

GOBP_REGULATION_OF_MICROTUBULE_POLYMERIZATION_OR_DEPOLYMERIZATION	0,0305581	0,3597322	0,3524879	0,5097344	1,3961659	82

GOBP_REGULATION_OF_PLATELET_AGGREGATION	0,0732357	0,5437677	0,1900233	0,6166856	1,3958582	22

GOBP_POSITIVE_REGULATION_OF_DNA_REPAIR	0,0239383	0,3028159	0,3524879	0,5039416	1,3946974	90

GOBP_TRNA_WOBBLE_BASE_MODIFICATION	0,0791946	0,5645811	0,1830239	0,6398892	1,3943641	17

GOBP_REGULATION_OF_VASCULAR_PERMEABILITY	0,0453972	0,4458081	0,2377938	0,5669221	1,3928974	36

GOBP_LOCOMOTORY_BEHAVIOR	0,0110911	0,1819611	0,3807304	0,4777291	1,3928005	161

GOBP_DIGESTIVE_TRACT_MORPHOGENESIS	0,0517677	0,4586792	0,222056	0,5719683	1,3915077	34

GOBP_POSITIVE_REGULATION_OF_DOUBLE_STRAND_BREAK_REPAIR	0,0361011	0,3918652	0,2616635	0,526351	1,3915022	61

GOBP_PROTEIN_LOCALIZATION_TO_CILIARY_MEMBRANE	0,0791946	0,5645811	0,1830239	0,6378353	1,3898886	17

GOBP_CELLULAR_PROCESS_INVOLVED_IN_REPRODUCTION_IN_MULTICELLULAR_ORGANISM	0,0022522	0,058354	0,4317077	0,4611781	1,3895606	274

GOBP_REGULATION_OF_RESPIRATORY_GASEOUS_EXCHANGE	0,0791946	0,5645811	0,1830239	0,6376511	1,3894872	17

GOBP_MRNA_SPLICE_SITE_SELECTION	0,0618687	0,4981249	0,2020717	0,5755746	1,3892273	32

GOBP_HEMATOPOIETIC_PROGENITOR_CELL_DIFFERENTIATION	0,0289518	0,3486277	0,3524879	0,4936183	1,3857862	105

GOBP_NEGATIVE_REGULATION_OF_ORGANELLE_ORGANIZATION	0,0017281	0,0457213	0,4550599	0,4577032	1,3855373	318

GOBP_REGULATION_OF_AMYLOID_PRECURSOR_PROTEIN_CATABOLIC_PROCESS	0,0515723	0,4583496	0,222056	0,5580783	1,3854191	40

GOBP_REGULATION_OF_CELL_CYCLE_G1_S_PHASE_TRANSITION	0,0198386	0,2700491	0,3524879	0,4790706	1,3851167	145

GOBP_POSITIVE_REGULATION_OF_ORGANELLE_ASSEMBLY	0,0329412	0,3750759	0,2712886	0,5126357	1,3843449	75

GOBP_POSITIVE_REGULATION_OF_SYNAPTIC_TRANSMISSION_GLUTAMATERGIC	0,0745672	0,5503912	0,1882041	0,611329	1,3837339	22

GOBP_AMMONIUM_ION_METABOLIC_PROCESS	0,0832215	0,5800799	0,1782199	0,6344243	1,3824559	17

GOBP_AMINO_ACID_IMPORT	0,0481481	0,4476616	0,2279872	0,5394151	1,3823033	49

GOBP_BLOOD_VESSEL_ENDOTHELIAL_CELL_PROLIFERATION_INVOLVED_IN_SPROUTING_ANGIOGENESIS	0,0758988	0,5563902	0,1864326	0,6104343	1,3817087	22

GOBP_REGULATION_OF_GLIOGENESIS	0,0324173	0,3747434	0,3217759	0,5049281	1,3813697	81

GOBP_NEGATIVE_REGULATION_OF_INFLAMMATORY_RESPONSE	0,0255435	0,3167219	0,3524879	0,4978754	1,3806166	95

GOBP_POSITIVE_REGULATION_OF_LYASE_ACTIVITY	0,0758808	0,5563902	0,1882041	0,6545287	1,3805864	15

GOBP_MAMMARY_GLAND_EPITHELIAL_CELL_PROLIFERATION	0,0758988	0,5563902	0,1864326	0,6098492	1,3803843	22

GOBP_POSITIVE_REGULATION_OF_TELOMERE_MAINTENANCE_VIA_TELOMERE_LENGTHENING	0,0554855	0,4725528	0,2139279	0,5600573	1,3799869	37

GOBP_SEX_DIFFERENTIATION	0,0088239	0,153084	0,3807304	0,4654703	1,3797057	215

GOBP_PEPTIDYL_TYROSINE_DEPHOSPHORYLATION	0,032	0,3710617	0,2712886	0,4983026	1,3794354	91

GOBP_VENTRICULAR_SEPTUM_MORPHOGENESIS	0,055	0,4717694	0,2139279	0,5562235	1,3792861	39

GOBP_AMYLOID_BETA_FORMATION	0,0530973	0,465004	0,2192503	0,5505932	1,3786665	42

GOBP_CARDIAC_SEPTUM_MORPHOGENESIS	0,0420168	0,4277971	0,24134	0,5211661	1,378363	63

GOBP_POSITIVE_REGULATION_OF_PROTEIN_MODIFICATION_BY_SMALL_PROTEIN_CONJUGATION_OR_REMOVAL	0,0187934	0,2624788	0,3524879	0,4811551	1,377994	130

GOBP_CENTRIOLE_ASSEMBLY	0,0641509	0,5002749	0,197822	0,5447191	1,3771881	44

GOBP_POSITIVE_REGULATION_OF_NERVOUS_SYSTEM_DEVELOPMENT	0,0052667	0,1046935	0,4070179	0,4614493	1,3766603	236

GOBP_REGULATION_OF_NEUROTRANSMITTER_LEVELS	0,0088805	0,153084	0,3807304	0,4681626	1,3764432	183

GOBP_RESPONSE_TO_NITRIC_OXIDE	0,0885906	0,5922329	0,1723243	0,63144	1,3759528	17

GOBP_NEUROBLAST_PROLIFERATION	0,0442804	0,4412775	0,2377938	0,5340432	1,3755905	51

GOBP_COCHLEA_DEVELOPMENT	0,0567465	0,4758855	0,2114002	0,5581907	1,3753875	37

GOBP_PYRIMIDINE_CONTAINING_COMPOUND_METABOLIC_PROCESS	0,0376471	0,3973034	0,2529611	0,5091175	1,3748442	75

GOBP_POSITIVE_REGULATION_OF_NON_CANONICAL_WNT_SIGNALING_PATHWAY	0,0813008	0,5719984	0,1813831	0,6516072	1,3744242	15

GOBP_POSITIVE_CHEMOTAXIS	0,0556258	0,4725528	0,2139279	0,5488562	1,3743172	42

GOBP_CELLULAR_RESPONSE_TO_ARSENIC_CONTAINING_SUBSTANCE	0,084	0,5801999	0,1766943	0,6073309	1,3732715	21

GOBP_ALPHA_AMINO_ACID_CATABOLIC_PROCESS	0,0361727	0,3918652	0,2572065	0,5064503	1,3724849	76

GOBP_POSITIVE_REGULATION_OF_NEUROGENESIS	0,0098091	0,167512	0,3807304	0,4647848	1,371633	198

GOBP_NEURAL_RETINA_DEVELOPMENT	0,0476773	0,4476616	0,2279872	0,5306522	1,3714086	52

GOBP_CELLULAR_RESPONSE_TO_LEPTIN_STIMULUS	0,0840108	0,5801999	0,1782199	0,6499568	1,3709429	15

GOBP_MALE_MEIOTIC_NUCLEAR_DIVISION	0,0668348	0,5166633	0,1938133	0,5609349	1,3705652	35

GOBP_CELLULAR_RESPONSE_TO_REACTIVE_OXYGEN_SPECIES	0,032909	0,3750759	0,3217759	0,4814139	1,3703514	121

GOBP_POSITIVE_REGULATION_OF_PHOSPHOLIPASE_ACTIVITY	0,059194	0,4845135	0,2065879	0,5539136	1,3685626	38

GOBP_NUCLEOBASE_BIOSYNTHETIC_PROCESS	0,0987984	0,6145457	0,1619789	0,6150718	1,3678815	19

GOBP_VENTRAL_SPINAL_CORD_DEVELOPMENT	0,0786082	0,5645811	0,1797823	0,5761712	1,3676256	29

GOBP_CHROMATIN_ORGANIZATION	0,0003851	0,0130351	0,4984931	0,441017	1,3675102	493

GOBP_OUTER_DYNEIN_ARM_ASSEMBLY	0,1001335	0,6157146	0,1608014	0,6147352	1,367133	19

GOBP_HETEROCHROMATIN_ORGANIZATION	0,0459364	0,4458081	0,2279872	0,506582	1,3670483	74

GOBP_SMOOTHENED_SIGNALING_PATHWAY	0,0244479	0,3072694	0,3524879	0,476622	1,3650116	130

GOBP_PERIPHERAL_NERVOUS_SYSTEM_DEVELOPMENT	0,052381	0,4619607	0,2139279	0,5096742	1,3649557	68

GOBP_RESPONSE_TO_CADMIUM_ION	0,0621827	0,4981249	0,2020717	0,5399994	1,3643239	46

GOBP_CELLULAR_AMINO_ACID_CATABOLIC_PROCESS	0,0331808	0,3750759	0,2663507	0,4939476	1,364277	92

GOBP_NEGATIVE_REGULATION_OF_NEURON_APOPTOTIC_PROCESS	0,0208352	0,2805513	0,3524879	0,4776256	1,3621736	126

GOBP_POSITIVE_REGULATION_OF_NERVOUS_SYSTEM_PROCESS	0,0980645	0,6130256	0,1596467	0,5802803	1,3621143	26

GOBP_REGULATION_OF_DNA_METABOLIC_PROCESS	0,0027681	0,0670951	0,4317077	0,4458929	1,3609311	359

GOBP_TELOMERE_ORGANIZATION	0,0275938	0,3365908	0,2878571	0,4752009	1,3608157	135

GOBP_NEGATIVE_REGULATION_OF_NEURON_DEATH	0,0141758	0,2140825	0,3807304	0,4624438	1,3592595	176

GOBP_NEGATIVE_REGULATION_OF_HISTONE_METHYLATION	0,1013333	0,6180346	0,1596467	0,6009165	1,3587674	21

GOBP_DNA_DOUBLE_STRAND_BREAK_PROCESSING	0,0948509	0,6050565	0,1669338	0,6441683	1,3587333	15

GOBP_HISTONE_EXCHANGE	0,0956873	0,6059163	0,1656567	0,6156888	1,3582685	18

GOBP_RESPONSE_TO_ACIDIC_PH	0,105474	0,6209246	0,1563124	0,6105936	1,3579224	19

GOBP_REGULATION_OF_NEUROBLAST_PROLIFERATION	0,1045161	0,6209246	0,154191	0,5778719	1,356461	26

GOBP_LUNG_CELL_DIFFERENTIATION	0,1058981	0,6209246	0,1563124	0,6316838	1,3562554	16

GOBP_TELOMERE_MAINTENANCE	0,0276855	0,3366162	0,2878571	0,475804	1,3562226	125

GOBP_OOCYTE_MATURATION	0,0932091	0,6018376	0,1669338	0,5990632	1,3559704	22

GOBP_ENDOCARDIAL_CUSHION_MORPHOGENESIS	0,0777917	0,5631281	0,1782199	0,5573814	1,3559666	33

GOBP_NEUROTROPHIN_TRK_RECEPTOR_SIGNALING_PATHWAY	0,0911458	0,5997765	0,1669338	0,588792	1,3557875	23

GOBP_POSITIVE_REGULATION_OF_NEURON_DIFFERENTIATION	0,0518868	0,4586792	0,2139279	0,5036914	1,3552178	72

GOBP_STEROID_CATABOLIC_PROCESS	0,1058981	0,6209246	0,1563124	0,6306971	1,3541369	16

GOBP_POSITIVE_REGULATION_OF_REPRODUCTIVE_PROCESS	0,0509709	0,4576243	0,2192503	0,5154145	1,353313	57

GOBP_NEURAL_TUBE_PATTERNING	0,0932312	0,6018376	0,1631801	0,5682263	1,3532574	30

GOBP_INTERLEUKIN_6_MEDIATED_SIGNALING_PATHWAY	0,1072386	0,6209246	0,155242	0,6301648	1,3529942	16

GOBP_POSITIVE_REGULATION_OF_MRNA_SPLICING_VIA_SPLICEOSOME	0,1068091	0,6209246	0,155242	0,6080996	1,3523759	19

GOBP_NEGATIVE_REGULATION_OF_GENE_EXPRESSION_EPIGENETIC	0,0496454	0,4544318	0,2192503	0,5018127	1,3508903	73

GOBP_REGULATION_OF_DOUBLE_STRAND_BREAK_REPAIR_VIA_HOMOLOGOUS_RECOMBINATION	0,0548926	0,4717694	0,208955	0,5053666	1,3508806	67

GOBP_CHONDROITIN_SULFATE_PROTEOGLYCAN_BIOSYNTHETIC_PROCESS	0,0958722	0,6059163	0,1644058	0,5965112	1,350194	22

GOBP_SPERM_FLAGELLUM_ASSEMBLY	0,0898438	0,5958215	0,1682382	0,5721811	1,3488042	28

GOBP_PHOSPHATIDYLCHOLINE_BIOSYNTHETIC_PROCESS	0,1021992	0,6192943	0,1563124	0,578757	1,3487163	25

GOBP_RESPONSE_TO_GAMMA_RADIATION	0,0560976	0,4725528	0,208955	0,5179999	1,3486926	54

GOBP_POSITIVE_REGULATION_OF_TRANSFERASE_ACTIVITY	0,0011153	0,0324835	0,4550599	0,4362632	1,3483656	467

GOBP_LABYRINTHINE_LAYER_DEVELOPMENT	0,0768262	0,5607311	0,1797823	0,5456204	1,3480725	38

GOBP_NEGATIVE_REGULATION_OF_RECEPTOR_SIGNALING_PATHWAY_VIA_STAT	0,0972037	0,6117159	0,1631801	0,5951085	1,3470189	22

GOBP_MATURE_B_CELL_DIFFERENTIATION	0,1103723	0,623409	0,1521449	0,5989337	1,3457552	20

GOBP_POSITIVE_REGULATION_OF_CYCLASE_ACTIVITY	0,104336	0,6209246	0,1585141	0,6374757	1,3446169	15

GOBP_NEURON_APOPTOTIC_PROCESS	0,0109489	0,1812109	0,3807304	0,45511	1,3445312	205

GOBP_PRODUCTION_OF_MOLECULAR_MEDIATOR_OF_IMMUNE_RESPONSE	0,0356757	0,3907683	0,2489111	0,4620675	1,3431376	155

GOBP_CENTRAL_NERVOUS_SYSTEM_NEURON_DIFFERENTIATION	0,0341786	0,3810357	0,2572065	0,4690476	1,3428889	136

GOBP_REGULATION_OF_NEURONAL_SYNAPTIC_PLASTICITY	0,0741206	0,5492527	0,1830239	0,5270045	1,3421388	48

GOBP_NUCLEOTIDE_EXCISION_REPAIR	0,0560976	0,4725528	0,208955	0,5134036	1,3421238	56

GOBP_MISMATCH_REPAIR	0,0914867	0,5997765	0,1644058	0,5610233	1,3418252	31

GOBP_CELLULAR_RESPONSE_TO_ESTRADIOL_STIMULUS	0,0896465	0,5958215	0,1656567	0,5558055	1,341512	32

GOBP_REGULATION_OF_CELLULAR_AMINE_METABOLIC_PROCESS	0,1132813	0,6289167	0,1482615	0,5720686	1,3412745	27

GOBP_REGULATION_OF_SYNAPSE_ASSEMBLY	0,047836	0,4476616	0,2192503	0,4849523	1,3411524	86

GOBP_REGULATION_OF_DNA_BINDING	0,0405405	0,4184363	0,2377938	0,477223	1,3397578	105

GOBP_NUCLEOSOME_ORGANIZATION	0,0355951	0,3907683	0,2529611	0,473456	1,3389696	114

GOBP_REGULATION_OF_GLIAL_CELL_DIFFERENTIATION	0,0612245	0,4946675	0,197822	0,5061423	1,3386285	63

GOBP_NEURON_DEATH	0,0051724	0,1033661	0,4070179	0,4443192	1,338412	297

GOBP_OLIGODENDROCYTE_DIFFERENTIATION	0,0535308	0,4666242	0,2065879	0,4832341	1,3378796	87

GOBP_ORGANELLE_LOCALIZATION	0,0017082	0,0455167	0,4550599	0,4315608	1,3376627	490

GOBP_POSITIVE_REGULATION_OF_NEUROBLAST_PROLIFERATION	0,1183511	0,6334603	0,1464162	0,5951926	1,3373492	20

GOBP_GLOMERULAR_EPITHELIAL_CELL_DIFFERENTIATION	0,11749	0,6334603	0,1473312	0,6012762	1,3372011	19

GOBP_CELLULAR_SODIUM_ION_HOMEOSTASIS	0,1179625	0,6334603	0,1473312	0,6227772	1,3371325	16

GOBP_TELOMERE_MAINTENANCE_VIA_RECOMBINATION	0,1138211	0,6297879	0,1511488	0,6339023	1,3370795	15

GOBP_HISTONE_DEACETYLATION	0,0469108	0,4461866	0,222056	0,4840139	1,3358958	93

GOBP_CELLULAR_RESPONSE_TO_GAMMA_RADIATION	0,1060026	0,6209246	0,1521449	0,5609313	1,335884	30

GOBP_ESTABLISHMENT_OF_CELL_POLARITY	0,0365449	0,3930483	0,2489111	0,4667945	1,3350675	133

GOBP_DEVELOPMENTAL_PIGMENTATION	0,0855346	0,5832186	0,1695706	0,5374917	1,3343131	40

GOBP_SIGNAL_TRANSDUCTION_INVOLVED_IN_REGULATION_OF_GENE_EXPRESSION	0,1179625	0,6334603	0,1473312	0,6212905	1,3339406	16

GOBP_SPINAL_CORD_DEVELOPMENT	0,0579196	0,4816976	0,2020717	0,4954224	1,3336876	73

GOBP_MICROTUBULE_BASED_TRANSPORT	0,0193944	0,2661509	0,3524879	0,4518766	1,3333014	195

GOBP_PURINERGIC_NUCLEOTIDE_RECEPTOR_SIGNALING_PATHWAY	0,1172507	0,6334603	0,1482615	0,6043366	1,3332245	18

GOBP_SPINAL_CORD_MOTOR_NEURON_DIFFERENTIATION	0,1236702	0,6435305	0,1429011	0,5933374	1,3331807	20

GOBP_PROTEOGLYCAN_METABOLIC_PROCESS	0,0591017	0,4845135	0,1999152	0,4951377	1,3329209	73

GOBP_ASYMMETRIC_CELL_DIVISION	0,1194631	0,6357264	0,1464162	0,6114795	1,3324575	17

GOBP_RESPONSE_TO_PH	0,0997475	0,6154476	0,1563124	0,5519593	1,3322284	32

GOBP_RESPONSE_TO_TOXIC_SUBSTANCE	0,0299465	0,3578315	0,2712886	0,4532215	1,3321584	182

GOBP_IN_UTERO_EMBRYONIC_DEVELOPMENT	0,0061196	0,1179047	0,4070179	0,4375637	1,3317944	345

GOBP_REGULATION_OF_PHOSPHOLIPASE_ACTIVITY	0,0777778	0,5631281	0,1766943	0,5190453	1,3301036	49

GOBP_PHOTORECEPTOR_CELL_DIFFERENTIATION	0,0799492	0,5678014	0,1766943	0,5241448	1,330088	47

GOBP_VOCALIZATION_BEHAVIOR	0,1193029	0,6357264	0,1464162	0,6191199	1,3292801	16

GOBP_NEGATIVE_REGULATION_OF_BMP_SIGNALING_PATHWAY	0,0880503	0,5913792	0,1669338	0,5256236	1,32891	44

GOBP_LYMPHOCYTE_MEDIATED_IMMUNITY	0,0331551	0,3750759	0,2572065	0,4520249	1,3288892	186

GOBP_REGULATION_OF_OLIGODENDROCYTE_DIFFERENTIATION	0,0920555	0,5997765	0,1631801	0,54085	1,3288395	36

GOBP_NEGATIVE_REGULATION_OF_AMINE_TRANSPORT	0,1201602	0,6385317	0,1455161	0,5974955	1,3287931	19

GOBP_AMYLOID_BETA_METABOLIC_PROCESS	0,080402	0,5678014	0,175204	0,5216805	1,3285799	48

GOBP_RESPONSE_TO_ZINC_ION	0,0920555	0,5997765	0,1631801	0,5435996	1,3282088	35

GOBP_HIPPOCAMPUS_DEVELOPMENT	0,0678571	0,5171182	0,1864326	0,4956226	1,3273242	68

GOBP_NEGATIVE_REGULATION_OF_CELL_SUBSTRATE_JUNCTION_ORGANIZATION	0,1261745	0,6484782	0,1420566	0,6090011	1,3270568	17

GOBP_POSITIVE_REGULATION_OF_CHROMATIN_ORGANIZATION	0,1205962	0,6395918	0,1464162	0,6291028	1,326956	15

GOBP_POSITIVE_REGULATION_OF_GLIOGENESIS	0,0695122	0,5265268	0,1864326	0,5085717	1,3261971	55

GOBP_POSITIVE_REGULATION_OF_ATP_DEPENDENT_ACTIVITY	0,0919395	0,5997765	0,1631801	0,5366374	1,325878	38

GOBP_REGULATION_OF_MICROTUBULE_BASED_MOVEMENT	0,0944584	0,6050565	0,1608014	0,5363927	1,3252734	38

GOBP_STRIATED_MUSCLE_CELL_PROLIFERATION	0,0743902	0,5501656	0,1797823	0,5088155	1,3247796	54

GOBP_CELL_FATE_DETERMINATION	0,1123883	0,6263281	0,1473312	0,5561819	1,3245731	30

GOBP_PROXIMAL_DISTAL_PATTERN_FORMATION	0,123057	0,6412276	0,1412251	0,569976	1,323953	24

GOBP_AXIS_SPECIFICATION	0,0678571	0,5171182	0,1864326	0,4943072	1,3238015	68

GOBP_REGULATION_OF_CATECHOLAMINE_METABOLIC_PROCESS	0,1246649	0,6442448	0,1429011	0,6163586	1,3233516	16

GOBP_EMBRYONIC_SKELETAL_SYSTEM_DEVELOPMENT	0,0516055	0,4583496	0,2114002	0,478886	1,3230855	94

GOBP_INNER_EAR_MORPHOGENESIS	0,0653442	0,5072278	0,1882041	0,4881622	1,3229241	76

GOBP_REGULATION_OF_MITOTIC_SPINDLE_ASSEMBLY	0,1306667	0,6552847	0,1388051	0,5849049	1,3225627	21

GOBP_CELLULAR_RESPONSE_TO_CADMIUM_ION	0,109375	0,6226089	0,1511488	0,5610101	1,3224709	28

GOBP_G1_TO_G0_TRANSITION	0,1241656	0,6435974	0,1429011	0,5946365	1,3224348	19

GOBP_NEGATIVE_REGULATION_OF_T_CELL_DIFFERENTIATION	0,0959596	0,6059163	0,1596467	0,5435354	1,3223348	34

GOBP_SPLICEOSOMAL_COMPLEX_ASSEMBLY	0,0825243	0,5762894	0,1695706	0,5035736	1,3222228	57

GOBP_NEURON_MIGRATION	0,0437637	0,4384537	0,2249661	0,4586967	1,3206524	142

GOBP_NUCLEOSIDE_MONOPHOSPHATE_BIOSYNTHETIC_PROCESS	0,09875	0,6145457	0,1563124	0,532542	1,3205624	39

GOBP_POSITIVE_REGULATION_OF_SMOOTHENED_SIGNALING_PATHWAY	0,1143583	0,6304365	0,1455161	0,551954	1,3201339	31

GOBP_SCHWANN_CELL_DIFFERENTIATION	0,1008827	0,6180254	0,155242	0,5372896	1,3200919	36

GOBP_ANIMAL_ORGAN_REGENERATION	0,0867347	0,588199	0,1695706	0,5234618	1,319356	45

GOBP_PRESYNAPSE_ORGANIZATION	0,0862944	0,5862715	0,1695706	0,5198926	1,3192976	47

GOBP_HEAD_MORPHOGENESIS	0,1098485	0,6232617	0,1482615	0,5466009	1,3192954	32

GOBP_NEGATIVE_REGULATION_OF_CELL_JUNCTION_ASSEMBLY	0,1106771	0,623409	0,1501698	0,5595999	1,3191467	28

GOBP_VENTRICULAR_SYSTEM_DEVELOPMENT	0,117268	0,6334603	0,1446305	0,5552559	1,3179799	29

GOBP_INTERMEMBRANE_LIPID_TRANSFER	0,0952986	0,6058153	0,1608014	0,5236206	1,3167796	43

| GOBP_NEGATIVE_REGULATION_OF_LIPID_CATABOLIC_PROCESS | 0,1353887 0,6623118 0,1364904 0,6128698 1,3158608 | 16 |
| --- | --- | --- |
| GOBP_REGULATION_OF_TELOMERASE_ACTIVITY | 0,0875635 0,5907538 0,1682382 0,51845 1,3156367 | 47 |
| GOBP_AMINO_ACID_IMPORT_ACROSS_PLASMA_MEMBRANE | 0,0993711 0,615061 0,1563124 0,5296189 1,3147691 | 40 |
| GOBP_NEGATIVE_REGULATION_OF_G_PROTEIN_COUPLED_RECEPTOR_SIGNALING_PATHWAY | 0,1099874 0,6232617 0,1482615 0,52428 1,3127793 | 42 |
| GOBP_RHYTHMIC_PROCESS | 0,0193438 0,2661509 0,3524879 0,4384145 1,3115976 | 248 |
| GOBP_APPENDAGE_DEVELOPMENT | 0,0458015 0,4458081 0,2192503 0,4532358 1,3104218 | 145 |
| GOBP_POSITIVE_REGULATION_OF_NEURON_APOPTOTIC_PROCESS | 0,1099874 0,6232617 0,1482615 0,523276 1,3102653 | 42 |
| GOBP_MESONEPHROS_DEVELOPMENT | 0,0629371 0,5000396 0,1918922 0,478873 1,3100888 | 81 |
| GOBP_POSITIVE_REGULATION_OF_STEM_CELL_PROLIFERATION | 0,1335113 0,6600028 0,1372508 0,5890762 1,3100689 | 19 |
| GOBP_POTASSIUM_ION_IMPORT_ACROSS_PLASMA_MEMBRANE | 0,1146096 0,6304365 0,1446305 0,5298387 1,3090804 | 38 |
| GOBP_NEGATIVE_REGULATION_OF_RESPONSE_TO_DNA_DAMAGE_STIMULUS | 0,0770128 0,5607311 0,1723243 0,4829072 1,3086831 | 76 |
| GOBP_REGULATION_OF_TELOMERE_MAINTENANCE_VIA_TELOMERE_LENGTHENING | 0,092233 0,5997765 0,1596467 0,4978916 1,3083436 | 58 |
| GOBP_RESPONSE_TO_NERVE_GROWTH_FACTOR | 0,0913706 0,5997765 0,1644058 0,5177264 1,3080504 | 46 |
| GOBP_NEGATIVE_REGULATION_OF_MICROTUBULE_POLYMERIZATION_OR_DEPOLYMERIZATION | 0,10625 0,6209246 0,1501698 0,5273132 1,3075964 | 39 |
| GOBP_RETINA_LAYER_FORMATION | 0,1422872 0,6810828 0,1321473 0,581782 1,3072166 | 20 |
| GOBP_MALE_SEX_DIFFERENTIATION | 0,0465116 0,4458081 0,2192503 0,4585559 1,3070589 | 125 |
| GOBP_NEUROTRANSMITTER_REUPTAKE | 0,1289911 0,6540073 0,1364904 0,5487246 1,3068134 | 30 |
| GOBP_CELLULAR_RESPONSE_TO_VITAMIN_D | 0,1449664 0,684728 0,1314576 0,5995417 1,3064441 | 17 |
| GOBP_RESPONSE_TO_MUSCLE_ACTIVITY | 0,1449664 0,684728 0,1314576 0,5995338 1,3064269 | 17 |
| GOBP_L_GLUTAMATE_TRANSMEMBRANE_TRANSPORT | 0,1367742 0,6680012 0,1328463 0,5565514 1,3064145 | 26 |
| GOBP_POSTSYNAPTIC_SPECIALIZATION_ASSEMBLY | 0,1401617 0,6759789 0,1342735 0,5921685 1,3063805 | 18 |
| GOBP_REGULATION_OF_HOMOTYPIC_CELL_CELL_ADHESION | 0,1315104 0,6552847 0,1364904 0,557108 1,3061978 | 27 |
| GOBP_DEOXYRIBONUCLEOSIDE_MONOPHOSPHATE_METABOLIC_PROCESS | 0,1388518 0,6731177 0,1342735 0,5869013 1,3052321 | 19 |
| GOBP_NEGATIVE_REGULATION_OF_CYCLIN_DEPENDENT_PROTEIN_SERINE_THREONINE_KINASE_ACTIVITY | 0,1289063 0,6540073 0,1380222 0,5536617 1,3051485 | 28 |
| GOBP_LEARNING | 0,0498339 0,4544318 0,2114002 0,4562272 1,304844 | 133 |
| GOBP_REGULATION_OF_MRNA_PROCESSING | 0,0553097 0,4725528 0,1999152 0,4555221 1,304583 | 130 |
| GOBP_REPRODUCTIVE_SYSTEM_DEVELOPMENT | 0,0125831 0,2003162 0,3807304 0,4283599 1,304407 | 349 |
| GOBP_DEOXYRIBOSE_PHOSPHATE_METABOLIC_PROCESS | 0,1056604 0,6209246 0,1511488 0,5252471 1,3039161 | 40 |
| GOBP_CELLULAR_RESPONSE_TO_LIGHT_STIMULUS | 0,064 0,5002749 0,1882041 0,4702829 1,3039038 | 97 |
| GOBP_POSITIVE_REGULATION_OF_LIPASE_ACTIVITY | 0,1067344 0,6209246 0,1511488 0,5183053 1,3034128 | 43 |
| GOBP_POSITIVE_REGULATION_OF_NEURON_DEATH | 0,0801887 0,5678014 0,1695706 0,4842162 1,3028184 | 72 |
| GOBP_PLACENTA_BLOOD_VESSEL_DEVELOPMENT | 0,1315104 0,6552847 0,1364904 0,5525468 1,3025202 | 28 |
| GOBP_NEGATIVE_REGULATION_OF_VASCULATURE_DEVELOPMENT | 0,0788863 0,5645811 0,1695706 0,4767763 1,3016749 | 79 |
| GOBP_RNA_SPLICING_VIA_TRANSESTERIFICATION_REACTIONS | 0,0260146 0,3215024 0,2878571 0,4322997 1,3016404 | 275 |
| GOBP_NEGATIVE_REGULATION_OF_INFLAMMATORY_RESPONSE_TO_ANTIGENIC_STIMULUS | 0,1455526 0,684728 0,1314576 0,5900049 1,3016073 | 18 |
| GOBP_NEURONAL_STEM_CELL_POPULATION_MAINTENANCE | 0,1344874 0,6613468 0,1364904 0,5749087 1,3012969 | 22 |
| GOBP_GLOMERULAR_EPITHELIUM_DEVELOPMENT | 0,1344874 0,6613468 0,1364904 0,5748537 1,3011726 | 22 |
| GOBP_RESPIRATORY_SYSTEM_DEVELOPMENT | 0,0506466 0,4574017 0,2065879 0,4441365 1,3006227 | 170 |
| GOBP_REGULATION_OF_MRNA_SPLICING_VIA_SPLICEOSOME | 0,0699541 0,5284543 0,1797823 0,4707515 1,3006113 | 94 |
| GOBP_EMBRYONIC_CRANIAL_SKELETON_MORPHOGENESIS | 0,1125 0,6263281 0,1455161 0,5243523 1,3002542 | 39 |
| GOBP_NEGATIVE_REGULATION_OF_BLOOD_PRESSURE | 0,1308767 0,6552847 0,135002 0,5430847 1,2989207 | 31 |
| GOBP_NEGATIVE_REGULATION_OF_SIGNALING_RECEPTOR_ACTIVITY | 0,1221662 0,6395918 0,1395997 0,5256343 1,2986924 | 38 |
| GOBP_REGULATION_OF_BMP_SIGNALING_PATHWAY | 0,0801833 0,5678014 0,1669338 0,4706473 1,2974529 | 84 |
| GOBP_POSITIVE_REGULATION_OF_BMP_SIGNALING_PATHWAY | 0,1186869 0,6337269 0,1420566 0,5326569 1,2958693 | 34 |
| GOBP_NEGATIVE_REGULATION_OF_DEFENSE_RESPONSE | 0,0541126 0,4706039 0,1999152 0,443537 1,2958014 | 165 |
| GOBP_TELOMERE_CAPPING | 0,11875 0,6337269 0,1412251 0,522546 1,2957751 | 39 |
| GOBP_POSITIVE_REGULATION_OF_SMOOTH_MUSCLE_CELL_MIGRATION | 0,1353768 0,6623118 0,1328463 0,5439908 1,2955396 | 30 |
| GOBP_POSITIVE_REGULATION_OF_SPROUTING_ANGIOGENESIS | 0,1495957 0,6878113 0,1294429 0,5870524 1,2950939 | 18 |
| GOBP_CELLULAR_RESPONSE_TO_STEROL | 0,1487936 0,6878113 0,1294429 0,6030092 1,2946897 | 16 |
| GOBP_REGULATION_OF_OSTEOBLAST_DIFFERENTIATION | 0,0685393 0,5210682 0,1797823 0,4599537 1,2946835 | 108 |
| GOBP_2_OXOGLUTARATE_METABOLIC_PROCESS | 0,150134 0,6878113 0,1287887 0,6026055 1,2938231 | 16 |
| GOBP_REGULATION_OF_NUCLEAR_TRANSCRIBED_MRNA_POLY_A_TAIL_SHORTENING | 0,1449864 0,684728 0,1321473 0,6132005 1,2934135 | 15 |
| GOBP_CELL_AGGREGATION | 0,1449864 0,684728 0,1321473 0,6129264 1,2928354 | 15 |
| GOBP_NEGATIVE_REGULATION_OF_MRNA_SPLICING_VIA_SPLICEOSOME | 0,1476064 0,6878113 0,1294429 0,5753355 1,2927319 | 20 |
| GOBP_DORSAL_VENTRAL_PATTERN_FORMATION | 0,0883055 0,5913792 0,1619789 0,483532 1,2925153 | 67 |
| GOBP_NEUROTRANSMITTER_TRANSPORT | 0,0513919 0,4583496 0,2042948 0,4396974 1,2924009 | 176 |
| GOBP_EATING_BEHAVIOR | 0,1476064 0,6878113 0,1294429 0,575053 1,292097 | 20 |
| GOBP_LEFT_RIGHT_PATTERN_FORMATION | 0,1533333 0,6935605 0,1268757 0,5714222 1,292076 | 21 |
| GOBP_METANEPHROS_DEVELOPMENT | 0,0943396 0,6050565 0,155242 0,4799525 1,2913466 | 72 |
| GOBP_REGULATION_OF_CENTROSOME_CYCLE | 0,1074074 0,6209246 0,1482615 0,503848 1,291159 | 49 |
| GOBP_ADULT_BEHAVIOR | 0,0723831 0,5412925 0,1737478 0,457029 1,2905897 | 112 |
| GOBP_RESPONSE_TO_ATP | 0,1489637 0,6878113 0,1268757 0,5555521 1,2904488 | 24 |
| GOBP_MITOCHONDRIAL_GENOME_MAINTENANCE | 0,1536458 0,6935605 0,1250334 0,5601304 1,2897896 | 23 |
| GOBP_POLYOL_METABOLIC_PROCESS | 0,0706956 0,5322709 0,1782199 0,4650621 1,2896131 | 96 |
| GOBP_CELLULAR_RESPONSE_TO_COPPER_ION | 0,1590296 0,7040708 0,1250334 0,5839199 1,2881832 | 18 |
| GOBP_ANTIGEN_PROCESSING_AND_PRESENTATION | 0,0785877 0,5645811 0,1682382 0,4657689 1,2881001 | 86 |
| GOBP_REGULATION_OF_ENDOTHELIAL_CELL_CHEMOTAXIS | 0,1597315 0,7043561 0,1244342 0,5910998 1,2880487 | 17 |
| GOBP_REGULATION_OF_ACUTE_INFLAMMATORY_RESPONSE | 0,1536458 0,6935605 0,1250334 0,5593545 1,288003 | 23 |
| GOBP_ORGANIC_HYDROXY_COMPOUND_CATABOLIC_PROCESS | 0,1085366 0,6209246 0,1464162 0,4938637 1,2878432 | 55 |
| GOBP_POSITIVE_REGULATION_OF_DOUBLE_STRAND_BREAK_REPAIR_VIA_HOMOLOGOUS_RECOMBINATION | 0,1311475 0,6552847 0,1342735 0,5268157 1,2871996 | 35 |
| GOBP_RESPONSE_TO_INTERLEUKIN_1 | 0,0732265 0,5437677 0,175204 0,4659899 1,2861489 | 93 |
| GOBP_NEGATIVE_REGULATION_OF_TELOMERE_MAINTENANCE_VIA_TELOMERE_LENGTHENING | 0,1541451 0,6935605 0,1244342 0,5535954 1,2859037 | 24 |
| GOBP_MRNA_CIS_SPLICING_VIA_SPLICEOSOME | 0,1541451 0,6935605 0,1244342 0,5532207 1,2850336 | 24 |
| GOBP_GLIAL_CELL_MIGRATION | 0,1248424 0,6442758 0,1380222 0,5148082 1,284092 | 41 |
| GOBP_CELL_DIFFERENTIATION_INVOLVED_IN_EMBRYONIC_PLACENTA_DEVELOPMENT | 0,150466 0,6878113 0,1281429 0,5672929 1,2840588 | 22 |
| GOBP_EMBRYONIC_SKELETAL_SYSTEM_MORPHOGENESIS | 0,0890736 0,5944043 0,1608014 0,4794278 1,2839282 | 70 |
| GOBP_POSITIVE_REGULATION_OF_MRNA_PROCESSING | 0,1458333 0,684728 0,1287887 0,5444469 1,2834264 | 28 |
| GOBP_POSITIVE_REGULATION_OF_ANION_TRANSPORT | 0,1325758 0,657107 0,133555 0,5272545 1,282726 | 34 |
| GOBP_PROTEIN_KINASE_C_SIGNALING | 0,1484375 0,6878113 0,1275053 0,5470521 1,2826208 | 27 |
| GOBP_REGULATION_OF_PROTEIN_MATURATION | 0,1124694 0,6263281 0,143759 0,4960603 1,2820097 | 52 |
| GOBP_NEGATIVE_REGULATION_OF_TISSUE_REMODELING | 0,1595174 0,7042385 0,1244342 0,5969653 1,2817131 | 16 |
| GOBP_DORSAL_VENTRAL_NEURAL_TUBE_PATTERNING | 0,1548732 0,6957407 0,126254 0,5762245 1,2814876 | 19 |
| GOBP_ADAPTIVE_IMMUNE_RESPONSE_BASED_ON_SOMATIC_RECOMBINATION_OF_IMMUNE_RECEPTORS_BUILT_FROM_IMMUNOGLOBULIN_SUPERFAMILY_DOMAINS | 0,0436635 0,4384537 0,222056 0,4333195 1,2808476 | 202 |
| GOBP_NEGATIVE_REGULATION_OF_HISTONE_MODIFICATION | 0,1219822 0,6395918 0,1404062 0,5091746 1,2804512 | 43 |
| GOBP_MAMMARY_GLAND_EPITHELIUM_DEVELOPMENT | 0,1067961 0,6209246 0,1473312 0,487538 1,2801182 | 57 |
| GOBP_SYNAPTIC_TRANSMISSION_GABAERGIC | 0,1298865 0,6552847 0,135002 0,5130172 1,2796247 | 41 |
| GOBP_POSITIVE_REGULATION_OF_CALCIUM_ION_IMPORT | 0,1571816 0,7001174 0,126254 0,606596 1,2794827 | 15 |
| GOBP_NEURON_FATE_COMMITMENT | 0,138539 0,6724692 0,1301056 0,5177786 1,2792832 | 38 |
| GOBP_DEVELOPMENTAL_MATURATION | 0,0446334 0,4436179 0,2192503 0,4316544 1,2792752 | 218 |
| GOBP_PULMONARY_VALVE_DEVELOPMENT | 0,1725067 0,7265238 0,1193484 0,579092 1,2775324 | 18 |
| GOBP_ENDOCARDIAL_CUSHION_DEVELOPMENT | 0,1325 0,657107 0,1328463 0,5151385 1,2774064 | 39 |
| GOBP_RESPONSE_TO_AUDITORY_STIMULUS | 0,1567358 0,6993542 0,1232572 0,5498699 1,27725 | 24 |
| GOBP_REGULATION_OF_NEUROTRANSMITTER_TRANSPORT | 0,0882016 0,5913792 0,1585141 0,4627534 1,2769809 | 85 |
| GOBP_MICROTUBULE_ORGANIZING_CENTER_LOCALIZATION | 0,1486658 0,6878113 0,1256399 0,5338876 1,2769236 | 31 |
| GOBP_COCHLEA_MORPHOGENESIS | 0,1689008 0,7227339 0,1204334 0,5944018 1,2762091 | 16 |
| GOBP_PHAGOSOME_MATURATION | 0,1630013 0,7137481 0,1204334 0,5471335 1,275022 | 25 |
| GOBP_FACE_DEVELOPMENT | 0,1257862 0,647368 0,1372508 0,504258 1,2748922 | 44 |
| GOBP_TELOMERE_MAINTENANCE_VIA_TELOMERE_LENGTHENING | 0,0952941 0,6058153 0,154191 0,4720903 1,2748544 | 75 |
| GOBP_PROTEIN_TETRAMERIZATION | 0,0997625 0,6154476 0,1511488 0,475869 1,2743976 | 70 |
| GOBP_STRIATUM_DEVELOPMENT | 0,1677852 0,7220723 0,1209851 0,5844286 1,2735115 | 17 |
| GOBP_REGULATION_OF_DNA_RECOMBINATION | 0,0822222 0,5752493 0,1619789 0,4494732 1,2731896 | 115 |
| GOBP_VISUAL_BEHAVIOR | 0,1276865 0,6535669 0,1364904 0,5084133 1,2730496 | 42 |
| GOBP_RETINA_VASCULATURE_DEVELOPMENT_IN_CAMERA_TYPE_EYE | 0,1752022 0,7308245 0,1182875 0,5765858 1,2720036 | 18 |
| GOBP_PROTEIN_CONTAINING_COMPLEX_LOCALIZATION | 0,0784314 0,5645811 0,1644058 0,4393506 1,2708819 | 147 |
| GOBP_REGULATION_OF_TELOMERE_MAINTENANCE | 0,1026838 0,6197349 0,1473312 0,4682678 1,2705651 | 77 |
| GOBP_ALCOHOL_CATABOLIC_PROCESS | 0,1276865 0,6535669 0,1364904 0,507216 1,2700515 | 42 |
| GOBP_REGULATION_OF_LIPASE_ACTIVITY | 0,1143201 0,6304365 0,1412251 0,480705 1,2700058 | 62 |
| GOBP_POSITIVE_REGULATION_OF_EXTRACELLULAR_MATRIX_ORGANIZATION | 0,1619171 0,7123215 0,1209851 0,5465934 1,2696394 | 24 |
| GOBP_POSITIVE_REGULATION_OF_TELOMERE_MAINTENANCE | 0,1283951 0,6540073 0,1342735 0,4938257 1,269405 | 50 |
| GOBP_EMBRYONIC_ORGAN_MORPHOGENESIS | 0,0465116 0,4458081 0,2139279 0,4266408 1,2691888 | 226 |
| GOBP_GLANDULAR_EPITHELIAL_CELL_DIFFERENTIATION | 0,1399748 0,6759789 0,1294429 0,5083854 1,2680717 | 41 |
| GOBP_IMMUNE_EFFECTOR_PROCESS | 0,0288949 0,3486277 0,3524879 0,4138485 1,2676807 | 383 |
| GOBP_NEGATIVE_REGULATION_OF_EPITHELIAL_CELL_MIGRATION | 0,1207317 0,6395918 0,1380222 0,4847576 1,2672385 | 56 |
| GOBP_BLASTOCYST_GROWTH | 0,1675532 0,7219006 0,1204334 0,563909 1,2670574 | 20 |
| GOBP_MESENCHYMAL_CELL_PROLIFERATION | 0,1525851 0,6931829 0,1232572 0,5184494 1,2667579 | 35 |
| GOBP_MICROTUBULE_POLYMERIZATION | 0,1032483 0,6209246 0,1464162 0,4639414 1,2666334 | 79 |
| GOBP_REGULATION_OF_REPRODUCTIVE_PROCESS | 0,0844444 0,5810582 0,1596467 0,4469951 1,2661701 | 115 |
| GOBP_MITOCHONDRIAL_DNA_METABOLIC_PROCESS | 0,1761518 0,7324279 0,1182875 0,6002438 1,2660841 | 15 |
| GOBP_POSITIVE_REGULATION_OF_CELL_DEVELOPMENT | 0,0530697 0,465004 0,197822 0,4213824 1,2655431 | 254 |
| GOBP_BENZENE_CONTAINING_COMPOUND_METABOLIC_PROCESS | 0,1735648 0,7269596 0,1182875 0,5687408 1,2648443 | 19 |
| GOBP_REGULATION_OF_DNA_DAMAGE_CHECKPOINT | 0,1788618 0,7387418 0,1172497 0,5993905 1,2642843 | 15 |
| GOBP_GLANDULAR_EPITHELIAL_CELL_DEVELOPMENT | 0,1627604 0,7137481 0,1209851 0,5390761 1,2639202 | 27 |
| GOBP_REGULATION_OF_NERVOUS_SYSTEM_DEVELOPMENT | 0,0193149 0,2661509 0,3524879 0,4124855 1,2630079 | 382 |
| GOBP_N_GLYCAN_PROCESSING | 0,1748999 0,7308245 0,1177658 0,5674508 1,2619754 | 19 |
| GOBP_DNA_MODIFICATION | 0,0948571 0,6050565 0,1521449 0,4564944 1,2610494 | 89 |
| GOBP_PROTEIN_DNA_COMPLEX_DISASSEMBLY | 0,176235 0,7324279 0,1172497 0,5669619 1,2608883 | 19 |
| GOBP_OUTFLOW_TRACT_SEPTUM_MORPHOGENESIS | 0,171875 0,7265238 0,1172497 0,5472161 1,2600525 | 23 |
| GOBP_NEGATIVE_REGULATION_OF_TELOMERE_MAINTENANCE | 0,1603535 0,7062699 0,1198878 0,5218471 1,2595486 | 32 |
| GOBP_REGULATION_OF_GLUCAN_BIOSYNTHETIC_PROCESS | 0,1696891 0,7244568 0,1177658 0,542137 1,2592881 | 24 |
| GOBP_NEUROMUSCULAR_PROCESS_CONTROLLING_BALANCE | 0,15 0,6878113 0,1238422 0,5076681 1,2588818 | 39 |
| GOBP_EMBRYONIC_HEMOPOIESIS | 0,1802403 0,7416899 0,1157344 0,566026 1,2588067 | 19 |
| GOBP_NEUROPEPTIDE_SIGNALING_PATHWAY | 0,1218593 0,6395918 0,1395997 0,4939949 1,2580722 | 48 |
| GOBP_EMBRYONIC_PLACENTA_DEVELOPMENT | 0,1120187 0,6263281 0,1404062 0,4641606 1,2578796 | 76 |
| GOBP_GLYCOSYLCERAMIDE_METABOLIC_PROCESS | 0,1876676 0,7580835 0,1133129 0,5855823 1,2572733 | 16 |
| GOBP_PHENOL_CONTAINING_COMPOUND_METABOLIC_PROCESS | 0,112529 0,6263281 0,1395997 0,4604925 1,2572173 | 79 |
| GOBP_POSITIVE_REGULATION_OF_ORGAN_GROWTH | 0,1639344 0,7153329 0,1182875 0,5141973 1,2563685 | 35 |
| GOBP_FOREBRAIN_DEVELOPMENT | 0,0414079 0,4246236 0,2249661 0,4146395 1,256293 | 320 |
| GOBP_HINDBRAIN_MORPHOGENESIS | 0,1525 0,6931829 0,1226792 0,506527 1,2560521 | 39 |
| GOBP_NUCLEUS_ORGANIZATION | 0,0875831 0,5907538 0,1563124 0,4393058 1,2559783 | 128 |
| GOBP_EMBRYONIC_EYE_MORPHOGENESIS | 0,1688144 0,7227339 0,1177658 0,5291157 1,2559325 | 29 |
| GOBP_RESPONSE_TO_CORTICOSTEROID | 0,110245 0,623409 0,1380222 0,4445041 1,2552213 | 112 |
| GOBP_HOMOPHILIC_CELL_ADHESION_VIA_PLASMA_MEMBRANE_ADHESION_MOLECULES | 0,0925926 0,5997765 0,1501698 0,4337993 1,2548241 | 147 |
| GOBP_DETOXIFICATION | 0,0852974 0,5826589 0,1596467 0,4468682 1,2547033 | 104 |
| GOBP_PURINE_NUCLEOBASE_METABOLIC_PROCESS | 0,1954178 0,7683417 0,1110115 0,5686121 1,2544127 | 18 |
| GOBP_UROGENITAL_SYSTEM_DEVELOPMENT | 0,0428422 0,4326394 0,222056 0,4164868 1,2543895 | 298 |
| GOBP_CRANIAL_SKELETAL_SYSTEM_DEVELOPMENT | 0,1225728 0,6395918 0,1364904 0,4771468 1,2538309 | 58 |
| GOBP_CELLULAR_RESPONSE_TO_ESTROGEN_STIMULUS | 0,1829268 0,7462064 0,1157344 0,5943457 1,2536435 | 15 |
| GOBP_POSITIVE_REGULATION_OF_TELOMERASE_ACTIVITY | 0,165404 0,7178712 0,1177658 0,5193092 1,2534232 | 32 |
| GOBP_NEGATIVE_REGULATION_OF_PHOSPHOPROTEIN_PHOSPHATASE_ACTIVITY | 0,196 0,7683417 0,1101223 0,5541944 1,2531213 | 21 |
| GOBP_PROSTATE_GLAND_DEVELOPMENT | 0,155 0,6957407 0,1215433 0,5052654 1,2529238 | 39 |
| GOBP_NEGATIVE_REGULATION_OF_SMOOTHENED_SIGNALING_PATHWAY | 0,1666667 0,7197318 0,1172497 0,519091 1,2528964 | 32 |
| GOBP_FOREBRAIN_CELL_MIGRATION | 0,1225728 0,6395918 0,1364904 0,4771643 1,2528804 | 57 |
| GOBP_ONE_CARBON_METABOLIC_PROCESS | 0,1643664 0,7155556 0,1177658 0,5149091 1,2526425 | 33 |
| GOBP_REGULATION_OF_NOTCH_SIGNALING_PATHWAY | 0,1053837 0,6209246 0,143759 0,4542352 1,2522089 | 84 |
| GOBP_CELLULAR_RESPONSE_TO_VITAMIN | 0,1848958 0,7509769 0,1123785 0,543749 1,2520689 | 23 |
| GOBP_RESPONSE_TO_ESTRADIOL | 0,0969213 0,6109621 0,1501698 0,4514363 1,2518401 | 95 |
| GOBP_CORTICAL_CYTOSKELETON_ORGANIZATION | 0,1317073 0,6553965 0,1314576 0,4799279 1,251503 | 55 |
| GOBP_EMBRYONIC_HEART_TUBE_MORPHOGENESIS | 0,1274272 0,6535669 0,133555 0,4765116 1,2511666 | 57 |
| GOBP_REGULATION_OF_RESPONSE_TO_DNA_DAMAGE_STIMULUS | 0,0607966 0,4922694 0,1847065 0,4192569 1,2507861 | 236 |
| GOBP_PROTEIN_LOCALIZATION_TO_MICROTUBULE | 0,1946309 0,7683417 0,1110115 0,5739414 1,2506592 | 17 |
| GOBP_PLATELET_DERIVED_GROWTH_FACTOR_RECEPTOR_SIGNALING_PATHWAY | 0,137037 0,6680012 0,1294429 0,4878302 1,250112 | 49 |
| GOBP_L_GLUTAMATE_IMPORT_ACROSS_PLASMA_MEMBRANE | 0,196477 0,7683417 0,1110115 0,5922806 1,2492876 | 15 |
| GOBP_AMYLOID_PRECURSOR_PROTEIN_CATABOLIC_PROCESS | 0,1374233 0,6680012 0,1287887 0,4820338 1,2490584 | 53 |
| GOBP_GLIAL_CELL_DEVELOPMENT | 0,0935006 0,6025414 0,1531588 0,450189 1,2483702 | 96 |
| GOBP_CARBOHYDRATE_BIOSYNTHETIC_PROCESS | 0,0770053 0,5607311 0,1644058 0,4249089 1,2483495 | 174 |
| GOBP_NEPHRON_DEVELOPMENT | 0,0922222 0,5997765 0,1521449 0,4385433 1,2480987 | 122 |
| GOBP_NEGATIVE_REGULATION_OF_TELOMERE_MAINTENANCE_VIA_TELOMERASE | 0,1975968 0,7692965 0,1096841 0,5609438 1,2475043 | 19 |
| GOBP_EMBRYONIC_ORGAN_DEVELOPMENT | 0,038105 0,4010102 0,2343926 0,4086074 1,246888 | 360 |
| GOBP_BEHAVIOR | 0,0215874 0,2849045 0,3524879 0,4026856 1,2465974 | 483 |
| GOBP_ALTERNATIVE_MRNA_SPLICING_VIA_SPLICEOSOME | 0,1223278 0,6395918 0,135002 0,4650952 1,2455449 | 70 |
| GOBP_SYNAPSE_ASSEMBLY | 0,0879479 0,5913792 0,154191 0,4277941 1,2450127 | 158 |
| GOBP_IRON_ION_TRANSMEMBRANE_TRANSPORT | 0,2008086 0,7744131 0,10925 0,5642826 1,2448616 | 18 |
| GOBP_CELLULAR_RESPONSE_TO_UV | 0,1138211 0,6297879 0,1388051 0,4540399 1,2436183 | 82 |
| GOBP_PROTEIN_LOCALIZATION_TO_CYTOSKELETON | 0,1414634 0,6791978 0,126254 0,4755285 1,2431119 | 56 |
| GOBP_POSITIVE_REGULATION_OF_DNA_BINDING | 0,143401 0,6837026 0,1281429 0,491969 1,2429735 | 46 |
| GOBP_NUCLEAR_MIGRATION | 0,208 0,7744131 0,1063233 0,5496747 1,2429015 | 21 |
| GOBP_POSITIVE_REGULATION_OF_INSULIN_SECRETION | 0,1488315 0,6878113 0,1232572 0,4823798 1,2425158 | 51 |
| GOBP_REGULATION_OF_CHROMATIN_ASSEMBLY | 0,1848958 0,7509769 0,1123785 0,526654 1,2414832 | 28 |
| GOBP_NEGATIVE_REGULATION_OF_ORGANELLE_ASSEMBLY | 0,159292 0,7040708 0,1204334 0,4956699 1,2411405 | 42 |
| GOBP_PHENOL_CONTAINING_COMPOUND_BIOSYNTHETIC_PROCESS | 0,1828499 0,7462064 0,1110115 0,5078955 1,2409708 | 35 |
| GOBP_POSITIVE_REGULATION_OF_TORC1_SIGNALING | 0,1981383 0,7706062 0,10925 0,5521227 1,2405746 | 20 |
| GOBP_MACROMOLECULE_DEACYLATION | 0,1053215 0,6209246 0,1412251 0,4340451 1,2403876 | 127 |
| GOBP_POSITIVE_REGULATION_OF_RECEPTOR_SIGNALING_PATHWAY_VIA_STAT | 0,1915709 0,7671945 0,1088201 0,5207676 1,2402324 | 30 |
| GOBP_REGULATION_OF_POSTSYNAPTIC_MEMBRANE_POTENTIAL | 0,1047297 0,6209246 0,1429011 0,4414863 1,2400919 | 106 |
| GOBP_SPROUTING_ANGIOGENESIS | 0,1144444 0,6304365 0,135002 0,4373175 1,2387569 | 115 |
| GOBP_GAMMA_AMINOBUTYRIC_ACID_SIGNALING_PATHWAY | 0,2034574 0,7744131 0,1075544 0,5512449 1,2386022 | 20 |
| GOBP_PYRIMIDINE_NUCLEOSIDE_MONOPHOSPHATE_METABOLIC_PROCESS | 0,2080537 0,7744131 0,1067299 0,568372 1,238523 | 17 |
| GOBP_REGULATION_OF_G_PROTEIN_COUPLED_RECEPTOR_SIGNALING_PATHWAY | 0,0987514 0,6145457 0,1482615 0,4439419 1,2376901 | 101 |
| GOBP_GENITALIA_DEVELOPMENT | 0,1806775 0,7426756 0,1114627 0,5081528 1,2362061 | 33 |
| GOBP_REGULATION_OF_INSULIN_SECRETION | 0,1222222 0,6395918 0,1301056 0,4360714 1,2352273 | 115 |
| GOBP_POSITIVE_REGULATION_OF_ERK1_AND_ERK2_CASCADE | 0,1083151 0,6209246 0,1380222 0,4290292 1,2344688 | 141 |
| GOBP_POSITIVE_REGULATION_OF_PLASMA_MEMBRANE_BOUNDED_CELL_PROJECTION_ASSEMBLY | 0,1151653 0,631644 0,1364904 0,4449472 1,2338458 | 95 |
| GOBP_BLASTOCYST_DEVELOPMENT | 0,1010216 0,6180254 0,1464162 0,4421916 1,2328105 | 101 |
| GOBP_CAMERA_TYPE_EYE_PHOTORECEPTOR_CELL_DIFFERENTIATION | 0,2029372 0,7744131 0,1079724 0,5543055 1,232741 | 19 |
| GOBP_FOREBRAIN_GENERATION_OF_NEURONS | 0,1731985 0,7265238 0,1147507 0,4921233 1,2322599 | 42 |
| GOBP_KIDNEY_EPITHELIUM_DEVELOPMENT | 0,1183628 0,6334603 0,1321473 0,4332054 1,2321183 | 119 |
| GOBP_REGULATION_OF_NEUROGENESIS | 0,0641822 0,5002749 0,1782199 0,4062189 1,2303127 | 319 |
| GOBP_NEGATIVE_REGULATION_OF_OSTEOBLAST_DIFFERENTIATION | 0,1813602 0,743854 0,1114627 0,4978565 1,2300614 | 38 |
| GOBP_LIVER_REGENERATION | 0,22 0,7888973 0,1028218 0,5439614 1,2299829 | 21 |
| GOBP_FILOPODIUM_ASSEMBLY | 0,1578313 0,7011945 0,1177658 0,4662151 1,2298727 | 60 |
| GOBP_REGULATION_OF_LIPID_CATABOLIC_PROCESS | 0,1775 0,7360568 0,1123785 0,4959531 1,2298317 | 39 |
| GOBP_POSITIVE_REGULATION_OF_NEURON_MIGRATION | 0,2181572 0,7873562 0,104344 0,5828644 1,2294261 | 15 |
| GOBP_CHROMATIN_DISASSEMBLY | 0,212938 0,7803554 0,1055209 0,5572414 1,2293279 | 18 |
| GOBP_INCLUSION_BODY_ASSEMBLY | 0,2213333 0,7889462 0,1024494 0,5434904 1,2289179 | 21 |
| GOBP_NUCLEIC_ACID_PHOSPHODIESTER_BOND_HYDROLYSIS | 0,0857741 0,5837919 0,1531588 0,4116678 1,228873 | 232 |
| GOBP_DETECTION_OF_LIGHT_STIMULUS | 0,1851385 0,7511506 0,1101223 0,4973302 1,2287612 | 38 |
| GOBP_REGULATION_OF_DNA_METHYLATION_DEPENDENT_HETEROCHROMATIN_ASSEMBLY | 0,2198391 0,7888973 0,1031975 0,5722214 1,2285868 | 16 |
| GOBP_OSTEOBLAST_DIFFERENTIATION | 0,0851064 0,5824129 0,155242 0,4159439 1,2279462 | 196 |
| GOBP_PYRIMIDINE_CONTAINING_COMPOUND_CATABOLIC_PROCESS | 0,1881313 0,7580835 0,10925 0,5086878 1,2277869 | 32 |
| GOBP_REGULATION_OF_ANIMAL_ORGAN_MORPHOGENESIS | 0,1217587 0,6395918 0,1314576 0,4366776 1,2277054 | 107 |
| GOBP_PROTEOGLYCAN_BIOSYNTHETIC_PROCESS | 0,1658537 0,7178712 0,11524 0,4714423 1,2274727 | 54 |
| GOBP_POSITIVE_REGULATION_OF_TRANSMEMBRANE_RECEPTOR_PROTEIN_SERINE_THREONINE_KINASE_SIGNALING_PATHWAY | 0,1081081 0,6209246 0,1404062 0,4371645 1,2272973 | 105 |
| GOBP_REGULATION_OF_PROTEIN_SERINE_THREONINE_KINASE_ACTIVITY | 0,0639835 0,5002749 0,1782199 0,4052398 1,2269483 | 315 |
| GOBP_HEART_FORMATION | 0,2137306 0,7803554 0,1028218 0,5279438 1,2263196 | 24 |
| GOBP_CELL_PROLIFERATION_IN_FOREBRAIN | 0,2201342 0,7888973 0,1031975 0,5626701 1,2260983 | 17 |
| GOBP_NUCLEUS_LOCALIZATION | 0,2122396 0,7803554 0,1035763 0,5324692 1,2260952 | 23 |
| GOBP_REGULATION_OF_GENE_EXPRESSION_EPIGENETIC | 0,1337017 0,6600254 0,1232572 0,4311183 1,2258063 | 117 |
| GOBP_NEGATIVE_REGULATION_OF_NUCLEOCYTOPLASMIC_TRANSPORT | 0,224734 0,7901992 0,1013507 0,5453635 1,2253872 | 20 |
| GOBP_LONG_CHAIN_FATTY_ACID_METABOLIC_PROCESS | 0,1392111 0,67399 0,1238422 0,4488229 1,2253577 | 79 |
| GOBP_AMINE_BIOSYNTHETIC_PROCESS | 0,2048969 0,7744131 0,1051251 0,5160224 1,2248538 | 29 |
| GOBP_CHLORIDE_TRANSMEMBRANE_TRANSPORT | 0,1687817 0,7227339 0,1167392 0,4844523 1,2239825 | 46 |
| GOBP_ENDOCRINE_SYSTEM_DEVELOPMENT | 0,1254276 0,6464081 0,1301056 0,4413727 1,2239228 | 96 |
| GOBP_REGULATION_OF_ENDOTHELIAL_CELL_MIGRATION | 0,1182213 0,6334603 0,1307771 0,4223388 1,2234222 | 149 |
| GOBP_RESPONSE_TO_CHOLESTEROL | 0,2149533 0,7825382 0,104344 0,5499198 1,2229876 | 19 |
| GOBP_REGULATION_OF_SMOOTH_MUSCLE_CELL_DIFFERENTIATION | 0,2048969 0,7744131 0,1051251 0,5151439 1,2227685 | 29 |
| GOBP_REGULATION_OF_CENTRIOLE_REPLICATION | 0,2157124 0,7830255 0,1039585 0,5400508 1,2223967 | 22 |
| GOBP_T_CELL_LINEAGE_COMMITMENT | 0,2255034 0,7903135 0,1017139 0,5608854 1,2222092 | 17 |
| GOBP_NON_MEMBRANE_BOUNDED_ORGANELLE_ASSEMBLY | 0,0588843 0,4845135 0,1864326 0,4009085 1,2217438 | 353 |
| GOBP_REGULATION_OF_ANIMAL_ORGAN_FORMATION | 0,2210386 0,7889462 0,1024494 0,5387144 1,2193717 | 22 |
| GOBP_VASCULAR_TRANSPORT | 0,1458576 0,684728 0,1209851 0,4495808 1,2183682 | 76 |
| GOBP_CYCLIC_NUCLEOTIDE_BIOSYNTHETIC_PROCESS | 0,2362416 0,7988836 0,0988903 0,5587814 1,2176245 | 17 |
| GOBP_NEGATIVE_REGULATION_OF_FAT_CELL_DIFFERENTIATION | 0,178934 0,7387418 0,1128434 0,4819131 1,2175672 | 46 |
| GOBP_REGULATION_OF_RNA_SPLICING | 0,128149 0,6540073 0,1256399 0,4235671 1,2175524 | 140 |
| GOBP_NEGATIVE_REGULATION_OF_TRANSFERASE_ACTIVITY | 0,0958333 0,6059163 0,143759 0,4069904 1,2173704 | 240 |
| GOBP_ACETYL_COA_BIOSYNTHETIC_PROCESS | 0,2362416 0,7988836 0,0988903 0,5585532 1,2171272 | 17 |
| GOBP_ENDOCRINE_PANCREAS_DEVELOPMENT | 0,2030265 0,7744131 0,104344 0,4980314 1,2168694 | 35 |
| GOBP_TELENCEPHALON_DEVELOPMENT | 0,1079365 0,6209246 0,1357409 0,4097796 1,2163106 | 219 |
| GOBP_CORTICAL_ACTIN_CYTOSKELETON_ORGANIZATION | 0,1954603 0,7683417 0,1067299 0,4936269 1,2163018 | 37 |
| GOBP_PATTERN_SPECIFICATION_PROCESS | 0,0630165 0,5000396 0,1797823 0,3988957 1,2156097 | 353 |
| GOBP_METAL_ION_EXPORT | 0,1992434 0,7738707 0,1055209 0,4932023 1,2152556 | 37 |
| GOBP_NEGATIVE_REGULATION_OF_MUSCLE_CELL_DIFFERENTIATION | 0,1754601 0,7308245 0,1119183 0,4687131 1,2145415 | 53 |
| GOBP_NLS_BEARING_PROTEIN_IMPORT_INTO_NUCLEUS | 0,230563 0,7968953 0,1002791 0,5656338 1,214443 | 16 |
| GOBP_NEGATIVE_REGULATION_OF_MRNA_PROCESSING | 0,2126289 0,7803554 0,1028218 0,511569 1,214283 | 29 |
| GOBP_LEUKOCYTE_MEDIATED_IMMUNITY | 0,0993724 0,615061 0,1412251 0,4058633 1,2139787 | 246 |
| GOBP_RESPONSE_TO_COCAINE | 0,220681 0,7888973 0,0992333 0,4940115 1,21376 | 36 |
| GOBP_LABYRINTHINE_LAYER_BLOOD_VESSEL_DEVELOPMENT | 0,2358491 0,7988836 0,0992333 0,5501642 1,2137151 | 18 |
| GOBP_POSITIVE_REGULATION_OF_PROTEIN_KINASE_ACTIVITY | 0,0724638 0,5412925 0,1669338 0,4005823 1,213702 | 320 |
| GOBP_EAR_MORPHOGENESIS | 0,1338673 0,6600254 0,1256399 0,4397368 1,2136893 | 93 |
| GOBP_INFLAMMATORY_RESPONSE_TO_ANTIGENIC_STIMULUS | 0,195 0,7683417 0,1063233 0,4893991 1,2135795 | 39 |
| GOBP_SENSORY_PERCEPTION_OF_CHEMICAL_STIMULUS | 0,2002519 0,7744131 0,1051251 0,4908895 1,2128481 | 38 |
| GOBP_VENTRICULAR_SEPTUM_DEVELOPMENT | 0,1722488 0,7265238 0,1114627 0,4564829 1,2128087 | 64 |
| GOBP_PROTEIN_DEPOLYMERIZATION | 0,1414141 0,6791978 0,1204334 0,4304755 1,2125287 | 109 |
| GOBP_NEGATIVE_REGULATION_OF_RNA_SPLICING | 0,2276843 0,7916784 0,0988903 0,5197307 1,2111635 | 25 |
| GOBP_RESPONSE_TO_REACTIVE_OXYGEN_SPECIES | 0,1178378 0,6334603 0,1307771 0,4143438 1,2109509 | 168 |
| GOBP_ADAPTATION_OF_SIGNALING_PATHWAY | 0,2345845 0,7983096 0,0992333 0,563997 1,2109286 | 16 |
| GOBP_REGULATION_OF_PRESYNAPSE_ORGANIZATION | 0,2177835 0,7873562 0,1013507 0,5100783 1,2107445 | 29 |
| GOBP_EAR_DEVELOPMENT | 0,124197 0,6435974 0,126254 0,4123175 1,210686 | 173 |
| GOBP_REGULATION_OF_CELL_JUNCTION_ASSEMBLY | 0,1165775 0,6334603 0,1307771 0,4120797 1,2106583 | 174 |
| GOBP_NEGATIVE_REGULATION_OF_PROTEIN_SERINE_THREONINE_KINASE_ACTIVITY | 0,1293984 0,6544984 0,1275053 0,4340947 1,2102367 | 101 |
| GOBP_CELL_MATURATION | 0,1406423 0,6774271 0,1198878 0,4241135 1,2095825 | 124 |
| GOBP_MRNA_TRANSCRIPTION | 0,20125 0,7744131 0,104344 0,4877661 1,2095301 | 39 |
| GOBP_ASSOCIATIVE_LEARNING | 0,172209 0,7265238 0,1110115 0,4514252 1,2089358 | 70 |
| GOBP_BODY_MORPHOGENESIS | 0,193426 0,7681836 0,1075544 0,4827421 1,2087697 | 42 |
| GOBP_CELLULAR_RESPONSE_TO_ZINC_ION | 0,2469799 0,8075292 0,0962406 0,5543998 1,2080767 | 17 |
| GOBP_COGNITION | 0,0979167 0,6130256 0,1420566 0,4010056 1,2072785 | 264 |
| GOBP_ERK1_AND_ERK2_CASCADE | 0,1069182 0,6209246 0,1357409 0,4043337 1,2062651 | 236 |
| GOBP_NEGATIVE_REGULATION_OF_PEPTIDE_SECRETION | 0,2146465 0,7825382 0,1009906 0,499725 1,2061539 | 32 |
| GOBP_REGULATION_OF_SYNAPTIC_TRANSMISSION_GLUTAMATERGIC | 0,1796117 0,7399133 0,1096841 0,4593673 1,2061512 | 57 |
| GOBP_STRIATED_MUSCLE_CELL_APOPTOTIC_PROCESS | 0,2070707 0,7744131 0,1031975 0,4957403 1,206057 | 34 |
| GOBP_REGULATION_OF_CHROMATIN_ASSEMBLY_OR_DISASSEMBLY | 0,2185515 0,7879614 0,1002791 0,5042311 1,2059927 | 31 |
| GOBP_NEUROTRANSMITTER_SECRETION | 0,1463415 0,6855422 0,1172497 0,421654 1,2055115 | 128 |
| GOBP_ANIMAL_ORGAN_FORMATION | 0,1888889 0,7589899 0,1075544 0,4704138 1,2054806 | 49 |
| GOBP_DEFINITIVE_HEMOPOIESIS | 0,2496644 0,8103969 0,0956031 0,5530818 1,2052047 | 17 |
| GOBP_REGULATION_OF_LONG_TERM_NEURONAL_SYNAPTIC_PLASTICITY | 0,2421875 0,8030878 0,0956031 0,5233689 1,2051405 | 23 |
| GOBP_POSITIVE_REGULATION_OF_NEUROTRANSMITTER_SECRETION | 0,2371274 0,7998324 0,0992333 0,5713086 1,2050515 | 15 |
| GOBP_RESPIRATORY_SYSTEM_PROCESS | 0,209596 0,7750512 0,1024494 0,4953121 1,2050154 | 34 |
| GOBP_HISTONE_H2A_ACETYLATION | 0,2421875 0,8030878 0,0956031 0,5230235 1,2043451 | 23 |
| GOBP_REGULATION_OF_HISTONE_METHYLATION | 0,1793349 0,7395843 0,1083943 0,4494954 1,2037679 | 70 |
| GOBP_NEURON_FATE_SPECIFICATION | 0,2426273 0,8038368 0,0972151 0,56053 1,2034849 | 16 |
| GOBP_REGULATION_OF_PROTEIN_STABILITY | 0,0925156 0,5997765 0,1464162 0,3992599 1,2032949 | 271 |
| GOBP_PALLIUM_DEVELOPMENT | 0,1428571 0,6828426 0,1177658 0,4161436 1,2031787 | 145 |
| GOBP_REGULATION_OF_SYNAPTIC_PLASTICITY | 0,1287879 0,6540073 0,1244342 0,4115734 1,2028549 | 167 |
| GOBP_REGULATION_OF_HISTONE_H3_K9_METHYLATION | 0,2506667 0,8104601 0,0949752 0,5319393 1,2027991 | 21 |
| GOBP_POSITIVE_REGULATION_OF_HISTONE_DEACETYLATION | 0,2506667 0,8104601 0,0949752 0,5318799 1,2026646 | 21 |
| GOBP_NEGATIVE_REGULATION_OF_PROTEIN_LOCALIZATION_TO_NUCLEUS | 0,2298851 0,7963341 0,0975449 0,5048627 1,2023541 | 30 |
| GOBP_ANATOMICAL_STRUCTURE_MATURATION | 0,1304813 0,6552847 0,1226792 0,4092484 1,2023403 | 174 |
| GOBP_RESPONSE_TO_BMP | 0,1503268 0,6878113 0,1142665 0,416301 1,2018947 | 144 |
| GOBP_POSITIVE_REGULATION_OF_PHAGOCYTOSIS | 0,1956798 0,7683417 0,1071402 0,4776468 1,2011666 | 43 |
| GOBP_CELL_FATE_SPECIFICATION | 0,1917476 0,7671945 0,1055209 0,4568319 1,2004482 | 58 |
| GOBP_REGULATION_OF_PLATELET_DERIVED_GROWTH_FACTOR_RECEPTOR_SIGNALING_PATHWAY | 0,2520216 0,8117205 0,095288 0,5438492 1,1997835 | 18 |
| GOBP_INORGANIC_ANION_TRANSMEMBRANE_TRANSPORT | 0,1925837 0,7680255 0,104344 0,4508063 1,199521 | 66 |
| GOBP_MATING | 0,2252604 0,7901992 0,0999277 0,5086444 1,1990291 | 28 |
| GOBP_REGULATION_OF_TRANS_SYNAPTIC_SIGNALING | 0,0781893 0,5645811 0,1596467 0,39245 1,1988475 | 366 |
| GOBP_CELLULAR_RESPONSE_TO_TOXIC_SUBSTANCE | 0,1533181 0,6935605 0,1162341 0,4339637 1,1986021 | 92 |
| GOBP_POSITIVE_REGULATION_OF_AMINO_ACID_TRANSPORT | 0,246996 0,8075292 0,0959207 0,5387239 1,1980886 | 19 |
| GOBP_ACETYL_COA_METABOLIC_PROCESS | 0,2236341 0,7901992 0,0988903 0,5009186 1,1980701 | 31 |
| GOBP_POSTSYNAPTIC_MEMBRANE_ORGANIZATION | 0,2283563 0,7921835 0,0968878 0,4922912 1,1976187 | 33 |
| GOBP_CORONARY_VASCULATURE_MORPHOGENESIS | 0,2590604 0,8179127 0,0934449 0,5495195 1,1974421 | 17 |
| GOBP_POLYOL_CATABOLIC_PROCESS | 0,2335484 0,7983096 0,0972151 0,5101162 1,1974154 | 26 |
| GOBP_POSITIVE_REGULATION_OF_TRANSCRIPTION_FROM_RNA_POLYMERASE_II_PROMOTER_IN_RESPONSE_TO_STRESS | 0,2473958 0,8075292 0,0943564 0,5200121 1,197411 | 23 |
| GOBP_REGULATION_OF_PROTEIN_DEACETYLATION | 0,1865854 0,7553892 0,1075544 0,4580078 1,1973098 | 56 |
| GOBP_GERM_CELL_DEVELOPMENT | 0,1332623 0,6596394 0,1209851 0,4052057 1,1970033 | 207 |
| GOBP_SYNAPTIC_TRANSMISSION_GLUTAMATERGIC | 0,1622248 0,7128404 0,1133129 0,4378871 1,196991 | 80 |
| GOBP_MORPHOGENESIS_OF_A_POLARIZED_EPITHELIUM | 0,1554286 0,6960014 0,11524 0,4331245 1,1964908 | 89 |
| GOBP_FAT_SOLUBLE_VITAMIN_METABOLIC_PROCESS | 0,2348387 0,7984227 0,0968878 0,5096756 1,1963812 | 26 |
| GOBP_REGULATION_OF_ALTERNATIVE_MRNA_SPLICING_VIA_SPLICEOSOME | 0,1939024 0,7683417 0,1051251 0,4582391 1,1949453 | 55 |
| GOBP_NEGATIVE_REGULATION_OF_PLASMA_MEMBRANE_BOUNDED_CELL_PROJECTION_ASSEMBLY | 0,2319588 0,7982628 0,0975449 0,503333 1,1947335 | 29 |
| GOBP_MEMBRANE_REPOLARIZATION_DURING_CARDIAC_MUSCLE_CELL_ACTION_POTENTIAL | 0,2533512 0,8131684 0,0946646 0,5564487 1,1947221 | 16 |
| GOBP_POSITIVE_REGULATION_OF_POSTTRANSCRIPTIONAL_GENE_SILENCING | 0,2345361 0,7983096 0,0968878 0,5026297 1,1930641 | 29 |
| GOBP_LIPID_EXPORT_FROM_CELL | 0,2345361 0,7983096 0,0968878 0,5025965 1,1929854 | 29 |
| GOBP_OLIGOSACCHARIDE_METABOLIC_PROCESS | 0,2068528 0,7744131 0,1035763 0,4699575 1,1925806 | 47 |
| GOBP_PYRIMIDINE_CONTAINING_COMPOUND_BIOSYNTHETIC_PROCESS | 0,2321205 0,7982628 0,0959207 0,4900841 1,1922496 | 33 |
| GOBP_REGULATION_OF_PROTEIN_MODIFICATION_BY_SMALL_PROTEIN_CONJUGATION_OR_REMOVAL | 0,1300211 0,6552847 0,1221079 0,4005176 1,1919373 | 229 |
| GOBP_CHLORIDE_TRANSPORT | 0,1728972 0,7265238 0,1096841 0,4376629 1,1919368 | 78 |
| GOBP_AMINO_ACID_TRANSMEMBRANE_TRANSPORT | 0,1682135 0,7227339 0,1110115 0,436381 1,1913892 | 79 |
| GOBP_SENSORY_PERCEPTION_OF_SMELL | 0,2641509 0,8201778 0,0925529 0,539946 1,1911727 | 18 |
| GOBP_NEGATIVE_REGULATION_OF_DNA_BINDING | 0,1967005 0,7683417 0,1067299 0,4714488 1,1911288 | 46 |
| GOBP_RESPONSE_TO_ANGIOTENSIN | 0,252996 0,8130932 0,0943564 0,5262167 1,1910834 | 22 |
| GOBP_SEGMENT_SPECIFICATION | 0,2547425 0,8131684 0,0949752 0,5646072 1,1909166 | 15 |
| GOBP_REGULATION_OF_CHROMATIN_ORGANIZATION | 0,208805 0,7744131 0,1024494 0,4794796 1,1902992 | 40 |
| GOBP_RNA_DEPENDENT_DNA_BIOSYNTHETIC_PROCESS | 0,202864 0,7744131 0,1009906 0,444822 1,1890406 | 67 |
| GOBP_MRNA_TRANSPORT | 0,1768632 0,7342265 0,1051251 0,4192702 1,1881114 | 116 |
| GOBP_REGULATION_OF_CELL_AGING | 0,2279597 0,7916784 0,0972151 0,4808213 1,1879724 | 38 |
| GOBP_POSITIVE_REGULATION_OF_PROTEIN_DEACETYLATION | 0,2464516 0,8075292 0,0940503 0,5060603 1,1878948 | 26 |
| GOBP_RESPONSE_TO_EPIDERMAL_GROWTH_FACTOR | 0,2100629 0,7760141 0,1020801 0,469607 1,1872857 | 44 |
| GOBP_OVULATION | 0,2613941 0,8179308 0,0928481 0,5528874 1,1870758 | 16 |
| GOBP_RESPONSE_TO_ISCHEMIA | 0,2053571 0,7744131 0,104344 0,4708382 1,1867213 | 45 |
| GOBP_POSITIVE_REGULATION_OF_KINASE_ACTIVITY | 0,0841889 0,5803628 0,1531588 0,3866802 1,185662 | 386 |
| GOBP_IMPORT_ACROSS_PLASMA_MEMBRANE | 0,1721133 0,7265238 0,1055209 0,4106417 1,1855559 | 144 |
| GOBP_HISTONE_MONOUBIQUITINATION | 0,246134 0,8075292 0,0940503 0,4994123 1,1854272 | 29 |
| GOBP_SMALL_MOLECULE_CATABOLIC_PROCESS | 0,1018711 0,6192943 0,1388051 0,3922908 1,1854205 | 311 |
| GOBP_POSITIVE_REGULATION_OF_RNA_SPLICING | 0,2373737 0,7998324 0,0949752 0,4911033 1,1853445 | 32 |
| GOBP_NEGATIVE_REGULATION_OF_CELLULAR_RESPONSE_TO_GROWTH_FACTOR_STIMULUS | 0,1728538 0,7265238 0,10925 0,4341239 1,1852269 | 79 |
| GOBP_REGULATION_OF_MORPHOGENESIS_OF_A_BRANCHING_STRUCTURE | 0,225 0,7901992 0,0975449 0,4779161 1,1851048 | 39 |
| GOBP_POSITIVE_REGULATION_OF_CALCIUM_ION_TRANSMEMBRANE_TRANSPORT | 0,2050971 0,7744131 0,1013507 0,451245 1,1848246 | 57 |
| GOBP_POLYOL_BIOSYNTHETIC_PROCESS | 0,207362 0,7744131 0,1013507 0,4570263 1,1842583 | 53 |
| GOBP_RESPONSE_TO_KETONE | 0,1780673 0,7367829 0,1031975 0,4092462 1,1841606 | 148 |
| GOBP_DOUBLE_STRAND_BREAK_REPAIR_VIA_NONHOMOLOGOUS_END_JOINING | 0,2064439 0,7744131 0,0999277 0,4429763 1,184107 | 67 |
| GOBP_POSITIVE_REGULATION_OF_EXCITATORY_POSTSYNAPTIC_POTENTIAL | 0,2609854 0,8179308 0,0925529 0,522955 1,1837005 | 22 |
| GOBP_REGULATION_OF_MACROPHAGE_ACTIVATION | 0,22625 0,7916122 0,0972151 0,4772754 1,1835159 | 39 |
| GOBP_MRNA_PROCESSING | 0,0837589 0,5801999 0,1531588 0,3838025 1,1831272 | 433 |
| GOBP_GLAND_DEVELOPMENT | 0,1001032 0,6157146 0,1395997 0,3891411 1,1830291 | 338 |
| GOBP_CHAPERONE_MEDIATED_PROTEIN_COMPLEX_ASSEMBLY | 0,2765957 0,8313362 0,0891647 0,5264875 1,1829743 | 20 |
| GOBP_POSITIVE_REGULATION_OF_PEPTIDYL_THREONINE_PHOSPHORYLATION | 0,2529032 0,8130932 0,0925529 0,5039229 1,1828776 | 26 |
| GOBP_REGULATION_OF_CELLULAR_RESPONSE_TO_GROWTH_FACTOR_STIMULUS | 0,1294363 0,6544984 0,1215433 0,3954668 1,1826257 | 245 |
| GOBP_REGULATION_OF_ORGANELLE_ASSEMBLY | 0,1509033 0,6888745 0,1123785 0,3998587 1,1823617 | 212 |
| GOBP_PANCREAS_DEVELOPMENT | 0,2123786 0,7803554 0,0992333 0,4499356 1,1823261 | 58 |
| GOBP_OSSIFICATION | 0,1091658 0,6226089 0,1328463 0,388401 1,1821167 | 342 |
| GOBP_GLYCOSPHINGOLIPID_METABOLIC_PROCESS | 0,207362 0,7744131 0,1013507 0,4560905 1,1818333 | 53 |
| GOBP_L_AMINO_ACID_TRANSPORT | 0,2079807 0,7744131 0,1002791 0,449507 1,1813681 | 59 |
| GOBP_REGULATION_OF_BINDING | 0,1085832 0,6209246 0,133555 0,3896708 1,1809038 | 321 |
| GOBP_REGULATION_OF_VASCULATURE_DEVELOPMENT | 0,1492063 0,6878113 0,1128434 0,3977635 1,1806443 | 219 |
| GOBP_POSITIVE_REGULATION_OF_FATTY_ACID_METABOLIC_PROCESS | 0,2590674 0,8179127 0,0913924 0,5079439 1,1798636 | 24 |
| GOBP_REGULATION_OF_GLYCOGEN_METABOLIC_PROCESS | 0,2408854 0,8030226 0,0959207 0,5005044 1,1798405 | 28 |
| GOBP_LIMBIC_SYSTEM_DEVELOPMENT | 0,1822323 0,7458028 0,1047328 0,4260623 1,1795939 | 87 |
| GOBP_ESTABLISHMENT_OF_PROTEIN_LOCALIZATION_TO_CHROMOSOME | 0,2603627 0,8179308 0,0911073 0,507722 1,1793481 | 24 |
| GOBP_REGULATION_OF_CELLULAR_EXTRAVASATION | 0,2421875 0,8030878 0,0956031 0,500153 1,1790121 | 28 |
| GOBP_NEGATIVE_REGULATION_OF_MAP_KINASE_ACTIVITY | 0,2223618 0,7901992 0,09855 0,4629007 1,1788836 | 48 |
| GOBP_PIGMENT_BIOSYNTHETIC_PROCESS | 0,2129964 0,7803554 0,09855 0,4461852 1,1788057 | 62 |
| GOBP_HISTONE_MODIFICATION | 0,0873984 0,5907538 0,1492075 0,3820418 1,1787918 | 449 |
| GOBP_PROTEIN_AUTOPROCESSING | 0,2746667 0,8289344 0,0897105 0,5212026 1,1785217 | 21 |
| GOBP_IMPORT_INTO_NUCLEUS | 0,1842966 0,7501651 0,1013507 0,4075351 1,1782891 | 145 |
| GOBP_REGULATION_OF_TRANSMEMBRANE_RECEPTOR_PROTEIN_SERINE_THREONINE_KINASE_SIGNALING_PATHWAY | 0,1489362 0,6878113 0,1133129 0,3973762 1,1775647 | 217 |
| GOBP_NEGATIVE_REGULATION_OF_FIBROBLAST_PROLIFERATION | 0,2421875 0,8030878 0,0956031 0,5020625 1,177138 | 27 |
| GOBP_THIOESTER_BIOSYNTHETIC_PROCESS | 0,2275601 0,7916784 0,0975449 0,4700327 1,1769457 | 42 |
| GOBP_POSITIVE_REGULATION_OF_ORGANIC_ACID_TRANSPORT | 0,2496863 0,8103969 0,0916795 0,4837888 1,1769347 | 33 |
| GOBP_CALCIUM_DEPENDENT_CELL_CELL_ADHESION_VIA_PLASMA_MEMBRANE_CELL_ADHESION_MOLECULES | 0,2411616 0,8030878 0,0940503 0,4837185 1,17681 | 34 |
| GOBP_CELL_CELL_ADHESION_VIA_PLASMA_MEMBRANE_ADHESION_MOLECULES | 0,1492063 0,6878113 0,1128434 0,3952546 1,1762303 | 228 |
| GOBP_ANOIKIS | 0,2515883 0,8113453 0,0919686 0,4916492 1,1758999 | 31 |
| GOBP_RETINA_MORPHOGENESIS_IN_CAMERA_TYPE_EYE | 0,2181122 0,7873562 0,1006334 0,4665058 1,1758017 | 45 |
| GOBP_POSITIVE_REGULATION_OF_MESENCHYMAL_CELL_PROLIFERATION | 0,2857143 0,8386161 0,0880945 0,5328435 1,1755039 | 18 |
| GOBP_SECONDARY_METABOLIC_PROCESS | 0,2338812 0,7983096 0,0959207 0,4693861 1,1753267 | 42 |
| GOBP_NUCLEOSIDE_METABOLIC_PROCESS | 0,2324723 0,7983096 0,0946646 0,45609 1,1747984 | 51 |
| GOBP_REGULATION_OF_HISTONE_DEACETYLATION | 0,2301887 0,7963341 0,096563 0,4731266 1,174528 | 40 |
| GOBP_INNER_EAR_RECEPTOR_CELL_DEVELOPMENT | 0,2436869 0,8052169 0,0934449 0,4827407 1,1744311 | 34 |
| GOBP_REGULATION_OF_CELL_DEVELOPMENT | 0,099182 0,615061 0,1395997 0,3811794 1,1742434 | 426 |
| GOBP_RESPONSE_TO_DIETARY_EXCESS | 0,276 0,8303154 0,0894367 0,5188679 1,1732426 | 21 |
| GOBP_RESPONSE_TO_ALKALOID | 0,192757 0,7680255 0,1028218 0,4306924 1,1729532 | 78 |
| GOBP_POSITIVE_REGULATION_OF_SYNAPSE_ASSEMBLY | 0,2359413 0,7988836 0,0934449 0,4537602 1,17269 | 52 |
| GOBP_REGULATION_OF_LIPID_METABOLIC_PROCESS | 0,1459854 0,684728 0,1133129 0,3897578 1,1726213 | 258 |
| GOBP_CHAPERONE_MEDIATED_AUTOPHAGY | 0,2774799 0,8326613 0,0894367 0,5461431 1,1725954 | 16 |
| GOBP_SOMATIC_DIVERSIFICATION_OF_IMMUNOGLOBULINS | 0,226506 0,7916122 0,0949752 0,4443807 1,1722737 | 60 |
| GOBP_RESPONSE_TO_FOOD | 0,2786667 0,8335594 0,0888945 0,5183157 1,1719939 | 21 |
| GOBP_NUCLEOSIDE_PHOSPHATE_BIOSYNTHETIC_PROCESS | 0,1572795 0,7001174 0,1096841 0,3953518 1,1718666 | 215 |
| GOBP_MONOSACCHARIDE_CATABOLIC_PROCESS | 0,2449495 0,8075292 0,0931455 0,4814109 1,1711959 | 34 |
| GOBP_CALCIUM_ION_IMPORT_INTO_CYTOSOL | 0,2815013 0,8359478 0,0886261 0,5449179 1,1699649 | 16 |
| GOBP_ORGANIC_ACID_CATABOLIC_PROCESS | 0,1655983 0,7178712 0,1067299 0,3959788 1,1698398 | 205 |
| GOBP_POSITIVE_REGULATION_OF_AXONOGENESIS | 0,2193396 0,7888973 0,0956031 0,4347307 1,1696742 | 72 |
| GOBP_BIOLOGICAL_PROCESS_INVOLVED_IN_INTRASPECIES_INTERACTION_BETWEEN_ORGANISMS | 0,2322335 0,7982628 0,096563 0,4628674 1,1694477 | 46 |
| GOBP_VASCULAR_ENDOTHELIAL_GROWTH_FACTOR_PRODUCTION | 0,2681992 0,8262377 0,0886261 0,4908864 1,1690688 | 30 |
| GOBP_IMPORT_INTO_CELL | 0,1636364 0,7148626 0,1075544 0,3975379 1,1687049 | 186 |
| GOBP_REGULATION_OF_MRNA_METABOLIC_PROCESS | 0,144641 0,684728 0,1137873 0,3878864 1,168467 | 269 |
| GOBP_REGULATION_OF_MICROTUBULE_DEPOLYMERIZATION | 0,2554003 0,8131684 0,0911073 0,4885344 1,1684502 | 31 |
| GOBP_HEPARAN_SULFATE_PROTEOGLYCAN_METABOLIC_PROCESS | 0,2554003 0,8131684 0,0911073 0,4885007 1,1683696 | 31 |
| GOBP_REGULATION_OF_NUCLEAR_TRANSCRIBED_MRNA_CATABOLIC_PROCESS_DEADENYLATION_DEPENDENT_DECAY | 0,2733161 0,8289344 0,0883594 0,5024878 1,1671899 | 24 |
| GOBP_DNA_METHYLATION | 0,2472325 0,8075292 0,0911073 0,4529124 1,1666135 | 51 |
| GOBP_POSITIVE_REGULATION_OF_SODIUM_ION_TRANSMEMBRANE_TRANSPORT | 0,2911051 0,8451947 0,0870516 0,5286649 1,1662854 | 18 |
| GOBP_CELL_RECOGNITION | 0,1880631 0,7580835 0,1020801 0,4153469 1,1660468 | 105 |
| GOBP_EXPORT_ACROSS_PLASMA_MEMBRANE | 0,2395062 0,8012686 0,0931455 0,4535167 1,1657885 | 50 |
| GOBP_HISTONE_H3_K9_TRIMETHYLATION | 0,2882038 0,8422118 0,0873098 0,5428761 1,1655811 | 16 |
| GOBP_REGULATION_OF_TRANSCRIPTION_INVOLVED_IN_G1_S_TRANSITION_OF_MITOTIC_CELL_CYCLE | 0,2866667 0,8397027 0,0873098 0,5154538 1,1655228 | 21 |
| GOBP_POSITIVE_REGULATION_OF_CELL_CELL_ADHESION | 0,1641791 0,7155556 0,1071402 0,3942782 1,1652847 | 203 |
| GOBP_SULFUR_COMPOUND_BIOSYNTHETIC_PROCESS | 0,1937984 0,7683417 0,0992333 0,4074257 1,165268 | 133 |
| GOBP_INSULIN_SECRETION | 0,2013129 0,7744131 0,0962406 0,40485 1,1648967 | 141 |
| GOBP_L_ALPHA_AMINO_ACID_TRANSMEMBRANE_TRANSPORT | 0,2330097 0,7983096 0,0937465 0,443141 1,1644715 | 58 |
| GOBP_REGULATION_OF_LIPOPOLYSACCHARIDE_MEDIATED_SIGNALING_PATHWAY | 0,2924528 0,8455459 0,086795 0,5277091 1,164177 | 18 |
| GOBP_POSITIVE_REGULATION_OF_CELL_DIVISION | 0,2394705 0,8012686 0,0916795 0,440218 1,1637945 | 61 |
| GOBP_INORGANIC_ANION_TRANSPORT | 0,2006615 0,7744131 0,0968878 0,4063502 1,1633855 | 136 |
| GOBP_NEGATIVE_REGULATION_OF_HEMOPOIESIS | 0,2152941 0,7830203 0,096563 0,4307967 1,1633433 | 75 |
| GOBP_ORGANIC_HYDROXY_COMPOUND_METABOLIC_PROCESS | 0,1054248 0,6209246 0,135002 0,3779636 1,1631491 | 415 |
| GOBP_CHONDROITIN_SULFATE_PROTEOGLYCAN_METABOLIC_PROCESS | 0,2720307 0,8282165 0,0878313 0,488099 1,1624306 | 30 |
| GOBP_SOMATIC_DIVERSIFICATION_OF_IMMUNE_RECEPTORS | 0,2264151 0,7916122 0,0937465 0,4317133 1,1615555 | 72 |
| GOBP_MALE_GAMETE_GENERATION | 0,1148718 0,6309552 0,1287887 0,3784417 1,1614804 | 392 |
| GOBP_POSITIVE_REGULATION_OF_RESPONSE_TO_DNA_DAMAGE_STIMULUS | 0,2017738 0,7744131 0,0968878 0,407484 1,1613505 | 123 |
| GOBP_REGULATION_OF_PROTEIN_IMPORT | 0,2404908 0,8024191 0,0925529 0,4479568 1,1607572 | 53 |
| GOBP_POLARIZED_EPITHELIAL_CELL_DIFFERENTIATION | 0,2849741 0,8386161 0,0860347 0,4995784 1,160432 | 24 |
| GOBP_MORPHOGENESIS_OF_A_BRANCHING_STRUCTURE | 0,2071584 0,7744131 0,0940503 0,4003766 1,1598026 | 149 |
| GOBP_NEURAL_TUBE_DEVELOPMENT | 0,2030568 0,7744131 0,0956031 0,4033301 1,1596165 | 139 |
| GOBP_CGMP_MEDIATED_SIGNALING | 0,2825521 0,8376329 0,086795 0,5031859 1,1586658 | 23 |
| GOBP_EXPLORATION_BEHAVIOR | 0,286285 0,8396351 0,0873098 0,5117678 1,1583786 | 22 |
| GOBP_POSITIVE_REGULATION_OF_NEUROTRANSMITTER_TRANSPORT | 0,3018617 0,8521327 0,0843144 0,5151247 1,1574431 | 20 |
| GOBP_POSITIVE_REGULATION_OF_GLUCOSE_TRANSMEMBRANE_TRANSPORT | 0,2634881 0,8194741 0,0886261 0,4757539 1,1573878 | 33 |
| GOBP_ADAPTIVE_IMMUNE_RESPONSE | 0,1540062 0,6935605 0,1096841 0,3840168 1,156262 | 275 |
| GOBP_REGULATION_OF_T_CELL_ACTIVATION | 0,1712185 0,7265238 0,1035763 0,3877264 1,1561028 | 235 |
| GOBP_REGULATION_OF_CAMP_DEPENDENT_PROTEIN_KINASE_ACTIVITY | 0,3100671 0,8581768 0,0833634 0,5302351 1,1554202 | 17 |
| GOBP_PHAGOLYSOSOME_ASSEMBLY | 0,2962466 0,8470309 0,0857844 0,5379518 1,1550082 | 16 |
| GOBP_ZINC_ION_HOMEOSTASIS | 0,2954839 0,8468136 0,0838361 0,4920336 1,1549694 | 26 |
| GOBP_ORGAN_GROWTH | 0,2119205 0,7803554 0,0937465 0,4032594 1,1547993 | 135 |
| GOBP_REGULATION_OF_T_HELPER_1_TYPE_IMMUNE_RESPONSE | 0,3100671 0,8581768 0,0833634 0,52981 1,1544938 | 17 |
| GOBP_CELLULAR_CARBOHYDRATE_BIOSYNTHETIC_PROCESS | 0,2449223 0,8075292 0,0899861 0,4343899 1,1542567 | 65 |
| GOBP_B_CELL_ACTIVATION_INVOLVED_IN_IMMUNE_RESPONSE | 0,2452153 0,8075292 0,0899861 0,434352 1,1540099 | 64 |
| GOBP_RESPONSE_TO_IRON_ION | 0,2954839 0,8468136 0,0838361 0,491535 1,1537991 | 26 |
| GOBP_DIENCEPHALON_DEVELOPMENT | 0,2656827 0,8222156 0,0870516 0,4479199 1,1537537 | 51 |
| GOBP_METANEPHRIC_NEPHRON_DEVELOPMENT | 0,2689394 0,8262377 0,0878313 0,4779429 1,1535799 | 32 |
| GOBP_SODIUM_ION_IMPORT_ACROSS_PLASMA_MEMBRANE | 0,3018868 0,8521327 0,0850427 0,5228766 1,153516 | 18 |
| GOBP_ROOF_OF_MOUTH_DEVELOPMENT | 0,2227378 0,7901992 0,0937465 0,422368 1,1531315 | 79 |
| GOBP_NEGATIVE_REGULATION_OF_ENDOTHELIAL_CELL_PROLIFERATION | 0,2731893 0,8289344 0,0873098 0,4817585 1,152244 | 31 |
| GOBP_RESPONSE_TO_AMINE | 0,2686003 0,8262377 0,0878313 0,4687844 1,1517782 | 36 |
| GOBP_DEFENSE_RESPONSE_TO_GRAM_NEGATIVE_BACTERIUM | 0,2747804 0,8289344 0,0862866 0,4731199 1,15098 | 33 |
| GOBP_ODONTOGENESIS_OF_DENTIN_CONTAINING_TOOTH | 0,24821 0,8080806 0,0891647 0,4303897 1,1504621 | 67 |
| GOBP_POSITIVE_REGULATION_OF_OSTEOBLAST_DIFFERENTIATION | 0,2506024 0,8104601 0,0891647 0,4360641 1,1503347 | 60 |
| GOBP_POSITIVE_REGULATION_OF_LEUKOCYTE_CELL_CELL_ADHESION | 0,1928879 0,7680255 0,0978773 0,3924986 1,1499698 | 171 |
| GOBP_RESPONSE_TO_STEROL | 0,3035952 0,8550677 0,0840746 0,5080335 1,149926 | 22 |
| GOBP_REGULATION_OF_CARDIAC_MUSCLE_CELL_PROLIFERATION | 0,2760351 0,8303154 0,0860347 0,4725651 1,1496302 | 33 |
| GOBP_NEGATIVE_REGULATION_OF_IMMUNE_EFFECTOR_PROCESS | 0,2508961 0,8105043 0,0886261 0,4326336 1,1495898 | 65 |
| GOBP_PORE_COMPLEX_ASSEMBLY | 0,308 0,857788 0,0833634 0,5083907 1,1495519 | 21 |
| GOBP_AMINO_ACID_TRANSPORT | 0,2319645 0,7982628 0,0888945 0,4037655 1,1492715 | 120 |
| GOBP_PERICARDIUM_DEVELOPMENT | 0,3130081 0,8598592 0,0833634 0,5446124 1,1487418 | 15 |
| GOBP_REGULATION_OF_MEIOTIC_NUCLEAR_DIVISION | 0,3138298 0,8598592 0,0822055 0,5112307 1,1486936 | 20 |
| GOBP_TYPE_2_IMMUNE_RESPONSE | 0,3062583 0,8571368 0,0835991 0,5073254 1,1483233 | 22 |
| GOBP_REGULATION_OF_PROTEIN_DEPHOSPHORYLATION | 0,2252042 0,7901992 0,0934449 0,4231282 1,1480865 | 77 |
| GOBP_CARDIAC_SEPTUM_DEVELOPMENT | 0,2201835 0,7888973 0,0937465 0,415527 1,1480348 | 94 |
| GOBP_ACTIVATION_OF_PROTEIN_KINASE_ACTIVITY | 0,2331492 0,7983096 0,0883594 0,4034956 1,1472659 | 117 |
| GOBP_RNA_MEDIATED_GENE_SILENCING_BY_INHIBITION_OF_TRANSLATION | 0,3151596 0,8605047 0,0819779 0,5104429 1,1469235 | 20 |
| GOBP_REGULATION_OF_PEPTIDYL_THREONINE_PHOSPHORYLATION | 0,268262 0,8262377 0,0878313 0,4638962 1,1461551 | 38 |
| GOBP_POSITIVE_REGULATION_OF_NUCLEAR_DIVISION | 0,2629583 0,8194741 0,0891647 0,4576764 1,146006 | 42 |
| GOBP_POSITIVE_REGULATION_OF_HEART_GROWTH | 0,2955729 0,8468136 0,0843144 0,485899 1,1454112 | 28 |
| GOBP_GLUTAMINE_FAMILY_AMINO_ACID_CATABOLIC_PROCESS | 0,310253 0,8581768 0,0828962 0,5059607 1,1452342 | 22 |
| GOBP_NEGATIVE_REGULATION_OF_CHEMOKINE_PRODUCTION | 0,3140162 0,8598592 0,0828962 0,5191104 1,1452073 | 18 |
| GOBP_ALPHA_AMINO_ACID_METABOLIC_PROCESS | 0,2064865 0,7744131 0,0940503 0,3917459 1,144907 | 168 |
| GOBP_ESTABLISHMENT_OF_TISSUE_POLARITY | 0,2577566 0,8179127 0,0870516 0,428278 1,1448173 | 67 |
| GOBP_DNA_METHYLATION_OR_DEMETHYLATION | 0,2550656 0,8131684 0,0875697 0,428219 1,1446823 | 69 |
| GOBP_POSITIVE_REGULATION_OF_CALCIUM_ION_TRANSPORT | 0,2268041 0,7916784 0,0919686 0,4135903 1,1446434 | 90 |
| GOBP_PIGMENTATION | 0,2391304 0,8012686 0,0888945 0,4137136 1,1441262 | 88 |
| GOBP_RNA_SPLICING | 0,1469681 0,6876205 0,1119183 0,3736791 1,1438956 | 380 |
| GOBP_POSITIVE_REGULATION_OF_HISTONE_METHYLATION | 0,2691824 0,8262377 0,0875697 0,4524212 1,1438355 | 44 |
| GOBP_RESPONSE_TO_STEROID_HORMONE | 0,1826722 0,7462064 0,0992333 0,3799058 1,1433642 | 259 |
| GOBP_REGULATION_OF_PHOSPHOPROTEIN_PHOSPHATASE_ACTIVITY | 0,2703046 0,8264451 0,0878313 0,4503297 1,1427723 | 47 |
| GOBP_REGULATION_OF_MEGAKARYOCYTE_DIFFERENTIATION | 0,3115846 0,8582281 0,0826646 0,5047253 1,142438 | 22 |
| GOBP_REGULATION_OF_NERVOUS_SYSTEM_PROCESS | 0,2233446 0,7901992 0,0916795 0,4067741 1,1421282 | 104 |
| GOBP_TRABECULA_FORMATION | 0,3115846 0,8582281 0,0826646 0,504541 1,1420209 | 22 |
| GOBP_NEGATIVE_REGULATION_OF_STRIATED_MUSCLE_CELL_APOPTOTIC_PROCESS | 0,3164219 0,861089 0,0819779 0,5134857 1,1419605 | 19 |
| GOBP_TRANSMISSION_OF_NERVE_IMPULSE | 0,2743902 0,8289344 0,0847985 0,438413 1,141476 | 54 |
| GOBP_INTESTINAL_EPITHELIAL_CELL_DIFFERENTIATION | 0,3252033 0,8660673 0,0813027 0,5410964 1,1413255 | 15 |
| GOBP_OLEFINIC_COMPOUND_METABOLIC_PROCESS | 0,2315551 0,7982628 0,0902635 0,4092828 1,1410622 | 101 |
| GOBP_NEGATIVE_REGULATION_OF_LIPID_METABOLIC_PROCESS | 0,237819 0,8006148 0,0899861 0,4179401 1,1410426 | 79 |
| GOBP_NEGATIVE_REGULATION_OF_NOTCH_SIGNALING_PATHWAY | 0,2777778 0,83289 0,0860347 0,4690163 1,1410418 | 34 |
| GOBP_NEGATIVE_REGULATION_OF_REPRODUCTIVE_PROCESS | 0,2754717 0,8299496 0,0862866 0,4595867 1,1409154 | 40 |
| GOBP_ENERGY_RESERVE_METABOLIC_PROCESS | 0,2630952 0,8194741 0,0857844 0,4259115 1,1406313 | 68 |
| GOBP_NEGATIVE_REGULATION_OF_DNA_BIOSYNTHETIC_PROCESS | 0,2840909 0,8386161 0,0847985 0,4724241 1,1402595 | 32 |
| GOBP_ALCOHOL_METABOLIC_PROCESS | 0,18125 0,743854 0,0995791 0,3787756 1,1399804 | 277 |
| GOBP_CEREBRAL_CORTEX_CELL_MIGRATION | 0,279597 0,8352792 0,0855357 0,4613347 1,1398265 | 38 |
| GOBP_OVARIAN_FOLLICLE_DEVELOPMENT | 0,2716049 0,8282165 0,0860347 0,4433727 1,1397128 | 50 |
| GOBP_ESTABLISHMENT_OR_MAINTENANCE_OF_CELL_POLARITY | 0,2036247 0,7744131 0,0940503 0,3861164 1,1394737 | 198 |
| GOBP_OOCYTE_DIFFERENTIATION | 0,2704403 0,8264451 0,0873098 0,4504435 1,1388354 | 44 |
| GOBP_CELLULAR_HORMONE_METABOLIC_PROCESS | 0,2394044 0,8012686 0,0888945 0,4123702 1,1379471 | 85 |
| GOBP_HISTONE_H4_K5_ACETYLATION | 0,3217158 0,8660673 0,0813027 0,5296067 1,1370911 | 16 |
| GOBP_NUCLEOSIDE_MONOPHOSPHATE_METABOLIC_PROCESS | 0,2588652 0,8179127 0,0862866 0,4233903 1,1370381 | 71 |
| GOBP_REGULATION_OF_NEUROTRANSMITTER_UPTAKE | 0,3292683 0,8660673 0,0806388 0,538717 1,1363068 | 15 |

| GOBP_POSITIVE_REGULATION_OF_OLIGODENDROCYTE_DIFFERENTIATION | 0,3257979 0,8660673 0,0802023 0,5054967 1,1358097 | 20 |
| --- | --- | --- |
| GOBP_TELENCEPHALON_GLIAL_CELL_MIGRATION | 0,3133333 0,8598592 0,0824344 0,5022184 1,1355953 | 21 |
| GOBP_PROTEIN_COMPLEX_OLIGOMERIZATION | 0,2114772 0,7803554 0,0916795 0,3842581 1,1347881 | 197 |
| GOBP_ENTEROENDOCRINE_CELL_DIFFERENTIATION | 0,3148387 0,8602538 0,08042 0,4831554 1,1341293 | 26 |
| GOBP_SENSORY_PERCEPTION | 0,1671827 0,7211312 0,1039585 0,3723478 1,1338434 | 349 |
| GOBP_CELL_FATE_COMMITMENT | 0,2235294 0,7901992 0,0888945 0,3859035 1,1337547 | 174 |
| GOBP_MONOCARBOXYLIC_ACID_TRANSPORT | 0,2707838 0,8264451 0,0840746 0,4233414 1,1337262 | 70 |
| GOBP_POSITIVE_REGULATION_OF_CELL_JUNCTION_ASSEMBLY | 0,2540046 0,8131684 0,0855357 0,4099311 1,1336657 | 88 |
| GOBP_CELL_AGING | 0,2434585 0,8052169 0,0875697 0,4068038 1,133512 | 100 |
| GOBP_GLUTAMINE_METABOLIC_PROCESS | 0,3182423 0,8622448 0,0815265 0,5007357 1,1334076 | 22 |
| GOBP_GENE_SILENCING_BY_RNA | 0,2599558 0,8179308 0,0824344 0,3980352 1,1320876 | 119 |
| GOBP_POSITIVE_REGULATION_OF_TRANSCRIPTION_REGULATORY_REGION_DNA_BINDING | 0,3274933 0,8660673 0,0806388 0,5131316 1,1320174 | 18 |
| GOBP_POSITIVE_REGULATION_OF_MITOTIC_NUCLEAR_DIVISION | 0,293601 0,8459041 0,0826646 0,465257 1,1318514 | 33 |
| GOBP_SIGNAL_TRANSDUCTION_BY_P53_CLASS_MEDIATOR | 0,2361863 0,7988836 0,08654 0,3886334 1,131836 | 159 |
| GOBP_POSITIVE_REGULATION_OF_BINDING | 0,2548807 0,8131684 0,0824344 0,3902355 1,1304259 | 149 |
| GOBP_CELLULAR_RESPONSE_TO_PROSTAGLANDIN_STIMULUS | 0,3297587 0,8660673 0,0799859 0,526295 1,1299806 | 16 |
| GOBP_REGULATION_OF_PROTEIN_BINDING | 0,2274678 0,7916784 0,0880945 0,3850974 1,1297769 | 172 |
| GOBP_COLUMNAR_CUBOIDAL_EPITHELIAL_CELL_DEVELOPMENT | 0,2884131 0,8422118 0,0838361 0,4572492 1,1297325 | 38 |
| GOBP_POSTSYNAPTIC_SPECIALIZATION_ORGANIZATION | 0,3131443 0,8598592 0,0806388 0,4758878 1,1295883 | 29 |
| GOBP_SCF_DEPENDENT_PROTEASOMAL_UBIQUITIN_DEPENDENT_PROTEIN_CATABOLIC_PROCESS | 0,286802 0,8397027 0,0845557 0,446913 1,1291385 | 46 |
| GOBP_PIGMENT_METABOLIC_PROCESS | 0,2647754 0,8207601 0,0850427 0,4194216 1,1290917 | 73 |
| GOBP_REGULATION_OF_MYELINATION | 0,2842767 0,8386161 0,0845557 0,4547986 1,1290291 | 40 |
| GOBP_RESPONSE_TO_PURINE_CONTAINING_COMPOUND | 0,2491582 0,8103969 0,0855357 0,4021062 1,1290218 | 104 |
| GOBP_NEGATIVE_REGULATION_OF_MYOBLAST_DIFFERENTIATION | 0,337766 0,8693282 0,0782955 0,5024546 1,1289745 | 20 |
| GOBP_HISTONE_METHYLATION | 0,2456332 0,8075292 0,0847985 0,3925172 1,1285282 | 139 |
| GOBP_REGULATION_OF_RECEPTOR_SIGNALING_PATHWAY_VIA_STAT | 0,2843489 0,8386161 0,0817516 0,4246608 1,1284046 | 65 |
| GOBP_POSITIVE_REGULATION_OF_LIPID_METABOLIC_PROCESS | 0,25 0,8103969 0,0855357 0,4016619 1,1282292 | 106 |
| GOBP_CENTRAL_NERVOUS_SYSTEM_NEURON_DEVELOPMENT | 0,2671395 0,8260436 0,0845557 0,4190947 1,1282118 | 73 |
| GOBP_LAMELLIPODIUM_MORPHOGENESIS | 0,3297587 0,8660673 0,0799859 0,5253406 1,1279314 | 16 |
| GOBP_GLUCAN_BIOSYNTHETIC_PROCESS | 0,2934509 0,8459041 0,0828962 0,4561274 1,1269607 | 38 |
| GOBP_POSITIVE_REGULATION_OF_BIOMINERALIZATION | 0,2893401 0,8430641 0,0840746 0,445968 1,1267509 | 46 |
| GOBP_PHOSPHATIDYLINOSITOL_PHOSPHATE_BIOSYNTHETIC_PROCESS | 0,290012 0,8439777 0,0810802 0,4264516 1,1266702 | 62 |
| GOBP_SEX_DETERMINATION | 0,3324397 0,8660673 0,0795565 0,5243844 1,1258785 | 16 |
| GOBP_POSTSYNAPSE_ASSEMBLY | 0,3242188 0,8660673 0,0793435 0,4801767 1,1258243 | 27 |
| GOBP_REGULATION_OF_ORGANIC_ACID_TRANSPORT | 0,2885086 0,8422118 0,0822055 0,4355598 1,1256534 | 52 |
| GOBP_LEUKOCYTE_HOMEOSTASIS | 0,2783019 0,8331316 0,0822055 0,4183662 1,1256442 | 72 |
| GOBP_PROTEIN_LOCALIZATION_TO_MICROTUBULE_ORGANIZING_CENTER | 0,2992424 0,8492127 0,0819779 0,4625872 1,125401 | 34 |
| GOBP_AORTA_MORPHOGENESIS | 0,3242188 0,8660673 0,0793435 0,4799948 1,125398 | 27 |
| GOBP_TYPE_B_PANCREATIC_CELL_DEVELOPMENT | 0,336 0,8693282 0,0787114 0,4975986 1,1251493 | 21 |
| GOBP_RECEPTOR_CLUSTERING | 0,291358 0,8452758 0,0822055 0,4375705 1,124798 | 50 |
| GOBP_CEREBRAL_CORTEX_DEVELOPMENT | 0,2599319 0,8179308 0,0838361 0,4034317 1,1247497 | 101 |
| GOBP_LEUKOTRIENE_METABOLIC_PROCESS | 0,3364611 0,8693282 0,078921 0,5235113 1,1240038 | 16 |
| GOBP_HINDBRAIN_DEVELOPMENT | 0,2613511 0,8179308 0,0822055 0,3940978 1,123977 | 124 |
| GOBP_BILE_ACID_BIOSYNTHETIC_PROCESS | 0,3255208 0,8660673 0,0791317 0,4793781 1,1239519 | 27 |
| GOBP_CELLULAR_COMPONENT_DISASSEMBLY_INVOLVED_IN_EXECUTION_PHASE_OF_APOPTOSIS | 0,3225389 0,8660673 0,0793435 0,4838646 1,1239316 | 24 |
| GOBP_EMBRYONIC_AXIS_SPECIFICATION | 0,3255208 0,8660673 0,0791317 0,4793649 1,1239212 | 27 |
| GOBP_NEGATIVE_REGULATION_OF_EXTRINSIC_APOPTOTIC_SIGNALING_PATHWAY_VIA_DEATH_DOMAIN_RECEPTORS | 0,3225389 0,8660673 0,0793435 0,4838284 1,1238473 | 24 |
| GOBP_N_TERMINAL_PROTEIN_AMINO_ACID_ACETYLATION | 0,3387534 0,8693282 0,0791317 0,5327377 1,1236946 | 15 |
| GOBP_POSITIVE_REGULATION_OF_SYNAPTIC_TRANSMISSION | 0,2654867 0,8222156 0,0813027 0,3922707 1,1234354 | 130 |
| GOBP_POSITIVE_REGULATION_OF_VIRAL_LIFE_CYCLE | 0,3328841 0,8660673 0,0797706 0,5091483 1,12323 | 18 |
| GOBP_POSITIVE_REGULATION_OF_ANIMAL_ORGAN_MORPHOGENESIS | 0,3255208 0,8660673 0,0791317 0,4790097 1,1230882 | 27 |
| GOBP_ADENYLATE_CYCLASE_INHIBITING_G_PROTEIN_COUPLED_RECEPTOR_SIGNALING_PATHWAY | 0,2956853 0,8468136 0,0828962 0,4425691 1,1230786 | 47 |
| GOBP_PROTEIN_LIPID_COMPLEX_ASSEMBLY | 0,3324433 0,8660673 0,0793435 0,5049802 1,1230448 | 19 |
| GOBP_REGULATION_OF_FATTY_ACID_METABOLIC_PROCESS | 0,2990431 0,8492127 0,0791317 0,4225766 1,1227246 | 64 |
| GOBP_BRANCHING_INVOLVED_IN_MAMMARY_GLAND_DUCT_MORPHOGENESIS | 0,3328841 0,8660673 0,0797706 0,5089048 1,1226928 | 18 |
| GOBP_MEMBRANE_REPOLARIZATION_DURING_ACTION_POTENTIAL | 0,3444149 0,8738784 0,0772747 0,4995794 1,1225142 | 20 |
| GOBP_ARACHIDONIC_ACID_METABOLIC_PROCESS | 0,3113088 0,8582281 0,0802023 0,4692828 1,1224052 | 31 |
| GOBP_RESPONSE_TO_INORGANIC_SUBSTANCE | 0,1820041 0,7456809 0,0982123 0,3641504 1,1221187 | 431 |
| GOBP_CARDIAC_MUSCLE_CELL_PROLIFERATION | 0,2869785 0,8397027 0,0843144 0,4481193 1,1220753 | 42 |
| GOBP_STEROL_METABOLIC_PROCESS | 0,2633333 0,8194741 0,0819779 0,3958798 1,1213794 | 115 |
| GOBP_PLATELET_AGGREGATION | 0,3060976 0,8571368 0,078921 0,429767 1,120699 | 55 |
| GOBP_MEMORY | 0,2719101 0,8282165 0,0808589 0,3979274 1,120091 | 108 |
| GOBP_RNA_PHOSPHODIESTER_BOND_HYDROLYSIS_ENDONUCLEOLYTIC | 0,2943981 0,8468136 0,0797706 0,4189787 1,1199819 | 69 |
| GOBP_NEURAL_TUBE_FORMATION | 0,2582857 0,8179127 0,0845557 0,4039443 1,1199737 | 97 |
| GOBP_PHOSPHOLIPID_DEPHOSPHORYLATION | 0,2986023 0,8492127 0,0824344 0,4452678 1,1197411 | 43 |
| GOBP_REGULATION_OF_CILIUM_ASSEMBLY | 0,2749141 0,8289344 0,0813027 0,4061317 1,1195999 | 84 |
| GOBP_POTASSIUM_ION_TRANSPORT | 0,2481283 0,8080806 0,0831291 0,3809599 1,1193447 | 175 |
| GOBP_ACTIVATION_OF_PHOSPHOLIPASE_C_ACTIVITY | 0,3351135 0,8693282 0,078921 0,5030814 1,1188218 | 19 |
| GOBP_POSITIVE_REGULATION_OF_BEHAVIOR | 0,3455285 0,8738784 0,0780892 0,5298958 1,1177002 | 15 |
| GOBP_POTASSIUM_ION_HOMEOSTASIS | 0,3294271 0,8660673 0,0785029 0,4851284 1,1170857 | 23 |
| GOBP_SENSORY_PERCEPTION_OF_MECHANICAL_STIMULUS | 0,2739274 0,8289344 0,0793435 0,3898607 1,1170595 | 137 |
| GOBP_CELLULAR_AMINO_ACID_METABOLIC_PROCESS | 0,2181628 0,7873562 0,0888945 0,373483 1,1168215 | 241 |
| GOBP_NEGATIVE_REGULATION_OF_STRIATED_MUSCLE_CELL_DIFFERENTIATION | 0,3269476 0,8660673 0,077884 0,4688697 1,116635 | 30 |
| GOBP_MYELIN_MAINTENANCE | 0,3503356 0,8764774 0,0768737 0,512329 1,1164013 | 17 |
| GOBP_ORGAN_INDUCTION | 0,3396226 0,8706299 0,0787114 0,5060229 1,1163351 | 18 |
| GOBP_ANTERIOR_POSTERIOR_AXIS_SPECIFICATION | 0,3102144 0,8581768 0,0799859 0,4543019 1,1161956 | 36 |
| GOBP_REGULATION_OF_CELL_ACTIVATION | 0,1956967 0,7683417 0,0940503 0,3627516 1,1160951 | 422 |
| GOBP_TRANSMEMBRANE_RECEPTOR_PROTEIN_SERINE_THREONINE_KINASE_SIGNALING_PATHWAY | 0,2087683 0,7744131 0,0913924 0,3698867 1,1156168 | 300 |
| GOBP_CORONARY_VASCULATURE_DEVELOPMENT | 0,3024142 0,8529807 0,0817516 0,4435863 1,1155125 | 43 |
| GOBP_REGULATION_OF_CELLULAR_KETONE_METABOLIC_PROCESS | 0,2639363 0,8201778 0,0831291 0,4002819 1,1153396 | 100 |
| GOBP_REGULATION_OF_B_CELL_APOPTOTIC_PROCESS | 0,349866 0,8764774 0,0768737 0,5191735 1,1146903 | 16 |
| GOBP_POSITIVE_REGULATION_OF_CELL_PROJECTION_ORGANIZATION | 0,20625 0,7744131 0,0919686 0,3690712 1,1143083 | 309 |
| GOBP_NEGATIVE_REGULATION_OF_COLD_INDUCED_THERMOGENESIS | 0,3144654 0,8598592 0,0791317 0,4405616 1,1138516 | 44 |
| GOBP_TRABECULA_MORPHOGENESIS | 0,295082 0,8468136 0,0826646 0,4464965 1,1137014 | 41 |
| GOBP_NEGATIVE_REGULATION_OF_WNT_SIGNALING_PATHWAY | 0,2816901 0,8359478 0,0770737 0,3842072 1,1135216 | 150 |
| GOBP_SPERMATID_DIFFERENTIATION | 0,2852386 0,8386161 0,0776799 0,3910259 1,1130097 | 120 |
| GOBP_RNA_LOCALIZATION | 0,2625938 0,8194741 0,0802023 0,3787851 1,112948 | 181 |
| GOBP_PLASMA_MEMBRANE_PHOSPHOLIPID_SCRAMBLING | 0,3409704 0,8714711 0,0785029 0,5044418 1,1128469 | 18 |
| GOBP_RESPONSE_TO_INTERFERON_BETA | 0,3510363 0,877063 0,0749279 0,4787853 1,1121332 | 24 |
| GOBP_CEREBELLAR_CORTEX_MORPHOGENESIS | 0,3278272 0,8660673 0,0774768 0,4649868 1,1121304 | 31 |
| GOBP_NEGATIVE_REGULATION_OF_GLUCOSE_TRANSMEMBRANE_TRANSPORT | 0,3525469 0,877063 0,0764767 0,5179715 1,1121097 | 16 |
| GOBP_SKELETAL_SYSTEM_MORPHOGENESIS | 0,2620321 0,8190138 0,0802023 0,3785121 1,1120394 | 174 |
| GOBP_SENSORY_ORGAN_DEVELOPMENT | 0,196738 0,7683417 0,0934449 0,3605719 1,1118325 | 446 |
| GOBP_REGULATION_OF_CELL_CELL_ADHESION | 0,2132505 0,7803554 0,0897105 0,366926 1,1117285 | 320 |
| GOBP_REGULATION_OF_POSITIVE_CHEMOTAXIS | 0,3590426 0,877063 0,0751182 0,4947312 1,1116207 | 20 |
| GOBP_GLYCOSPHINGOLIPID_BIOSYNTHETIC_PROCESS | 0,3307791 0,8660673 0,0772747 0,4664275 1,1108188 | 30 |
| GOBP_PYRIMIDINE_RIBONUCLEOTIDE_BIOSYNTHETIC_PROCESS | 0,3590426 0,877063 0,0751182 0,494349 1,1107619 | 20 |
| GOBP_RESPONSE_TO_ALCOHOL | 0,2633833 0,8194741 0,0799859 0,3777377 1,110283 | 176 |
| GOBP_NEGATIVE_REGULATION_OF_INTRACELLULAR_PROTEIN_TRANSPORT | 0,31125 0,8582281 0,0793435 0,44752 1,1097304 | 39 |
| GOBP_POSITIVE_REGULATION_OF_CELLULAR_CARBOHYDRATE_METABOLIC_PROCESS | 0,318239 0,8622448 0,0785029 0,4387818 1,1093518 | 44 |
| GOBP_STARTLE_RESPONSE | 0,3590426 0,877063 0,0751182 0,4935905 1,1090575 | 20 |
| GOBP_PROTEIN_AUTOPHOSPHORYLATION | 0,2707889 0,8264451 0,0782955 0,375077 1,1084504 | 201 |
| GOBP_NUCLEOTIDE_SUGAR_METABOLIC_PROCESS | 0,3282828 0,8660673 0,0770737 0,4592197 1,1083891 | 32 |
| GOBP_NEGATIVE_REGULATION_OF_SMOOTH_MUSCLE_CELL_DIFFERENTIATION | 0,357047 0,877063 0,0758887 0,5086448 1,1083733 | 17 |
| GOBP_POSITIVE_REGULATION_OF_NITRIC_OXIDE_METABOLIC_PROCESS | 0,3324742 0,8660673 0,0774768 0,4667536 1,107907 | 29 |
| GOBP_ESTABLISHMENT_OF_RNA_LOCALIZATION | 0,3006536 0,8499289 0,0739901 0,3829426 1,1077141 | 147 |
| GOBP_LIPOSACCHARIDE_METABOLIC_PROCESS | 0,2894737 0,8430641 0,0785029 0,4012393 1,107435 | 93 |
| GOBP_G_PROTEIN_COUPLED_RECEPTOR_SIGNALING_PATHWAY | 0,1993896 0,7738707 0,0925529 0,3581884 1,1068716 | 466 |
| GOBP_KIDNEY_MESENCHYME_DEVELOPMENT | 0,3605898 0,8785577 0,0753094 0,5155185 1,106843 | 16 |
| GOBP_ACUTE_PHASE_RESPONSE | 0,3586667 0,877063 0,0753094 0,4894663 1,106761 | 21 |
| GOBP_RETINAL_GANGLION_CELL_AXON_GUIDANCE | 0,3504043 0,8764774 0,0770737 0,5015668 1,1065045 | 18 |
| GOBP_LONG_CHAIN_FATTY_ACYL_COA_BIOSYNTHETIC_PROCESS | 0,351752 0,877063 0,0768737 0,5014964 1,1063492 | 18 |
| GOBP_NEURONAL_ACTION_POTENTIAL | 0,3329098 0,8660673 0,0766747 0,4624739 1,10612 | 31 |
| GOBP_FATTY_ACID_DERIVATIVE_BIOSYNTHETIC_PROCESS | 0,3203026 0,8657389 0,0782955 0,4526392 1,10596 | 35 |
| GOBP_SOMATIC_DIVERSIFICATION_OF_IMMUNOGLOBULINS_INVOLVED_IN_IMMUNE_RESPONSE | 0,3245283 0,8660673 0,0774768 0,4371841 1,1053124 | 44 |
| GOBP_CELLULAR_RESPONSE_TO_INTERLEUKIN_1 | 0,3087886 0,8581768 0,0770737 0,4126038 1,1049705 | 70 |
| GOBP_POSITIVE_T_CELL_SELECTION | 0,3661784 0,8810414 0,0741759 0,4880437 1,1046793 | 22 |
| GOBP_LUNG_EPITHELIUM_DEVELOPMENT | 0,3450521 0,8738784 0,0760837 0,4711344 1,1046238 | 27 |
| GOBP_MUSCLE_CELL_APOPTOTIC_PROCESS | 0,3192261 0,8634503 0,0762797 0,4203045 1,1046198 | 59 |
| GOBP_OUTFLOW_TRACT_MORPHOGENESIS | 0,3169856 0,861733 0,0760837 0,4151252 1,1045795 | 66 |
| GOBP_PROTEIN_PROCESSING | 0,2780749 0,8331158 0,0770737 0,3754567 1,1035404 | 185 |
| GOBP_NEGATIVE_REGULATION_OF_PROTEIN_DEPHOSPHORYLATION | 0,3427835 0,8738784 0,0758887 0,4649113 1,1035342 | 29 |
| GOBP_POSITIVE_REGULATION_OF_SMOOTH_MUSCLE_CELL_PROLIFERATION | 0,325 0,8660673 0,0745501 0,4120444 1,1034938 | 68 |
| GOBP_CELLULAR_OXIDANT_DETOXIFICATION | 0,2990544 0,8492127 0,0785029 0,4098627 1,1033589 | 73 |
| GOBP_ELECTRON_TRANSPORT_CHAIN | 0,2935378 0,8459041 0,0755015 0,3838276 1,1033204 | 140 |
| GOBP_PROTEIN_METHYLATION | 0,2805139 0,8352792 0,0766747 0,3752288 1,103239 | 177 |
| GOBP_DOSAGE_COMPENSATION | 0,3497997 0,8764774 0,0766747 0,4960732 1,1032361 | 19 |
| GOBP_NEGATIVE_REGULATION_OF_CELL_ACTIVATION | 0,2976847 0,8492037 0,0751182 0,3852124 1,1028678 | 136 |
| GOBP_VENTRICULAR_CARDIAC_MUSCLE_CELL_MEMBRANE_REPOLARIZATION | 0,3643617 0,880326 0,0743625 0,490489 1,1020886 | 20 |
| GOBP_HISTONE_H2A_MONOUBIQUITINATION | 0,3610738 0,8785974 0,0753094 0,5056003 1,1017391 | 17 |
| GOBP_NEGATIVE_REGULATION_OF_EPITHELIAL_CELL_PROLIFERATION | 0,3063973 0,8571368 0,0745501 0,392213 1,101244 | 104 |
| GOBP_PYRIMIDINE_NUCLEOSIDE_TRIPHOSPHATE_BIOSYNTHETIC_PROCESS | 0,3625337 0,8797754 0,0753094 0,4991218 1,1011106 | 18 |
| GOBP_MYELOID_CELL_HOMEOSTASIS | 0,3082873 0,8579521 0,0734381 0,3846095 1,1007177 | 131 |
| GOBP_NEGATIVE_REGULATION_OF_CELLULAR_MACROMOLECULE_BIOSYNTHETIC_PROCESS | 0,2546973 0,8131684 0,08042 0,3678857 1,100084 | 241 |
| GOBP_RESPIRATORY_GASEOUS_EXCHANGE_BY_RESPIRATORY_SYSTEM | 0,3317073 0,8660673 0,0747385 0,4218439 1,1000379 | 55 |
| GOBP_REGULATION_OF_EXTRINSIC_APOPTOTIC_SIGNALING_PATHWAY_VIA_DEATH_DOMAIN_RECEPTORS | 0,3165195 0,861089 0,078921 0,4410129 1,1000236 | 41 |
| GOBP_CIRCADIAN_RHYTHM | 0,3006466 0,8499289 0,0734381 0,3756331 1,1000152 | 170 |
| GOBP_SECONDARY_METABOLITE_BIOSYNTHETIC_PROCESS | 0,3693333 0,8814579 0,0738053 0,4864165 1,0998649 | 21 |
| GOBP_CELLULAR_MONOVALENT_INORGANIC_CATION_HOMEOSTASIS | 0,3005714 0,8499289 0,0764767 0,3972995 1,0998316 | 91 |
| GOBP_HEMATOPOIETIC_STEM_CELL_HOMEOSTASIS | 0,3686327 0,8814579 0,0741759 0,5121211 1,0995486 | 16 |
| GOBP_NEGATIVE_REGULATION_OF_NERVOUS_SYSTEM_DEVELOPMENT | 0,3070953 0,8571747 0,0738053 0,3847617 1,0995483 | 127 |
| GOBP_REGULATION_OF_ESTABLISHMENT_OF_PLANAR_POLARITY | 0,3276074 0,8660673 0,0756946 0,4241762 1,0991363 | 53 |
| GOBP_AMYLOID_PRECURSOR_PROTEIN_METABOLIC_PROCESS | 0,3309524 0,8660673 0,0736213 0,4102362 1,0986513 | 68 |
| GOBP_LYMPHOCYTE_ACTIVATION_INVOLVED_IN_IMMUNE_RESPONSE | 0,307947 0,857788 0,0734381 0,3836338 1,0985983 | 135 |
| GOBP_RESPONSE_TO_LEUKEMIA_INHIBITORY_FACTOR | 0,3104238 0,8581768 0,0749279 0,3978882 1,0979836 | 85 |
| GOBP_B_CELL_DIFFERENTIATION | 0,3062857 0,8571368 0,0755015 0,3966271 1,0979702 | 91 |
| GOBP_LOW_DENSITY_LIPOPROTEIN_PARTICLE_CLEARANCE | 0,3564753 0,877063 0,0756946 0,4934638 1,097433 | 19 |
| GOBP_NEGATIVE_REGULATION_OF_NEURAL_PRECURSOR_CELL_PROLIFERATION | 0,3683511 0,8814579 0,0738053 0,488395 1,0973837 | 20 |
| GOBP_FATTY_ACID_TRANSPORT | 0,3309266 0,8660673 0,0741759 0,415346 1,0973298 | 62 |
| GOBP_NEGATIVE_REGULATION_OF_ESTABLISHMENT_OF_PROTEIN_LOCALIZATION | 0,3164414 0,861089 0,0730744 0,3906606 1,0967423 | 105 |
| GOBP_RESPONSE_TO_DEXAMETHASONE | 0,3450439 0,8738784 0,0741759 0,450759 1,0965816 | 33 |
| GOBP_MICROGLIAL_CELL_ACTIVATION | 0,3580729 0,877063 0,0741759 0,4676088 1,0963576 | 27 |
| GOBP_REGULATION_OF_STEM_CELL_PROLIFERATION | 0,3462986 0,8738784 0,0739901 0,4503973 1,0957017 | 33 |
| GOBP_PROTEIN_IMPORT | 0,2992545 0,8492127 0,0730744 0,3721178 1,0955274 | 188 |
| GOBP_HOMEOSTASIS_OF_NUMBER_OF_CELLS_WITHIN_A_TISSUE | 0,3808256 0,8862233 0,0721798 0,4838732 1,0952395 | 22 |
| GOBP_SENSORY_PERCEPTION_OF_TASTE | 0,3726542 0,8832291 0,0736213 0,509913 1,0948076 | 16 |
| GOBP_REGULATION_OF_SYNAPTIC_TRANSMISSION_GABAERGIC | 0,3569588 0,877063 0,0738053 0,4609945 1,094237 | 29 |
| GOBP_COPULATION | 0,3726542 0,8832291 0,0736213 0,5096357 1,0942124 | 16 |
| GOBP_SECRETORY_GRANULE_ORGANIZATION | 0,3569588 0,877063 0,0738053 0,4609768 1,0941949 | 29 |
| GOBP_REGULATION_OF_MICROTUBULE_POLYMERIZATION | 0,3247232 0,8660673 0,0762797 0,4247111 1,0939724 | 51 |
| GOBP_POSITIVE_REGULATION_OF_SMALL_MOLECULE_METABOLIC_PROCESS | 0,3231982 0,8660673 0,0720033 0,3895904 1,0937376 | 105 |
| GOBP_ENDOTHELIAL_CELL_CHEMOTAXIS | 0,373057 0,8832291 0,0718276 0,470731 1,0934244 | 24 |
| GOBP_GLYCOSYL_COMPOUND_METABOLIC_PROCESS | 0,3191489 0,8634503 0,0751182 0,4060548 1,093108 | 73 |
| GOBP_FAT_CELL_DIFFERENTIATION | 0,3003195 0,8499289 0,0728939 0,3703274 1,0930931 | 199 |
| GOBP_PYRIMIDINE_DEOXYRIBONUCLEOTIDE_METABOLIC_PROCESS | 0,38 0,8862233 0,0723571 0,4834013 1,0930469 | 21 |
| GOBP_GASTRULATION_WITH_MOUTH_FORMING_SECOND | 0,3861518 0,8862233 0,0714786 0,4824385 1,0919921 | 22 |
| GOBP_RESPONSE_TO_PLATELET_DERIVED_GROWTH_FACTOR | 0,3861518 0,8862233 0,0714786 0,4822562 1,0915794 | 22 |
| GOBP_REGULATION_OF_EMBRYONIC_DEVELOPMENT | 0,3398773 0,8706299 0,0738053 0,4211782 1,0913679 | 53 |
| GOBP_TAXIS | 0,2464213 0,8075292 0,0810802 0,3539408 1,091311 | 438 |
| GOBP_EXCITATORY_SYNAPSE_ASSEMBLY | 0,3861518 0,8862233 0,0714786 0,482057 1,0911285 | 22 |
| GOBP_CELL_CHEMOTAXIS | 0,3049041 0,8565046 0,0721798 0,3705447 1,090733 | 189 |
| GOBP_FC_RECEPTOR_SIGNALING_PATHWAY | 0,3375315 0,8693282 0,0755015 0,4412642 1,090238 | 38 |
| GOBP_HEPARAN_SULFATE_PROTEOGLYCAN_BIOSYNTHETIC_PROCESS | 0,3803364 0,8862233 0,0707899 0,4677733 1,0900835 | 25 |
| GOBP_NEURON_RECOGNITION | 0,3417085 0,8727389 0,0747385 0,4279579 1,0898937 | 48 |
| GOBP_REGULATION_OF_INTRACELLULAR_STEROID_HORMONE_RECEPTOR_SIGNALING_PATHWAY | 0,3433373 0,8738784 0,0721798 0,4120632 1,0898113 | 63 |
| GOBP_NEGATIVE_REGULATION_OF_TOLL_LIKE_RECEPTOR_SIGNALING_PATHWAY | 0,3842173 0,8862233 0,0702813 0,46746 1,0893535 | 25 |
| GOBP_NEGATIVE_REGULATION_OF_STRESS_ACTIVATED_PROTEIN_KINASE_SIGNALING_CASCADE | 0,3405337 0,8714711 0,0755015 0,4331625 1,0892991 | 43 |
| GOBP_RETINOIC_ACID_METABOLIC_PROCESS | 0,3744966 0,8832291 0,0734381 0,4998003 1,0891005 | 17 |
| GOBP_POSITIVE_REGULATION_OF_PATHWAY_RESTRICTED_SMAD_PROTEIN_PHOSPHORYLATION | 0,3324905 0,8660673 0,0764767 0,4349471 1,0890928 | 42 |
| GOBP_RRNA_CATABOLIC_PROCESS | 0,3684913 0,8814579 0,0739901 0,4895404 1,0887075 | 19 |
| GOBP_AMINO_ACID_BETAINE_METABOLIC_PROCESS | 0,3744966 0,8832291 0,0734381 0,4995713 1,0886015 | 17 |
| GOBP_GLYCOLIPID_BIOSYNTHETIC_PROCESS | 0,3381123 0,8693282 0,0727141 0,409512 1,0881515 | 65 |
| GOBP_AORTA_DEVELOPMENT | 0,3284133 0,8660673 0,0756946 0,4222732 1,0876929 | 51 |
| GOBP_MYELOID_CELL_APOPTOTIC_PROCESS | 0,3802083 0,8862233 0,0711327 0,4723608 1,0876862 | 23 |
| GOBP_REGULATION_OF_AMINO_ACID_TRANSPORT | 0,3494282 0,8764774 0,0741759 0,454592 1,0872687 | 31 |
| GOBP_ASPARTATE_FAMILY_AMINO_ACID_METABOLIC_PROCESS | 0,3534591 0,877063 0,0730744 0,4295045 1,0858965 | 44 |
| GOBP_COLUMNAR_CUBOIDAL_EPITHELIAL_CELL_DIFFERENTIATION | 0,3321513 0,8660673 0,0730744 0,4042818 1,0857211 | 71 |
| GOBP_REGULATION_OF_CIRCADIAN_RHYTHM | 0,3321957 0,8660673 0,0711327 0,3893226 1,0848027 | 100 |
| GOBP_REACTIVE_NITROGEN_SPECIES_METABOLIC_PROCESS | 0,3609756 0,8785974 0,0704501 0,4149351 1,0847107 | 56 |
| GOBP_POSITIVE_REGULATION_OF_RELEASE_OF_SEQUESTERED_CALCIUM_ION_INTO_CYTOSOL | 0,3537676 0,877063 0,0738053 0,4554616 1,0847031 | 30 |
| GOBP_REGULATION_OF_PROTEIN_POLYUBIQUITINATION | 0,3729032 0,8832291 0,0716527 0,4620934 1,0846898 | 26 |
| GOBP_POSITIVE_REGULATION_OF_CANONICAL_WNT_SIGNALING_PATHWAY | 0,3375143 0,8693282 0,0704501 0,3901987 1,0846738 | 98 |
| GOBP_POSITIVE_REGULATION_OF_TRANSMEMBRANE_TRANSPORT | 0,3206934 0,8660673 0,0704501 0,3713842 1,0841974 | 166 |
| GOBP_CELLULAR_RESPONSE_TO_KETONE | 0,3302433 0,8660673 0,0723571 0,396613 1,0841657 | 80 |
| GOBP_HISTONE_DEUBIQUITINATION | 0,34625 0,8738784 0,0738053 0,4372031 1,0841474 | 39 |
| GOBP_NEPHRON_EPITHELIUM_DEVELOPMENT | 0,3302857 0,8660673 0,0716527 0,3916103 1,0840824 | 91 |
| GOBP_DETECTION_OF_VISIBLE_LIGHT | 0,3710938 0,8831254 0,0723571 0,4598486 1,0840026 | 28 |
| GOBP_MAMMARY_GLAND_DUCT_MORPHOGENESIS | 0,388601 0,8862233 0,0697793 0,466552 1,0837175 | 24 |
| GOBP_NEGATIVE_REGULATION_OF_SECRETION | 0,3348165 0,8693196 0,0696133 0,382383 1,0835819 | 116 |
| GOBP_CONNECTIVE_TISSUE_DEVELOPMENT | 0,3036093 0,8550677 0,0721798 0,3654086 1,0828825 | 216 |
| GOBP_T_CELL_DIFFERENTIATION_INVOLVED_IN_IMMUNE_RESPONSE | 0,3546012 0,877063 0,0716527 0,4177528 1,0824919 | 53 |
| GOBP_POSITIVE_REGULATION_OF_GLUCONEOGENESIS | 0,3766938 0,8849204 0,0736213 0,513201 1,0824862 | 15 |
| GOBP_PLACENTA_DEVELOPMENT | 0,3300111 0,8660673 0,0701132 0,3794932 1,0823243 | 124 |
| GOBP_NEGATIVE_REGULATION_OF_OXIDOREDUCTASE_ACTIVITY | 0,3827493 0,8862233 0,0725352 0,49046 1,0820018 | 18 |
| GOBP_HISTONE_UBIQUITINATION | 0,357868 0,877063 0,0728939 0,4263718 1,0819759 | 47 |
| GOBP_CELLULAR_GLUCAN_METABOLIC_PROCESS | 0,3518072 0,877063 0,0711327 0,4101233 1,0819029 | 60 |
| GOBP_POSITIVE_REGULATION_OF_FATTY_ACID_TRANSPORT | 0,3887399 0,8862233 0,0714786 0,5038952 1,0818873 | 16 |
| GOBP_REGULATION_OF_KETONE_BIOSYNTHETIC_PROCESS | 0,3887399 0,8862233 0,0714786 0,5038891 1,0818741 | 16 |
| GOBP_SOMITE_DEVELOPMENT | 0,3552632 0,877063 0,0702813 0,4071899 1,0818444 | 64 |
| GOBP_REGULATION_OF_CARBOHYDRATE_BIOSYNTHETIC_PROCESS | 0,3462415 0,8738784 0,0691198 0,3911716 1,0817988 | 86 |
| GOBP_INNER_EAR_RECEPTOR_CELL_STEREOCILIUM_ORGANIZATION | 0,3867188 0,8862233 0,0702813 0,4697945 1,0817768 | 23 |
| GOBP_PHOTOTRANSDUCTION | 0,3898964 0,8869191 0,0696133 0,4656744 1,081679 | 24 |
| GOBP_REGULATION_OF_MYELOID_CELL_DIFFERENTIATION | 0,3383948 0,8693282 0,0678338 0,373162 1,0809677 | 149 |
| GOBP_POSITIVE_REGULATION_OF_FILOPODIUM_ASSEMBLY | 0,3736979 0,8832291 0,0720033 0,4608721 1,0805628 | 27 |
| GOBP_EYE_PHOTORECEPTOR_CELL_DIFFERENTIATION | 0,3623737 0,8797754 0,0720033 0,4440264 1,0802454 | 34 |
| GOBP_POSITIVE_REGULATION_OF_HISTONE_MODIFICATION | 0,3501144 0,8764774 0,0687943 0,3909866 1,0798999 | 92 |
| GOBP_POSITIVE_REGULATION_OF_MRNA_METABOLIC_PROCESS | 0,3377778 0,8693282 0,0691198 0,3811973 1,0797896 | 115 |
| GOBP_PURINE_NUCLEOSIDE_MONOPHOSPHATE_BIOSYNTHETIC_PROCESS | 0,3906667 0,8869191 0,0709609 0,477507 1,079719 | 21 |
| GOBP_POSITIVE_REGULATION_OF_ION_TRANSPORT | 0,3102345 0,8581768 0,0713053 0,3654826 1,0796588 | 207 |
| GOBP_POSITIVE_REGULATION_OF_MUSCLE_TISSUE_DEVELOPMENT | 0,3821138 0,8862233 0,0728939 0,5116846 1,0792877 | 15 |
| GOBP_LAMELLIPODIUM_ASSEMBLY | 0,3586041 0,877063 0,0701132 0,4083955 1,0789668 | 62 |
| GOBP_MUSCLE_CELL_PROLIFERATION | 0,3449024 0,8738784 0,0668966 0,3703384 1,0786493 | 160 |
| GOBP_ORGANOPHOSPHATE_CATABOLIC_PROCESS | 0,3399558 0,8706299 0,0684715 0,376551 1,0783154 | 135 |
| GOBP_LEUKOCYTE_CELL_CELL_ADHESION | 0,2992701 0,8492127 0,0720033 0,3583973 1,0782706 | 258 |
| GOBP_EMBRYONIC_DIGIT_MORPHOGENESIS | 0,3680203 0,8814579 0,0714786 0,4248225 1,0780443 | 47 |
| GOBP_AMINOGLYCAN_BIOSYNTHETIC_PROCESS | 0,3486455 0,8764774 0,0704501 0,3994742 1,0780101 | 74 |
| GOBP_CARDIAC_CHAMBER_MORPHOGENESIS | 0,3447099 0,8738784 0,0692836 0,3866347 1,0773132 | 100 |
| GOBP_RESPONSE_TO_INTERLEUKIN_4 | 0,3958603 0,8896911 0,0687943 0,4622535 1,0772204 | 25 |
| GOBP_GLUTATHIONE_METABOLIC_PROCESS | 0,3654321 0,8810414 0,0704501 0,4188258 1,0766137 | 50 |
| GOBP_PROTEIN_HOMOOLIGOMERIZATION | 0,3453159 0,8738784 0,0670513 0,3721707 1,0765549 | 147 |
| GOBP_POSITIVE_REGULATION_OF_ORGANELLE_ORGANIZATION | 0,2811861 0,8359478 0,0741759 0,3493596 1,0763723 | 432 |
| GOBP_PATHWAY_RESTRICTED_SMAD_PROTEIN_PHOSPHORYLATION | 0,3719512 0,8832291 0,0689567 0,4127362 1,0762878 | 55 |
| GOBP_REGULATION_OF_WNT_SIGNALING_PATHWAY | 0,2933194 0,8459041 0,0730744 0,3572107 1,0757858 | 294 |
| GOBP_PROTEIN_MATURATION | 0,3159541 0,861089 0,0692836 0,3588782 1,0757635 | 252 |
| GOBP_REGULATION_OF_LYASE_ACTIVITY | 0,3764115 0,8849204 0,0697793 0,4418975 1,0750238 | 33 |
| GOBP_POSITIVE_REGULATION_OF_WNT_SIGNALING_PATHWAY | 0,3503326 0,8764774 0,0672065 0,3761153 1,0748392 | 127 |
| GOBP_LONG_CHAIN_FATTY_ACID_BIOSYNTHETIC_PROCESS | 0,3858478 0,8862233 0,0716527 0,483129 1,074449 | 19 |
| GOBP_ORGANIC_ACID_TRANSPORT | 0,3050314 0,8565046 0,0713053 0,3598672 1,0741047 | 234 |
| GOBP_REGULATION_OF_ORGAN_GROWTH | 0,3546099 0,877063 0,0697793 0,3989822 1,0740685 | 73 |
| GOBP_NEGATIVE_REGULATION_OF_INTERLEUKIN_1_BETA_PRODUCTION | 0,3834688 0,8862233 0,0727141 0,5092038 1,0740551 | 15 |
| GOBP_POSITIVE_REGULATION_OF_HISTONE_H3_K4_METHYLATION | 0,3984476 0,8910522 0,0684715 0,460835 1,0739147 | 25 |
| GOBP_SENSORY_PERCEPTION_OF_LIGHT_STIMULUS | 0,3526201 0,877063 0,0661326 0,3733882 1,0735304 | 139 |
| GOBP_NEGATIVE_REGULATION_OF_ORGANIC_ACID_TRANSPORT | 0,3981233 0,8910522 0,0702813 0,4996938 1,0728666 | 16 |
| GOBP_EPITHELIAL_CELL_PROLIFERATION | 0,3061013 0,8571368 0,0704501 0,3536706 1,0721751 | 323 |
| GOBP_HEPATICOBILIARY_SYSTEM_DEVELOPMENT | 0,3501684 0,8764774 0,0678338 0,3806072 1,0720638 | 109 |
| GOBP_MICROTUBULE_NUCLEATION | 0,3801757 0,8862233 0,0692836 0,4405975 1,0718613 | 33 |
| GOBP_RIBONUCLEOSIDE_MONOPHOSPHATE_BIOSYNTHETIC_PROCESS | 0,3754789 0,8844353 0,0707899 0,450066 1,0718531 | 30 |
| GOBP_CAMP_MEDIATED_SIGNALING | 0,3664987 0,8810414 0,0713053 0,4338145 1,0718319 | 38 |
| GOBP_NEGATIVE_REGULATION_OF_RESPONSE_TO_EXTERNAL_STIMULUS | 0,3069719 0,8571747 0,0706196 0,3556659 1,0716244 | 281 |
| GOBP_REGULATION_OF_HUMORAL_IMMUNE_RESPONSE | 0,3973333 0,8910522 0,0701132 0,4737725 1,0712746 | 21 |
| GOBP_PHOSPHOLIPID_CATABOLIC_PROCESS | 0,36625 0,8810414 0,0709609 0,431703 1,0705086 | 39 |
| GOBP_RNA_PHOSPHODIESTER_BOND_HYDROLYSIS | 0,3599114 0,8774738 0,0658312 0,3742687 1,0704365 | 133 |
| GOBP_ANTERIOR_POSTERIOR_PATTERN_SPECIFICATION | 0,3539631 0,877063 0,0656814 0,3699025 1,0703192 | 148 |
| GOBP_T_CELL_ACTIVATION_INVOLVED_IN_IMMUNE_RESPONSE | 0,3652482 0,8810414 0,0683111 0,3984997 1,0701928 | 71 |
| GOBP_NEGATIVE_REGULATION_OF_MUSCLE_CONTRACTION | 0,400266 0,8929924 0,0696133 0,4762278 1,0700451 | 20 |
| GOBP_LAMELLIPODIUM_ORGANIZATION | 0,3573086 0,877063 0,0684715 0,391894 1,0699328 | 79 |
| GOBP_REGULATION_OF_NITRIC_OXIDE_METABOLIC_PROCESS | 0,3770492 0,8849204 0,0699459 0,428884 1,0697703 | 41 |
| GOBP_TUBE_FORMATION | 0,3576159 0,877063 0,0659816 0,373433 1,0693866 | 135 |
| GOBP_REGULATION_OF_CYTOSKELETON_ORGANIZATION | 0,3135853 0,8598592 0,0686326 0,346182 1,0691389 | 468 |
| GOBP_POSITIVE_REGULATION_OF_TYPE_I_INTERFERON_MEDIATED_SIGNALING_PATHWAY | 0,4021448 0,89378 0,0697793 0,4977631 1,0687213 | 16 |
| GOBP_POSITIVE_REGULATION_OF_CALCIUM_ION_TRANSMEMBRANE_TRANSPORTER_ACTIVITY | 0,369483 0,8814579 0,0709609 0,4373225 1,0685356 | 35 |
| GOBP_PYRIMIDINE_NUCLEOTIDE_METABOLIC_PROCESS | 0,3813776 0,8862233 0,0699459 0,423924 1,0684766 | 45 |
| GOBP_CELLULAR_SENESCENCE | 0,3730632 0,8832291 0,067676 0,3996344 1,0682722 | 69 |
| GOBP_CELLULAR_CARBOHYDRATE_METABOLIC_PROCESS | 0,3256785 0,8660673 0,0678338 0,3568446 1,0670679 | 241 |
| GOBP_SIGNAL_RELEASE | 0,3133402 0,8598592 0,0692836 0,3502912 1,0662103 | 347 |
| GOBP_NEGATIVE_REGULATION_OF_RESPONSE_TO_BIOTIC_STIMULUS | 0,3668571 0,8812547 0,0664364 0,3851288 1,06614 | 91 |
| GOBP_UNSATURATED_FATTY_ACID_BIOSYNTHETIC_PROCESS | 0,3778338 0,8860933 0,0697793 0,4314599 1,0660143 | 38 |
| GOBP_REGULATION_OF_CYCLASE_ACTIVITY | 0,3783102 0,8862233 0,0697793 0,4325868 1,0658984 | 37 |
| GOBP_AXIS_ELONGATION | 0,4026667 0,8941009 0,0694481 0,4712512 1,0655737 | 21 |
| GOBP_CYCLIC_NUCLEOTIDE_MEDIATED_SIGNALING | 0,3871359 0,8862233 0,0667426 0,4054818 1,0655119 | 58 |
| GOBP_LABYRINTHINE_LAYER_MORPHOGENESIS | 0,4147651 0,9 0,0683111 0,4888784 1,0653009 | 17 |
| GOBP_NEURON_PROJECTION_GUIDANCE | 0,3436499 0,8738784 0,0662842 0,3606037 1,0651234 | 206 |
| GOBP_REGULATION_OF_T_HELPER_CELL_DIFFERENTIATION | 0,3893229 0,8869191 0,0699459 0,4542673 1,0650772 | 27 |
| GOBP_PEPTIDYL_LYSINE_METHYLATION | 0,3713969 0,8831254 0,0643583 0,3733847 1,0648817 | 126 |
| GOBP_NEGATIVE_REGULATION_OF_MYELOID_CELL_DIFFERENTIATION | 0,3814681 0,8862233 0,0670513 0,4028011 1,0648761 | 61 |
| GOBP_NEGATIVE_REGULATION_OF_CELL_ADHESION | 0,3386243 0,8693282 0,0665892 0,3587552 1,0648594 | 219 |
| GOBP_ACIDIC_AMINO_ACID_TRANSPORT | 0,3878049 0,8862233 0,0668966 0,4088869 1,0646 | 54 |
| GOBP_RIBONUCLEOPROTEIN_COMPLEX_SUBUNIT_ORGANIZATION | 0,362955 0,8797754 0,0637845 0,3621535 1,0644764 | 176 |
| GOBP_NEGATIVE_REGULATION_OF_PRODUCTION_OF_MOLECULAR_MEDIATOR_OF_IMMUNE_RESPONSE | 0,3978638 0,8910522 0,0701132 0,4785863 1,0643464 | 19 |
| GOBP_REGULATION_OF_GLUCOSE_IMPORT | 0,3861635 0,8862233 0,0686326 0,4209624 1,0642997 | 44 |
| GOBP_REGULATION_OF_ANION_TRANSPORT | 0,3791866 0,8862233 0,0670513 0,4004546 1,0639495 | 64 |
| GOBP_CELLULAR_RESPONSE_TO_HORMONE_STIMULUS | 0,3329928 0,8660673 0,0656814 0,3443227 1,0633969 | 468 |
| GOBP_REGULATION_OF_PEPTIDE_TRANSPORT | 0,3697572 0,8814579 0,0643583 0,3713257 1,0633519 | 135 |
| GOBP_NEGATIVE_REGULATION_OF_TRANSMEMBRANE_RECEPTOR_PROTEIN_SERINE_THREONINE_KINASE_SIGNALING_PATHWAY | 0,3703284 0,8822599 0,0655321 0,3802823 1,0627473 | 102 |
| GOBP_SKELETAL_SYSTEM_DEVELOPMENT | 0,3183214 0,8622448 0,0679923 0,3453619 1,062726 | 418 |
| GOBP_AGING | 0,3409802 0,8714711 0,0655321 0,355212 1,0624178 | 242 |
| GOBP_MUSCLE_CELL_MIGRATION | 0,3885648 0,8862233 0,0646485 0,3919231 1,0621154 | 76 |
| GOBP_REGIONALIZATION | 0,3333333 0,8660673 0,0667426 0,3527785 1,0614793 | 260 |
| GOBP_DEVELOPMENTAL_INDUCTION | 0,3958333 0,8896911 0,0691198 0,4527127 1,0614322 | 27 |
| GOBP_POSITIVE_REGULATION_OF_PROTEIN_DEPHOSPHORYLATION | 0,38 0,8862233 0,0691198 0,4280319 1,0614053 | 39 |
| GOBP_MESONEPHRIC_TUBULE_MORPHOGENESIS | 0,3982412 0,8910522 0,0670513 0,4167622 1,0613814 | 48 |
| GOBP_POSITIVE_REGULATION_OF_NOTCH_SIGNALING_PATHWAY | 0,3786164 0,8862233 0,0696133 0,4275323 1,0613411 | 40 |
| GOBP_MONOCYTE_CHEMOTAXIS | 0,4087969 0,8971086 0,0672065 0,4553506 1,061134 | 25 |
| GOBP_NEGATIVE_REGULATION_OF_SIGNAL_TRANSDUCTION_IN_ABSENCE_OF_LIGAND | 0,4106218 0,8974438 0,0670513 0,4568161 1,0611027 | 24 |
| GOBP_POSTSYNAPTIC_SIGNAL_TRANSDUCTION | 0,3958333 0,8896911 0,0691198 0,4501331 1,0611002 | 28 |
| GOBP_POSITIVE_REGULATION_OF_VASCULAR_ENDOTHELIAL_GROWTH_FACTOR_PRODUCTION | 0,4106218 0,8974438 0,0670513 0,4567401 1,0609261 | 24 |
| GOBP_SPECIFICATION_OF_ANIMAL_ORGAN_IDENTITY | 0,3958333 0,8896911 0,0691198 0,4524396 1,0607919 | 27 |
| GOBP_NEGATIVE_REGULATION_OF_ADENYLATE_CYCLASE_ACTIVITY | 0,4105691 0,8974438 0,0692836 0,502901 1,0607607 | 15 |
| GOBP_CELL_MIGRATION_INVOLVED_IN_HEART_DEVELOPMENT | 0,4142091 0,9 0,0683111 0,4939399 1,0605127 | 16 |
| GOBP_CEREBELLAR_CORTEX_DEVELOPMENT | 0,3966837 0,8908192 0,0679923 0,4207093 1,0603743 | 45 |
| GOBP_FOREBRAIN_NEURON_DIFFERENTIATION | 0,395202 0,8896911 0,067676 0,4355126 1,0595328 | 34 |
| GOBP_ADULT_LOCOMOTORY_BEHAVIOR | 0,3744019 0,8832291 0,067676 0,398173 1,0594723 | 66 |
| GOBP_REGULATION_OF_CELL_PROJECTION_ASSEMBLY | 0,3620873 0,8797754 0,0636424 0,3588797 1,0593031 | 199 |
| GOBP_REPLACEMENT_OSSIFICATION | 0,4087969 0,8971086 0,0672065 0,4545475 1,0592626 | 25 |
| GOBP_REGULATION_OF_GLUCOSE_TRANSMEMBRANE_TRANSPORT | 0,3956311 0,8896911 0,0656814 0,4030878 1,059221 | 58 |
| GOBP_TRANSMEMBRANE_RECEPTOR_PROTEIN_TYROSINE_KINASE_SIGNALING_PATHWAY | 0,338726 0,8693282 0,0643583 0,3413665 1,0587743 | 497 |
| GOBP_REGULATION_OF_EPITHELIAL_CELL_MIGRATION | 0,3695421 0,8814579 0,0626618 0,3581743 1,0587261 | 202 |
| GOBP_REGULATION_OF_GENE_SILENCING_BY_RNA | 0,4022843 0,89378 0,0670513 0,4189625 1,0585207 | 46 |
| GOBP_FC_GAMMA_RECEPTOR_SIGNALING_PATHWAY | 0,4167776 0,9009229 0,067676 0,4676439 1,0585046 | 22 |
| GOBP_REGULATION_OF_PHOSPHATIDYLINOSITOL_3_KINASE_SIGNALING | 0,3844367 0,8862233 0,0649408 0,3863655 1,0582577 | 82 |
| GOBP_PHOSPHATIDYLCHOLINE_METABOLIC_PROCESS | 0,4 0,8929924 0,0653834 0,405749 1,0580674 | 55 |
| GOBP_SPLICEOSOMAL_SNRNP_ASSEMBLY | 0,3883985 0,8862233 0,0684715 0,4291851 1,0575164 | 37 |
| GOBP_T_HELPER_1_TYPE_IMMUNE_RESPONSE | 0,3926768 0,8896911 0,0679923 0,4379883 1,0571441 | 32 |
| GOBP_PEPTIDYL_L_CYSTEINE_S_PALMITOYLATION | 0,4114583 0,8982272 0,0672065 0,4590958 1,0571413 | 23 |
| GOBP_EXECUTION_PHASE_OF_APOPTOSIS | 0,4002418 0,8929924 0,0649408 0,4021928 1,0570196 | 59 |
| GOBP_HISTONE_H3_K14_ACETYLATION | 0,4202128 0,9037217 0,0672065 0,4703004 1,0567266 | 20 |
| GOBP_REGULATION_OF_ASTROCYTE_DIFFERENTIATION | 0,4113842 0,8982272 0,0668966 0,4530347 1,0557373 | 25 |
| GOBP_REGULATION_OF_EXTRACELLULAR_MATRIX_ORGANIZATION | 0,3855879 0,8862233 0,0689567 0,4215901 1,0556473 | 42 |
| GOBP_APOPTOTIC_PROCESS_INVOLVED_IN_DEVELOPMENT | 0,3883985 0,8862233 0,0684715 0,4319407 1,0553861 | 35 |
| GOBP_ORGANIC_CYCLIC_COMPOUND_CATABOLIC_PROCESS | 0,3384458 0,8693282 0,0649408 0,3422721 1,0547229 | 430 |
| GOBP_PLATELET_ACTIVATION | 0,386545 0,8862233 0,0637845 0,3802915 1,0545451 | 96 |
| GOBP_NUCLEOSIDE_BISPHOSPHATE_BIOSYNTHETIC_PROCESS | 0,407767 0,8971086 0,0642141 0,4016089 1,0544962 | 57 |
| GOBP_REGULATION_OF_LYMPHOCYTE_ACTIVATION | 0,3557394 0,877063 0,063079 0,3475368 1,0537204 | 324 |
| GOBP_CELLULAR_RESPONSE_TO_INORGANIC_SUBSTANCE | 0,3818182 0,8862233 0,0613026 0,3583598 1,0533304 | 182 |
| GOBP_HOMEOSTASIS_OF_NUMBER_OF_CELLS | 0,362963 0,8797754 0,0632191 0,3538744 1,0530877 | 228 |
| GOBP_VASCULAR_ASSOCIATED_SMOOTH_MUSCLE_CELL_MIGRATION | 0,4173333 0,9009229 0,067676 0,4655589 1,0527025 | 21 |
| GOBP_REGULATORY_T_CELL_DIFFERENTIATION | 0,4152652 0,9 0,0664364 0,4516338 1,0524726 | 25 |
| GOBP_RESPONSE_TO_OXIDATIVE_STRESS | 0,3528807 0,877063 0,0632191 0,3444184 1,0524241 | 368 |
| GOBP_REGULATION_OF_REACTIVE_OXYGEN_SPECIES_METABOLIC_PROCESS | 0,3904338 0,8869191 0,0621124 0,3719949 1,0520298 | 114 |
| GOBP_MONOCARBOXYLIC_ACID_BIOSYNTHETIC_PROCESS | 0,3911159 0,8873324 0,0607719 0,3623538 1,0520295 | 152 |
| GOBP_NEGATIVE_REGULATION_OF_CANONICAL_WNT_SIGNALING_PATHWAY | 0,3860619 0,8862233 0,0623861 0,3698014 1,0517851 | 119 |
| GOBP_VESICLE_FUSION_TO_PLASMA_MEMBRANE | 0,4098798 0,8974438 0,0686326 0,4729354 1,051779 | 19 |
| GOBP_PROTEIN_REFOLDING | 0,4322148 0,9090275 0,0662842 0,4825787 1,0515735 | 17 |
| GOBP_STEROID_METABOLIC_PROCESS | 0,358351 0,877063 0,0637845 0,35358 1,0515214 | 224 |
| GOBP_EMBRYONIC_MORPHOGENESIS | 0,3637285 0,8799278 0,0610364 0,3386454 1,0495419 | 491 |
| GOBP_POSITIVE_REGULATION_OF_T_HELPER_CELL_DIFFERENTIATION | 0,4214092 0,9043534 0,0679923 0,4975683 1,0495126 | 15 |
| GOBP_REGULATION_OF_HEMOPOIESIS | 0,3649635 0,8810414 0,062249 0,3487695 1,0494575 | 267 |
| GOBP_REGULATION_OF_DNA_METHYLATION | 0,433557 0,9090275 0,0661326 0,4815305 1,0492894 | 17 |
| GOBP_MEMBRANE_REPOLARIZATION | 0,3934426 0,8896911 0,0678338 0,4270202 1,049166 | 36 |
| GOBP_METENCEPHALON_DEVELOPMENT | 0,4057143 0,8971086 0,0615707 0,3796424 1,0487484 | 89 |
| GOBP_NEGATIVE_REGULATION_OF_IMMUNE_RESPONSE | 0,3955801 0,8896911 0,0611693 0,3686806 1,0482761 | 117 |
| GOBP_ORGANIC_HYDROXY_COMPOUND_BIOSYNTHETIC_PROCESS | 0,3923241 0,8895363 0,0598603 0,3558286 1,0477827 | 190 |
| GOBP_MODULATION_OF_EXCITATORY_POSTSYNAPTIC_POTENTIAL | 0,4035309 0,8954906 0,0665892 0,4286885 1,0474397 | 35 |
| GOBP_MORPHOGENESIS_OF_EMBRYONIC_EPITHELIUM | 0,4015487 0,8935596 0,0605093 0,3657289 1,0474216 | 130 |
| GOBP_C4_DICARBOXYLATE_TRANSPORT | 0,4230272 0,9043534 0,0655321 0,4494387 1,0473572 | 25 |
| GOBP_CARBOXYLIC_ACID_TRANSPORT | 0,3855165 0,8862233 0,0606404 0,3548159 1,047308 | 199 |
| GOBP_CELLULAR_KETONE_METABOLIC_PROCESS | 0,4069264 0,8971086 0,0588438 0,3591467 1,0470782 | 161 |
| GOBP_NUCLEOBASE_CONTAINING_SMALL_MOLECULE_CATABOLIC_PROCESS | 0,421875 0,9043534 0,0659816 0,4546173 1,0468289 | 23 |
| GOBP_REGULATION_OF_FATTY_ACID_BIOSYNTHETIC_PROCESS | 0,4149485 0,9 0,0662842 0,4409826 1,0467358 | 29 |
| GOBP_REGULATION_OF_MORPHOGENESIS_OF_AN_EPITHELIUM | 0,408313 0,8971086 0,0645031 0,4049101 1,0464427 | 52 |
| GOBP_HEART_VALVE_DEVELOPMENT | 0,4097561 0,8974438 0,0642141 0,4018286 1,0462227 | 54 |
| GOBP_LYTIC_VACUOLE_ORGANIZATION | 0,3978622 0,8910522 0,0643583 0,3905912 1,04602 | 70 |
| GOBP_POSITIVE_REGULATION_OF_MIRNA_TRANSCRIPTION | 0,40875 0,8971086 0,0655321 0,4217828 1,0459091 | 39 |
| GOBP_NEGATIVE_REGULATION_OF_CELLULAR_AMIDE_METABOLIC_PROCESS | 0,39381 0,8896911 0,0597318 0,3547881 1,0458204 | 192 |
| GOBP_NUCLEOBASE_CONTAINING_SMALL_MOLECULE_METABOLIC_PROCESS | 0,3798371 0,8862233 0,0592219 0,3379331 1,0456257 | 480 |
| GOBP_RESPONSE_TO_METAL_ION | 0,3802083 0,8862233 0,0602484 0,3468627 1,0453242 | 284 |
| GOBP_ORGANOPHOSPHATE_BIOSYNTHETIC_PROCESS | 0,3767821 0,8849204 0,0596037 0,3374039 1,0448133 | 484 |
| GOBP_DSRNA_PROCESSING | 0,4135802 0,9 0,0643583 0,4064308 1,0447519 | 50 |
| GOBP_ENDOTHELIAL_CELL_MIGRATION | 0,3955224 0,8896911 0,059476 0,3548371 1,0444963 | 189 |
| GOBP_POSITIVE_REGULATION_OF_CHOLESTEROL_EFFLUX | 0,4287617 0,9075254 0,0662842 0,4612083 1,0439379 | 22 |
| GOBP_INOSITOL_PHOSPHATE_CATABOLIC_PROCESS | 0,423231 0,9043534 0,0670513 0,4694022 1,0439214 | 19 |
| GOBP_NEGATIVE_REGULATION_OF_LYASE_ACTIVITY | 0,4366577 0,9093808 0,0659816 0,4725412 1,0424711 | 18 |
| GOBP_UNSATURATED_FATTY_ACID_METABOLIC_PROCESS | 0,4176471 0,9009229 0,0615707 0,3858761 1,0420375 | 75 |
| GOBP_HEART_GROWTH | 0,4135876 0,9 0,0626618 0,3896465 1,0415732 | 69 |
| GOBP_POSITIVE_REGULATION_OF_ENDOTHELIAL_CELL_PROLIFERATION | 0,4159907 0,9002749 0,0610364 0,3808671 1,0411235 | 80 |
| GOBP_NEGATIVE_REGULATION_OF_KINASE_ACTIVITY | 0,3959445 0,8896911 0,059476 0,3524087 1,0409175 | 206 |
| GOBP_WALKING_BEHAVIOR | 0,4153646 0,9 0,0667426 0,4437921 1,0405169 | 27 |
| GOBP_REGULATION_OF_PRODUCTION_OF_SMALL_RNA_INVOLVED_IN_GENE_SILENCING_BY_RNA | 0,4299065 0,9084133 0,0662842 0,4676984 1,0401322 | 19 |
| GOBP_REGULATION_OF_FATTY_ACID_TRANSPORT | 0,4335938 0,9090275 0,0646485 0,451672 1,0400469 | 23 |
| GOBP_NEUROTRANSMITTER_RECEPTOR_TRANSPORT | 0,4354194 0,9090275 0,0655321 0,4592323 1,0394652 | 22 |
| GOBP_NATURAL_KILLER_CELL_DIFFERENTIATION | 0,4375 0,9105785 0,0652353 0,4626177 1,0394642 | 20 |
| GOBP_MUSCLE_HYPERTROPHY | 0,4054374 0,8970719 0,0632191 0,3869708 1,0392314 | 71 |
| GOBP_VASCULOGENESIS | 0,4043062 0,8958579 0,0639272 0,3905605 1,0392168 | 66 |
| GOBP_CELLULAR_RESPONSE_TO_FLUID_SHEAR_STRESS | 0,4420485 0,9125737 0,0653834 0,4708988 1,0388478 | 18 |
| GOBP_TYPE_B_PANCREATIC_CELL_PROLIFERATION | 0,4325768 0,9090275 0,0659816 0,4669185 1,0383978 | 19 |
| GOBP_POSITIVE_REGULATION_OF_DENDRITE_DEVELOPMENT | 0,4477212 0,9158429 0,0645031 0,4836255 1,0383672 | 16 |
| GOBP_POSITIVE_REGULATION_OF_EPITHELIAL_CELL_PROLIFERATION | 0,4118919 0,8986515 0,0582216 0,3569359 1,0383132 | 156 |
| GOBP_POSITIVE_REGULATION_OF_PROTEIN_POLYUBIQUITINATION | 0,4477212 0,9158429 0,0645031 0,4835149 1,0381296 | 16 |
| GOBP_ORGANIC_ACID_BIOSYNTHETIC_PROCESS | 0,4044118 0,8958579 0,0577309 0,3482482 1,0380383 | 231 |
| GOBP_REGULATION_OF_KIDNEY_DEVELOPMENT | 0,4326425 0,9090275 0,0645031 0,4468838 1,0380317 | 24 |
| GOBP_NEGATIVE_REGULATION_OF_PHOSPHATASE_ACTIVITY | 0,4269377 0,9062536 0,0642141 0,4338829 1,0377377 | 31 |
| GOBP_BILE_ACID_METABOLIC_PROCESS | 0,4224464 0,9043534 0,0643583 0,4245127 1,0372368 | 35 |
| GOBP_NUCLEAR_TRANSCRIBED_MRNA_POLY_A_TAIL_SHORTENING | 0,4179688 0,9009229 0,0664364 0,4422337 1,036863 | 27 |
| GOBP_POSITIVE_REGULATION_OF_PROTEIN_BINDING | 0,4247059 0,9045465 0,0607719 0,3837402 1,0362697 | 75 |
| GOBP_POSITIVE_REGULATION_OF_PROTEIN_LOCALIZATION_TO_CELL_PERIPHERY | 0,440534 0,9118932 0,0605093 0,394319 1,0361786 | 58 |
| GOBP_REGULATION_OF_EXTRINSIC_APOPTOTIC_SIGNALING_PATHWAY_IN_ABSENCE_OF_LIGAND | 0,4303639 0,9088685 0,0632191 0,4258622 1,036014 | 33 |
| GOBP_POSITIVE_REGULATION_OF_DNA_RECOMBINATION | 0,4259928 0,9062536 0,0617054 0,3921159 1,0359566 | 62 |
| GOBP_REGULATION_OF_ARP2_3_COMPLEX_MEDIATED_ACTIN_NUCLEATION | 0,4414894 0,9125737 0,0647943 0,4609995 1,0358283 | 20 |
| GOBP_REGULATION_OF_DNA_TEMPLATED_TRANSCRIPTION_IN_RESPONSE_TO_STRESS | 0,4317343 0,9090275 0,0621124 0,4020948 1,0357173 | 51 |
| GOBP_POSITIVE_REGULATION_OF_GLIAL_CELL_DIFFERENTIATION | 0,4199243 0,9037217 0,0646485 0,4202146 1,0354132 | 37 |
| GOBP_LOCOMOTORY_EXPLORATION_BEHAVIOR | 0,4430894 0,9132893 0,0655321 0,4908139 1,0352656 | 15 |
| GOBP_POSITIVE_REGULATION_OF_LIPID_TRANSPORT | 0,4241338 0,9043534 0,0615707 0,3895981 1,0352364 | 65 |
| GOBP_SENSORY_ORGAN_MORPHOGENESIS | 0,4083156 0,8971086 0,0579755 0,3503735 1,035192 | 208 |
| GOBP_ESTABLISHMENT_OR_MAINTENANCE_OF_MONOPOLAR_CELL_POLARITY | 0,4359638 0,9090275 0,0640704 0,4439128 1,0344799 | 25 |
| GOBP_POSITIVE_REGULATION_OF_CATION_CHANNEL_ACTIVITY | 0,4417476 0,9125737 0,0603786 0,3936126 1,0343225 | 58 |
| GOBP_PYRIDINE_NUCLEOTIDE_METABOLIC_PROCESS | 0,4378238 0,9105785 0,0639272 0,4452796 1,0343053 | 24 |
| GOBP_CELL_DIFFERENTIATION_INVOLVED_IN_KIDNEY_DEVELOPMENT | 0,4352792 0,9090275 0,0632191 0,409293 1,0340904 | 46 |
| GOBP_PROTEIN_CATABOLIC_PROCESS_IN_THE_VACUOLE | 0,4420772 0,9125737 0,0647943 0,4567148 1,0337669 | 22 |
| GOBP_PROTEIN_AUTOUBIQUITINATION | 0,4078014 0,8971086 0,0629395 0,3849318 1,0337555 | 71 |
| GOBP_REGULATION_OF_CYSTEINE_TYPE_ENDOPEPTIDASE_ACTIVITY_INVOLVED_IN_APOPTOTIC_SIGNALING_PATHWAY | 0,4571046 0,9210223 0,0635008 0,4814519 1,0337004 | 16 |
| GOBP_REGULATION_OF_OXIDATIVE_STRESS_INDUCED_INTRINSIC_APOPTOTIC_SIGNALING_PATHWAY | 0,4348387 0,9090275 0,0640704 0,4403263 1,033595 | 26 |
| GOBP_SENSORY_SYSTEM_DEVELOPMENT | 0,4066736 0,8971086 0,0571259 0,3424551 1,0335822 | 307 |
| GOBP_NEGATIVE_REGULATION_OF_PROTEIN_MODIFICATION_PROCESS | 0,4159021 0,9002749 0,0550211 0,3350811 1,0333748 | 445 |
| GOBP_REGULATION_OF_TISSUE_REMODELING | 0,4390244 0,9107756 0,0609039 0,3967225 1,0329281 | 54 |
| GOBP_CARDIAC_MUSCLE_CELL_MEMBRANE_REPOLARIZATION | 0,4283854 0,9075254 0,0652353 0,4381548 1,0328636 | 28 |
| GOBP_CAMERA_TYPE_EYE_DEVELOPMENT | 0,408142 0,8971086 0,057006 0,3431436 1,0325743 | 263 |
| GOBP_METHYLATION | 0,4173554 0,9009229 0,0554793 0,3402733 1,0324177 | 326 |
| GOBP_POSITIVE_REGULATION_OF_CD4_POSITIVE_ALPHA_BETA_T_CELL_DIFFERENTIATION | 0,4447403 0,9145536 0,0645031 0,4560579 1,0322799 | 22 |
| GOBP_EPITHELIAL_TUBE_FORMATION | 0,4235033 0,9043534 0,0580984 0,3621338 1,0321001 | 123 |
| GOBP_NEGATIVE_REGULATION_OF_CELL_DEVELOPMENT | 0,4237838 0,9043534 0,0568864 0,3550238 1,0319829 | 155 |
| GOBP_CHOLESTEROL_EFFLUX | 0,43125 0,9090275 0,0629395 0,4159027 1,0313279 | 39 |
| GOBP_REGULATION_OF_NEURON_PROJECTION_DEVELOPMENT | 0,4153846 0,9 0,0553643 0,3361086 1,0312072 | 396 |
| GOBP_INTRINSIC_APOPTOTIC_SIGNALING_PATHWAY_IN_RESPONSE_TO_DNA_DAMAGE | 0,4446978 0,9145536 0,0571259 0,371563 1,030341 | 96 |
| GOBP_POSITIVE_REGULATION_OF_CELLULAR_RESPONSE_TO_TRANSFORMING_GROWTH_FACTOR_BETA_STIMULUS | 0,4355045 0,9090275 0,0635008 0,4324536 1,0299084 | 30 |
| GOBP_POSITIVE_REGULATION_OF_GLUCOSE_IMPORT | 0,4296875 0,9084133 0,0650878 0,4366846 1,029398 | 28 |
| GOBP_THYMUS_DEVELOPMENT | 0,4402516 0,9118932 0,062249 0,4146648 1,0293976 | 40 |
| GOBP_TRANSPORT_ALONG_MICROTUBULE | 0,4335135 0,9090275 0,0558265 0,3542864 1,0293586 | 154 |
| GOBP_PEPTIDYL_THREONINE_MODIFICATION | 0,4266667 0,9062536 0,057853 0,3633918 1,029353 | 115 |
| GOBP_MATURATION_OF_5_8S_RRNA_FROM_TRICISTRONIC_RRNA_TRANSCRIPT_SSU_RRNA_5_8S_RRNA_LSU_RRNA | 0,4417098 0,9125737 0,0635008 0,4430544 1,0291368 | 24 |
| GOBP_REGULATION_OF_IMMUNE_EFFECTOR_PROCESS | 0,4216102 0,9043534 0,0561767 0,3462558 1,0285011 | 222 |
| GOBP_RHYTHMIC_BEHAVIOR | 0,4381313 0,9105785 0,0626618 0,4227214 1,0284138 | 34 |
| GOBP_MONOAMINE_TRANSPORT | 0,4502427 0,9158429 0,059476 0,3913585 1,0283992 | 58 |
| GOBP_ENDOTHELIAL_CELL_PROLIFERATION | 0,4345898 0,9090275 0,0568864 0,3606824 1,0279635 | 123 |
| GOBP_POSITIVE_REGULATION_OF_OSSIFICATION | 0,4334171 0,9090275 0,0629395 0,4035157 1,027646 | 48 |
| GOBP_RESPONSE_TO_ACETYLCHOLINE | 0,4500666 0,9158429 0,0639272 0,453822 1,0272192 | 22 |
| GOBP_POSITIVE_REGULATION_OF_CELL_DEATH | 0,4342508 0,9090275 0,0530213 0,3322218 1,0270673 | 472 |
| GOBP_FATTY_ACYL_COA_BIOSYNTHETIC_PROCESS | 0,4479167 0,9158429 0,063079 0,446034 1,0270646 | 23 |
| GOBP_PHOTORECEPTOR_CELL_DEVELOPMENT | 0,4325347 0,9090275 0,0632191 0,4177694 1,0264372 | 36 |
| GOBP_RESPONSE_TO_XENOBIOTIC_STIMULUS | 0,4202294 0,9037217 0,0555947 0,3402142 1,0261561 | 301 |
| GOBP_CARTILAGE_DEVELOPMENT | 0,4404762 0,9118932 0,0551352 0,3519042 1,025963 | 161 |
| GOBP_SULFUR_COMPOUND_METABOLIC_PROCESS | 0,425 0,9046601 0,0550211 0,340393 1,025866 | 286 |
| GOBP_ANION_TRANSPORT | 0,4269547 0,9062536 0,0542316 0,3352476 1,0253914 | 378 |
| GOBP_OLIGOSACCHARIDE_BIOSYNTHETIC_PROCESS | 0,4534574 0,917415 0,0635008 0,4562496 1,0251556 | 20 |
| GOBP_POSITIVE_REGULATION_OF_PHOSPHATIDYLINOSITOL_3_KINASE_SIGNALING | 0,4490291 0,9158429 0,0596037 0,3902975 1,0247959 | 57 |
| GOBP_PROTEIN_ACETYLATION | 0,4348291 0,9090275 0,0551352 0,347504 1,0247908 | 194 |
| GOBP_POSITIVE_REGULATION_OF_CELL_ACTIVATION | 0,4245578 0,9045465 0,0550211 0,3411933 1,0247102 | 254 |
| GOBP_SULFUR_AMINO_ACID_BIOSYNTHETIC_PROCESS | 0,4647696 0,9252407 0,0632191 0,4857994 1,0246886 | 15 |
| GOBP_REGULATION_OF_VESICLE_FUSION | 0,4534574 0,917415 0,0635008 0,4559883 1,0245686 | 20 |
| GOBP_DENDRITE_SELF_AVOIDANCE | 0,4758713 0,9291869 0,0615707 0,4765714 1,0232217 | 16 |
| GOBP_MALE_MEIOSIS_I | 0,4574468 0,9210223 0,063079 0,4552414 1,0228902 | 20 |
| GOBP_RESPONSE_TO_MAGNESIUM_ION | 0,4758713 0,9291869 0,0615707 0,4762406 1,0225115 | 16 |
| GOBP_NEUROTROPHIN_SIGNALING_PATHWAY | 0,4388398 0,9107756 0,0625237 0,415782 1,0215543 | 36 |
| GOBP_POSITIVE_REGULATION_OF_ALPHA_BETA_T_CELL_DIFFERENTIATION | 0,4388398 0,9107756 0,0625237 0,4156921 1,0213334 | 36 |
| GOBP_PROTEIN_POLYMERIZATION | 0,4363257 0,9091933 0,0538979 0,3414353 1,0210469 | 245 |
| GOBP_CELLULAR_RESPONSE_TO_GROWTH_HORMONE_STIMULUS | 0,4649596 0,9252407 0,0629395 0,4627936 1,020967 | 18 |
| GOBP_CELLULAR_RESPONSE_TO_HYDROGEN_PEROXIDE | 0,4515753 0,9170639 0,0574877 0,3767272 1,0209341 | 76 |
| GOBP_PHOTOPERIODISM | 0,4471649 0,9158429 0,0626618 0,430051 1,0207882 | 29 |
| GOBP_CELLULAR_RESPONSE_TO_ABIOTIC_STIMULUS | 0,4360042 0,9090275 0,0537873 0,3387547 1,0199794 | 275 |
| GOBP_CELLULAR_RESPONSE_TO_ATP | 0,4688347 0,9267843 0,0628004 0,4835289 1,0198993 | 15 |
| GOBP_POSITIVE_REGULATION_OF_RECEPTOR_INTERNALIZATION | 0,4544271 0,9188819 0,0623861 0,4428353 1,0196989 | 23 |
| GOBP_RECEPTOR_SIGNALING_PATHWAY_VIA_STAT | 0,4476615 0,9158429 0,0557104 0,3610315 1,0195054 | 112 |
| GOBP_POSITIVE_REGULATION_OF_MYELOID_LEUKOCYTE_DIFFERENTIATION | 0,4483627 0,9158429 0,0614364 0,4126001 1,0194171 | 38 |
| GOBP_BRANCHING_INVOLVED_IN_SALIVARY_GLAND_MORPHOGENESIS | 0,4765101 0,9291869 0,0615707 0,4677883 1,019344 | 17 |
| GOBP_REGULATION_OF_SYSTEMIC_ARTERIAL_BLOOD_PRESSURE_BY_RENIN_ANGIOTENSIN | 0,4812332 0,9326074 0,0610364 0,4746538 1,0191044 | 16 |
| GOBP_REGULATION_OF_CANONICAL_WNT_SIGNALING_PATHWAY | 0,4344609 0,9090275 0,0546808 0,3424234 1,0186553 | 226 |
| GOBP_NEGATIVE_REGULATION_OF_SYSTEMIC_ARTERIAL_BLOOD_PRESSURE | 0,4728997 0,9291869 0,0623861 0,4829345 1,0186456 | 15 |
| GOBP_CYTOSKELETON_DEPENDENT_INTRACELLULAR_TRANSPORT | 0,4494143 0,9158429 0,0534572 0,3459741 1,0185594 | 188 |
| GOBP_HYDROGEN_PEROXIDE_METABOLIC_PROCESS | 0,4574333 0,9210223 0,0609039 0,4258221 1,0184583 | 31 |
| GOBP_REGULATION_OF_BROWN_FAT_CELL_DIFFERENTIATION | 0,467655 0,9266771 0,0626618 0,461581 1,0182919 | 18 |
| GOBP_RESPONSE_TO_PROGESTERONE | 0,4558081 0,9206833 0,0607719 0,418458 1,0180416 | 34 |
| GOBP_PROTEIN_DEPHOSPHORYLATION | 0,447341 0,9158429 0,0526973 0,3389936 1,0180217 | 255 |
| GOBP_ENSHEATHMENT_OF_NEURONS | 0,4584718 0,9223114 0,0543434 0,3571393 1,0179832 | 125 |
| GOBP_POSITIVE_REGULATION_OF_MYELOID_CELL_DIFFERENTIATION | 0,4433962 0,9132893 0,0588438 0,3783364 1,0179411 | 72 |
| GOBP_ERYTHROCYTE_HOMEOSTASIS | 0,4617978 0,9242226 0,0546808 0,3613201 1,0170484 | 108 |
| GOBP_HISTONE_H4_K16_ACETYLATION | 0,4492188 0,9158429 0,0629395 0,4337542 1,016982 | 27 |
| GOBP_INDOLE_CONTAINING_COMPOUND_METABOLIC_PROCESS | 0,4783198 0,9291869 0,0618406 0,4820441 1,0167675 | 15 |
| GOBP_NEGATIVE_REGULATION_OF_PROTEIN_LOCALIZATION | 0,4612069 0,9241356 0,052805 0,3475449 1,0166548 | 169 |
| GOBP_REGULATION_OF_BRANCHING_INVOLVED_IN_URETERIC_BUD_MORPHOGENESIS | 0,4852547 0,9382609 0,0606404 0,4733443 1,0162929 | 16 |
| GOBP_REGULATION_OF_MESENCHYMAL_CELL_PROLIFERATION | 0,4624352 0,9244178 0,0613026 0,4374769 1,016181 | 24 |
| GOBP_PURINE_CONTAINING_COMPOUND_BIOSYNTHETIC_PROCESS | 0,4568966 0,9210223 0,0532386 0,3469242 1,0159433 | 170 |
| GOBP_REGULATION_OF_INFLAMMATORY_RESPONSE_TO_ANTIGENIC_STIMULUS | 0,4632258 0,9248678 0,0610364 0,4326221 1,0155105 | 26 |
| GOBP_RESPONSE_TO_VITAMIN | 0,460241 0,9231849 0,0580984 0,384888 1,0153324 | 60 |
| GOBP_HEART_TRABECULA_MORPHOGENESIS | 0,4518229 0,9170711 0,0626618 0,4330109 1,0152392 | 27 |
| GOBP_VASCULAR_PROCESS_IN_CIRCULATORY_SYSTEM | 0,4471718 0,9158429 0,0537873 0,3434443 1,0150197 | 204 |
| GOBP_EPITHELIAL_TUBE_MORPHOGENESIS | 0,4505723 0,9160174 0,0522693 0,3370792 1,0149344 | 275 |
| GOBP_BLOOD_VESSEL_MORPHOGENESIS | 0,4752275 0,9291869 0,048506 0,3271098 1,0147244 | 496 |
| GOBP_FC_RECEPTOR_MEDIATED_STIMULATORY_SIGNALING_PATHWAY | 0,453125 0,917415 0,0625237 0,4326778 1,0144582 | 27 |
| GOBP_POSITIVE_REGULATION_OF_STEROID_METABOLIC_PROCESS | 0,4734043 0,9291869 0,0614364 0,4508717 1,013072 | 20 |
| GOBP_POLYSACCHARIDE_BIOSYNTHETIC_PROCESS | 0,4741897 0,9291869 0,05653 0,3830243 1,0130102 | 63 |
| GOBP_RESPONSE_TO_ELECTRICAL_STIMULUS | 0,4633838 0,9248678 0,0599892 0,4195955 1,0127505 | 32 |
| GOBP_REGULATION_OF_B_CELL_DIFFERENTIATION | 0,4760638 0,9291869 0,0611693 0,4502989 1,0117849 | 20 |
| GOBP_PHOTORECEPTOR_CELL_MAINTENANCE | 0,4659091 0,9253885 0,0597318 0,4188952 1,0110604 | 32 |
| GOBP_BONE_MORPHOGENESIS | 0,4756381 0,9291869 0,054794 0,3703065 1,0109953 | 79 |
| GOBP_PROTEIN_LOCALIZATION_TO_CELL_PERIPHERY | 0,4635417 0,9248678 0,0510114 0,3349448 1,0109395 | 306 |
| GOBP_ORGANELLE_INHERITANCE | 0,4864499 0,9382609 0,0610364 0,4792706 1,0109174 | 15 |
| GOBP_POSITIVE_REGULATION_OF_VIRAL_PROCESS | 0,4731707 0,9291869 0,0573667 0,3882647 1,010907 | 54 |
| GOBP_ANION_TRANSMEMBRANE_TRANSPORT | 0,466951 0,9262358 0,0517405 0,3418396 1,0102253 | 201 |
| GOBP_AXON_DEVELOPMENT | 0,4887526 0,9387054 0,0477342 0,3277523 1,0099795 | 430 |
| GOBP_MAINTENANCE_OF_CELL_NUMBER | 0,4644444 0,9252407 0,0538979 0,3548147 1,0098063 | 122 |

| GOBP_POSITIVE_REGULATION_OF_ENDOCYTOSIS | 0,4889663 0,9387054 0,0535669 0,368593 1,0095786 | 82 |
| --- | --- | --- |
| GOBP_REGULATION_OF_TRANSMEMBRANE_TRANSPORT | 0,4846939 0,9376905 0,048022 0,3273209 1,0095692 | 439 |
| GOBP_NEGATIVE_REGULATION_OF_ALPHA_BETA_T_CELL_DIFFERENTIATION | 0,4773936 0,9291869 0,0610364 0,4492767 1,009488 | 20 |
| GOBP_REGULATION_OF_SMALL_MOLECULE_METABOLIC_PROCESS | 0,462578 0,9244178 0,0510114 0,3348622 1,0092739 | 270 |
| GOBP_EXCRETION | 0,4702908 0,9279845 0,0593488 0,4030165 1,0091397 | 42 |
| GOBP_SECONDARY_ALCOHOL_METABOLIC_PROCESS | 0,4725644 0,9291869 0,0534572 0,3577406 1,0084495 | 111 |
| GOBP_BRANCHING_MORPHOGENESIS_OF_AN_EPITHELIAL_TUBE | 0,4649611 0,9252407 0,0538979 0,3558049 1,0082659 | 116 |
| GOBP_STEM_CELL_PROLIFERATION | 0,4878049 0,9382609 0,0559429 0,3856372 1,008121 | 56 |
| GOBP_PROSTATE_GLAND_MORPHOGENESIS | 0,4765625 0,9291869 0,0601186 0,4378066 1,0081197 | 23 |
| GOBP_REGULATION_OF_COLLAGEN_METABOLIC_PROCESS | 0,4570313 0,9210223 0,0621124 0,4274605 1,0076539 | 28 |
| GOBP_REGULATION_OF_FAT_CELL_DIFFERENTIATION | 0,4671858 0,9262358 0,053677 0,3554046 1,0071315 | 116 |
| GOBP_REGULATION_OF_HISTONE_MODIFICATION | 0,4707158 0,9283356 0,052163 0,3457384 1,0069991 | 160 |
| GOBP_NOTCH_SIGNALING_PATHWAY | 0,4691892 0,9267843 0,052163 0,3466812 1,0061672 | 151 |
| GOBP_RESPONSE_TO_SALT | 0,4912752 0,9407344 0,0601186 0,4615103 1,0056638 | 17 |
| GOBP_EMBRYONIC_HEART_TUBE_DEVELOPMENT | 0,4689737 0,9267843 0,0567672 0,3761689 1,005526 | 67 |
| GOBP_T_HELPER_1_CELL_DIFFERENTIATION | 0,4946381 0,941623 0,0597318 0,4682944 1,0054506 | 16 |
| GOBP_ICOSANOID_TRANSPORT | 0,4587629 0,9223114 0,0614364 0,4234989 1,0052358 | 29 |
| GOBP_CARDIAC_VENTRICLE_DEVELOPMENT | 0,4818182 0,9326074 0,0532386 0,3611954 1,0051422 | 99 |
| GOBP_INFLAMMATORY_RESPONSE | 0,5096057 0,9479394 0,0453152 0,3239859 1,005034 | 496 |
| GOBP_DETERMINATION_OF_ADULT_LIFESPAN | 0,4959786 0,941623 0,0596037 0,4679418 1,0046934 | 16 |
| GOBP_GLYCOPROTEIN_METABOLIC_PROCESS | 0,4880829 0,9382609 0,0484088 0,3315606 1,0044261 | 331 |
| GOBP_CELLULAR_RESPONSE_TO_ALCOHOL | 0,4793875 0,9307798 0,0551352 0,3719735 1,0037975 | 74 |
| GOBP_NEGATIVE_REGULATION_OF_GROWTH | 0,4776119 0,9291869 0,0507028 0,3396525 1,0035163 | 208 |
| GOBP_RESPONSE_TO_MOLECULE_OF_BACTERIAL_ORIGIN | 0,4751848 0,9291869 0,0504983 0,3372818 1,0034696 | 225 |
| GOBP_REGULATION_OF_LEUKOCYTE_DIFFERENTIATION | 0,4781217 0,9291869 0,0507028 0,3395162 1,0034106 | 204 |
| GOBP_REGULATION_OF_ACTIN_NUCLEATION | 0,4652062 0,9252407 0,0607719 0,4225999 1,0031019 | 29 |
| GOBP_CEREBRAL_CORTEX_RADIALLY_ORIENTED_CELL_MIGRATION | 0,4596354 0,9229558 0,0618406 0,4254994 1,003031 | 28 |
| GOBP_MITOCHONDRIAL_DNA_REPLICATION | 0,495935 0,941623 0,0601186 0,4754273 1,0028109 | 15 |
| GOBP_REGULATION_OF_TOLL_LIKE_RECEPTOR_4_SIGNALING_PATHWAY | 0,4878706 0,9382609 0,0606404 0,4544515 1,0025635 | 18 |
| GOBP_POSITIVE_REGULATION_OF_TRANSCRIPTION_FROM_RNA_POLYMERASE_II_PROMOTER_INVOLVED_IN_CELLULAR_RESPONSE_TO_CHEMICAL_STIMULUS | 0,4766839 0,9291869 0,0598603 0,431588 1,0025022 | 24 |
| GOBP_RESPONSE_TO_FIBROBLAST_GROWTH_FACTOR | 0,4954442 0,941623 0,052057 0,361806 1,0016943 | 87 |
| GOBP_REGULATION_OF_MIRNA_TRANSCRIPTION | 0,4932515 0,941623 0,0557104 0,3864139 1,0012857 | 53 |
| GOBP_REGULATION_OF_PROTEIN_LOCALIZATION_TO_CELL_PERIPHERY | 0,4870932 0,9382609 0,052163 0,3554525 1,00121 | 109 |
| GOBP_SMOOTH_MUSCLE_CELL_MIGRATION | 0,4969988 0,9416872 0,0543434 0,3783534 1,0006566 | 63 |
| GOBP_SMAD_PROTEIN_SIGNAL_TRANSDUCTION | 0,4817432 0,9326074 0,0549074 0,3707908 1,000606 | 74 |
| GOBP_POSITIVE_REGULATION_OF_VASCULATURE_DEVELOPMENT | 0,4877778 0,9382609 0,0516356 0,351435 1,0001877 | 122 |
| GOBP_POSITIVE_REGULATION_OF_PHOSPHOPROTEIN_PHOSPHATASE_ACTIVITY | 0,495302 0,941623 0,0597318 0,458907 0,9999912 | 17 |
| GOBP_ODONTOGENESIS | 0,4982896 0,9416872 0,0518458 0,3597112 0,9999246 | 98 |
| GOBP_NUCLEOTIDE_BINDING_DOMAIN_LEUCINE_RICH_REPEAT_CONTAINING_RECEPTOR_SIGNALING_PATHWAY | 0,4869792 0,9382609 0,0590955 0,4340991 0,9995824 | 23 |
| GOBP_CELL_ACTIVATION_INVOLVED_IN_IMMUNE_RESPONSE | 0,4936034 0,941623 0,0491928 0,33901 0,9987909 | 191 |
| GOBP_REGULATION_OF_CELLULAR_CARBOHYDRATE_METABOLIC_PROCESS | 0,4916944 0,9410574 0,0511148 0,350379 0,9987137 | 125 |
| GOBP_REGULATION_OF_RELEASE_OF_SEQUESTERED_CALCIUM_ION_INTO_CYTOSOL | 0,486747 0,9382609 0,0554793 0,3785804 0,998693 | 60 |
| GOBP_RESPONSE_TO_PROSTAGLANDIN | 0,4893333 0,9389302 0,0599892 0,4415656 0,9984498 | 21 |
| GOBP_SMOOTH_MUSCLE_CELL_DIFFERENTIATION | 0,5006135 0,9437054 0,0550211 0,3852162 0,998182 | 53 |
| GOBP_REGULATION_OF_HEMATOPOIETIC_PROGENITOR_CELL_DIFFERENTIATION | 0,4712644 0,9289298 0,0597318 0,4186759 0,9970962 | 30 |
| GOBP_DICARBOXYLIC_ACID_METABOLIC_PROCESS | 0,5017182 0,9439459 0,0517405 0,36168 0,9970583 | 84 |
| GOBP_NEPHRON_MORPHOGENESIS | 0,4969916 0,9416872 0,0544556 0,3770582 0,9968203 | 61 |
| GOBP_NEGATIVE_REGULATION_OF_ANTIGEN_RECEPTOR_MEDIATED_SIGNALING_PATHWAY | 0,4908854 0,9404674 0,0587186 0,4328781 0,996771 | 23 |
| GOBP_GLYCOSYL_COMPOUND_CATABOLIC_PROCESS | 0,4967742 0,9416872 0,0577309 0,4245876 0,9966508 | 26 |
| GOBP_REGULATION_OF_ANDROGEN_RECEPTOR_SIGNALING_PATHWAY | 0,4864166 0,9382609 0,0588438 0,4273974 0,9959929 | 25 |
| GOBP_CELLULAR_RESPONSE_TO_ACID_CHEMICAL | 0,4882353 0,9382609 0,0542316 0,3684884 0,9950829 | 75 |
| GOBP_REGULATION_OF_MYELOID_LEUKOCYTE_DIFFERENTIATION | 0,5137931 0,9485329 0,0508054 0,3614495 0,9937873 | 83 |
| GOBP_HISTONE_H3_K9_MODIFICATION | 0,501269 0,9439459 0,05653 0,3932133 0,9934645 | 46 |
| GOBP_CELL_REDOX_HOMEOSTASIS | 0,4854981 0,9382609 0,0577309 0,4064869 0,9931934 | 35 |
| GOBP_ANATOMICAL_STRUCTURE_HOMEOSTASIS | 0,5026178 0,9441675 0,0475434 0,3326551 0,9930928 | 233 |
| GOBP_CELLULAR_RESPONSE_TO_ORGANIC_CYCLIC_COMPOUND | 0,5361876 0,9525709 0,0433637 0,3219768 0,9930437 | 444 |
| GOBP_CARBOHYDRATE_HOMEOSTASIS | 0,5069519 0,9477823 0,0481184 0,337791 0,9928722 | 182 |
| GOBP_BRANCHED_CHAIN_AMINO_ACID_CATABOLIC_PROCESS | 0,5039894 0,9457982 0,0584693 0,4417732 0,9926282 | 20 |
| GOBP_ACUTE_INFLAMMATORY_RESPONSE | 0,5072816 0,9477823 0,0538979 0,3779338 0,9923329 | 57 |
| GOBP_CELLULAR_GLUCOSE_HOMEOSTASIS | 0,4972376 0,9416872 0,0504983 0,3489666 0,9922229 | 117 |
| GOBP_POSITIVE_REGULATION_OF_MUSCLE_HYPERTROPHY | 0,4896373 0,9390339 0,0585938 0,4271022 0,9920826 | 24 |
| GOBP_CELL_JUNCTION_ASSEMBLY | 0,5216049 0,9520965 0,0450443 0,3245243 0,9911586 | 363 |
| GOBP_POSITIVE_REGULATION_OF_CELLULAR_COMPONENT_BIOGENESIS | 0,5470348 0,9525767 0,0425878 0,3215906 0,9906771 | 426 |
| GOBP_MUSCLE_ADAPTATION | 0,5091116 0,9479394 0,0508054 0,3577743 0,9905321 | 87 |
| GOBP_REGULATION_OF_LEUKOCYTE_ADHESION_TO_VASCULAR_ENDOTHELIAL_CELL | 0,4973958 0,9416872 0,0580984 0,429872 0,989849 | 23 |
| GOBP_REGULATION_OF_VIRAL_ENTRY_INTO_HOST_CELL | 0,4987374 0,9420595 0,05653 0,4101042 0,989842 | 32 |
| GOBP_REGULATION_OF_PROTEIN_SUMOYLATION | 0,5019973 0,9439459 0,0587186 0,4372885 0,9897959 | 22 |
| GOBP_FC_EPSILON_RECEPTOR_SIGNALING_PATHWAY | 0,5046729 0,9461358 0,0585938 0,4448989 0,9894275 | 19 |
| GOBP_NUCLEOLUS_ORGANIZATION | 0,5147453 0,9489196 0,057853 0,4606968 0,9891381 | 16 |
| GOBP_POSITIVE_REGULATION_OF_ACTIN_NUCLEATION | 0,5160858 0,9496706 0,0577309 0,4603236 0,9883369 | 16 |
| GOBP_ACTIVATION_OF_IMMUNE_RESPONSE | 0,5197439 0,9515973 0,046882 0,3345901 0,9882864 | 206 |
| GOBP_BIOMINERALIZATION | 0,5099778 0,9479394 0,0494905 0,3458043 0,9882183 | 127 |
| GOBP_ORGANISM_EMERGENCE_FROM_PROTECTIVE_STRUCTURE | 0,4947917 0,941623 0,0583453 0,4214783 0,9881999 | 27 |
| GOBP_RESPONSE_TO_LEPTIN | 0,506008 0,9472208 0,0584693 0,4442835 0,9880591 | 19 |
| GOBP_MONOCARBOXYLIC_ACID_METABOLIC_PROCESS | 0,5422177 0,9525709 0,0427591 0,3203113 0,9879651 | 451 |
| GOBP_MAINTENANCE_OF_BLOOD_BRAIN_BARRIER | 0,4930114 0,941623 0,0573667 0,4130416 0,9878907 | 31 |
| GOBP_PROTEIN_STABILIZATION | 0,5178765 0,9512582 0,0477342 0,3382987 0,9876095 | 166 |
| GOBP_MEMBRANE_INVAGINATION | 0,5158924 0,9496706 0,0534572 0,3820702 0,9874158 | 52 |
| GOBP_TRANSPOSITION | 0,496 0,941623 0,0593488 0,4365918 0,9872032 | 21 |
| GOBP_STRESS_ACTIVATED_PROTEIN_KINASE_SIGNALING_CASCADE | 0,5079702 0,9479394 0,0477342 0,3337154 0,9869884 | 211 |
| GOBP_REGULATION_OF_INFLAMMATORY_RESPONSE | 0,5208768 0,9520965 0,0457704 0,3299986 0,9867907 | 241 |
| GOBP_MONOUBIQUITINATED_PROTEIN_DEUBIQUITINATION | 0,4949495 0,941623 0,0568864 0,4052055 0,9858003 | 34 |
| GOBP_RESPONSE_TO_ESTROGEN | 0,5146341 0,9489196 0,0534572 0,3785897 0,9857167 | 54 |
| GOBP_THIOESTER_METABOLIC_PROCESS | 0,5185615 0,9512869 0,0508054 0,3609838 0,9855429 | 79 |
| GOBP_LEUKOCYTE_ADHESION_TO_VASCULAR_ENDOTHELIAL_CELL | 0,5018916 0,9439459 0,0561767 0,4010871 0,9854495 | 36 |
| GOBP_RELAXATION_OF_MUSCLE | 0,5039063 0,9457982 0,0574877 0,4180357 0,9854369 | 28 |
| GOBP_PROTEIN_HOMOTETRAMERIZATION | 0,518239 0,9512582 0,0545681 0,3896575 0,9851531 | 44 |
| GOBP_POSITIVE_REGULATION_OF_G_PROTEIN_COUPLED_RECEPTOR_SIGNALING_PATHWAY | 0,5121951 0,9479394 0,0585938 0,4669887 0,9850115 | 15 |
| GOBP_LIPID_MODIFICATION | 0,5230439 0,9520965 0,0467883 0,3351141 0,9846338 | 181 |
| GOBP_NEGATIVE_REGULATION_OF_PROTEIN_CATABOLIC_PROCESS | 0,525196 0,9520965 0,0486034 0,3492612 0,9845467 | 111 |
| GOBP_HYDROGEN_PEROXIDE_CATABOLIC_PROCESS | 0,5121951 0,9479394 0,0585938 0,4666582 0,9843143 | 15 |
| GOBP_SPHINGOLIPID_BIOSYNTHETIC_PROCESS | 0,5313569 0,9525709 0,0488971 0,3549564 0,9842909 | 96 |
| GOBP_RESPONSE_TO_ACID_CHEMICAL | 0,5230079 0,9520965 0,0488971 0,3493523 0,9840276 | 109 |
| GOBP_NON_CANONICAL_WNT_SIGNALING_PATHWAY | 0,5119617 0,9479394 0,052805 0,3697087 0,9837336 | 66 |
| GOBP_PEPTIDYL_GLUTAMIC_ACID_MODIFICATION | 0,505992 0,9472208 0,0583453 0,4344639 0,9834023 | 22 |
| GOBP_RIBOSE_PHOSPHATE_BIOSYNTHETIC_PROCESS | 0,524377 0,9520965 0,0471642 0,3368491 0,9833779 | 166 |
| GOBP_POLY_A_PLUS_MRNA_EXPORT_FROM_NUCLEUS | 0,5134771 0,9484432 0,0582216 0,4456929 0,9832413 | 18 |
| GOBP_REGULATION_OF_SYNAPSE_STRUCTURE_OR_ACTIVITY | 0,5283422 0,9525709 0,0462303 0,3343228 0,9826782 | 182 |
| GOBP_REGULATION_OF_CHOLESTEROL_EFFLUX | 0,498094 0,9416872 0,0568864 0,4107676 0,9824518 | 31 |
| GOBP_KIDNEY_MORPHOGENESIS | 0,5250875 0,9520965 0,0504983 0,3619224 0,9820149 | 77 |
| GOBP_REGULATION_OF_FILOPODIUM_ASSEMBLY | 0,5140306 0,9485329 0,0555947 0,3895779 0,9819092 | 45 |
| GOBP_REGULATION_OF_METAL_ION_TRANSPORT | 0,5572917 0,9586744 0,0425878 0,3253082 0,9818231 | 303 |
| GOBP_CELLULAR_EXTRAVASATION | 0,5268949 0,9520965 0,0524828 0,3798552 0,9816912 | 52 |
| GOBP_POSITIVE_REGULATION_OF_ION_TRANSMEMBRANE_TRANSPORT | 0,5177384 0,9512582 0,048799 0,3434628 0,9815268 | 127 |
| GOBP_MAINTENANCE_OF_LOCATION_IN_CELL | 0,5161638 0,9496706 0,0476387 0,3350047 0,98152 | 171 |
| GOBP_ESTABLISHMENT_OF_EPITHELIAL_CELL_APICAL_BASAL_POLARITY | 0,5194631 0,9515789 0,0574877 0,4504242 0,9815063 | 17 |
| GOBP_RNA_EXPORT_FROM_NUCLEUS | 0,5117647 0,9479394 0,052057 0,3634196 0,981395 | 75 |
| GOBP_REGULATION_OF_HISTONE_H3_K4_METHYLATION | 0,5044136 0,9461218 0,0559429 0,4015525 0,9811367 | 35 |
| GOBP_NEGATIVE_REGULATION_OF_BINDING | 0,5217865 0,9520965 0,0476387 0,3402992 0,9809639 | 143 |
| GOBP_MYELOID_LEUKOCYTE_ACTIVATION | 0,525974 0,9520965 0,0469759 0,3372556 0,9796043 | 153 |
| GOBP_POSITIVE_REGULATION_OF_NUCLEOTIDE_BIOSYNTHETIC_PROCESS | 0,5243902 0,9520965 0,0574877 0,4643436 0,9794322 | 15 |
| GOBP_NEGATIVE_REGULATION_OF_VIRAL_LIFE_CYCLE | 0,5241287 0,9520965 0,057006 0,4560663 0,9791962 | 16 |
| GOBP_P38MAPK_CASCADE | 0,5266497 0,9520965 0,0542316 0,3857903 0,9789947 | 47 |
| GOBP_MAMMARY_GLAND_MORPHOGENESIS | 0,5094578 0,9479394 0,0554793 0,4005469 0,9786797 | 35 |
| GOBP_RNA_STABILIZATION | 0,5317073 0,9525709 0,0519512 0,3742413 0,9783301 | 56 |
| GOBP_POSITIVE_REGULATION_OF_IMMUNE_RESPONSE | 0,556701 0,95855 0,0421619 0,3214564 0,9782161 | 343 |
| GOBP_REGULATION_OF_AMPA_RECEPTOR_ACTIVITY | 0,5133333 0,9484432 0,0577309 0,4326052 0,978189 | 21 |
| GOBP_POSITIVE_REGULATION_OF_AXON_EXTENSION | 0,5258512 0,9520965 0,0540088 0,3969894 0,978186 | 37 |
| GOBP_EMBRYONIC_PLACENTA_MORPHOGENESIS | 0,5133333 0,9484432 0,0577309 0,4325765 0,9781239 | 21 |
| GOBP_OSTEOCLAST_DIFFERENTIATION | 0,5190476 0,9515789 0,0519512 0,3651865 0,9780041 | 68 |
| GOBP_PHOSPHATIDYLINOSITOL_3_KINASE_SIGNALING | 0,5319865 0,9525709 0,0481184 0,3471713 0,9778843 | 109 |
| GOBP_REGULATION_OF_CALCIUM_ION_TRANSMEMBRANE_TRANSPORT | 0,5265487 0,9520965 0,0479259 0,3437287 0,9776297 | 119 |
| GOBP_BONE_RESORPTION | 0,5283019 0,9525709 0,053677 0,3866402 0,9775246 | 44 |
| GOBP_HORMONE_METABOLIC_PROCESS | 0,5308776 0,9525709 0,0466015 0,3365533 0,9771225 | 152 |
| GOBP_REGULATION_OF_LIPID_TRANSPORT | 0,5300113 0,9525709 0,0487011 0,3494792 0,9766641 | 102 |
| GOBP_REGULATION_OF_INTRACELLULAR_ESTROGEN_RECEPTOR_SIGNALING_PATHWAY | 0,5121328 0,9479394 0,0558265 0,4100118 0,9764623 | 30 |
| GOBP_AORTIC_VALVE_MORPHOGENESIS | 0,5090674 0,9479394 0,0567672 0,4202499 0,976166 | 24 |
| GOBP_MEMBRANE_DEPOLARIZATION_DURING_ACTION_POTENTIAL | 0,5151515 0,9492027 0,0550211 0,4043901 0,9760502 | 32 |
| GOBP_NEGATIVE_REGULATION_OF_CYTOKINE_PRODUCTION_INVOLVED_IN_INFLAMMATORY_RESPONSE | 0,5288591 0,9525709 0,0566484 0,4478632 0,9759258 | 17 |
| GOBP_POSITIVE_REGULATION_OF_DEPHOSPHORYLATION | 0,5252153 0,9520965 0,052913 0,3788277 0,9757858 | 51 |
| GOBP_NUCLEOBASE_CONTAINING_COMPOUND_TRANSPORT | 0,5469083 0,9525767 0,0445071 0,3306713 0,9756746 | 195 |
| GOBP_NEGATIVE_REGULATION_OF_INTERLEUKIN_1_PRODUCTION | 0,5288591 0,9525709 0,0566484 0,4477257 0,9756263 | 17 |
| GOBP_STEROL_TRANSPORT | 0,536 0,9525709 0,0486034 0,3531701 0,9756196 | 89 |
| GOBP_REGULATION_OF_JUN_KINASE_ACTIVITY | 0,531407 0,9525709 0,0533478 0,3829686 0,975318 | 48 |
| GOBP_MEMBRANE_LIPID_BIOSYNTHETIC_PROCESS | 0,5348066 0,9525709 0,0471642 0,3406537 0,9749202 | 131 |
| GOBP_CERAMIDE_BIOSYNTHETIC_PROCESS | 0,5351942 0,9525709 0,0514265 0,3709894 0,974874 | 58 |
| GOBP_POSITIVE_REGULATION_OF_STEROL_TRANSPORT | 0,51341 0,9484432 0,0557104 0,4093281 0,974834 | 30 |
| GOBP_RESPIRATORY_ELECTRON_TRANSPORT_CHAIN | 0,5423341 0,9525709 0,0481184 0,3530682 0,9744809 | 93 |
| GOBP_SMALL_GTPASE_MEDIATED_SIGNAL_TRANSDUCTION | 0,5861366 0,9690661 0,0392759 0,3160642 0,9743071 | 454 |
| GOBP_NEGATIVE_REGULATION_OF_RESPONSE_TO_CYTOKINE_STIMULUS | 0,5376214 0,9525709 0,0512184 0,3709312 0,9739462 | 57 |
| GOBP_NEGATIVE_REGULATION_OF_LYMPHOCYTE_ACTIVATION | 0,5402951 0,9525709 0,0479259 0,3490666 0,973182 | 101 |
| GOBP_CHEMICAL_SYNAPTIC_TRANSMISSION_POSTSYNAPTIC | 0,5390805 0,9525709 0,0486034 0,3539223 0,9730917 | 83 |
| GOBP_CELLULAR_NITROGEN_COMPOUND_CATABOLIC_PROCESS | 0,5979487 0,9706649 0,0386372 0,316882 0,9730337 | 402 |
| GOBP_REGULATION_OF_CELL_SHAPE | 0,5404208 0,9525709 0,0467883 0,3401871 0,9729607 | 133 |
| GOBP_DETECTION_OF_STIMULUS | 0,540154 0,9525709 0,0465084 0,3395599 0,9729338 | 137 |
| GOBP_POSITIVE_REGULATION_OF_DEFENSE_RESPONSE_TO_VIRUS_BY_HOST | 0,5109961 0,9479394 0,05653 0,41724 0,9723225 | 25 |
| GOBP_SMALL_MOLECULE_BIOSYNTHETIC_PROCESS | 0,5912334 0,9706649 0,0388759 0,315243 0,9721948 | 445 |
| GOBP_SKELETAL_MUSCLE_CELL_DIFFERENTIATION | 0,5415648 0,9525709 0,0512184 0,3760735 0,971918 | 52 |
| GOBP_POSITIVE_REGULATION_OF_TRANSPORTER_ACTIVITY | 0,5497143 0,9535954 0,0474483 0,3503414 0,9713546 | 97 |
| GOBP_MULTICELLULAR_ORGANISMAL_SIGNALING | 0,5393134 0,9525709 0,046882 0,3406455 0,9709694 | 125 |
| GOBP_RETINA_HOMEOSTASIS | 0,5389447 0,9525709 0,0526973 0,3811357 0,9706503 | 48 |
| GOBP_NEUROINFLAMMATORY_RESPONSE | 0,5251613 0,9520965 0,0551352 0,41343 0,9704601 | 26 |
| GOBP_NEUROMUSCULAR_PROCESS | 0,5422222 0,9525709 0,0467883 0,3408893 0,9701745 | 122 |
| GOBP_REGULATION_OF_PROTEIN_SECRETION | 0,5651709 0,9620966 0,0431037 0,3288812 0,9698722 | 194 |
| GOBP_POSITIVE_REGULATION_OF_BONE_MINERALIZATION | 0,5245283 0,9520965 0,0540088 0,3906479 0,9697762 | 40 |
| GOBP_NEURAL_NUCLEUS_DEVELOPMENT | 0,5436893 0,9525767 0,0507028 0,3693385 0,9697643 | 57 |
| GOBP_CELLULAR_TRANSITION_METAL_ION_HOMEOSTASIS | 0,5451429 0,9525767 0,04783 0,3510224 0,9696868 | 89 |
| GOBP_POSITIVE_REGULATION_OF_CALCIUM_ION_TRANSPORT_INTO_CYTOSOL | 0,53375 0,9525709 0,052913 0,3908995 0,9693267 | 39 |
| GOBP_VASCULAR_ASSOCIATED_SMOOTH_MUSCLE_CELL_DIFFERENTIATION | 0,5251613 0,9520965 0,0551352 0,4129052 0,9692283 | 26 |
| GOBP_REGULATION_OF_VASCULAR_ASSOCIATED_SMOOTH_MUSCLE_CELL_DIFFERENTIATION | 0,5402145 0,9525709 0,0555947 0,4513761 0,9691262 | 16 |
| GOBP_HORMONE_TRANSPORT | 0,5662778 0,9621995 0,0426734 0,3268242 0,9690166 | 214 |
| GOBP_DETECTION_OF_BIOTIC_STIMULUS | 0,524 0,9520965 0,0567672 0,428378 0,9686306 | 21 |
| GOBP_RESPONSE_TO_DSRNA | 0,5271122 0,9520965 0,0538979 0,3942338 0,9686115 | 36 |
| GOBP_REGULATION_OF_SECRETION | 0,605102 0,9711099 0,0378488 0,3138345 0,9684661 | 461 |
| GOBP_NEGATIVE_REGULATION_OF_IMMUNE_SYSTEM_PROCESS | 0,5677083 0,9629256 0,0417397 0,3217815 0,9684482 | 277 |
| GOBP_PROTEIN_SUMOYLATION | 0,5414573 0,9525709 0,0524828 0,3802211 0,9683209 | 48 |
| GOBP_POSITIVE_REGULATION_OF_MACROPHAGE_ACTIVATION | 0,5355705 0,9525709 0,0560596 0,444346 0,9682616 | 17 |
| GOBP_MIDBRAIN_DEVELOPMENT | 0,5500589 0,9535954 0,048799 0,3586987 0,9679745 | 74 |
| GOBP_REGULATION_OF_HEART_GROWTH | 0,5387454 0,9525709 0,0517405 0,3757526 0,9678649 | 51 |
| GOBP_EXTRINSIC_APOPTOTIC_SIGNALING_PATHWAY_VIA_DEATH_DOMAIN_RECEPTORS | 0,5460993 0,9525767 0,0492918 0,3594777 0,9677215 | 73 |
| GOBP_ATP_SYNTHESIS_COUPLED_ELECTRON_TRANSPORT | 0,5518868 0,9546219 0,0487011 0,3596421 0,9676428 | 72 |
| GOBP_ORGANIC_HYDROXY_COMPOUND_TRANSPORT | 0,565635 0,9621995 0,0430173 0,3282215 0,9675093 | 192 |
| GOBP_NEGATIVE_REGULATION_OF_PHOSPHORUS_METABOLIC_PROCESS | 0,5969072 0,9706649 0,0389556 0,3168542 0,9674595 | 370 |
| GOBP_REGULATION_OF_LYMPHOCYTE_DIFFERENTIATION | 0,5480663 0,9532801 0,0460458 0,3379257 0,9671127 | 131 |
| GOBP_REGULATION_OF_CALCIUM_ION_TRANSPORT | 0,5667022 0,9621995 0,0429311 0,3285784 0,96637 | 187 |
| GOBP_MRNA_CLEAVAGE | 0,5327103 0,9525709 0,0560596 0,4345038 0,9663095 | 19 |
| GOBP_POSITIVE_REGULATION_OF_PROTEIN_IMPORT | 0,5308953 0,9525709 0,0535669 0,3932563 0,9662097 | 36 |
| GOBP_REGULATION_OF_CALCINEURIN_MEDIATED_SIGNALING | 0,5249042 0,9520965 0,0546808 0,4056693 0,9661203 | 30 |
| GOBP_TROPHECTODERMAL_CELL_DIFFERENTIATION | 0,5369128 0,9525709 0,0559429 0,4432486 0,9658703 | 17 |
| GOBP_CYTOKINE_PRODUCTION_INVOLVED_IN_IMMUNE_RESPONSE | 0,5333333 0,9525709 0,0507028 0,360476 0,9653888 | 68 |
| GOBP_NEGATIVE_REGULATION_OF_PROTEIN_SECRETION | 0,5473555 0,9525767 0,0510114 0,3745158 0,9646791 | 51 |
| GOBP_MAST_CELL_ACTIVATION_INVOLVED_IN_IMMUNE_RESPONSE | 0,5334174 0,9525709 0,0533478 0,3924817 0,9643067 | 36 |
| GOBP_NEGATIVE_REGULATION_OF_NECROTIC_CELL_DEATH | 0,5367156 0,9525709 0,0557104 0,4335244 0,9641313 | 19 |
| GOBP_NEUROEPITHELIAL_CELL_DIFFERENTIATION | 0,5339547 0,9525709 0,0558265 0,4257769 0,9637395 | 22 |
| GOBP_SIGNAL_TRANSDUCTION_IN_ABSENCE_OF_LIGAND | 0,5485855 0,9535954 0,0509083 0,3740383 0,9634492 | 51 |
| GOBP_MEMBRANE_BIOGENESIS | 0,5501814 0,9535954 0,0499914 0,366485 0,9631744 | 59 |
| GOBP_INTERFERON_GAMMA_MEDIATED_SIGNALING_PATHWAY | 0,5352863 0,9525709 0,0557104 0,4254809 0,9630695 | 22 |
| GOBP_INTRINSIC_APOPTOTIC_SIGNALING_PATHWAY_IN_RESPONSE_TO_DNA_DAMAGE_BY_P53_CLASS_MEDIATOR | 0,5474083 0,9525767 0,0522693 0,3845738 0,9629596 | 42 |
| GOBP_GLUTAMINE_FAMILY_AMINO_ACID_METABOLIC_PROCESS | 0,5406699 0,9525709 0,0502948 0,3617978 0,962684 | 66 |
| GOBP_REGULATION_OF_PHAGOCYTOSIS | 0,5559567 0,95855 0,0492918 0,3642068 0,9622218 | 62 |
| GOBP_PRIMARY_ALCOHOL_METABOLIC_PROCESS | 0,5625 0,9614707 0,04783 0,3576119 0,9621804 | 72 |
| GOBP_NOTOCHORD_DEVELOPMENT | 0,5469169 0,9525767 0,0550211 0,4480543 0,9619941 | 16 |
| GOBP_REGULATION_OF_ION_TRANSMEMBRANE_TRANSPORT | 0,6166495 0,9790646 0,037303 0,3146141 0,9613496 | 367 |
| GOBP_REGULATION_OF_SUPEROXIDE_METABOLIC_PROCESS | 0,5379494 0,9525709 0,0554793 0,4246575 0,9612056 | 22 |
| GOBP_NEGATIVE_REGULATION_OF_DEFENSE_RESPONSE_TO_VIRUS | 0,5303226 0,9525709 0,0546808 0,4094561 0,9611321 | 26 |
| GOBP_NEGATIVE_REGULATION_OF_EXOCYTOSIS | 0,546875 0,9525767 0,053677 0,4099062 0,9610678 | 27 |
| GOBP_TOOTH_MINERALIZATION | 0,5409396 0,9525709 0,0555947 0,4407657 0,9604599 | 17 |
| GOBP_POSITIVE_REGULATION_OF_VIRAL_GENOME_REPLICATION | 0,5455729 0,9525767 0,0537873 0,4072173 0,9599347 | 28 |
| GOBP_EXTERNAL_ENCAPSULATING_STRUCTURE_ORGANIZATION | 0,5868201 0,9690661 0,0404114 0,3208132 0,9595852 | 246 |
| GOBP_CELL_SURFACE_RECEPTOR_SIGNALING_PATHWAY_INVOLVED_IN_CELL_CELL_SIGNALING | 0,6289501 0,9808052 0,0359969 0,3103897 0,9595732 | 472 |
| GOBP_ASPARTATE_FAMILY_AMINO_ACID_BIOSYNTHETIC_PROCESS | 0,5447263 0,9525767 0,0550211 0,431341 0,9592756 | 19 |
| GOBP_POSITIVE_REGULATION_OF_INSULIN_SECRETION_INVOLVED_IN_CELLULAR_RESPONSE_TO_GLUCOSE_STIMULUS | 0,5346667 0,9525709 0,0558265 0,4240662 0,9588808 | 21 |
| GOBP_REGULATION_OF_CELLULAR_RESPONSE_TO_TRANSFORMING_GROWTH_FACTOR_BETA_STIMULUS | 0,5696629 0,9641632 0,0450443 0,3405318 0,9588042 | 110 |
| GOBP_PEPTIDYL_LYSINE_ACETYLATION | 0,5646552 0,9620966 0,0435378 0,3270585 0,9582387 | 171 |
| GOBP_NEGATIVE_REGULATION_OF_EXTRINSIC_APOPTOTIC_SIGNALING_PATHWAY | 0,5707657 0,9641936 0,0464155 0,3503783 0,9565881 | 79 |
| GOBP_INNATE_IMMUNE_RESPONSE | 0,6353535 0,9813188 0,0350901 0,3081587 0,9563174 | 495 |
| GOBP_RNA_DESTABILIZATION | 0,5606407 0,9613497 0,0466015 0,3457932 0,9562921 | 88 |
| GOBP_LYMPHOCYTE_HOMEOSTASIS | 0,5680982 0,963152 0,0491928 0,369015 0,9562012 | 53 |
| GOBP_POSITIVE_REGULATION_OF_EMBRYONIC_DEVELOPMENT | 0,5463087 0,9525767 0,0551352 0,4387858 0,9561456 | 17 |
| GOBP_NCRNA_3_END_PROCESSING | 0,5528967 0,9559287 0,0516356 0,3869844 0,9561282 | 38 |
| GOBP_BILE_ACID_AND_BILE_SALT_TRANSPORT | 0,5444744 0,9525767 0,0554793 0,4333922 0,9561048 | 18 |
| GOBP_DEPHOSPHORYLATION | 0,6278351 0,9808052 0,0366082 0,3129895 0,9556593 | 370 |
| GOBP_ENERGY_DERIVATION_BY_OXIDATION_OF_ORGANIC_COMPOUNDS | 0,603125 0,9706649 0,0389556 0,3172215 0,9556142 | 272 |
| GOBP_POSITIVE_REGULATION_OF_DEVELOPMENTAL_GROWTH | 0,5772627 0,9664677 0,0436251 0,3336938 0,9555869 | 135 |
| GOBP_NEGATIVE_REGULATION_OF_CILIUM_ASSEMBLY | 0,5463087 0,9525767 0,0551352 0,4383712 0,955242 | 17 |
| GOBP_SODIUM_ION_TRANSPORT | 0,5884861 0,9697116 0,0411544 0,3244372 0,9550114 | 189 |
| GOBP_POSITIVE_REGULATION_OF_GROWTH | 0,5778252 0,9664677 0,0419926 0,322974 0,954546 | 203 |
| GOBP_CELLULAR_COMPONENT_DISASSEMBLY | 0,6448311 0,9816504 0,0350151 0,3102048 0,9545428 | 418 |
| GOBP_CAMP_METABOLIC_PROCESS | 0,5386667 0,9525709 0,0554793 0,4221266 0,954495 | 21 |
| GOBP_MEGAKARYOCYTE_DIFFERENTIATION | 0,55375 0,9567971 0,0512184 0,3848258 0,9542655 | 39 |
| GOBP_NUCLEOSIDE_PHOSPHATE_CATABOLIC_PROCESS | 0,5647059 0,9620966 0,0475434 0,3533588 0,9542264 | 75 |
| GOBP_NEGATIVE_REGULATION_OF_LEUKOCYTE_APOPTOTIC_PROCESS | 0,556116 0,95855 0,0514265 0,3871566 0,9539578 | 37 |
| GOBP_CELLULAR_RESPONSE_TO_INTERFERON_BETA | 0,5503356 0,9535954 0,054794 0,4377479 0,9538839 | 17 |
| GOBP_CELLULAR_RESPONSE_TO_CHEMICAL_STRESS | 0,6104167 0,9761348 0,0383996 0,3165009 0,9538242 | 284 |
| GOBP_POSITIVE_REGULATION_OF_MAPK_CASCADE | 0,6322314 0,9808052 0,0363783 0,3129043 0,9536274 | 354 |
| GOBP_SOMATIC_STEM_CELL_POPULATION_MAINTENANCE | 0,5510719 0,9536513 0,0518458 0,390268 0,9535646 | 35 |
| GOBP_APICAL_PROTEIN_LOCALIZATION | 0,5555556 0,95855 0,054794 0,4519917 0,9533786 | 15 |
| GOBP_NADP_METABOLIC_PROCESS | 0,5510719 0,9536513 0,0518458 0,3900941 0,9531399 | 35 |
| GOBP_NUCLEAR_TRANSCRIBED_MRNA_CATABOLIC_PROCESS_DEADENYLATION_DEPENDENT_DECAY | 0,5722892 0,9645986 0,048022 0,361303 0,9531152 | 60 |
| GOBP_FIBROBLAST_GROWTH_FACTOR_RECEPTOR_SIGNALING_PATHWAY | 0,5702281 0,9641936 0,048022 0,3603534 0,9530509 | 63 |
| GOBP_PROTEIN_LOCALIZATION_TO_CHROMOSOME_TELOMERIC_REGION | 0,5502577 0,9535954 0,052913 0,4013398 0,9526379 | 29 |
| GOBP_ANTIGEN_PROCESSING_AND_PRESENTATION_OF_ENDOGENOUS_PEPTIDE_ANTIGEN | 0,5539084 0,9567971 0,0546808 0,4316926 0,9523554 | 18 |
| GOBP_G_PROTEIN_COUPLED_RECEPTOR_SIGNALING_PATHWAY_COUPLED_TO_CYCLIC_NUCLEOTIDE_SECOND_MESSENGER | 0,5388601 0,9525709 0,0541201 0,4099893 0,9523323 | 24 |
| GOBP_REGULATION_OF_OSTEOCLAST_DIFFERENTIATION | 0,5651075 0,9620966 0,0508054 0,3801542 0,9518932 | 42 |
| GOBP_EXTRINSIC_APOPTOTIC_SIGNALING_PATHWAY | 0,5997866 0,9706649 0,0403295 0,3236224 0,9517943 | 187 |
| GOBP_ANTIGEN_PROCESSING_AND_PRESENTATION_OF_ENDOGENOUS_ANTIGEN | 0,5505319 0,9535954 0,0543434 0,4233878 0,9513179 | 20 |
| GOBP_RESPONSE_TO_CARBOHYDRATE | 0,5925134 0,9706649 0,0409884 0,3236978 0,950999 | 174 |
| GOBP_REGULATION_OF_HORMONE_SECRETION | 0,5888651 0,9698138 0,041321 0,3238585 0,9509441 | 173 |
| GOBP_CYCLIC_NUCLEOTIDE_METABOLIC_PROCESS | 0,5470514 0,9525767 0,0519512 0,3908113 0,950744 | 33 |
| GOBP_MYELOID_LEUKOCYTE_MEDIATED_IMMUNITY | 0,5619048 0,9613497 0,0483118 0,3547771 0,9501265 | 68 |
| GOBP_REGULATION_OF_MUSCLE_CELL_DIFFERENTIATION | 0,5808989 0,9678213 0,0441523 0,3374479 0,950121 | 110 |
| GOBP_LONG_CHAIN_FATTY_ACYL_COA_METABOLIC_PROCESS | 0,5401554 0,9525709 0,0540088 0,4089898 0,9500107 | 24 |
| GOBP_NEGATIVE_REGULATION_OF_PHOSPHORYLATION | 0,6359876 0,9813188 0,0361492 0,313329 0,9498769 | 323 |
| GOBP_REGULATION_OF_ADENYLATE_CYCLASE_ACTIVITY | 0,5403646 0,9525709 0,0542316 0,4120848 0,948891 | 23 |
| GOBP_DETECTION_OF_ABIOTIC_STIMULUS | 0,5782857 0,9664677 0,0451344 0,3426705 0,948604 | 91 |
| GOBP_REGENERATION | 0,583878 0,9684899 0,0425023 0,3277965 0,9481963 | 147 |
| GOBP_POLYSACCHARIDE_METABOLIC_PROCESS | 0,5876289 0,9691491 0,0445071 0,3425887 0,9481408 | 90 |
| GOBP_PROTEIN_ADP_RIBOSYLATION | 0,5542929 0,9570214 0,0516356 0,3928216 0,9481282 | 32 |
| GOBP_PYRIDINE_NUCLEOTIDE_BIOSYNTHETIC_PROCESS | 0,5585106 0,9598923 0,053677 0,4217835 0,9477132 | 20 |
| GOBP_POSITIVE_REGULATION_OF_CELL_ADHESION | 0,6452282 0,9816504 0,0356176 0,3126269 0,9468875 | 329 |
| GOBP_HISTONE_H3_DEACETYLATION | 0,5610738 0,9613497 0,0538979 0,4342618 0,9462875 | 17 |
| GOBP_HOMOTYPIC_CELL_CELL_ADHESION | 0,5945626 0,9706649 0,0454059 0,3512637 0,9456093 | 73 |
| GOBP_MYELOID_CELL_DIFFERENTIATION | 0,6541275 0,9840117 0,0353157 0,3138471 0,9452541 | 296 |
| GOBP_REGULATION_OF_PROTEIN_CATABOLIC_PROCESS | 0,6498455 0,9819096 0,0349401 0,3097612 0,9452535 | 360 |
| GOBP_ACTIVATION_OF_INNATE_IMMUNE_RESPONSE | 0,563522 0,9619047 0,0507028 0,380548 0,9447034 | 40 |
| GOBP_SMOOTH_MUSCLE_CELL_PROLIFERATION | 0,592881 0,9706649 0,0427591 0,3332311 0,9442971 | 116 |
| GOBP_REGULATION_OF_DEFENSE_RESPONSE | 0,6690501 0,9872205 0,0331626 0,306302 0,9442208 | 433 |
| GOBP_CELLULAR_IRON_ION_HOMEOSTASIS | 0,5829268 0,9680138 0,0477342 0,3625768 0,9440247 | 54 |
| GOBP_REGULATION_OF_CD4_POSITIVE_ALPHA_BETA_T_CELL_DIFFERENTIATION | 0,5662043 0,9621995 0,0506004 0,386341 0,9439696 | 35 |
| GOBP_CARDIAC_RIGHT_VENTRICLE_MORPHOGENESIS | 0,5670241 0,9621995 0,0533478 0,43965 0,9439497 | 16 |
| GOBP_REGULATION_OF_PROTEIN_LOCALIZATION_TO_NUCLEUS | 0,5913621 0,9706649 0,0426734 0,3318837 0,9439459 | 118 |
| GOBP_DENDRITE_DEVELOPMENT | 0,607864 0,9738784 0,0395172 0,3183819 0,9437194 | 215 |
| GOBP_TOLL_LIKE_RECEPTOR_4_SIGNALING_PATHWAY | 0,5618687 0,9613497 0,0510114 0,3909763 0,9436743 | 32 |
| GOBP_AROMATIC_AMINO_ACID_FAMILY_METABOLIC_PROCESS | 0,5613333 0,9613497 0,0535669 0,417294 0,9435678 | 21 |
| GOBP_MACROPHAGE_ACTIVATION | 0,5630952 0,9616328 0,048215 0,3521721 0,9431503 | 68 |
| GOBP_DNA_DAMAGE_RESPONSE_SIGNAL_TRANSDUCTION_BY_P53_CLASS_MEDIATOR | 0,5775656 0,9664677 0,0471642 0,3528109 0,9430883 | 67 |
| GOBP_GLYCOPROTEIN_BIOSYNTHETIC_PROCESS | 0,6326743 0,9808052 0,036685 0,3132456 0,9430544 | 276 |
| GOBP_NEGATIVE_REGULATION_OF_OSSIFICATION | 0,5619174 0,9613497 0,0534572 0,4166178 0,9430079 | 22 |
| GOBP_CELLULAR_RESPONSE_TO_PEPTIDE | 0,6499478 0,9819096 0,0356176 0,3131839 0,942844 | 295 |
| GOBP_POSITIVE_REGULATION_OF_PROTEIN_SERINE_THREONINE_KINASE_ACTIVITY | 0,6032086 0,9706649 0,0401661 0,3208772 0,9427124 | 174 |
| GOBP_PROTEIN_CONTAINING_COMPLEX_DISASSEMBLY | 0,6271008 0,9808052 0,0375363 0,3162632 0,9426994 | 231 |
| GOBP_RESPONSE_TO_LEAD_ION | 0,5670241 0,9621995 0,0533478 0,4390273 0,9426127 | 16 |
| GOBP_VASODILATION | 0,5694444 0,9641632 0,0503964 0,3872339 0,9420783 | 34 |
| GOBP_SPHINGOLIPID_METABOLIC_PROCESS | 0,6026201 0,9706649 0,0411544 0,327649 0,9420254 | 139 |
| GOBP_T_HELPER_17_CELL_DIFFERENTIATION | 0,5606469 0,9613497 0,0541201 0,4268092 0,941582 | 18 |
| GOBP_NEURON_PROJECTION_REGENERATION | 0,5829146 0,9680138 0,049094 0,3696401 0,9413739 | 48 |
| GOBP_RESPONSE_TO_AXON_INJURY | 0,5770609 0,9664677 0,0472587 0,3541042 0,9409224 | 65 |
| GOBP_REGULATION_OF_CELL_SUBSTRATE_JUNCTION_ORGANIZATION | 0,5770609 0,9664677 0,0472587 0,354071 0,9408342 | 65 |
| GOBP_REGULATION_OF_HORMONE_LEVELS | 0,6642562 0,9872205 0,0340464 0,3086843 0,9406963 | 353 |
| GOBP_RNA_3_END_PROCESSING | 0,6015891 0,9706649 0,0430173 0,3374048 0,9406696 | 101 |
| GOBP_REGULATION_OF_MONOCYTE_DIFFERENTIATION | 0,5697051 0,9641632 0,0531298 0,4380467 0,9405072 | 16 |
| GOBP_POSITIVE_REGULATION_OF_CALCIUM_MEDIATED_SIGNALING | 0,5706667 0,9641936 0,052805 0,4158338 0,940266 | 21 |
| GOBP_REGULATION_OF_VOLTAGE_GATED_CALCIUM_CHANNEL_ACTIVITY | 0,5721584 0,9645986 0,0507028 0,394796 0,9402251 | 30 |
| GOBP_ADIPOSE_TISSUE_DEVELOPMENT | 0,5737705 0,9650381 0,0499914 0,3826465 0,9401419 | 36 |
| GOBP_NEGATIVE_REGULATION_OF_DEPHOSPHORYLATION | 0,5750315 0,9654878 0,0498907 0,3768801 0,9400565 | 41 |
| GOBP_INTEGRIN_ACTIVATION | 0,5718085 0,9645986 0,0525899 0,4183235 0,9399387 | 20 |
| GOBP_CARDIAC_CHAMBER_DEVELOPMENT | 0,6141125 0,9776358 0,0407404 0,3281435 0,9394789 | 136 |
| GOBP_NEGATIVE_REGULATION_OF_CYTOSKELETON_ORGANIZATION | 0,6074155 0,9735751 0,0407404 0,3249179 0,9394215 | 145 |
| GOBP_ICOSANOID_METABOLIC_PROCESS | 0,5714286 0,9645986 0,0475434 0,3507212 0,9392646 | 68 |
| GOBP_NEGATIVE_REGULATION_OF_MRNA_METABOLIC_PROCESS | 0,5942529 0,9706649 0,0441523 0,3415498 0,9390742 | 83 |
| GOBP_POSITIVE_REGULATION_OF_ENDOTHELIAL_CELL_MIGRATION | 0,6147727 0,9781472 0,0420772 0,3373972 0,9389161 | 99 |
| GOBP_CELLULAR_AMINO_ACID_BIOSYNTHETIC_PROCESS | 0,5873206 0,9690661 0,0465084 0,3533112 0,9386964 | 64 |
| GOBP_CYTOKINE_MEDIATED_SIGNALING_PATHWAY | 0,6522188 0,982332 0,0348652 0,3100269 0,9386712 | 315 |
| GOBP_T_CELL_ACTIVATION | 0,6687307 0,9872205 0,033677 0,3081508 0,9385128 | 350 |
| GOBP_NEGATIVE_REGULATION_OF_CELL_GROWTH | 0,6117137 0,9766609 0,0401661 0,3221886 0,9384078 | 160 |
| GOBP_EXOCYTIC_PROCESS | 0,5918854 0,9706649 0,0460458 0,3508129 0,9377477 | 67 |
| GOBP_COLLAGEN_FIBRIL_ORGANIZATION | 0,5975309 0,9706649 0,0471642 0,3658358 0,9374895 | 49 |
| GOBP_CERAMIDE_METABOLIC_PROCESS | 0,6006865 0,9706649 0,0434507 0,3389567 0,9373859 | 88 |
| GOBP_T_CELL_MEDIATED_CYTOTOXICITY | 0,5730623 0,9647049 0,0503964 0,3918493 0,937204 | 31 |
| GOBP_3_UTR_MEDIATED_MRNA_STABILIZATION | 0,5754339 0,9655673 0,0524828 0,4209124 0,9360832 | 19 |
| GOBP_REGULATION_OF_INSULIN_SECRETION_INVOLVED_IN_CELLULAR_RESPONSE_TO_GLUCOSE_STIMULUS | 0,5895807 0,9698138 0,049094 0,3722285 0,9360649 | 43 |
| GOBP_ENERGY_HOMEOSTASIS | 0,5775536 0,9664677 0,0496901 0,3798201 0,9358806 | 37 |
| GOBP_SEMI_LUNAR_VALVE_DEVELOPMENT | 0,5719697 0,9645986 0,0501934 0,3876801 0,9357185 | 32 |
| GOBP_REGULATION_OF_CALCIUM_MEDIATED_SIGNALING | 0,5987805 0,9706649 0,0465084 0,3585982 0,9351129 | 55 |
| GOBP_INTRINSIC_APOPTOTIC_SIGNALING_PATHWAY_BY_P53_CLASS_MEDIATOR | 0,6105882 0,9761348 0,0439759 0,3460769 0,934562 | 75 |
| GOBP_HISTONE_H3_K4_METHYLATION | 0,599278 0,9706649 0,045862 0,3536826 0,934417 | 62 |
| GOBP_REGULATION_OF_TOR_SIGNALING | 0,6157355 0,9781472 0,0421619 0,3368566 0,9341088 | 95 |
| GOBP_CALCIUM_MEDIATED_SIGNALING | 0,6187364 0,9804 0,0398407 0,3240422 0,9341006 | 143 |
| GOBP_MAMMARY_GLAND_DEVELOPMENT | 0,6044944 0,9711099 0,0423318 0,3317356 0,9337735 | 108 |
| GOBP_CD4_POSITIVE_ALPHA_BETA_T_CELL_DIFFERENTIATION | 0,5946602 0,9706649 0,0466015 0,3552545 0,9335263 | 58 |
| GOBP_SNRNA_PROCESSING | 0,5640362 0,9620966 0,0519512 0,4005423 0,9334107 | 25 |
| GOBP_MRNA_3_END_PROCESSING | 0,6016847 0,9706649 0,045679 0,353029 0,9332949 | 61 |
| GOBP_MECHANORECEPTOR_DIFFERENTIATION | 0,6 0,9706649 0,0469759 0,3641951 0,9332852 | 49 |
| GOBP_POSITIVE_REGULATION_OF_CD4_POSITIVE_ALPHA_BETA_T_CELL_ACTIVATION | 0,5833333 0,9680138 0,0507028 0,3979701 0,9330824 | 27 |
| GOBP_POSITIVE_REGULATION_OF_MICROTUBULE_POLYMERIZATION_OR_DEPOLYMERIZATION | 0,584596 0,9688575 0,0491928 0,3834953 0,932983 | 34 |
| GOBP_MONOCYTE_DIFFERENTIATION | 0,5743329 0,9654446 0,0502948 0,3900296 0,9328518 | 31 |
| GOBP_VESICLE_MEDIATED_TRANSPORT_IN_SYNAPSE | 0,628877 0,9808052 0,0382417 0,3173494 0,9327881 | 182 |
| GOBP_PROTEIN_LOCALIZATION_TO_NUCLEUS | 0,6642336 0,9872205 0,0344919 0,3098101 0,9322446 | 268 |
| GOBP_MONOCARBOXYLIC_ACID_CATABOLIC_PROCESS | 0,6137387 0,9776358 0,0417397 0,3320251 0,9321286 | 105 |
| GOBP_CELLULAR_RESPONSE_TO_LIPID | 0,7017365 1 0,0308468 0,3023056 0,9319012 | 433 |
| GOBP_NEGATIVE_REGULATION_OF_OXIDATIVE_STRESS_INDUCED_INTRINSIC_APOPTOTIC_SIGNALING_PATHWAY | 0,5768194 0,9664677 0,052805 0,4223578 0,9317617 | 18 |
| GOBP_RESPONSE_TO_AMPHETAMINE | 0,5582902 0,9598923 0,0524828 0,4010304 0,9315224 | 24 |
| GOBP_SCHWANN_CELL_DEVELOPMENT | 0,5823755 0,9680138 0,0498907 0,3910943 0,9314094 | 30 |
| GOBP_MONOVALENT_INORGANIC_CATION_HOMEOSTASIS | 0,6157254 0,9781472 0,0408229 0,3273941 0,9311766 | 118 |
| GOBP_REGULATION_OF_RESPONSE_TO_OXIDATIVE_STRESS | 0,6156542 0,9781472 0,0432769 0,3418657 0,9310415 | 78 |
| GOBP_INOSITOL_LIPID_MEDIATED_SIGNALING | 0,6225383 0,9808052 0,0397596 0,3232229 0,9306044 | 142 |
| GOBP_REGULATION_OF_MYELOID_CELL_APOPTOTIC_PROCESS | 0,5795148 0,967661 0,0525899 0,4216912 0,9302913 | 18 |
| GOBP_POSITIVE_REGULATION_OF_AMINE_TRANSPORT | 0,5833333 0,9680138 0,0492918 0,3854124 0,9302451 | 32 |
| GOBP_MITOCHONDRION_LOCALIZATION | 0,5989848 0,9706649 0,0483118 0,3665448 0,9301569 | 47 |
| GOBP_NEGATIVE_REGULATION_OF_EPITHELIAL_CELL_DIFFERENTIATION | 0,5883838 0,9697116 0,0488971 0,3821103 0,9296134 | 34 |
| GOBP_CELLULAR_RESPONSE_TO_EXOGENOUS_DSRNA | 0,5790885 0,967379 0,0523759 0,4329475 0,9295591 | 16 |
| GOBP_CELLULAR_RESPONSE_TO_CARBOHYDRATE_STIMULUS | 0,6135857 0,9776358 0,0412376 0,3289781 0,9289908 | 112 |
| GOBP_PROTEIN_K48_LINKED_UBIQUITINATION | 0,6016746 0,9706649 0,0454059 0,3489901 0,928605 | 66 |
| GOBP_ACTIN_CYTOSKELETON_REORGANIZATION | 0,624 0,9808052 0,0416557 0,3348961 0,928531 | 97 |
| GOBP_REGULATION_OF_NUCLEOBASE_CONTAINING_COMPOUND_TRANSPORT | 0,5853659 0,9690661 0,0523759 0,4402037 0,9285144 | 15 |
| GOBP_NUCLEOSIDE_CATABOLIC_PROCESS | 0,5808625 0,9678213 0,0524828 0,4208327 0,9283973 | 18 |
| GOBP_POSITIVE_REGULATION_OF_PROTEIN_LOCALIZATION | 0,7081192 1 0,0307037 0,3028179 0,9283761 | 388 |
| GOBP_REGULATION_OF_ESTABLISHMENT_OF_PROTEIN_LOCALIZATION | 0,7094801 1 0,0302042 0,3008569 0,9279168 | 442 |
| GOBP_FATTY_ACID_METABOLIC_PROCESS | 0,6753653 0,9911514 0,0337507 0,3074225 0,9272181 | 300 |
| GOBP_RESPONSE_TO_VITAMIN_D | 0,59375 0,9706649 0,0498907 0,3954116 0,9270837 | 27 |
| GOBP_PROTEIN_ACYLATION | 0,6670168 0,9872205 0,034641 0,3108387 0,926843 | 235 |
| GOBP_RESPONSE_TO_PEPTIDE_HORMONE | 0,6853002 0,9963394 0,0326513 0,3059349 0,9265832 | 319 |
| GOBP_EPITHELIAL_TUBE_BRANCHING_INVOLVED_IN_LUNG_MORPHOGENESIS | 0,5818908 0,9680138 0,0518458 0,4093444 0,9265447 | 22 |
| GOBP_B_CELL_ACTIVATION | 0,6495726 0,9819096 0,036685 0,3147552 0,9257861 | 184 |
| GOBP_KERATINOCYTE_PROLIFERATION | 0,593947 0,9706649 0,0484088 0,3756483 0,9256012 | 37 |
| GOBP_REGULATION_OF_CHROMATIN_BINDING | 0,587217 0,9690661 0,0514265 0,4088931 0,9255232 | 22 |
| GOBP_REGULATION_OF_INTRINSIC_APOPTOTIC_SIGNALING_PATHWAY_IN_RESPONSE_TO_DNA_DAMAGE_BY_P53_CLASS_MEDIATOR | 0,5857909 0,9690661 0,0518458 0,4308955 0,9251533 | 16 |
| GOBP_STEROID_HORMONE_BIOSYNTHETIC_PROCESS | 0,5963542 0,9706649 0,0496901 0,394514 0,9249793 | 27 |
| GOBP_NEGATIVE_REGULATION_OF_LEUKOCYTE_CELL_CELL_ADHESION | 0,6304348 0,9808052 0,0412376 0,3344485 0,9249184 | 88 |
| GOBP_MITOCHONDRIAL_FISSION | 0,6020151 0,9706649 0,0477342 0,3743458 0,9249018 | 38 |
| GOBP_NEGATIVE_REGULATION_OF_CD4_POSITIVE_ALPHA_BETA_T_CELL_DIFFERENTIATION | 0,5862534 0,9690661 0,052057 0,4190874 0,9245471 | 18 |
| GOBP_PEPTIDYL_LYSINE_MODIFICATION | 0,7110423 1 0,0307037 0,3032573 0,9245429 | 355 |
| GOBP_BONE_DEVELOPMENT | 0,6417112 0,9813188 0,037303 0,3145909 0,9243381 | 175 |
| GOBP_CELLULAR_RESPONSE_TO_PEPTIDE_HORMONE_STIMULUS | 0,6729167 0,9908895 0,0338245 0,3089909 0,924239 | 240 |
| GOBP_REGULATION_OF_DNA_BINDING_TRANSCRIPTION_FACTOR_ACTIVITY | 0,7069143 1 0,0309901 0,303299 0,9242379 | 352 |
| GOBP_EXOCRINE_SYSTEM_DEVELOPMENT | 0,6032746 0,9706649 0,0476387 0,3740659 0,9242102 | 38 |
| GOBP_PURINE_CONTAINING_COMPOUND_CATABOLIC_PROCESS | 0,6168342 0,9790646 0,0465084 0,3628889 0,9241805 | 48 |
| GOBP_SODIUM_ION_HOMEOSTASIS | 0,5863808 0,9690661 0,0489954 0,3781719 0,9240095 | 35 |
| GOBP_ETHER_METABOLIC_PROCESS | 0,5738342 0,9650381 0,0512184 0,3977501 0,9239028 | 24 |
| GOBP_NAD_METABOLIC_PROCESS | 0,5751295 0,9654878 0,0511148 0,397446 0,9231964 | 24 |
| GOBP_CIRCULATORY_SYSTEM_PROCESS | 0,7206932 1 0,0294228 0,2993039 0,9230395 | 445 |
| GOBP_SENSORY_PERCEPTION_OF_PAIN | 0,6150179 0,9781472 0,0442407 0,3452875 0,9229962 | 69 |
| GOBP_MYELOID_LEUKOCYTE_MIGRATION | 0,635255 0,9813188 0,0394367 0,3227925 0,9228658 | 128 |
| GOBP_SODIUM_ION_TRANSMEMBRANE_TRANSPORT | 0,6405733 0,9813188 0,0387962 0,3221529 0,9223276 | 136 |
| GOBP_DNA_CATABOLIC_PROCESS_ENDONUCLEOLYTIC | 0,5912117 0,9706649 0,0511148 0,4074195 0,9221878 | 22 |
| GOBP_POSITIVE_REGULATION_OF_CREB_TRANSCRIPTION_FACTOR_ACTIVITY | 0,5892617 0,9698138 0,0516356 0,423169 0,9221154 | 17 |
| GOBP_NEGATIVE_REGULATION_OF_INTRACELLULAR_STEROID_HORMONE_RECEPTOR_SIGNALING_PATHWAY | 0,5895807 0,9698138 0,049094 0,3854848 0,9219817 | 31 |
| GOBP_REGULATION_OF_EXOCYTOSIS | 0,6523605 0,982332 0,036685 0,3141837 0,9217343 | 172 |
| GOBP_REGULATION_OF_RESPONSE_TO_CYTOKINE_STIMULUS | 0,639779 0,9813188 0,0389556 0,3219464 0,9213815 | 131 |
| GOBP_3_UTR_MEDIATED_MRNA_DESTABILIZATION | 0,5951743 0,9706649 0,0511148 0,4290666 0,9212267 | 16 |
| GOBP_NEGATIVE_REGULATION_OF_CELL_MIGRATION_INVOLVED_IN_SPROUTING_ANGIOGENESIS | 0,596206 0,9706649 0,0515309 0,4365428 0,9207924 | 15 |
| GOBP_EPITHELIAL_CELL_DIFFERENTIATION_INVOLVED_IN_KIDNEY_DEVELOPMENT | 0,6070529 0,9734092 0,0473534 0,3726692 0,9207593 | 38 |
| GOBP_REGULATION_OF_GLYCOPROTEIN_METABOLIC_PROCESS | 0,6242119 0,9808052 0,0461379 0,3689441 0,9202616 | 41 |
| GOBP_REGULATION_OF_DEPHOSPHORYLATION | 0,6304591 0,9808052 0,0402478 0,3262406 0,9196529 | 111 |
| GOBP_PROTEIN_LOCALIZATION_TO_EXTRACELLULAR_REGION | 0,6878252 0,9963394 0,0327242 0,3051508 0,919315 | 280 |
| GOBP_POSITIVE_REGULATION_OF_PROTEIN_SECRETION | 0,6477273 0,9816504 0,0396787 0,3302247 0,9189562 | 99 |
| GOBP_REGULATION_OF_CARDIAC_CONDUCTION | 0,5933333 0,9706649 0,0510114 0,4063981 0,9189304 | 21 |
| GOBP_RESPONSE_TO_HYDROGEN_PEROXIDE | 0,6373874 0,9813188 0,0400032 0,3270087 0,9185356 | 106 |
| GOBP_RIBOSOMAL_LARGE_SUBUNIT_ASSEMBLY | 0,5803109 0,9676999 0,0507028 0,3952718 0,9181461 | 24 |
| GOBP_TERPENOID_METABOLIC_PROCESS | 0,6267943 0,9808052 0,0435378 0,3455163 0,9179864 | 64 |
| GOBP_REGULATION_OF_GTP_BINDING | 0,601626 0,9706649 0,0511148 0,4351368 0,9178268 | 15 |
| GOBP_DOPAMINERGIC_NEURON_DIFFERENTIATION | 0,6013333 0,9706649 0,0503964 0,4057721 0,917515 | 21 |
| GOBP_DNA_TEMPLATED_TRANSCRIPTION_ELONGATION | 0,6376471 0,9813188 0,0419926 0,3396093 0,9170965 | 75 |
| GOBP_SUPEROXIDE_METABOLIC_PROCESS | 0,6281726 0,9808052 0,0461379 0,361389 0,9170733 | 47 |
| GOBP_AMEBOIDAL_TYPE_CELL_MIGRATION | 0,7319588 1 0,0292103 0,3001369 0,9166079 | 365 |
| GOBP_RESPONSE_TO_FLUID_SHEAR_STRESS | 0,6108247 0,9761348 0,0481184 0,3860933 0,9164481 | 29 |
| GOBP_SYNAPSE_ORGANIZATION | 0,7443299 1 0,0283628 0,300019 0,9160561 | 370 |
| GOBP_REGULATION_OF_PRODUCTION_OF_MOLECULAR_MEDIATOR_OF_IMMUNE_RESPONSE | 0,6418242 0,9813188 0,0391155 0,3231604 0,9157593 | 116 |
| GOBP_REGULATION_OF_BLOOD_PRESSURE | 0,6415094 0,9813188 0,0390355 0,3216964 0,9156715 | 120 |
| GOBP_APOPTOTIC_CELL_CLEARANCE | 0,6108312 0,9761348 0,04707 0,3704589 0,9152983 | 38 |
| GOBP_POSITIVE_REGULATION_OF_NEURON_PROJECTION_DEVELOPMENT | 0,6633554 0,9872205 0,0372254 0,3192883 0,9143345 | 135 |
| GOBP_HEART_DEVELOPMENT | 0,745694 1 0,0273777 0,2949858 0,9142863 | 492 |
| GOBP_REGULATION_OF_CATION_TRANSMEMBRANE_TRANSPORT | 0,7016632 1 0,0317091 0,3033992 0,914162 | 274 |
| GOBP_REGULATION_OF_DNA_TEMPLATED_TRANSCRIPTION_ELONGATION | 0,6331288 0,9808052 0,0442407 0,3527473 0,9140478 | 53 |
| GOBP_CAMERA_TYPE_EYE_MORPHOGENESIS | 0,6579545 0,9860042 0,0389556 0,3284212 0,9139375 | 99 |
| GOBP_LONG_TERM_MEMORY | 0,6174242 0,9791739 0,0466948 0,3755316 0,9136085 | 34 |
| GOBP_REGULATION_OF_PROTEIN_ACETYLATION | 0,6371158 0,9813188 0,0422468 0,339994 0,9130726 | 71 |
| GOBP_ATRIOVENTRICULAR_VALVE_DEVELOPMENT | 0,5992011 0,9706649 0,0504983 0,4033787 0,9130414 | 22 |
| GOBP_AXONAL_TRANSPORT_OF_MITOCHONDRION | 0,5956873 0,9706649 0,0513223 0,4137492 0,9127704 | 18 |
| GOBP_NEGATIVE_REGULATION_OF_TRANSPORT | 0,7362409 1 0,0292811 0,3010421 0,9123938 | 334 |
| GOBP_REGULATION_OF_PHOSPHATASE_ACTIVITY | 0,636472 0,9813188 0,0426734 0,3412426 0,9121836 | 69 |
| GOBP_SOMITOGENESIS | 0,638191 0,9813188 0,0449543 0,3581696 0,9121616 | 48 |
| GOBP_HISTONE_H2A_UBIQUITINATION | 0,603871 0,9711099 0,0487011 0,388579 0,9121265 | 26 |
| GOBP_CELL_CELL_SIGNALING_BY_WNT | 0,7349028 1 0,028645 0,2970539 0,9118314 | 395 |
| GOBP_DIVALENT_INORGANIC_CATION_HOMEOSTASIS | 0,7281153 1 0,0294228 0,2995186 0,9115991 | 342 |
| GOBP_TUBULIN_DEACETYLATION | 0,6005326 0,9706649 0,0503964 0,4026996 0,9115043 | 22 |
| GOBP_REGULATION_OF_PROTEIN_KINASE_C_SIGNALING | 0,5973154 0,9706649 0,0510114 0,4182663 0,911432 | 17 |
| GOBP_LONG_TERM_SYNAPTIC_POTENTIATION | 0,6476079 0,9816504 0,0409056 0,3362229 0,9111672 | 76 |
| GOBP_OLEFINIC_COMPOUND_BIOSYNTHETIC_PROCESS | 0,5973154 0,9706649 0,0510114 0,4180513 0,9109637 | 17 |
| GOBP_NEGATIVE_REGULATION_OF_RELEASE_OF_CYTOCHROME_C_FROM_MITOCHONDRIA | 0,5997305 0,9706649 0,0510114 0,4129129 0,9109255 | 18 |
| GOBP_REGULATION_OF_MRNA_CATABOLIC_PROCESS | 0,6663055 0,9872205 0,0361492 0,3130894 0,9107953 | 157 |
| GOBP_ASTROCYTE_DIFFERENTIATION | 0,6353791 0,9813188 0,0431902 0,3443486 0,9097569 | 62 |
| GOBP_DIACYLGLYCEROL_METABOLIC_PROCESS | 0,6031957 0,9706649 0,0501934 0,4019219 0,9097441 | 22 |
| GOBP_NEGATIVE_REGULATION_OF_WOUND_HEALING | 0,6236162 0,9808052 0,0450443 0,35318 0,9097225 | 51 |
| GOBP_MORPHOGENESIS_OF_AN_EPITHELIUM | 0,7535787 1 0,0273075 0,2951699 0,9095757 | 430 |
| GOBP_RNA_POLYMERASE_II_PREINITIATION_COMPLEX_ASSEMBLY | 0,6132813 0,9776358 0,0484088 0,3879242 0,9095288 | 27 |
| GOBP_INTRACELLULAR_ESTROGEN_RECEPTOR_SIGNALING_PATHWAY | 0,6402516 0,9813188 0,0448645 0,3596876 0,9093816 | 44 |
| GOBP_REGULATION_OF_SPROUTING_ANGIOGENESIS | 0,6191677 0,9804 0,0465084 0,3700976 0,9093102 | 36 |
| GOBP_REGULATION_OF_MYOTUBE_DIFFERENTIATION | 0,6257982 0,9808052 0,0466015 0,3818044 0,9092849 | 30 |
| GOBP_REGULATION_OF_MRNA_3_END_PROCESSING | 0,6276042 0,9808052 0,0473534 0,3857084 0,9092316 | 28 |
| GOBP_HEART_VALVE_MORPHOGENESIS | 0,6428571 0,9816504 0,0453152 0,3606541 0,9090083 | 45 |
| GOBP_REGULATION_OF_PEPTIDYL_LYSINE_ACETYLATION | 0,6326531 0,9808052 0,0432769 0,3436806 0,9089552 | 63 |
| GOBP_REGULATION_OF_MAST_CELL_ACTIVATION_INVOLVED_IN_IMMUNE_RESPONSE | 0,6093333 0,9758164 0,0497903 0,4019686 0,9089147 | 21 |
| GOBP_MONONUCLEAR_CELL_DIFFERENTIATION | 0,7291667 1 0,0299196 0,3009995 0,9083851 | 304 |
| GOBP_NEGATIVE_REGULATION_OF_PROTEASOMAL_PROTEIN_CATABOLIC_PROCESS | 0,6427673 0,9816504 0,0446854 0,3592092 0,908172 | 44 |
| GOBP_ACID_SECRETION | 0,628866 0,9808052 0,0467883 0,382494 0,9079048 | 29 |
| GOBP_SECOND_MESSENGER_MEDIATED_SIGNALING | 0,7013815 1 0,0327971 0,3062351 0,9077148 | 215 |
| GOBP_ESTABLISHMENT_OF_PROTEIN_LOCALIZATION_TO_TELOMERE | 0,6112601 0,9764048 0,0498907 0,4226441 0,9074371 | 16 |
| GOBP_NEGATIVE_REGULATION_OF_LYMPHOCYTE_APOPTOTIC_PROCESS | 0,6197917 0,9805897 0,0479259 0,3868875 0,9070981 | 27 |
| GOBP_CELLULAR_ION_HOMEOSTASIS | 0,756371 1 0,0269563 0,293302 0,9069788 | 474 |
| GOBP_RESPONSE_TO_TUMOR_CELL | 0,6213469 0,9808052 0,0466948 0,3792023 0,9069555 | 31 |
| GOBP_TORC1_SIGNALING | 0,6209877 0,9808052 0,0454059 0,3527787 0,9068362 | 50 |
| GOBP_REGULATION_OF_BONE_REMODELING | 0,6275253 0,9808052 0,0459538 0,3726758 0,9066608 | 34 |
| GOBP_POSITIVE_REGULATION_OF_HEMOPOIESIS | 0,6466667 0,9816504 0,0387167 0,3199914 0,9064159 | 115 |
| GOBP_STEROL_HOMEOSTASIS | 0,6507177 0,9819096 0,0418239 0,3410099 0,9060136 | 64 |
| GOBP_RESPONSE_TO_PEPTIDE | 0,7543679 1 0,0275183 0,2954899 0,9059099 | 388 |
| GOBP_PLASMA_LIPOPROTEIN_PARTICLE_CLEARANCE | 0,6054334 0,9712268 0,0487011 0,3886884 0,9057868 | 25 |
| GOBP_REGULATION_OF_ESTABLISHMENT_OR_MAINTENANCE_OF_CELL_POLARITY | 0,6049223 0,9711099 0,048799 0,3899115 0,9056952 | 24 |
| GOBP_VITAMIN_TRANSMEMBRANE_TRANSPORT | 0,6192412 0,9804 0,0497903 0,4292921 0,9054987 | 15 |
| GOBP_NEGATIVE_REGULATION_OF_CATABOLIC_PROCESS | 0,7211238 1 0,0304181 0,3003964 0,9049919 | 280 |
| GOBP_CLEAVAGE_INVOLVED_IN_RRNA_PROCESSING | 0,6236979 0,9808052 0,0476387 0,3857818 0,9045056 | 27 |
| GOBP_ADHERENS_JUNCTION_ORGANIZATION | 0,6466837 0,9816504 0,0450443 0,3586043 0,903842 | 45 |
| GOBP_DENDRITIC_CELL_DIFFERENTIATION | 0,625 0,9808052 0,0475434 0,385412 0,9036387 | 27 |
| GOBP_PURINE_NUCLEOTIDE_CATABOLIC_PROCESS | 0,6518424 0,982332 0,0445071 0,3592554 0,9034406 | 43 |
| GOBP_MULTICELLULAR_ORGANISM_GROWTH | 0,6662971 0,9872205 0,0372254 0,3169466 0,9033144 | 123 |
| GOBP_HORMONE_MEDIATED_SIGNALING_PATHWAY | 0,6721311 0,9906617 0,0361492 0,3126359 0,903313 | 146 |
| GOBP_AUTONOMIC_NERVOUS_SYSTEM_DEVELOPMENT | 0,6375 0,9813188 0,0447749 0,3641121 0,9029011 | 39 |
| GOBP_NEGATIVE_REGULATION_OF_PROTEASOMAL_UBIQUITIN_DEPENDENT_PROTEIN_CATABOLIC_PROCESS | 0,6262626 0,9808052 0,0460458 0,3740628 0,9028513 | 32 |
| GOBP_REGULATION_OF_SYNAPTIC_VESICLE_EXOCYTOSIS | 0,6395062 0,9813188 0,044064 0,3523021 0,9028081 | 49 |
| GOBP_CANONICAL_WNT_SIGNALING_PATHWAY | 0,7273673 1 0,0299907 0,2997316 0,9027655 | 265 |
| GOBP_RESPONSE_TO_BACTERIUM | 0,7746914 1 0,0261842 0,2952436 0,9026113 | 374 |
| GOBP_AMIDE_TRANSPORT | 0,7161017 1 0,031637 0,3038336 0,9024923 | 222 |
| GOBP_REGULATION_OF_CHOLESTEROL_BIOSYNTHETIC_PROCESS | 0,6107383 0,9761348 0,0499914 0,414047 0,9022379 | 17 |
| GOBP_FATTY_ACID_BIOSYNTHETIC_PROCESS | 0,6603563 0,9872205 0,0378488 0,3192967 0,9018734 | 113 |
| GOBP_RESPONSE_TO_HEAT | 0,674685 0,9911514 0,0381629 0,3258025 0,9016838 | 90 |
| GOBP_INTRACELLULAR_STEROL_TRANSPORT | 0,6404639 0,9813188 0,0459538 0,3797553 0,9014039 | 29 |
| GOBP_REGULATION_OF_STEROL_TRANSPORT | 0,6497462 0,9819096 0,0445962 0,3551232 0,9011729 | 47 |
| GOBP_TOR_SIGNALING | 0,66 0,9871099 0,0377705 0,318109 0,901084 | 115 |
| GOBP_TRANSFORMING_GROWTH_FACTOR_BETA_RECEPTOR_SIGNALING_PATHWAY | 0,6890574 0,9963394 0,0345664 0,3086085 0,9009337 | 166 |
| GOBP_CELLULAR_RESPONSE_TO_BIOTIC_STIMULUS | 0,6916395 0,9978839 0,0344919 0,3084106 0,9003085 | 163 |
| GOBP_LIPID_CATABOLIC_PROCESS | 0,7330553 1 0,0297065 0,3009763 0,9001609 | 243 |
| GOBP_REGULATION_OF_RESPONSE_TO_REACTIVE_OXYGEN_SPECIES | 0,6417526 0,9813188 0,045862 0,3791269 0,8999125 | 29 |
| GOBP_PROTEIN_DESTABILIZATION | 0,6556544 0,9849234 0,0442407 0,3578024 0,8997868 | 43 |
| GOBP_BONE_REMODELING | 0,6542617 0,9840117 0,0417397 0,3400876 0,8994526 | 63 |
| GOBP_CALCIUM_ION_TRANSPORT_INTO_CYTOSOL | 0,6740331 0,9911514 0,0365315 0,3163093 0,8993676 | 117 |
| GOBP_CELL_CELL_SIGNALING_INVOLVED_IN_CARDIAC_CONDUCTION | 0,6132813 0,9776358 0,0484088 0,3905595 0,8993257 | 23 |
| GOBP_RAS_PROTEIN_SIGNAL_TRANSDUCTION | 0,7559709 1 0,0279403 0,2973388 0,898921 | 312 |
| GOBP_NEGATIVE_REGULATION_OF_TUMOR_NECROSIS_FACTOR_MEDIATED_SIGNALING_PATHWAY | 0,6226415 0,9808052 0,0492918 0,407105 0,8981128 | 18 |
| GOBP_REGULATION_OF_BIOLOGICAL_PROCESS_INVOLVED_IN_SYMBIOTIC_INTERACTION | 0,6620076 0,9872205 0,0438002 0,3571302 0,8980962 | 43 |
| GOBP_REGULATION_OF_CLATHRIN_DEPENDENT_ENDOCYTOSIS | 0,6226415 0,9808052 0,0492918 0,4065582 0,8969063 | 18 |
| GOBP_REGULATION_OF_CHEMOTAXIS | 0,6931447 0,9992881 0,0344919 0,3075533 0,8969024 | 162 |
| GOBP_PEPTIDE_TRANSPORT | 0,7071353 1 0,0325058 0,3046414 0,8968745 | 188 |
| GOBP_CEREBELLAR_CORTEX_FORMATION | 0,620506 0,9808052 0,0488971 0,3962022 0,8967977 | 22 |
| GOBP_REGULATION_OF_EXTRINSIC_APOPTOTIC_SIGNALING_PATHWAY | 0,6773836 0,9923443 0,0364549 0,313734 0,8965698 | 127 |
| GOBP_REGULATION_OF_ALPHA_BETA_T_CELL_DIFFERENTIATION | 0,6573604 0,9855311 0,044064 0,3532643 0,8964558 | 47 |
| GOBP_REGULATION_OF_SMAD_PROTEIN_SIGNAL_TRANSDUCTION | 0,6507732 0,9819096 0,0452247 0,3776502 0,8964073 | 29 |
| GOBP_MEMBRANE_LIPID_METABOLIC_PROCESS | 0,7095391 1 0,0326513 0,3048782 0,8957946 | 181 |
| GOBP_ALCOHOL_BIOSYNTHETIC_PROCESS | 0,6751947 0,9911514 0,036762 0,3160581 0,8956329 | 116 |
| GOBP_AMINO_SUGAR_METABOLIC_PROCESS | 0,6460957 0,9816504 0,0445071 0,3624637 0,8955444 | 38 |
| GOBP_B_CELL_APOPTOTIC_PROCESS | 0,6266667 0,9808052 0,048506 0,3960044 0,8954287 | 21 |
| GOBP_REGULATION_OF_PROTEIN_DEPOLYMERIZATION | 0,6864802 0,9963394 0,0381629 0,3272227 0,8952078 | 81 |
| GOBP_REGULATION_OF_CYTOSOLIC_CALCIUM_ION_CONCENTRATION | 0,7293869 1 0,0306323 0,3007306 0,8943513 | 224 |
| GOBP_FOREBRAIN_REGIONALIZATION | 0,6253369 0,9808052 0,049094 0,4053991 0,8943493 | 18 |
| GOBP_FORELIMB_MORPHOGENESIS | 0,6419271 0,9813188 0,0463228 0,3813798 0,8941848 | 27 |
| GOBP_NEGATIVE_REGULATION_OF_RESPONSE_TO_OXIDATIVE_STRESS | 0,6253369 0,9808052 0,049094 0,4052632 0,8940495 | 18 |
| GOBP_GLUCOSAMINE_CONTAINING_COMPOUND_METABOLIC_PROCESS | 0,6243523 0,9808052 0,0473534 0,3848538 0,893947 | 24 |
| GOBP_METHYLGUANOSINE_CAP_DECAPPING | 0,6327913 0,9808052 0,048799 0,4237976 0,8939092 | 15 |
| GOBP_LYMPHOID_PROGENITOR_CELL_DIFFERENTIATION | 0,6241611 0,9808052 0,0489954 0,4100917 0,893619 | 17 |
| GOBP_AMINE_TRANSPORT | 0,679669 0,9931997 0,0392759 0,3319491 0,8936139 | 73 |

| GOBP_T_CELL_HOMEOSTASIS | 0,6401515 0,9813188 0,0450443 0,370156 0,8934216 | | 32 |
| --- | --- | --- | --- |
| GOBP_NEGATIVE_REGULATION_OF_COAGULATION | 0,6323714 0,9808052 0,0453152 0,3669022 0,8925794 | | 33 |
| GOBP_LEUKOCYTE_TETHERING_OR_ROLLING | 0,6306667 0,9808052 0,048215 0,3947039 0,892488 | | 21 |
| GOBP_REGULATION_OF_POLYSACCHARIDE_METABOLIC_PROCESS | 0,6519546 0,982332 0,0441523 0,3631934 0,8923469 | | 36 |
| GOBP_METAL_ION_HOMEOSTASIS | 0,7887755 1 0,024779 0,2892438 0,8919656 | | 459 |
| GOBP_CENTRAL_NERVOUS_SYSTEM_NEURON_AXONOGENESIS | 0,6323714 0,9808052 0,0453152 0,366601 0,8918467 | | 33 |
| GOBP_RESPONSE_TO_ETHANOL | 0,6879271 0,9963394 0,0369933 0,3220543 0,891638 | | 87 |
| GOBP_NEGATIVE_REGULATION_OF_CELL_CYCLE_G1_S_PHASE_TRANSITION | 0,6686321 0,9872205 0,0399219 0,331332 0,8914724 | | 72 |
| GOBP_MRNA_EXPORT_FROM_NUCLEUS | 0,6658537 0,9872205 0,0416557 0,3408666 0,8910829 | | 56 |
| GOBP_NUCLEOTIDE_SALVAGE | 0,6355014 0,9813188 0,0486034 0,4224318 0,8910283 | | 15 |
| GOBP_CENTRAL_NERVOUS_SYSTEM_PROJECTION_NEURON_AXONOGENESIS | 0,6235446 0,9808052 0,0473534 0,3823456 0,8910056 | | 25 |
| GOBP_BARBED_END_ACTIN_FILAMENT_CAPPING | 0,6284953 0,9808052 0,0483118 0,3935752 0,8908514 | | 22 |
| GOBP_RESPONSE_TO_CAMP | 0,6761905 0,991889 0,0398407 0,3325268 0,8905381 | | 68 |
| GOBP_CALCINEURIN_MEDIATED_SIGNALING | 0,65 0,9819096 0,043888 0,3590825 0,8904289 | | 39 |
| GOBP_CELL_GROWTH | 0,7930328 1 0,0247086 0,2890647 0,8899077 | | 425 |
| GOBP_MULTICELLULAR_ORGANISMAL_HOMEOSTASIS | 0,7890947 1 0,025201 0,2901854 0,8898697 | | 399 |
| GOBP_CELLULAR_RESPONSE_TO_RETINOIC_ACID | 0,6634146 0,9872205 0,0418239 0,3411968 0,8897353 | | 55 |
| GOBP_REGULATION_OF_ESTABLISHMENT_OF_PROTEIN_LOCALIZATION_TO_MITOCHONDRION | 0,6763959 0,991889 0,0427591 0,3520819 0,889545 | | 46 |
| GOBP_CARBOHYDRATE_DERIVATIVE_CATABOLIC_PROCESS | 0,7080132 1 0,0338984 0,3099049 0,8893935 | | 138 |
| GOBP_LIPID_OXIDATION | 0,7023945 1 0,036073 0,3205559 0,8889065 | | 95 |
| GOBP_AMINOGLYCAN_METABOLIC_PROCESS | 0,6898876 0,9965975 0,0362255 0,3157385 0,8887448 | | 108 |
| GOBP_NEGATIVE_REGULATION_OF_RECEPTOR_SIGNALING_PATHWAY_VIA_JAK_STAT | 0,6329787 0,9808052 0,0479259 0,3955392 0,8887442 | | 20 |
| GOBP_STRIATED_MUSCLE_ADAPTATION | 0,6519546 0,982332 0,0441523 0,3605881 0,8884928 | | 37 |
| GOBP_REACTIVE_OXYGEN_SPECIES_METABOLIC_PROCESS | 0,719697 1 0,0324331 0,3039198 0,8882291 | | 167 |
| GOBP_CELLULAR_METABOLIC_COMPOUND_SALVAGE | 0,6287193 0,9808052 0,0469759 0,3810928 0,8880861 | | 25 |
| GOBP_INTRINSIC_APOPTOTIC_SIGNALING_PATHWAY_IN_RESPONSE_TO_OXIDATIVE_STRESS | 0,6687737 0,9872205 0,0431037 0,3545425 0,8877622 | | 42 |
| GOBP_SMOOTH_MUSCLE_CELL_APOPTOTIC_PROCESS | 0,6329787 0,9808052 0,0479259 0,3950518 0,8876493 | | 20 |
| GOBP_REGULATION_OF_COAGULATION | 0,6683417 0,9872205 0,0428451 0,3485319 0,8876172 | | 48 |
| GOBP_PROTEIN_NEDDYLATION | 0,6295302 0,9808052 0,0486034 0,4072858 0,8875049 | | 17 |
| GOBP_CELLULAR_RESPONSE_TO_STEROID_HORMONE_STIMULUS | 0,7156757 1 0,0326513 0,3036206 0,887354 | | 168 |
| GOBP_REGULATION_OF_SIGNAL_TRANSDUCTION_BY_P53_CLASS_MEDIATOR | 0,7115165 1 0,0354665 0,3191353 0,8871318 | | 98 |
| GOBP_POSITIVE_REGULATION_OF_VASCULAR_ASSOCIATED_SMOOTH_MUSCLE_CELL_PROLIFERATION | 0,6662371 0,9872205 0,0441523 0,3733518 0,8862043 | | 29 |
| GOBP_POSITIVE_REGULATION_OF_CARBOHYDRATE_METABOLIC_PROCESS | 0,6734694 0,9910789 0,0404114 0,335032 0,8860818 | | 63 |
| GOBP_POSITIVE_REGULATION_OF_PHOSPHATASE_ACTIVITY | 0,6614583 0,9872205 0,0449543 0,3756138 0,8854357 | | 28 |
| GOBP_INHIBITORY_SYNAPSE_ASSEMBLY | 0,6463415 0,9816504 0,04783 0,4196859 0,8852364 | | 15 |
| GOBP_REGULATION_OF_VESICLE_MEDIATED_TRANSPORT | 0,8040816 1 0,0237209 0,2868916 0,8848715 | | 439 |
| GOBP_POSITIVE_REGULATION_OF_CYTOSKELETON_ORGANIZATION | 0,7198276 1 0,0322153 0,3020752 0,8846061 | | 170 |
| GOBP_RESPONSE_TO_ISOQUINOLINE_ALKALOID | 0,6351531 0,9813188 0,04783 0,3907344 0,8844212 | | 22 |
| GOBP_LIPID_LOCALIZATION | 0,7850467 1 0,0259736 0,2917351 0,8843218 | | 335 |
| GOBP_PYRIMIDINE_RIBONUCLEOTIDE_METABOLIC_PROCESS | 0,6403622 0,9813188 0,0461379 0,3794182 0,8841838 | | 25 |
| GOBP_RESPONSE_TO_HYPEROXIA | 0,6476965 0,9816504 0,0477342 0,4186328 0,883015 | | 15 |
| GOBP_MESENCHYMAL_TO_EPITHELIAL_TRANSITION | 0,6476965 0,9816504 0,0477342 0,4183324 0,8823814 | | 15 |
| GOBP_MAINTENANCE_OF_LOCATION | 0,7653806 1 0,0275183 0,2932777 0,8823524 | | 258 |
| GOBP_TISSUE_REMODELING | 0,71 1 0,0343431 0,3098871 0,8820979 | | 121 |
| GOBP_REGULATION_OF_MUSCLE_HYPERTROPHY | 0,6654321 0,9872205 0,0422468 0,3430907 0,8819327 | | 50 |
| GOBP_REGULATION_OF_TUMOR_NECROSIS_FACTOR_MEDIATED_SIGNALING_PATHWAY | 0,6772554 0,9923443 0,0427591 0,3503328 0,8810024 | | 43 |
| GOBP_RESPONSE_TO_INSULIN | 0,7569002 1 0,0289981 0,2969773 0,8800874 | | 216 |
| GOBP_MITOCHONDRION_MORPHOGENESIS | 0,6409574 0,9813188 0,0473534 0,3916807 0,8800745 | | 20 |
| GOBP_TRANSCRIPTION_INITIATION_FROM_RNA_POLYMERASE_II_PROMOTER | 0,7070588 1 0,0372254 0,3258293 0,8798843 | | 75 |
| GOBP_NUCLEOSIDE_BISPHOSPHATE_METABOLIC_PROCESS | 0,7184685 1 0,0344175 0,313826 0,8798839 | | 103 |
| GOBP_DNA_CATABOLIC_PROCESS | 0,6541935 0,9840117 0,0450443 0,3745254 0,8791379 | | 26 |
| GOBP_REGULATION_OF_SODIUM_ION_TRANSPORT | 0,7082847 1 0,036762 0,324402 0,8791325 | | 76 |
| GOBP_REGULATION_OF_VIRAL_PROCESS | 0,7123894 1 0,0339724 0,3064968 0,8777852 | | 130 |
| GOBP_EXTRACELLULAR_MATRIX_DISASSEMBLY | 0,6887755 0,9963394 0,0421619 0,3482199 0,8776686 | | 45 |
| GOBP_MIRNA_METABOLIC_PROCESS | 0,6507115 0,9819096 0,0454059 0,3766068 0,8776322 | | 25 |
| GOBP_CELLULAR_RESPONSE_TO_DSRNA | 0,6506667 0,9819096 0,0467883 0,3878633 0,8770203 | | 21 |
| GOBP_REGULATION_OF_BIOMINERALIZATION | 0,705951 1 0,0369161 0,3232034 0,8769576 | | 77 |
| GOBP_RNA_POLYADENYLATION | 0,6840102 0,9963394 0,0422468 0,3455355 0,876843 | | 47 |
| GOBP_ORGANIC_ANION_TRANSPORT | 0,7816092 1 0,0265351 0,2912108 0,8762275 | | 260 |
| GOBP_PROTEIN_LOCALIZATION_TO_SYNAPSE | 0,6892382 0,9963394 0,0396787 0,3333482 0,8760862 | | 59 |
| GOBP_RETINOIC_ACID_RECEPTOR_SIGNALING_PATHWAY | 0,6645161 0,9872205 0,0443293 0,3730485 0,8756712 | | 26 |
| GOBP_RESPONSE_TO_TRANSFORMING_GROWTH_FACTOR_BETA | 0,7677625 1 0,0282219 0,2953092 0,8755763 | | 214 |
| GOBP_SPLEEN_DEVELOPMENT | 0,6645161 0,9872205 0,0443293 0,3729735 0,8754951 | | 26 |
| GOBP_REGULATION_OF_T_CELL_DIFFERENTIATION | 0,7194164 1 0,0341946 0,3104803 0,874536 | | 109 |
| GOBP_PEPTIDYL_LYSINE_DIMETHYLATION | 0,6484375 0,9819096 0,045862 0,3796601 0,8742279 | | 23 |
| GOBP_REGULATION_OF_CALCIUM_ION_TRANSPORT_INTO_CYTOSOL | 0,7208481 1 0,0363783 0,3239211 0,8741248 | | 74 |
| GOBP_REGULATION_OF_ATP_DEPENDENT_ACTIVITY | 0,6883273 0,9963394 0,0395172 0,330678 0,8736397 | | 62 |
| GOBP_SERINE_FAMILY_AMINO_ACID_BIOSYNTHETIC_PROCESS | 0,6353887 0,9813188 0,0481184 0,406877 0,8735845 | | 16 |
| GOBP_FATTY_ACID_DERIVATIVE_METABOLIC_PROCESS | 0,6858191 0,9963394 0,0404114 0,3380184 0,8735691 | | 52 |
| GOBP_REGULATION_OF_ANOIKIS | 0,6626667 0,9872205 0,0459538 0,3862878 0,8734578 | | 21 |
| GOBP_PEPTIDYL_TYROSINE_MODIFICATION | 0,8002081 1 0,0250604 0,2899168 0,8734203 | | 280 |
| GOBP_OSTEOBLAST_PROLIFERATION | 0,6692708 0,9872205 0,0444181 0,3724788 0,8733153 | | 27 |
| GOBP_BLOOD_VESSEL_REMODELING | 0,6742424 0,9911514 0,0426734 0,3588695 0,8730723 | | 34 |
| GOBP_CELLULAR_RESPONSE_TO_MOLECULE_OF_BACTERIAL_ORIGIN | 0,7382256 1 0,0317813 0,3036516 0,8728528 | | 140 |
| GOBP_FOCAL_ADHESION_ASSEMBLY | 0,7219003 1 0,035542 0,3191994 0,8725511 | | 80 |
| GOBP_POSITIVE_REGULATION_OF_B_CELL_MEDIATED_IMMUNITY | 0,685567 0,9963394 0,0428451 0,3674607 0,8722209 | | 29 |
| GOBP_CARTILAGE_DEVELOPMENT_INVOLVED_IN_ENDOCHONDRAL_BONE_MORPHOGENESIS | 0,6645161 0,9872205 0,0443293 0,3715531 0,872161 | | 26 |
| GOBP_PROTEIN_POLYUBIQUITINATION | 0,7790698 1 0,0273075 0,2930263 0,8720441 | | 229 |
| GOBP_NEURAL_CREST_CELL_DIFFERENTIATION | 0,7149425 1 0,0356176 0,3171649 0,8720291 | | 83 |
| GOBP_REGULATION_OF_HAIR_CYCLE | 0,6551265 0,9849181 0,0464155 0,3851817 0,8718527 | | 22 |
| GOBP_POSITIVE_REGULATION_OF_RECEPTOR_MEDIATED_ENDOCYTOSIS | 0,6899619 0,9965975 0,0419082 0,346624 0,8716758 | | 43 |
| GOBP_CIRCADIAN_REGULATION_OF_GENE_EXPRESSION | 0,6907341 0,9969604 0,0393562 0,3292642 0,8704684 | | 61 |
| GOBP_ADENYLATE_CYCLASE_MODULATING_G_PROTEIN_COUPLED_RECEPTOR_SIGNALING_PATHWAY | 0,7467249 1 0,0310618 0,3025032 0,8697285 | | 139 |
| GOBP_NEGATIVE_REGULATION_OF_CALCIUM_MEDIATED_SIGNALING | 0,6461126 0,9816504 0,0473534 0,4047558 0,8690302 | | 16 |
| GOBP_NEGATIVE_REGULATION_OF_MRNA_CATABOLIC_PROCESS | 0,704878 1 0,0390355 0,3332354 0,8689744 | | 55 |
| GOBP_NUCLEOSIDE_MONOPHOSPHATE_CATABOLIC_PROCESS | 0,6639566 0,9872205 0,0466015 0,4118805 0,8687727 | | 15 |
| GOBP_EMBRYONIC_FORELIMB_MORPHOGENESIS | 0,6666667 0,9872205 0,045679 0,3841746 0,8686796 | | 21 |
| GOBP_ENDOCHONDRAL_BONE_MORPHOGENESIS | 0,6876543 0,9963394 0,0407404 0,3389111 0,8684924 | | 49 |
| GOBP_MORPHOGENESIS_OF_AN_ENDOTHELIUM | 0,6487936 0,9819096 0,0471642 0,4044973 0,8684751 | | 16 |
| GOBP_POSITIVE_REGULATION_OF_DNA_BINDING_TRANSCRIPTION_FACTOR_ACTIVITY | 0,781084 1 0,027448 0,2928525 0,8680471 | | 215 |
| GOBP_NUCLEAR_ENVELOPE_ORGANIZATION | 0,7060976 1 0,0389556 0,3328734 0,8680305 | | 55 |
| GOBP_REGULATION_OF_DEVELOPMENTAL_GROWTH | 0,8024948 1 0,0248494 0,2880719 0,86798 | | 274 |
| GOBP_EXOCYTOSIS | 0,8129572 1 0,0244269 0,2881092 0,8678643 | | 297 |
| GOBP_REGULATION_OF_MAP_KINASE_ACTIVITY | 0,7421452 1 0,0309901 0,2994003 0,8677314 | | 150 |
| GOBP_POSITIVE_REGULATION_OF_T_CELL_MEDIATED_CYTOTOXICITY | 0,6582109 0,9860042 0,0463228 0,3901018 0,8675622 | | 19 |
| GOBP_CELLULAR_RESPONSE_TO_PH | 0,6569149 0,9855311 0,0462303 0,3859816 0,8672692 | | 20 |
| GOBP_REGULATION_OF_MUSCLE_ORGAN_DEVELOPMENT | 0,6947637 1 0,0418239 0,364125 0,8671807 | | 30 |
| GOBP_PHAGOCYTOSIS | 0,7497291 1 0,0304894 0,2970354 0,8671479 | | 166 |
| GOBP_MEMBRANE_DEPOLARIZATION_DURING_CARDIAC_MUSCLE_CELL_ACTION_POTENTIAL | 0,6595745 0,9871099 0,0460458 0,3857943 0,8668483 | | 20 |
| GOBP_FIBROBLAST_APOPTOTIC_PROCESS | 0,6608812 0,9872205 0,0461379 0,3894914 0,8662048 | | 19 |
| GOBP_REGULATION_OF_MUSCLE_ADAPTATION | 0,7190476 1 0,0369933 0,3233137 0,8658646 | | 68 |
| GOBP_POSITIVE_REGULATION_OF_LIPID_LOCALIZATION | 0,7346466 1 0,0347157 0,3163755 0,8648317 | | 80 |
| GOBP_RIBONUCLEOPROTEIN_COMPLEX_BIOGENESIS | 0,8302658 1 0,0220116 0,2807465 0,864231 | | 414 |
| GOBP_GLYCOSYLATION | 0,7959831 1 0,0261842 0,290535 0,8640301 | | 224 |
| GOBP_REGULATION_OF_STEM_CELL_DIFFERENTIATION | 0,702439 1 0,0391956 0,331703 0,86364 | | 54 |
| GOBP_REGULATION_OF_DENDRITE_DEVELOPMENT | 0,7471396 1 0,0333093 0,3126863 0,8636356 | | 92 |
| GOBP_CARDIAC_CONDUCTION | 0,7362894 1 0,0349401 0,3181887 0,863351 | | 77 |
| GOBP_NEGATIVE_REGULATION_OF_MUSCLE_CELL_APOPTOTIC_PROCESS | 0,7045455 1 0,0406579 0,3576775 0,8633031 | | 32 |
| GOBP_NEGATIVE_REGULATION_OF_UBIQUITIN_DEPENDENT_PROTEIN_CATABOLIC_PROCESS | 0,7044025 1 0,0404935 0,3413098 0,8629177 | | 44 |
| GOBP_REGULATION_OF_RESPONSE_TO_BIOTIC_STIMULUS | 0,7977059 1 0,0253416 0,2867725 0,8627808 | | 258 |
| GOBP_MAINTENANCE_OF_PROTEIN_LOCATION | 0,7331042 1 0,0342688 0,3125848 0,8625864 | | 85 |
| GOBP_I_KAPPAB_KINASE_NF_KAPPAB_SIGNALING | 0,8048017 1 0,0249197 0,2884184 0,8625021 | | 245 |
| GOBP_TRANSCRIPTION_PREINITIATION_COMPLEX_ASSEMBLY | 0,695 1 0,0408229 0,3477394 0,8623012 | | 39 |
| GOBP_ARTERY_DEVELOPMENT | 0,7447552 1 0,0343431 0,3151874 0,8622819 | | 81 |
| GOBP_NEGATIVE_REGULATION_OF_LOCOMOTION | 0,8075314 1 0,0248494 0,286776 0,8621266 | | 262 |
| GOBP_VIRAL_GENOME_REPLICATION | 0,7441602 1 0,0321428 0,3046384 0,8615406 | | 114 |
| GOBP_PYRIDINE_CONTAINING_COMPOUND_BIOSYNTHETIC_PROCESS | 0,6692708 0,9872205 0,0444181 0,3740699 0,8613557 | | 23 |
| GOBP_POSITIVE_REGULATION_OF_SODIUM_ION_TRANSPORT | 0,706258 1 0,0410713 0,3615856 0,8611329 | | 30 |
| GOBP_REGULATION_OF_T_CELL_DIFFERENTIATION_IN_THYMUS | 0,670227 0,9882428 0,0454968 0,3871701 0,8610424 | | 19 |
| GOBP_RESPONSE_TO_PAIN | 0,677763 0,9923443 0,0448645 0,3803236 0,8608565 | | 22 |
| GOBP_POSITIVE_REGULATION_OF_INNATE_IMMUNE_RESPONSE | 0,7388826 1 0,033677 0,3104207 0,8608015 | | 95 |
| GOBP_MESENCHYMAL_CELL_DEVELOPMENT | 0,73979 1 0,0347157 0,3170838 0,8603529 | | 77 |
| GOBP_NUCLEOLAR_LARGE_RRNA_TRANSCRIPTION_BY_RNA_POLYMERASE_I | 0,6684636 0,9872205 0,0460458 0,3895696 0,8594278 | | 18 |
| GOBP_NEGATIVE_REGULATION_OF_OXIDATIVE_STRESS_INDUCED_CELL_DEATH | 0,6981132 1 0,0409056 0,3459234 0,8587485 | | 40 |
| GOBP_NEGATIVE_REGULATION_OF_INTRACELLULAR_TRANSPORT | 0,7092025 1 0,0390355 0,3313517 0,8586069 | | 53 |
| GOBP_NEGATIVE_REGULATION_OF_CELL_CELL_ADHESION | 0,7582781 1 0,0308468 0,2994198 0,8579047 | | 132 |
| GOBP_EYE_MORPHOGENESIS | 0,7763012 1 0,0298485 0,3005837 0,8572724 | | 124 |
| GOBP_REGULATION_OF_FATTY_ACID_OXIDATION | 0,7126289 1 0,0410713 0,3609424 0,8567488 | | 29 |
| GOBP_CYTOSOLIC_CALCIUM_ION_TRANSPORT | 0,7604857 1 0,0307037 0,2991017 0,8565267 | | 135 |
| GOBP_VASCULAR_ASSOCIATED_SMOOTH_MUSCLE_CELL_PROLIFERATION | 0,7098765 1 0,0392759 0,3331086 0,8562732 | | 50 |
| GOBP_REGULATION_OF_TRANSCRIPTION_BY_RNA_POLYMERASE_III | 0,7126289 1 0,0410713 0,3606921 0,8561547 | | 29 |
| GOBP_CELL_DEATH_IN_RESPONSE_TO_HYDROGEN_PEROXIDE | 0,6791721 0,9928927 0,0434507 0,3673439 0,8560461 | | 25 |
| GOBP_NEUTROPHIL_CHEMOTAXIS | 0,7097171 1 0,0391155 0,3323222 0,8559966 | | 51 |
| GOBP_NEGATIVE_REGULATION_OF_REGULATED_SECRETORY_PATHWAY | 0,6671141 0,9872205 0,0459538 0,3926649 0,8556449 | | 17 |
| GOBP_REGULATION_OF_B_CELL_MEDIATED_IMMUNITY | 0,7142857 1 0,0400846 0,3413187 0,8546503 | | 42 |
| GOBP_REGULATION_OF_NUCLEOTIDE_BIOSYNTHETIC_PROCESS | 0,7057292 1 0,0419926 0,3624854 0,854488 | | 28 |
| GOBP_GLYCOSIDE_METABOLIC_PROCESS | 0,6752022 0,9911514 0,0455878 0,3872707 0,8543564 | | 18 |
| GOBP_ACTION_POTENTIAL | 0,7552602 1 0,0312053 0,3003821 0,8543489 | | 118 |
| GOBP_NEGATIVE_REGULATION_OF_MAPK_CASCADE | 0,7690632 1 0,0294936 0,2963086 0,8541544 | | 143 |
| GOBP_RRNA_TRANSCRIPTION | 0,7032828 1 0,0407404 0,3510698 0,8540968 | | 34 |
| GOBP_NEGATIVE_REGULATION_OF_MUSCLE_HYPERTROPHY | 0,6864516 0,9963394 0,0428451 0,3638505 0,8540803 | | 26 |
| GOBP_GLYCEROPHOSPHOLIPID_CATABOLIC_PROCESS | 0,6787565 0,9928927 0,0435378 0,3676799 0,8540552 | | 24 |
| GOBP_TRANSITION_METAL_ION_HOMEOSTASIS | 0,7553311 1 0,0318535 0,3031728 0,8539527 | | 109 |
| GOBP_POSITIVE_REGULATION_OF_PROTEIN_CATABOLIC_PROCESS | 0,8097768 1 0,0255523 0,2887411 0,8537927 | | 212 |
| GOBP_POSITIVE_REGULATION_OF_SECRETION | 0,8181818 1 0,0247086 0,2866604 0,8532144 | | 227 |
| GOBP_TRANSITION_METAL_ION_TRANSPORT | 0,7517241 1 0,0332359 0,3101839 0,8528351 | | 83 |
| GOBP_MESENCHYMAL_CELL_DIFFERENTIATION | 0,8093717 1 0,0256927 0,2884799 0,8527167 | | 202 |
| GOBP_CYTOKINE_PRODUCTION_INVOLVED_IN_INFLAMMATORY_RESPONSE | 0,7099622 1 0,0402478 0,3417987 0,8525524 | | 41 |
| GOBP_DNA_TEMPLATED_TRANSCRIPTION_INITIATION | 0,752784 1 0,031637 0,3017 0,8519612 | | 112 |
| GOBP_RNA_PHOSPHODIESTER_BOND_HYDROLYSIS_EXONUCLEOLYTIC | 0,7099622 1 0,0402478 0,3415208 0,8518593 | | 41 |
| GOBP_RESPONSE_TO_GONADOTROPIN | 0,68 0,993297 0,0447749 0,3764503 0,8512138 | | 21 |
| GOBP_DETECTION_OF_STIMULUS_INVOLVED_IN_SENSORY_PERCEPTION | 0,7390777 1 0,0366082 0,3239888 0,8506905 | | 57 |
| GOBP_REGULATION_OF_CELLULAR_AMIDE_METABOLIC_PROCESS | 0,8495394 | 1 0,0207088 0,276388 0,8504838 | 418 |
| GOBP_SERINE_FAMILY_AMINO_ACID_METABOLIC_PROCESS | 0,7179161 | 1 0,0400846 0,3554975 0,8502596 | 31 |
| GOBP_NEGATIVE_REGULATION_OF_I_KAPPAB_KINASE_NF_KAPPAB_SIGNALING | 0,7166247 | 1 0,0397596 0,3439516 0,8498063 | 38 |
| GOBP_NUCLEAR_TRANSPORT | 0,8418314 | 1 0,0222268 0,2819276 0,8497343 | 282 |
| GOBP_REGULATION_OF_LIPID_LOCALIZATION | 0,7655556 | 1 0,0307037 0,2982711 0,8490329 | 121 |
| GOBP_REGULATION_OF_MEMBRANE_POTENTIAL | 0,8544892 | 1 0,0208547 0,2792276 0,8488807 | 338 |
| GOBP_POSITIVE_REGULATION_OF_NF_KAPPAB_TRANSCRIPTION_FACTOR_ACTIVITY | 0,7838137 | 1 0,0294228 0,2976236 0,8488133 | 126 |
| GOBP_SULFUR_COMPOUND_TRANSPORT | 0,7195431 | 1 0,0399219 0,3342529 0,8482117 | 47 |
| GOBP_COLLAGEN_METABOLIC_PROCESS | 0,7511792 | 1 0,0344919 0,3150378 0,8476319 | 72 |
| GOBP_CELLULAR_RESPONSE_TO_INSULIN_STIMULUS | 0,8008565 | 1 0,0265351 0,2882738 0,8475759 | 177 |
| GOBP_DICARBOXYLIC_ACID_TRANSPORT | 0,7658824 | 1 0,0334562 0,3138075 0,84742 | 75 |
| GOBP_ISOPRENOID_METABOLIC_PROCESS | 0,7586207 | 1 0,0327971 0,3080544 0,8469801 | 83 |
| GOBP_NEGATIVE_REGULATION_OF_T_CELL_RECEPTOR_SIGNALING_PATHWAY | 0,6808511 | 0,993958 0,0445962 0,376916 0,8468994 | 20 |
| GOBP_RIBONUCLEOTIDE_CATABOLIC_PROCESS | 0,7242694 | 1 0,0396787 0,3366699 0,8466437 | 43 |
| GOBP_REGULATION_OF_INTEGRIN_MEDIATED_SIGNALING_PATHWAY | 0,696477 | 1 0,0444181 0,4013254 0,846509 | 15 |
| GOBP_POSITIVE_REGULATION_OF_CATABOLIC_PROCESS | 0,8795918 | 1 0,0183209 0,2742754 0,8464288 | 462 |
| GOBP_REGULATION_OF_CELLULAR_PH | 0,7535377 | 1 0,0343431 0,3144571 0,8460694 | 72 |
| GOBP_MAMMARY_GLAND_EPITHELIAL_CELL_DIFFERENTIATION | 0,6729223 | 0,9908895 0,0454968 0,3939235 0,8457726 | 16 |
| GOBP_COLLAGEN_BIOSYNTHETIC_PROCESS | 0,7234848 | 1 0,0394367 0,3503033 0,8455045 | 32 |
| GOBP_GENERATION_OF_PRECURSOR_METABOLITES_AND_ENERGY | 0,863869 | 1 0,0196776 0,2745136 0,8447859 | 420 |
| GOBP_MULTICELLULAR_ORGANISMAL_RESPONSE_TO_STRESS | 0,7337349 | 1 0,0366082 0,3199434 0,844009 | 60 |
| GOBP_LEUKOCYTE_DIFFERENTIATION | 0,8676923 | 1 0,0195288 0,2749486 0,8438484 | 392 |
| GOBP_C21_STEROID_HORMONE_METABOLIC_PROCESS | 0,6993548 | 1 0,0419926 0,3593949 0,8436215 | 26 |
| GOBP_POSITIVE_REGULATION_OF_ISOTYPE_SWITCHING | 0,6981865 | 1 0,0422468 0,3630666 0,8433392 | 24 |
| GOBP_CHONDROCYTE_DIFFERENTIATION | 0,7748571 | 1 0,0314929 0,3041279 0,8432234 | 97 |
| GOBP_MAST_CELL_ACTIVATION | 0,7281922 | 1 0,0391956 0,3364686 0,8425058 | 42 |
| GOBP_ESTABLISHMENT_OF_MITOCHONDRION_LOCALIZATION | 0,7280928 | 1 0,0400846 0,3549191 0,8424518 | 29 |
| GOBP_INSULIN_SECRETION_INVOLVED_IN_CELLULAR_RESPONSE_TO_GLUCOSE_STIMULUS | 0,7359413 | 1 0,0371479 0,3259243 0,8423133 | 52 |
| GOBP_TRICARBOXYLIC_ACID_CYCLE | 0,7266922 | 1 0,0397596 0,3536166 0,8421544 | 30 |
| GOBP_NEGATIVE_REGULATION_OF_AUTOPHAGOSOME_ASSEMBLY | 0,7059621 | 1 0,0438002 0,3988322 0,8412502 | 15 |
| GOBP_REGULATION_OF_GTPASE_ACTIVITY | 0,8644421 | 1 0,0207818 0,2787665 0,8411421 | 305 |
| GOBP_BLOOD_VESSEL_ENDOTHELIAL_CELL_MIGRATION | 0,7900114 | 1 0,0302042 0,301653 0,8409951 | 101 |
| GOBP_KERATAN_SULFATE_METABOLIC_PROCESS | 0,6791946 | 0,9928927 0,0451344 0,3858017 0,8406895 | 17 |
| GOBP_POSITIVE_REGULATION_OF_ESTABLISHMENT_OF_PROTEIN_LOCALIZATION | 0,8447917 | 1 0,0220834 0,2791935 0,8405476 | 264 |
| GOBP_ZYMOGEN_ACTIVATION | 0,7411168 | 1 0,0385579 0,3326274 0,8403925 | 46 |
| GOBP_FATTY_ACYL_COA_METABOLIC_PROCESS | 0,7263556 | 1 0,0391956 0,3438449 0,8401363 | 35 |
| GOBP_ANTEROGRADE_AXONAL_TRANSPORT | 0,7273869 | 1 0,0389556 0,3298665 0,8400814 | 48 |
| GOBP_POSITIVE_REGULATION_OF_RHO_PROTEIN_SIGNAL_TRANSDUCTION | 0,7057292 | 1 0,0419926 0,3646615 0,8396913 | 23 |
| GOBP_POSITIVE_REGULATION_OF_HORMONE_SECRETION | 0,7757437 | 1 0,0314929 0,3036167 0,8396529 | 88 |
| GOBP_VITAMIN_BIOSYNTHETIC_PROCESS | 0,6823056 | 0,9955038 0,0448645 0,3907435 0,8389451 | 16 |
| GOBP_REGULATION_OF_GENERATION_OF_PRECURSOR_METABOLITES_AND_ENERGY | 0,7953281 | 1 0,0288568 0,2959326 0,8386022 | 116 |
| GOBP_RENAL_WATER_HOMEOSTASIS | 0,7127371 | 1 0,0433637 0,3974181 0,8382672 | 15 |
| GOBP_STEM_CELL_DIFFERENTIATION | 0,8181818 | 1 0,0253416 0,2849328 0,8381344 | 178 |
| GOBP_REGULATION_OF_TRANSPOSITION | 0,6872483 | 0,9963394 0,0445962 0,3843058 0,8374296 | 17 |
| GOBP_SYNAPTIC_VESICLE_EXOCYTOSIS | 0,7856328 | 1 0,0307037 0,3018921 0,8371443 | 96 |
| GOBP_ESTABLISHMENT_OF_PROTEIN_LOCALIZATION_TO_ORGANELLE | 0,8769231 | 1 0,0188533 0,2724723 0,836161 | 397 |
| GOBP_REGULATION_OF_PROTEIN_CONTAINING_COMPLEX_ASSEMBLY | 0,8728025 | 1 0,0196776 0,2744203 0,8352416 | 345 |
| GOBP_POSITIVE_REGULATION_OF_POTASSIUM_ION_TRANSPORT | 0,7383648 | 1 0,0383206 0,3361999 0,8346099 | 40 |
| GOBP_POSITIVE_REGULATION_OF_NUCLEOTIDE_METABOLIC_PROCESS | 0,7369759 | 1 0,0388759 0,348707 0,8340185 | 31 |
| GOBP_VIRAL_TRANSCRIPTION | 0,7371069 | 1 0,0383996 0,3297748 0,8337545 | 44 |
| GOBP_RESPONSE_TO_STEROL_DEPLETION | 0,6885906 | 0,9963394 0,0445071 0,382616 0,8337475 | 17 |
| GOBP_POSITIVE_REGULATION_OF_SMALL_GTPASE_MEDIATED_SIGNAL_TRANSDUCTION | 0,7416974 | 1 0,0370705 0,323416 0,8330561 | 51 |
| GOBP_TRACHEA_DEVELOPMENT | 0,7181572 | 1 0,0430173 0,3949295 0,8330181 | 15 |
| GOBP_PURINE_CONTAINING_COMPOUND_METABOLIC_PROCESS | 0,878225 | 1 0,0191547 0,2733853 0,8326377 | 348 |
| GOBP_REGULATION_OF_INSULIN_RECEPTOR_SIGNALING_PATHWAY | 0,754878 | 1 0,0358449 0,319627 0,8321982 | 54 |
| GOBP_RESPONSE_TO_ACTIVITY | 0,7395062 | 1 0,0373806 0,3246562 0,8319627 | 49 |
| GOBP_RECEPTOR_LOCALIZATION_TO_SYNAPSE | 0,7503075 | 1 0,0365315 0,3228625 0,8316303 | 51 |
| GOBP_NEGATIVE_REGULATION_OF_RESPONSE_TO_WOUNDING | 0,7634409 | 1 0,0343431 0,3128965 0,8314255 | 65 |
| GOBP_POSITIVE_REGULATION_OF_RESPONSE_TO_BIOTIC_STIMULUS | 0,8022099 | 1 0,0280811 0,2923705 0,8313021 | 117 |
| GOBP_CD4_POSITIVE_ALPHA_BETA_T_CELL_ACTIVATION | 0,7777778 | 1 0,0329431 0,3091747 0,8303057 | 71 |
| GOBP_POSITIVE_REGULATION_OF_CHONDROCYTE_DIFFERENTIATION | 0,7102426 | 1 0,0432769 0,3763638 0,8302946 | 18 |
| GOBP_GMP_METABOLIC_PROCESS | 0,716129 | 1 0,0409056 0,3534741 0,8297235 | 26 |
| GOBP_VESICLE_TARGETING_ROUGH_ER_TO_CIS_GOLGI | 0,7208672 | 1 0,0428451 0,3933531 0,8296931 | 15 |
| GOBP_NUCLEOSIDE_TRIPHOSPHATE_BIOSYNTHETIC_PROCESS | 0,7818396 | 1 0,0325785 0,3083424 0,8296174 | 72 |
| GOBP_CELL_DIFFERENTIATION_IN_HINDBRAIN | 0,7115903 | 1 0,0431902 0,3760506 0,8296038 | 18 |
| GOBP_TUMOR_NECROSIS_FACTOR_MEDIATED_SIGNALING_PATHWAY | 0,7862069 | 1 0,0310618 0,3016275 0,8293097 | 83 |
| GOBP_CARDIAC_CONDUCTION_SYSTEM_DEVELOPMENT | 0,7448454 | 1 0,0390355 0,3492708 0,8290446 | 29 |
| GOBP_HEMATOPOIETIC_STEM_CELL_DIFFERENTIATION | 0,7192755 | 1 0,0408229 0,3557012 0,8289145 | 25 |
| GOBP_POSITIVE_REGULATION_OF_HYDROLASE_ACTIVITY | 0,8960245 | 1 0,0169971 0,268786 0,8285662 | 454 |
| GOBP_REGULATION_OF_INNATE_IMMUNE_RESPONSE | 0,8244854 | 1 0,0256225 0,2834013 0,8282185 | 164 |
| GOBP_NEGATIVE_REGULATION_OF_FATTY_ACID_METABOLIC_PROCESS | 0,7283622 | 1 0,0415718 0,3657842 0,8279468 | 22 |
| GOBP_PEPTIDYL_SERINE_MODIFICATION | 0,8813736 | 1 0,0194542 0,2747881 0,8279389 | 281 |
| GOBP_CELL_COMMUNICATION_BY_ELECTRICAL_COUPLING_INVOLVED_IN_CARDIAC_CONDUCTION | 0,7283622 | 1 0,0415718 0,3657783 0,8279335 | 22 |
| GOBP_GLYCEROPHOSPHOLIPID_BIOSYNTHETIC_PROCESS | 0,8427205 | 1 0,0233667 0,2801227 0,8272563 | 197 |
| GOBP_ADENYLATE_CYCLASE_ACTIVATING_G_PROTEIN_COUPLED_RECEPTOR_SIGNALING_PATHWAY | 0,798627 | 1 0,0300618 0,2989879 0,826852 | 88 |
| GOBP_REGULATION_OF_PROTEIN_AUTOPHOSPHORYLATION | 0,7490542 | 1 0,0377705 0,335291 0,8261604 | 37 |
| GOBP_REGULATION_OF_IMMUNOGLOBULIN_PRODUCTION | 0,7619632 | 1 0,0356933 0,3188112 0,8261118 | 53 |
| GOBP_NEGATIVE_REGULATION_OF_PROTEOLYSIS_INVOLVED_IN_CELLULAR_PROTEIN_CATABOLIC_PROCESS | 0,776699 | 1 0,0342688 0,3142288 0,8257202 | 58 |
| GOBP_NEGATIVE_REGULATION_OF_DNA_BINDING_TRANSCRIPTION_FACTOR_ACTIVITY | 0,8194748 | 1 0,0264649 0,2869693 0,8257121 | 141 |
| GOBP_NEGATIVE_REGULATION_OF_MONONUCLEAR_CELL_MIGRATION | 0,7183288 | 1 0,0427591 0,3742262 0,8255789 | 18 |
| GOBP_V_D_J_RECOMBINATION | 0,7116155 | 1 0,0427591 0,3711586 0,8254336 | 19 |
| GOBP_RNA_CATABOLIC_PROCESS | 0,8665276 | 1 0,0206358 0,2742902 0,8252268 | 258 |
| GOBP_NEGATIVE_REGULATION_OF_ANION_TRANSPORT | 0,7050938 | 1 0,0433637 0,3840856 0,8246503 | 16 |
| GOBP_POSITIVE_REGULATION_BY_HOST_OF_VIRAL_TRANSCRIPTION | 0,7050938 | 1 0,0433637 0,3840647 0,8246053 | 16 |
| GOBP_NUCLEAR_BODY_ORGANIZATION | 0,7050938 | 1 0,0433637 0,3840424 0,8245575 | 16 |
| GOBP_VASCULAR_ENDOTHELIAL_CELL_PROLIFERATION | 0,7073826 | 1 0,0432769 0,3783282 0,8244041 | 17 |
| GOBP_RNA_SURVEILLANCE | 0,7262873 | 1 0,0425023 0,3908451 0,824403 | 15 |
| GOBP_VESICLE_DOCKING_INVOLVED_IN_EXOCYTOSIS | 0,7553594 | 1 0,0373806 0,3373744 0,8243268 | 35 |
| GOBP_NEGATIVE_REGULATION_OF_SUPRAMOLECULAR_FIBER_ORGANIZATION | 0,8135224 | 1 0,0266755 0,2850029 0,8240171 | 145 |
| GOBP_INSULIN_RECEPTOR_SIGNALING_PATHWAY | 0,815402 | 1 0,0285039 0,2948525 0,8240029 | 102 |
| GOBP_POSITIVE_REGULATION_OF_PROTEOLYSIS | 0,8850932 | 1 0,0188533 0,2718811 0,8237572 | 320 |
| GOBP_TYROSINE_PHOSPHORYLATION_OF_STAT_PROTEIN | 0,7628362 | 1 0,0354665 0,3186808 0,8235932 | 52 |
| GOBP_CARDIAC_EPITHELIAL_TO_MESENCHYMAL_TRANSITION | 0,7575758 | 1 0,037303 0,3411946 0,8235193 | 32 |
| GOBP_RESPONSE_TO_MONOSACCHARIDE | 0,8223185 | 1 0,025763 0,2836431 0,8235071 | 152 |
| GOBP_POSITIVE_REGULATION_OF_CHEMOTAXIS | 0,8165345 | 1 0,0284333 0,2946514 0,8234409 | 102 |
| GOBP_VERY_LONG_CHAIN_FATTY_ACID_METABOLIC_PROCESS | 0,7484036 | 1 0,0383996 0,3456088 0,8230834 | 30 |
| GOBP_POSITIVE_REGULATION_OF_INTERFERON_GAMMA_PRODUCTION | 0,7604938 | 1 0,036073 0,3201315 0,8229149 | 50 |
| GOBP_STEROID_HORMONE_MEDIATED_SIGNALING_PATHWAY | 0,8175751 | 1 0,027448 0,290854 0,8225572 | 114 |
| GOBP_NEGATIVE_REGULATION_OF_RNA_CATABOLIC_PROCESS | 0,7840095 | 1 0,0330163 0,3077141 0,8225414 | 67 |
| GOBP_CALCIUM_ION_TRANSMEMBRANE_TRANSPORT | 0,8704284 | 1 0,0204894 0,2752386 0,8224388 | 238 |
| GOBP_SEQUESTERING_OF_CALCIUM_ION | 0,8004587 | 1 0,0300618 0,2976686 0,8224108 | 94 |
| GOBP_REGULATION_OF_SMALL_GTPASE_MEDIATED_SIGNAL_TRANSDUCTION | 0,8761707 | 1 0,0198261 0,2729556 0,8221186 | 265 |
| GOBP_BIOLOGICAL_PROCESS_INVOLVED_IN_INTERACTION_WITH_SYMBIONT | 0,7852029 | 1 0,0329431 0,3075318 0,8220541 | 67 |
| GOBP_NEGATIVE_REGULATION_OF_NIK_NF_KAPPAB_SIGNALING | 0,7330729 | 1 0,0402478 0,3568909 0,8217982 | 23 |
| GOBP_ERBB_SIGNALING_PATHWAY | 0,8175676 | 1 0,0280811 0,2929151 0,8212553 | 103 |
| GOBP_REGULATION_OF_POTASSIUM_ION_TRANSMEMBRANE_TRANSPORTER_ACTIVITY | 0,7639594 | 1 0,0371479 0,3248658 0,8207826 | 46 |
| GOBP_POSITIVE_REGULATION_OF_STEROID_BIOSYNTHETIC_PROCESS | 0,7317073 | 1 0,0421619 0,3891043 0,8207311 | 15 |
| GOBP_TRANSCRIPTION_ELONGATION_FROM_RNA_POLYMERASE_II_PROMOTER | 0,7864078 | 1 0,033677 0,3123172 0,8206971 | 58 |
| GOBP_CELLULAR_RESPONSE_TO_OXYGEN_LEVELS | 0,8224917 | 1 0,0266755 0,286605 0,8205536 | 136 |
| GOBP_REGULATION_OF_MRNA_POLYADENYLATION | 0,7140957 | 1 0,042417 0,3650057 0,820138 | 20 |
| GOBP_POSITIVE_REGULATION_OF_DEFENSE_RESPONSE | 0,8580576 | 1 0,0225843 0,2787311 0,8197659 | 187 |
| GOBP_REGULATION_OF_CARBOHYDRATE_METABOLIC_PROCESS | 0,8367568 | 1 0,0247086 0,2817273 0,8195342 | 156 |
| GOBP_PROSTANOID_METABOLIC_PROCESS | 0,7604035 | 1 0,0370705 0,3353704 0,8194301 | 35 |
| GOBP_ACTIVIN_RECEPTOR_SIGNALING_PATHWAY | 0,7651515 | 1 0,036839 0,336778 0,8193272 | 34 |
| GOBP_REGULATION_OF_HORMONE_METABOLIC_PROCESS | 0,7585769 | 1 0,0375363 0,3425376 0,8192628 | 31 |
| GOBP_POST_EMBRYONIC_DEVELOPMENT | 0,8058824 | 1 0,0309901 0,302871 0,8178867 | 75 |
| GOBP_REGULATION_OF_OSSIFICATION | 0,8182857 | 1 0,0287861 0,2949799 0,8178597 | 97 |
| GOBP_REGULATION_OF_MAST_CELL_ACTIVATION | 0,7473958 | 1 0,0393562 0,3465736 0,8169791 | 28 |
| GOBP_POSITIVE_REGULATION_OF_COLLAGEN_METABOLIC_PROCESS | 0,7180851 | 1 0,0421619 0,3634247 0,8165856 | 20 |
| GOBP_AXONAL_FASCICULATION | 0,7336884 | 1 0,0412376 0,3606221 0,8162625 | 22 |
| GOBP_MYELOID_LEUKOCYTE_DIFFERENTIATION | 0,8385699 | 1 0,0247086 0,2805308 0,8160806 | 157 |
| GOBP_NEGATIVE_REGULATION_OF_PROTEIN_MODIFICATION_BY_SMALL_PROTEIN_CONJUGATION_OR_REMOVAL | 0,8109339 | 1 0,0290688 0,294557 0,8155089 | 87 |
| GOBP_NEGATIVE_REGULATION_OF_ATP_METABOLIC_PROCESS | 0,742228 | 1 0,0394367 0,3509933 0,815295 | 24 |
| GOBP_POSITIVE_REGULATION_OF_PROTEIN_DEPOLYMERIZATION | 0,7345013 | 1 0,0417397 0,3694762 0,8151 | 18 |
| GOBP_NEGATIVE_REGULATION_OF_PROTEIN_BINDING | 0,8137931 | 1 0,0293519 0,2964342 0,815031 | 83 |
| GOBP_POSITIVE_REGULATION_OF_PEPTIDE_SECRETION | 0,7834928 | 1 0,0331626 0,3065989 0,8145886 | 64 |
| GOBP_GLAND_MORPHOGENESIS | 0,8215909 | 1 0,0282924 0,2925762 0,8141873 | 99 |
| GOBP_REGULATION_OF_LAMELLIPODIUM_ORGANIZATION | 0,7647799 | 1 0,036685 0,3217697 0,8135155 | 44 |
| GOBP_NEGATIVE_REGULATION_OF_PROTEIN_MATURATION | 0,7263017 | 1 0,0418239 0,3656867 0,8132647 | 19 |
| GOBP_LYMPH_VESSEL_DEVELOPMENT | 0,7347995 | 1 0,0398407 0,3488115 0,8128588 | 25 |
| GOBP_TOLL_LIKE_RECEPTOR_SIGNALING_PATHWAY | 0,8176606 | 1 0,0289981 0,2938677 0,8119095 | 94 |
| GOBP_CELLULAR_MODIFIED_AMINO_ACID_METABOLIC_PROCESS | 0,8450704 | 1 0,0242859 0,279249 0,8107496 | 152 |
| GOBP_REGULATION_OF_PLASMA_LIPOPROTEIN_PARTICLE_LEVELS | 0,7806122 | 1 0,0363783 0,3216333 0,8106587 | 45 |
| GOBP_POSITIVE_REGULATION_OF_AMYLOID_BETA_FORMATION | 0,7252011 | 1 0,0420772 0,3771239 0,8097031 | 16 |
| GOBP_ACTIVATION_OF_GTPASE_ACTIVITY | 0,8211009 | 1 0,0287861 0,292985 0,8094706 | 94 |
| GOBP_RESPONSE_TO_OXYGEN_LEVELS | 0,89375 | 1 0,0186259 0,2688451 0,8093927 | 264 |
| GOBP_POST_TRANSLATIONAL_PROTEIN_MODIFICATION | 0,7831021 | 1 0,0356933 0,3283467 0,8090496 | 37 |
| GOBP_NEGATIVE_REGULATION_OF_APOPTOTIC_SIGNALING_PATHWAY | 0,8729989 | 1 0,0215798 0,2743155 0,8086089 | 192 |
| GOBP_CELLULAR_PIGMENTATION | 0,7889447 | 1 0,0351652 0,3173751 0,8082692 | 48 |
| GOBP_GLYCEROLIPID_METABOLIC_PROCESS | 0,9039256 | 1 0,017313 0,266385 0,8082346 | 326 |
| GOBP_PHOSPHOLIPID_METABOLIC_PROCESS | 0,913811 | 1 0,0169177 0,2665405 0,8079505 | 335 |
| GOBP_NEGATIVE_REGULATION_OF_INTRACELLULAR_SIGNAL_TRANSDUCTION | 0,9154786 | 1 0,0153676 0,2618906 0,8078339 | 443 |
| GOBP_NEGATIVE_REGULATION_OF_DNA_TEMPLATED_TRANSCRIPTION_ELONGATION | 0,7398922 | 1 0,0414044 0,3661605 0,8077853 | 18 |
| GOBP_REGULATION_OF_CHONDROCYTE_DIFFERENTIATION | 0,7750953 | 1 0,0365315 0,3211714 0,8076686 | 43 |
| GOBP_LONG_CHAIN_FATTY_ACID_TRANSPORT | 0,7749684 | 1 0,0363018 0,3225546 0,8076657 | 42 |
| GOBP_REGULATION_OF_T_CELL_PROLIFERATION | 0,8453838 | 1 0,0256927 0,2855492 0,8075548 | 114 |
| GOBP_DEFENSE_RESPONSE_TO_BACTERIUM | 0,8582503 | 1 0,0246382 0,2831448 0,8075363 | 124 |
| GOBP_MYOTUBE_CELL_DEVELOPMENT | 0,7726692 | 1 0,0369161 0,3382416 0,8055382 | 30 |
| GOBP_RNA_DECAPPING | 0,7439353 | 1 0,0411544 0,3650767 0,8053941 | 18 |
| GOBP_MRNA_CATABOLIC_PROCESS | 0,8925532 | 1 0,020048 0,2716753 0,805069 | 217 |
| GOBP_NEGATIVE_REGULATION_OF_OSTEOCLAST_DIFFERENTIATION | 0,7565104 | 1 0,0387962 0,349512 0,8048071 | 23 |
| GOBP_MATURATION_OF_5_8S_RRNA | 0,7881463 | 1 0,0353911 0,3290698 0,8040354 | 35 |
| GOBP_NEUTRAL_AMINO_ACID_TRANSPORT | 0,7924528 | 1 0,0350151 0,323514 0,8031174 | 40 |
| GOBP_MYOTUBE_DIFFERENTIATION | 0,8331429 | 1 0,0278699 0,290152 0,8015344 | 89 |
| GOBP_NEGATIVE_REGULATION_OF_CELL_PROJECTION_ORGANIZATION | 0,8674569 | 1 0,0225129 0,2735948 0,8012031 | 170 |
| GOBP_MICROVILLUS_ASSEMBLY | 0,7413333 | 1 0,0408229 0,3542892 0,801104 | 21 |
| GOBP_NEGATIVE_REGULATION_OF_INTRINSIC_APOPTOTIC_SIGNALING_PATHWAY_IN_RESPONSE_TO_DNA_DAMAGE | 0,7695313 | 1 0,0380056 0,3398146 0,801046 | 28 |
| GOBP_NEGATIVE_REGULATION_OF_MYELOID_LEUKOCYTE_DIFFERENTIATION | 0,7818411 | 1 0,0357691 0,3260238 0,801023 | 36 |
| GOBP_PHOSPHATIDYLINOSITOL_METABOLIC_PROCESS | 0,8495093 | 1 0,0243564 0,2770311 0,8009685 | 145 |
| GOBP_NEGATIVE_REGULATION_OF_B_CELL_ACTIVATION | 0,7536618 | 1 0,0400032 0,3538004 0,8008218 | 22 |
| GOBP_EPITHELIAL_TO_MESENCHYMAL_TRANSITION | 0,8493909 | 1 0,025201 0,2799334 0,8006304 | 133 |
| GOBP_REGULATION_OF_AXONOGENESIS | 0,8507625 | 1 0,0242154 0,2765084 0,7998387 | 147 |
| GOBP_NEGATIVE_REGULATION_OF_ANOIKIS | 0,7574526 | 1 0,0405756 0,3791823 0,7998028 | 15 |
| GOBP_NEGATIVE_REGULATION_OF_CYSTEINE_TYPE_ENDOPEPTIDASE_ACTIVITY | 0,8067227 | 1 0,0319257 0,3023724 0,7997046 | 63 |
| GOBP_TONGUE_DEVELOPMENT | 0,7302013 | 1 0,0418239 0,3669364 0,7995805 | 17 |
| GOBP_NCRNA_CATABOLIC_PROCESS | 0,7987421 | 1 0,034641 0,3220285 0,7994297 | 40 |
| GOBP_RESPONSE_TO_RETINOIC_ACID | 0,8455378 | 1 0,027167 0,2893865 0,7987173 | 93 |
| GOBP_PITUITARY_GLAND_DEVELOPMENT | 0,7496774 | 1 0,0387962 0,3401158 0,7983669 | 26 |
| GOBP_T_CELL_PROLIFERATION | 0,8577681 | 1 0,0240036 0,2772641 0,7982825 | 142 |
| GOBP_NITRIC_OXIDE_SYNTHASE_BIOSYNTHETIC_PROCESS | 0,7412869 | 1 0,0410713 0,3716535 0,7979579 | 16 |
| GOBP_PROTEIN_LOCALIZATION_TO_PLASMA_MEMBRANE | 0,9049112 | 1 0,0180135 0,2654205 0,7979085 | 257 |
| GOBP_PEPTIDYL_LYSINE_TRIMETHYLATION | 0,8056581 | 1 0,0331626 0,3096976 0,7977202 | 51 |
| GOBP_BONE_MINERALIZATION | 0,8506271 | 1 0,0266755 0,2867581 0,7971295 | 98 |
| GOBP_PHOSPHOLIPID_BIOSYNTHETIC_PROCESS | 0,907001 | 1 0,017859 0,2668019 0,7970062 | 237 |
| GOBP_POSITIVE_REGULATION_OF_IMMUNE_EFFECTOR_PROCESS | 0,8648649 | 1 0,0228694 0,2741371 0,7968615 | 155 |
| GOBP_REACTIVE_OXYGEN_SPECIES_BIOSYNTHETIC_PROCESS | 0,8020177 | 1 0,0345664 0,3194206 0,7967346 | 41 |
| GOBP_NUCLEAR_TRANSCRIBED_MRNA_CATABOLIC_PROCESS_EXONUCLEOLYTIC | 0,7503338 | 1 0,0403295 0,358215 0,7966479 | 19 |
| GOBP_RESPONSE_TO_THYROID_HORMONE | 0,7734375 | 1 0,0377705 0,3459451 0,7965939 | 23 |
| GOBP_POSITIVE_REGULATION_OF_REACTIVE_OXYGEN_SPECIES_METABOLIC_PROCESS | 0,802439 | 1 0,0329431 0,3058393 0,7962997 | 54 |
| GOBP_DEVELOPMENTAL_GROWTH_INVOLVED_IN_MORPHOGENESIS | 0,892895 | 1 0,0198261 0,2685357 0,7961941 | 214 |
| GOBP_CELL_SUBSTRATE_ADHESION | 0,9237996 | 1 0,0165176 0,2637989 0,795645 | 300 |
| GOBP_STEROID_HORMONE_SECRETION | 0,7615176 | 1 0,0403295 0,3771124 0,7954368 | 15 |
| GOBP_CLATHRIN_DEPENDENT_ENDOCYTOSIS | 0,7984694 | 1 0,0353157 0,3155475 0,7953197 | 45 |
| GOBP_EYE_PHOTORECEPTOR_CELL_DEVELOPMENT | 0,7580854 | 1 0,0383996 0,3412308 0,795193 | 25 |
| GOBP_METANEPHRIC_NEPHRON_MORPHOGENESIS | 0,7493333 | 1 0,0403295 0,3516335 0,7950989 | 21 |
| GOBP_REGULATION_OF_TRANSCRIPTION_ELONGATION_FROM_RNA_POLYMERASE_II_PROMOTER | 0,80375 | 1 0,0340464 0,3206121 0,7950327 | 39 |
| GOBP_RESPONSE_TO_TUMOR_NECROSIS_FACTOR | 0,8811563 | 1 0,0212182 0,2703961 0,7950124 | 177 |
| GOBP_OSTEOBLAST_DEVELOPMENT | 0,7628726 | 1 0,0402478 0,376675 0,7945143 | 15 |
| GOBP_STEROID_BIOSYNTHETIC_PROCESS | 0,8669623 | 1 0,0241448 0,2778924 0,794496 | 128 |
| GOBP_HIPPO_SIGNALING | 0,8110831 | 1 0,0339724 0,3213668 0,7940058 | 38 |
| GOBP_RENAL_TUBULE_DEVELOPMENT | 0,8399533 | 1 0,0285744 0,2914963 0,7938646 | 78 |
| GOBP_POSITIVE_REGULATION_OF_PROTEIN_LOCALIZATION_TO_NUCLEUS | 0,8329412 | 1 0,0293519 0,2939471 0,7937882 | 75 |
| GOBP_REGULATION_OF_APOPTOTIC_SIGNALING_PATHWAY | 0,9305699 | 1 0,0154514 0,2621164 0,792948 | 313 |
| GOBP_POSITIVE_REGULATION_OF_I_KAPPAB_KINASE_NF_KAPPAB_SIGNALING | 0,8825431 | 1 0,0215076 0,2707657 0,7920566 | 169 |
| GOBP_REGULATION_OF_HIPPO_SIGNALING | 0,753004 | 1 0,0401661 0,3561175 0,7919833 | 19 |
| GOBP_SNO_S_RNA_METABOLIC_PROCESS | 0,7493298 | 1 0,0405756 0,3687069 0,7916314 | 16 |
| GOBP_PRODUCTION_OF_MOLECULAR_MEDIATOR_INVOLVED_IN_INFLAMMATORY_RESPONSE | 0,8228155 | 1 0,0314929 0,3010387 0,7910596 | 58 |
| GOBP_REGULATION_OF_TOLL_LIKE_RECEPTOR_SIGNALING_PATHWAY | 0,8097561 | 1 0,0325058 0,3038101 0,7910165 | 54 |
| GOBP_NEGATIVE_REGULATION_OF_MITOCHONDRION_ORGANIZATION | 0,7966963 | 1 0,0352404 0,3145104 0,7909178 | 43 |
| GOBP_NEGATIVE_REGULATION_OF_PROTEIN_CONTAINING_COMPLEX_ASSEMBLY | 0,8803987 | 1 0,0232248 0,2772786 0,7908056 | 124 |
| GOBP_POSITIVE_REGULATION_OF_DOUBLE_STRAND_BREAK_REPAIR_VIA_NONHOMOLOGOUS_END_JOINING | 0,7682927 | 1 0,0399219 0,3748384 0,7906402 | 15 |
| GOBP_ANDROGEN_METABOLIC_PROCESS | 0,7560647 | 1 0,0404114 0,3583847 0,7906311 | 18 |
| GOBP_REGULATION_OF_DNA_TEMPLATED_TRANSCRIPTION_INITIATION | 0,8208589 | 1 0,0321428 0,3048154 0,7898455 | 53 |
| GOBP_REGULATION_OF_HEART_CONTRACTION | 0,8808234 | 1 0,0219398 0,2715332 0,7883482 | 152 |
| GOBP_METANEPHROS_MORPHOGENESIS | 0,7773438 | 1 0,0375363 0,3361626 0,7881684 | 27 |
| GOBP_CELLULAR_RESPONSE_TO_STEROL_DEPLETION | 0,7710027 | 1 0,0397596 0,3735769 0,7879794 | 15 |
| GOBP_ACTIN_NUCLEATION | 0,8096447 | 1 0,0344175 0,3104575 0,7878277 | 47 |
| GOBP_CELLULAR_RESPONSE_TO_XENOBIOTIC_STIMULUS | 0,8682432 | 1 0,0249197 0,2804212 0,7876757 | 106 |
| GOBP_NEGATIVE_REGULATION_OF_LIPID_STORAGE | 0,747651 | 1 0,0407404 0,3612205 0,7871251 | 17 |
| GOBP_NEGATIVE_REGULATION_OF_INNATE_IMMUNE_RESPONSE | 0,8280488 | 1 0,0314209 0,3018142 0,7870377 | 55 |
| GOBP_ARP2_3_COMPLEX_MEDIATED_ACTIN_NUCLEATION | 0,8058008 | 1 0,0343431 0,3194076 0,7870237 | 37 |
| GOBP_EPHRIN_RECEPTOR_SIGNALING_PATHWAY | 0,8134518 | 1 0,0341946 0,311067 0,7859197 | 46 |
| GOBP_LYMPHOCYTE_COSTIMULATION | 0,7774903 | 1 0,0372254 0,3370387 0,7854239 | 25 |
| GOBP_REGULATION_OF_BONE_RESORPTION | 0,7838542 | 1 0,0371479 0,3331728 0,7853893 | 28 |
| GOBP_POSITIVE_REGULATION_OF_ALPHA_BETA_T_CELL_ACTIVATION | 0,8283951 | 1 0,031998 0,3049244 0,7838242 | 50 |
| GOBP_CATECHOLAMINE_SECRETION | 0,8221942 | 1 0,0333827 0,3142 0,7837128 | 41 |
| GOBP_POSITIVE_REGULATION_OF_LYMPHOCYTE_MEDIATED_IMMUNITY | 0,8435294 | 1 0,0287155 0,2902154 0,7837109 | 75 |
| GOBP_HISTONE_H4_ACETYLATION | 0,8416076 | 1 0,0290688 0,2912988 0,7822989 | 71 |
| GOBP_REGULATION_OF_LAMELLIPODIUM_ASSEMBLY | 0,8130489 | 1 0,033677 0,3214273 0,7819505 | 33 |
| GOBP_POSITIVE_REGULATION_OF_PROTEIN_KINASE_B_SIGNALING | 0,8568156 | 1 0,0265351 0,2835856 0,7817721 | 84 |
| GOBP_NEGATIVE_REGULATION_OF_PROTEIN_CONTAINING_COMPLEX_DISASSEMBLY | 0,8439716 | 1 0,0289274 0,2897866 0,7801116 | 73 |
| GOBP_GLUTAMATE_METABOLIC_PROCESS | 0,7873711 | 1 0,0364549 0,328636 0,780065 | 29 |
| GOBP_MAINTENANCE_OF_PROTEIN_LOCATION_IN_CELL | 0,8349398 | 1 0,0304181 0,2953804 0,7792118 | 60 |
| GOBP_NEUTROPHIL_MIGRATION | 0,8337321 | 1 0,030133 0,2927559 0,7789749 | 66 |
| GOBP_POSITIVE_REGULATION_OF_LIPID_BIOSYNTHETIC_PROCESS | 0,8361446 | 1 0,0303467 0,295241 0,7788442 | 60 |
| GOBP_REGULATION_OF_PROTEIN_CONTAINING_COMPLEX_DISASSEMBLY | 0,8887653 | 1 0,0229406 0,2748433 0,77884 | 116 |
| GOBP_NEGATIVE_REGULATION_OF_HORMONE_SECRETION | 0,8274112 | 1 0,0333827 0,308167 0,7785927 | 46 |
| GOBP_ENDOCYTOSIS | 0,9542218 | 1 0,0118236 0,2518634 0,7783067 | 466 |
| GOBP_HISTONE_MRNA_METABOLIC_PROCESS | 0,7914508 | 1 0,0364549 0,3349846 0,7781098 | 24 |
| GOBP_POSITIVE_REGULATION_OF_ERBB_SIGNALING_PATHWAY | 0,7899485 | 1 0,0363018 0,3277693 0,7780077 | 29 |
| GOBP_REGULATION_OF_TRANSPORTER_ACTIVITY | 0,9312896 | 1 0,0168381 0,2613287 0,7777124 | 229 |
| GOBP_HEMATOPOIETIC_STEM_CELL_PROLIFERATION | 0,7914508 | 1 0,0364549 0,3346483 0,7773287 | 24 |
| GOBP_CARBOHYDRATE_TRANSMEMBRANE_TRANSPORT | 0,8662857 | 1 0,0258332 0,2806979 0,777047 | 91 |
| GOBP_CELL_JUNCTION_DISASSEMBLY | 0,7818428 | 1 0,0391155 0,3682891 0,7768261 | 15 |
| GOBP_POSITIVE_REGULATION_OF_ADAPTIVE_IMMUNE_RESPONSE | 0,8586207 | 1 0,0266053 0,2825308 0,7768045 | 83 |
| GOBP_BRANCHED_CHAIN_AMINO_ACID_METABOLIC_PROCESS | 0,7929688 | 1 0,0366082 0,3372743 0,7766279 | 23 |
| GOBP_RETINOL_METABOLIC_PROCESS | 0,819568 | 1 0,0338984 0,3244671 0,7760428 | 31 |
| GOBP_POSITIVE_REGULATION_OF_RESPONSE_TO_EXTERNAL_STIMULUS | 0,934375 | 1 0,0155349 0,2573637 0,7760073 | 291 |
| GOBP_POSITIVE_REGULATION_OF_TRANSCRIPTION_BY_RNA_POLYMERASE_III | 0,7749326 | 1 0,0392759 0,3515739 0,7756057 | 18 |
| GOBP_CALCIUM_ION_TRANSPORT | 0,9462254 | 1 0,013987 0,2560127 0,7747766 | 314 |
| GOBP_POSITIVE_REGULATION_OF_IMMUNOGLOBULIN_PRODUCTION | 0,8196721 | 1 0,0335297 0,3143737 0,77462 | 37 |
| GOBP_CELLULAR_MODIFIED_AMINO_ACID_BIOSYNTHETIC_PROCESS | 0,8322825 | 1 0,0327971 0,3103005 0,7739862 | 41 |
| GOBP_REGULATION_OF_PROTON_TRANSPORT | 0,7712766 | 1 0,0388759 0,3444423 0,7739336 | 20 |
| GOBP_PROTEIN_LIPID_COMPLEX_SUBUNIT_ORGANIZATION | 0,7955729 | 1 0,0364549 0,328279 0,7738532 | 28 |
| GOBP_RIBONUCLEOSIDE_METABOLIC_PROCESS | 0,7896774 | 1 0,0363783 0,3296476 0,7737945 | 26 |
| GOBP_REGULATION_OF_LYMPHOCYTE_APOPTOTIC_PROCESS | 0,8348045 | 1 0,0326513 0,3101021 0,7734912 | 41 |
| GOBP_REGULATION_OF_REACTIVE_OXYGEN_SPECIES_BIOSYNTHETIC_PROCESS | 0,8230866 | 1 0,0330894 0,3178749 0,7733084 | 33 |
| GOBP_ACTIVATION_OF_PROTEIN_KINASE_B_ACTIVITY | 0,7942708 | 1 0,0365315 0,3295924 0,7727637 | 27 |
| GOBP_POSITIVE_REGULATION_OF_EPITHELIAL_TO_MESENCHYMAL_TRANSITION | 0,8324873 | 1 0,0330894 0,3058102 0,7726381 | 46 |
| GOBP_POSITIVE_REGULATION_OF_SMAD_PROTEIN_SIGNAL_TRANSDUCTION | 0,7762803 | 1 0,0391956 0,350225 0,77263 | 18 |
| GOBP_NEGATIVE_REGULATION_OF_PROTEIN_POLYMERIZATION | 0,8510638 | 1 0,0285039 0,2876979 0,7726285 | 71 |
| GOBP_REGULATION_OF_STRESS_ACTIVATED_PROTEIN_KINASE_SIGNALING_CASCADE | 0,9014085 | 1 0,0205626 0,2645748 0,7723842 | 166 |
| GOBP_LIPOPOLYSACCHARIDE_MEDIATED_SIGNALING_PATHWAY | 0,8367347 | 1 0,0330894 0,3064271 0,7723323 | 45 |
| GOBP_ESTABLISHMENT_OF_PIGMENT_GRANULE_LOCALIZATION | 0,7726064 | 1 0,0387962 0,3436293 0,772107 | 20 |
| GOBP_NEGATIVE_REGULATION_OF_NUCLEOTIDE_METABOLIC_PROCESS | 0,7774799 | 1 0,0388759 0,3595487 0,7719683 | 16 |
| GOBP_MYOBLAST_DIFFERENTIATION | 0,8458781 | 1 0,0293519 0,2904148 0,7716875 | 65 |
| GOBP_REGULATION_OF_INTRACELLULAR_PROTEIN_TRANSPORT | 0,9220918 | 1 0,0181675 0,2610957 0,7712043 | 206 |
| GOBP_REGULATION_OF_VIRAL_GENOME_REPLICATION | 0,8515901 | 1 0,0282924 0,2857774 0,7711911 | 74 |
| GOBP_CELLULAR_RESPONSE_TO_DEXAMETHASONE_STIMULUS | 0,8007813 | 1 0,0361492 0,3348188 0,7709737 | 23 |
| GOBP_REGULATED_EXOCYTOSIS | 0,9200426 | 1 0,0182442 0,2616752 0,7709471 | 191 |
| GOBP_REGULATION_OF_SUPRAMOLECULAR_FIBER_ORGANIZATION | 0,9492228 | 1 0,0138978 0,2545165 0,7709127 | 330 |
| GOBP_POSITIVE_REGULATION_OF_EPITHELIAL_CELL_MIGRATION | 0,8976898 | 1 0,021724 0,2690122 0,770795 | 137 |
| GOBP_NEGATIVE_REGULATION_OF_CELLULAR_PROTEIN_CATABOLIC_PROCESS | 0,8539192 | 1 0,0285744 0,2877197 0,7705257 | 70 |
| GOBP_DENDRITE_MORPHOGENESIS | 0,8903654 | 1 0,0225843 0,2693842 0,7704592 | 133 |
| GOBP_REGULATION_OF_SYSTEM_PROCESS | 0,9559877 | 1 0,0122193 0,2503004 0,7702771 | 415 |
| GOBP_REGULATION_OF_SENSORY_PERCEPTION | 0,7739362 | 1 0,0387167 0,3426747 0,7699621 | 20 |
| GOBP_ORGANIC_ACID_TRANSMEMBRANE_TRANSPORT | 0,8946785 | 1 0,0223699 0,2699066 0,7697652 | 126 |
| GOBP_LYMPHANGIOGENESIS | 0,7828418 | 1 0,0385579 0,3584445 0,7695975 | 16 |
| GOBP_NEGATIVE_REGULATION_OF_EPITHELIAL_TO_MESENCHYMAL_TRANSITION | 0,7994792 | 1 0,0362255 0,3264562 0,7695563 | 28 |
| GOBP_JNK_CASCADE | 0,9002169 | 1 0,0207088 0,2656495 0,7695279 | 149 |
| GOBP_SKIN_EPIDERMIS_DEVELOPMENT | 0,8664303 | 1 0,0275886 0,2854835 0,7685275 | 73 |
| GOBP_HETEROPHILIC_CELL_CELL_ADHESION_VIA_PLASMA_MEMBRANE_CELL_ADHESION_MOLECULES | 0,8409091 | 1 0,0323604 0,3157857 0,7682562 | 34 |
| GOBP_SLEEP | 0,7841823 | 1 0,0384787 0,3577402 0,7680855 | 16 |
| GOBP_NEUTRAL_LIPID_METABOLIC_PROCESS | 0,8922018 | 1 0,0244269 0,2779452 0,7679181 | 94 |
| GOBP_VACUOLE_ORGANIZATION | 0,9146211 | 1 0,0187018 0,2608406 0,767802 | 179 |
| GOBP_REGULATION_OF_B_CELL_ACTIVATION | 0,8832952 | 1 0,0248494 0,2779202 0,7676119 | 92 |
| GOBP_CELLULAR_RESPONSE_TO_CORTICOSTEROID_STIMULUS | 0,8388325 | 1 0,0327242 0,30243 0,7674569 | 47 |
| GOBP_REGULATION_OF_POSTSYNAPTIC_NEUROTRANSMITTER_RECEPTOR_ACTIVITY | 0,7868633 | 1 0,0383206 0,3573027 0,767146 | 16 |
| GOBP_INTRACELLULAR_RECEPTOR_SIGNALING_PATHWAY | 0,9244681 | 1 0,0177815 0,2588673 0,7671145 | 217 |
| GOBP_IRON_ION_HOMEOSTASIS | 0,8586698 | 1 0,0282924 0,2862324 0,7665426 | 70 |
| GOBP_AUDITORY_RECEPTOR_CELL_DEVELOPMENT | 0,7771812 | 1 0,0389556 0,3517126 0,7664068 | 17 |
| GOBP_REGULATION_OF_MITOCHONDRIAL_FISSION | 0,8002577 | 1 0,0356933 0,3227318 0,7660506 | 29 |
| GOBP_INTRINSIC_APOPTOTIC_SIGNALING_PATHWAY | 0,9447341 | 1 0,0147735 0,254568 0,7660025 | 267 |
| GOBP_REGULATION_OF_OXIDATIVE_STRESS_INDUCED_CELL_DEATH | 0,8592233 | 1 0,0293519 0,2914589 0,7658862 | 58 |
| GOBP_REGULATION_OF_RECEPTOR_INTERNALIZATION | 0,8585486 | 1 0,0300618 0,2973027 0,7657934 | 51 |
| GOBP_NEGATIVE_REGULATION_OF_ENDOCYTOSIS | 0,8451777 | 1 0,0323604 0,3030957 0,7657798 | 46 |
| GOBP_CARDIAC_ATRIUM_MORPHOGENESIS | 0,7779255 | 1 0,0384787 0,3404925 0,7650589 | 20 |
| GOBP_LEUKOCYTE_PROLIFERATION | 0,9309245 | 1 0,0172343 0,2585871 0,7647907 | 211 |
| GOBP_REGULATION_OF_VASCULAR_ENDOTHELIAL_GROWTH_FACTOR_RECEPTOR_SIGNALING_PATHWAY | 0,8108808 | 1 0,0353157 0,3291165 0,7644792 | 24 |
| GOBP_PATTERN_RECOGNITION_RECEPTOR_SIGNALING_PATHWAY | 0,9066959 | 1 0,0210003 0,2663636 0,7644348 | 138 |
| GOBP_CELLULAR_RESPONSE_TO_HEAT | 0,854878 | 1 0,0298485 0,2924026 0,7643899 | 56 |
| GOBP_AXO_DENDRITIC_TRANSPORT | 0,8763127 | 1 0,0263245 0,2820264 0,7642942 | 76 |
| GOBP_T_CELL_SELECTION | 0,8199234 | 1 0,0341205 0,3205977 0,7635185 | 30 |
| GOBP_CELL_DEATH_IN_RESPONSE_TO_OXIDATIVE_STRESS | 0,8725377 | 1 0,0261842 0,2792456 0,7633348 | 80 |
| GOBP_NEGATIVE_REGULATION_OF_VASCULAR_ASSOCIATED_SMOOTH_MUSCLE_CELL_PROLIFERATION | 0,7779255 | 1 0,0384787 0,3396495 0,7631647 | 20 |
| GOBP_REGULATION_OF_OXIDOREDUCTASE_ACTIVITY | 0,8786464 | 1 0,0261842 0,2812605 0,7631525 | 77 |
| GOBP_RIBOSE_PHOSPHATE_METABOLIC_PROCESS | 0,9626168 | 1 0,0128902 0,2516338 0,7628017 | 336 |
| GOBP_REGULATION_OF_DENDRITE_MORPHOGENESIS | 0,8578313 | 1 0,0290688 0,2891127 0,7626775 | 60 |
| GOBP_NEGATIVE_REGULATION_OF_SMALL_MOLECULE_METABOLIC_PROCESS | 0,8770302 | 1 0,0259736 0,2793273 0,7626079 | 79 |
| GOBP_CELL_SUBSTRATE_JUNCTION_ORGANIZATION | 0,8916762 | 1 0,0241448 0,2748236 0,7620902 | 95 |
| GOBP_PH_REDUCTION | 0,8436318 | 1 0,0321428 0,3054547 0,7618992 | 41 |
| GOBP_REGULATION_OF_SODIUM_ION_TRANSMEMBRANE_TRANSPORT | 0,8634146 | 1 0,0293519 0,2921182 0,7617535 | 55 |
| GOBP_REGULATION_OF_MYOBLAST_DIFFERENTIATION | 0,845283 | 1 0,0319257 0,3066756 0,7613164 | 40 |
| GOBP_NEGATIVE_REGULATION_OF_PEPTIDYL_THREONINE_PHOSPHORYLATION | 0,800813 | 1 0,0380056 0,3608977 0,7612355 | 15 |
| GOBP_REGULATION_OF_CARDIAC_MUSCLE_CONTRACTION_BY_CALCIUM_ION_SIGNALING | 0,792 | 1 0,0377705 0,3363946 0,7606416 | 21 |
| GOBP_REGULATION_OF_ACTIVIN_RECEPTOR_SIGNALING_PATHWAY | 0,8111979 | 1 0,035542 0,3302627 0,7604826 | 23 |
| GOBP_POSITIVE_REGULATION_OF_TYROSINE_PHOSPHORYLATION_OF_STAT_PROTEIN | 0,84625 | 1 0,0315649 0,3066303 0,7603616 | 39 |
| GOBP_GLYCEROLIPID_BIOSYNTHETIC_PROCESS | 0,935518 | 1 0,0165176 0,2556672 0,760336 | 224 |
| GOBP_LEUKOCYTE_MIGRATION | 0,94375 | 1 0,0147735 0,2541078 0,760215 | 239 |
| GOBP_SYNAPTIC_VESICLE_MEMBRANE_ORGANIZATION | 0,8173575 | 1 0,0349401 0,3272757 0,7602034 | 24 |
| GOBP_POSITIVE_REGULATION_OF_BLOOD_VESSEL_ENDOTHELIAL_CELL_MIGRATION | 0,8683887 | 1 0,0294936 0,2950045 0,7598737 | 51 |
| GOBP_PIGMENT_GRANULE_LOCALIZATION | 0,8002663 | 1 0,0372254 0,3355118 0,7594258 | 22 |
| GOBP_PHOSPHATIDYLINOSITOL_3_PHOSPHATE_BIOSYNTHETIC_PROCESS | 0,7957276 | 1 0,0376143 0,3411265 0,7586442 | 19 |
| GOBP_REGULATION_OF_DOUBLE_STRAND_BREAK_REPAIR_VIA_NONHOMOLOGOUS_END_JOINING | 0,8054124 | 1 0,0353911 0,3195558 0,7585118 | 29 |
| GOBP_REGULATION_OF_ANION_TRANSMEMBRANE_TRANSPORT | 0,7933333 | 1 0,0376923 0,3352823 0,7581265 | 21 |
| GOBP_POSITIVE_REGULATION_OF_CELL_MATRIX_ADHESION | 0,836478 | 1 0,0324331 0,2997056 0,7577318 | 44 |
| GOBP_NEGATIVE_REGULATION_OF_INTERLEUKIN_2_PRODUCTION | 0,7983979 | 1 0,0374584 0,3401945 0,7565715 | 19 |
| GOBP_REGULATION_OF_ERYTHROCYTE_DIFFERENTIATION | 0,8425693 | 1 0,0321428 0,3058185 0,7555903 | 38 |
| GOBP_DETECTION_OF_MECHANICAL_STIMULUS_INVOLVED_IN_SENSORY_PERCEPTION | 0,8103226 | 1 0,0351652 0,3215831 0,7548644 | 26 |
| GOBP_BONE_MATURATION | 0,8203125 | 1 0,0350151 0,3276089 0,7543719 | 23 |
| GOBP_NEGATIVE_REGULATION_OF_SYNAPTIC_TRANSMISSION | 0,8652226 | 1 0,0285744 0,2853465 0,754364 | 61 |
| GOBP_REGULATION_OF_CELL_SIZE | 0,9274892 | 1 0,0187018 0,2580618 0,7542055 | 167 |
| GOBP_POSITIVE_REGULATION_OF_LEUKOCYTE_ADHESION_TO_VASCULAR_ENDOTHELIAL_CELL | 0,7973154 | 1 0,0377705 0,3456414 0,7531773 | 17 |
| GOBP_REPRODUCTIVE_BEHAVIOR | 0,7951482 | 1 0,0380842 0,3413325 0,7530123 | 18 |
| GOBP_PHOSPHATIDYLINOSITOL_BIOSYNTHETIC_PROCESS | 0,9124169 | 1 0,0212182 0,2641183 0,7527509 | 123 |
| GOBP_REGULATION_OF_LYMPHOCYTE_MEDIATED_IMMUNITY | 0,9089888 | 1 0,0222268 0,2674213 0,7527409 | 108 |
| GOBP_POSITIVE_REGULATION_OF_GLUCOSE_METABOLIC_PROCESS | 0,8522727 | 1 0,0317091 0,3091618 0,7521414 | 34 |
| GOBP_RRNA_METABOLIC_PROCESS | 0,9593326 | 1 0,0135373 0,2499967 0,7521375 | 258 |
| GOBP_MODIFICATION_OF_SYNAPTIC_STRUCTURE | 0,7898936 | 1 0,0377705 0,3346962 0,7520351 | 20 |
| GOBP_NUCLEAR_TRANSCRIBED_MRNA_CATABOLIC_PROCESS | 0,9114064 | 1 0,0212182 0,2643831 0,7519601 | 118 |
| GOBP_POSITIVE_REGULATION_OF_PROTEIN_LOCALIZATION_TO_MEMBRANE | 0,9028571 | 1 0,0235793 0,2716324 0,7519513 | 91 |
| GOBP_T_CELL_DIFFERENTIATION | 0,9423693 | 1 0,0166782 0,2549918 0,7516476 | 192 |
| GOBP_REGULATION_OF_STRIATED_MUSCLE_CELL_DIFFERENTIATION | 0,8696172 | 1 0,0280107 0,2822933 0,7511356 | 66 |
| GOBP_LYSOSOMAL_PROTEIN_CATABOLIC_PROCESS | 0,8077437 | 1 0,0369161 0,3376738 0,7509656 | 19 |
| GOBP_PROTEIN_LOCALIZATION_TO_CELL_SURFACE | 0,8688327 | 1 0,0283628 0,2840602 0,7509635 | 61 |
| GOBP_LEUKOCYTE_APOPTOTIC_PROCESS | 0,8943089 | 1 0,0249901 0,274083 0,7507151 | 82 |
| GOBP_REGULATION_OF_REGULATED_SECRETORY_PATHWAY | 0,9148936 | 1 0,0216519 0,2663061 0,7507011 | 111 |

| GOBP_RESPONSE_TO_MINERALOCORTICOID | 0,8163001 | 1 0,0349401 0,3221032 0,7506188 | 25 |
| --- | --- | --- | --- |
| GOBP_PROTEIN_KINASE_B_SIGNALING | 0,932973 | 1 0,0182442 0,2581806 0,7504792 | 155 |
| GOBP_GLYCEROPHOSPHOLIPID_METABOLIC_PROCESS | 0,9593326 | 1 0,0135373 0,2493961 0,7504536 | 268 |
| GOBP_ANIMAL_ORGAN_MATURATION | 0,8212435 | 1 0,0347157 0,3229827 0,7502316 | 24 |
| GOBP_REGULATION_OF_CATION_CHANNEL_ACTIVITY | 0,9228225 | 1 0,0201955 0,2620236 0,7501768 | 136 |
| GOBP_NEGATIVE_REGULATION_OF_VIRAL_PROCESS | 0,8877069 | 1 0,0263245 0,2792837 0,7500317 | 71 |
| GOBP_SYNAPTIC_VESICLE_CYTOSKELETAL_TRANSPORT | 0,8018868 | 1 0,0376923 0,339944 0,749949 | 18 |
| GOBP_REGULATION_OF_EPITHELIAL_TO_MESENCHYMAL_TRANSITION | 0,8966318 | 1 0,0248494 0,2736821 0,7496172 | 82 |
| GOBP_REGULATION_OF_GLUCOSE_METABOLIC_PROCESS | 0,9121622 | 1 0,0221551 0,2673217 0,7494982 | 103 |
| GOBP_POSITIVE_REGULATION_OF_FOCAL_ADHESION_ASSEMBLY | 0,8225389 | 1 0,034641 0,3225032 0,7491178 | 24 |
| GOBP_NADH_METABOLIC_PROCESS | 0,8216146 | 1 0,0349401 0,3175703 0,7486095 | 28 |
| GOBP_GLYCEROL_ETHER_METABOLIC_PROCESS | 0,7952128 | 1 0,0374584 0,333096 0,7484394 | 20 |
| GOBP_ACTIVATION_OF_CYSTEINE_TYPE_ENDOPEPTIDASE_ACTIVITY | 0,8018868 | 1 0,0376923 0,338977 0,7478157 | 18 |
| GOBP_REGULATION_OF_RENAL_SYSTEM_PROCESS | 0,8268229 | 1 0,034641 0,3244984 0,7472093 | 23 |
| GOBP_LIPOPROTEIN_METABOLIC_PROCESS | 0,9201774 | 1 0,0207088 0,2619567 0,7465901 | 123 |
| GOBP_KETONE_BIOSYNTHETIC_PROCESS | 0,8625473 | 1 0,0310618 0,3031526 0,7448298 | 36 |
| GOBP_ESTABLISHMENT_OR_MAINTENANCE_OF_BIPOLAR_CELL_POLARITY | 0,8806533 | 1 0,0298485 0,2923623 0,7445681 | 48 |
| GOBP_POSITIVE_REGULATION_OF_PEPTIDYL_SERINE_PHOSPHORYLATION | 0,8949825 | 1 0,025201 0,2746966 0,7444303 | 76 |
| GOBP_REGULATION_OF_ERBB_SIGNALING_PATHWAY | 0,8735084 | 1 0,0276589 0,2784113 0,7442131 | 67 |
| GOBP_VASCULAR_ENDOTHELIAL_GROWTH_FACTOR_RECEPTOR_SIGNALING_PATHWAY | 0,8769036 | 1 0,0305608 0,2944585 0,7439578 | 46 |
| GOBP_REGULATION_OF_CARTILAGE_DEVELOPMENT | 0,8762136 | 1 0,0283628 0,2830931 0,7439028 | 58 |
| GOBP_REGULATION_OF_RAS_PROTEIN_SIGNAL_TRANSDUCTION | 0,936078 | 1 0,0181675 0,2542094 0,7429074 | 164 |
| GOBP_REGULATION_OF_PH | 0,9053738 | 1 0,0246382 0,2727058 0,7426903 | 78 |
| GOBP_REGULATION_OF_CYSTEINE_TYPE_ENDOPEPTIDASE_ACTIVITY | 0,9454545 | 1 0,016598 0,2524567 0,7421869 | 186 |
| GOBP_T_HELPER_17_TYPE_IMMUNE_RESPONSE | 0,8270968 | 1 0,0341946 0,3159286 0,7415914 | 26 |
| GOBP_POSITIVE_REGULATION_OF_CELL_GROWTH | 0,9350935 | 1 0,0192297 0,2587232 0,7413142 | 137 |
| GOBP_REGULATION_OF_LIPID_BIOSYNTHETIC_PROCESS | 0,9238411 | 1 0,0201955 0,2587108 0,7412642 | 132 |
| GOBP_BROWN_FAT_CELL_DIFFERENTIATION | 0,8602015 | 1 0,0311335 0,2996039 0,7402359 | 38 |
| GOBP_POSITIVE_REGULATION_OF_CYSTEINE_TYPE_ENDOPEPTIDASE_ACTIVITY | 0,9255556 | 1 0,0204894 0,2600469 0,7402271 | 121 |
| GOBP_POSITIVE_REGULATION_OF_OXIDOREDUCTASE_ACTIVITY | 0,8807107 | 1 0,0303467 0,2916454 0,7400894 | 47 |
| GOBP_MICROVILLUS_ORGANIZATION | 0,8283871 | 1 0,0341205 0,3151086 0,7396665 | 26 |
| GOBP_NEGATIVE_REGULATION_OF_SIGNAL_TRANSDUCTION_BY_P53_CLASS_MEDIATOR | 0,8475222 | 1 0,0322878 0,3090932 0,7392724 | 31 |
| GOBP_POSITIVE_REGULATION_OF_ACTIN_CYTOSKELETON_REORGANIZATION | 0,8174497 | 1 0,0366082 0,3389988 0,7387025 | 17 |
| GOBP_POSTTRANSCRIPTIONAL_REGULATION_OF_GENE_EXPRESSION | 0,9828109 | 1 0,0078803 0,2381153 0,7386556 | 496 |
| GOBP_PIGMENT_GRANULE_ORGANIZATION | 0,8411458 | 1 0,0338245 0,3148096 0,738104 | 27 |
| GOBP_TRNA_METHYLATION | 0,8675914 | 1 0,0307753 0,2956589 0,7374655 | 41 |
| GOBP_METANEPHRIC_EPITHELIUM_DEVELOPMENT | 0,8151596 | 1 0,0363018 0,3281125 0,7372419 | 20 |
| GOBP_BONE_GROWTH | 0,84375 | 1 0,033677 0,3144382 0,7372332 | 27 |
| GOBP_REGULATION_OF_LONG_TERM_SYNAPTIC_POTENTIATION | 0,8663304 | 1 0,0308468 0,2991324 0,7370653 | 37 |
| GOBP_NEGATIVE_REGULATION_OF_PROTEIN_LOCALIZATION_TO_CELL_PERIPHERY | 0,8380829 | 1 0,0337507 0,3170335 0,7364126 | 24 |
| GOBP_POSITIVE_REGULATION_OF_PROTEIN_POLYMERIZATION | 0,9061414 | 1 0,0241448 0,2692155 0,7359171 | 80 |
| GOBP_ACTIN_POLYMERIZATION_OR_DEPOLYMERIZATION | 0,9454545 | 1 0,016598 0,2503677 0,7355608 | 174 |
| GOBP_POSITIVE_REGULATION_OF_CELL_KILLING | 0,8500635 | 1 0,0321428 0,3074217 0,7352747 | 31 |
| GOBP_PYRIMIDINE_NUCLEOTIDE_CATABOLIC_PROCESS | 0,8241611 | 1 0,0362255 0,3373723 0,7351584 | 17 |
| GOBP_REGULATION_OF_ENDOCYTOSIS | 0,9496788 | 1 0,0163561 0,2499424 0,7348749 | 177 |
| GOBP_REGULATION_OF_HORMONE_BIOSYNTHETIC_PROCESS | 0,8218085 | 1 0,0359209 0,3270434 0,7348399 | 20 |
| GOBP_REGULATION_OF_PROTEIN_POLYMERIZATION | 0,9507495 | 1 0,016275 0,249863 0,7346413 | 177 |
| GOBP_POSITIVE_REGULATION_OF_PROTEIN_CONTAINING_COMPLEX_DISASSEMBLY | 0,8619824 | 1 0,0308468 0,3014101 0,7332538 | 33 |
| GOBP_PYRIMIDINE_NUCLEOSIDE_TRIPHOSPHATE_METABOLIC_PROCESS | 0,8424479 | 1 0,0337507 0,3183231 0,7329897 | 23 |
| GOBP_PHOSPHATIDYLGLYCEROL_METABOLIC_PROCESS | 0,8476563 | 1 0,0334562 0,3108823 0,7328438 | 28 |
| GOBP_EMBRYONIC_BRAIN_DEVELOPMENT | 0,8281879 | 1 0,0359969 0,3359093 0,7319702 | 17 |
| GOBP_RESPONSE_TO_SALT_STRESS | 0,8221024 | 1 0,0365315 0,3314177 0,7311392 | 18 |
| GOBP_ERYTHROCYTE_DEVELOPMENT | 0,84 | 1 0,0334562 0,311337 0,7308134 | 26 |
| GOBP_CALCIUM_MEDIATED_SIGNALING_USING_INTRACELLULAR_CALCIUM_SOURCE | 0,8364611 | 1 0,0354665 0,339346 0,7285922 | 16 |
| GOBP_PROTEASOME_MEDIATED_UBIQUITIN_DEPENDENT_PROTEIN_CATABOLIC_PROCESS | 0,9805128 | 1 0,0098982 0,2373594 0,728365 | 391 |
| GOBP_SYNAPTIC_TRANSMISSION_CHOLINERGIC | 0,8306667 | 1 0,035542 0,3220703 0,728252 | 21 |
| GOBP_IMMUNE_RESPONSE_REGULATING_SIGNALING_PATHWAY | 0,9770115 | 1 0,0121213 0,2419908 0,7282257 | 288 |
| GOBP_DEOXYRIBOSE_PHOSPHATE_CATABOLIC_PROCESS | 0,8408797 | 1 0,0335297 0,3124167 0,7280456 | 25 |
| GOBP_POSITIVE_REGULATION_OF_MORPHOGENESIS_OF_AN_EPITHELIUM | 0,8518519 | 1 0,0322878 0,3054678 0,7274858 | 30 |
| GOBP_PURINE_DEOXYRIBONUCLEOTIDE_METABOLIC_PROCESS | 0,8401084 | 1 0,0357691 0,3445068 0,7266624 | 15 |
| GOBP_RUFFLE_ORGANIZATION | 0,9007538 | 1 0,0287155 0,2849093 0,7255874 | 48 |
| GOBP_GLUTAMATE_SECRETION | 0,8397864 | 1 0,0350901 0,3262596 0,7255812 | 19 |
| GOBP_DNA_TEMPLATED_TRANSCRIPTION_TERMINATION | 0,8333333 | 1 0,0353911 0,3206142 0,7249595 | 21 |
| GOBP_NEGATIVE_REGULATION_OF_CELLULAR_CATABOLIC_PROCESS | 0,9653361 | 1 0,013628 0,2430984 0,7246138 | 231 |
| GOBP_VESICLE_DOCKING | 0,9 | 1 0,0272372 0,2777671 0,7243304 | 55 |
| GOBP_POSITIVE_REGULATION_OF_TOR_SIGNALING | 0,8855346 | 1 0,0296355 0,291776 0,7243285 | 40 |
| GOBP_REGULATION_OF_ENDOCYTIC_RECYCLING | 0,833557 | 1 0,0356933 0,3322743 0,7240494 | 17 |
| GOBP_REGULATION_OF_BEHAVIOR | 0,9030612 | 1 0,0293519 0,2871152 0,7236578 | 45 |
| GOBP_REGULATION_OF_MULTICELLULAR_ORGANISM_GROWTH | 0,900978 | 1 0,0273075 0,2794614 0,7222354 | 52 |
| GOBP_RESPIRATORY_BURST | 0,8468708 | 1 0,0345664 0,3189686 0,7219806 | 22 |
| GOBP_XENOBIOTIC_METABOLIC_PROCESS | 0,9 | 1 0,0272372 0,2761604 0,7219301 | 56 |
| GOBP_NUCLEOSIDE_TRIPHOSPHATE_METABOLIC_PROCESS | 0,9350057 | 1 0,0214354 0,2603275 0,7218923 | 95 |
| GOBP_POSITIVE_REGULATION_OF_CELLULAR_AMIDE_METABOLIC_PROCESS | 0,9541985 | 1 0,017313 0,2495448 0,7214983 | 145 |
| GOBP_RESPONSE_TO_TEMPERATURE_STIMULUS | 0,9540984 | 1 0,0174698 0,2496752 0,7213978 | 146 |
| GOBP_POSITIVE_REGULATION_OF_MEMBRANE_PERMEABILITY | 0,8867925 | 1 0,0295645 0,290594 0,7213941 | 40 |
| GOBP_ICOSANOID_BIOSYNTHETIC_PROCESS | 0,8770389 | 1 0,0299907 0,2962776 0,7207678 | 33 |
| GOBP_TAIL_ANCHORED_MEMBRANE_PROTEIN_INSERTION_INTO_ER_MEMBRANE | 0,8362416 | 1 0,035542 0,3304341 0,7200394 | 17 |
| GOBP_ORGANOPHOSPHATE_ESTER_TRANSPORT | 0,9457965 | 1 0,0188533 0,2528921 0,7192732 | 119 |
| GOBP_CELLULAR_LIPID_CATABOLIC_PROCESS | 0,9625668 | 1 0,0152836 0,2442358 0,7185461 | 180 |
| GOBP_RESPONSE_TO_MISFOLDED_PROTEIN | 0,8510363 | 1 0,0330163 0,3093117 0,7184761 | 24 |
| GOBP_GLUCOSE_METABOLIC_PROCESS | 0,9567568 | 1 0,0165176 0,2457828 0,7183187 | 168 |
| GOBP_CELLULAR_ALDEHYDE_METABOLIC_PROCESS | 0,8968447 | 1 0,027167 0,2734468 0,7179836 | 57 |
| GOBP_REGULATION_OF_ISOTYPE_SWITCHING | 0,8795483 | 1 0,0298485 0,2949578 0,717557 | 33 |
| GOBP_NEGATIVE_REGULATION_OF_ACTIN_FILAMENT_POLYMERIZATION | 0,9004854 | 1 0,0269563 0,2726792 0,715968 | 57 |
| GOBP_NEGATIVE_REGULATION_OF_NF_KAPPAB_TRANSCRIPTION_FACTOR_ACTIVITY | 0,9148936 | 1 0,0247086 0,2658229 0,7156009 | 73 |
| GOBP_POSITIVE_REGULATION_OF_CARTILAGE_DEVELOPMENT | 0,8671875 | 1 0,0323604 0,3033513 0,715091 | 28 |
| GOBP_NEGATIVE_REGULATION_OF_AUTOPHAGY | 0,9311552 | 1 0,0230117 0,2634676 0,7148745 | 77 |
| GOBP_INTRACELLULAR_LIPID_TRANSPORT | 0,8918239 | 1 0,0292811 0,2826581 0,7146313 | 44 |
| GOBP_INFLAMMATORY_CELL_APOPTOTIC_PROCESS | 0,8389262 | 1 0,0353911 0,3279401 0,7146049 | 17 |
| GOBP_VESICLE_TARGETING_TO_FROM_OR_WITHIN_GOLGI | 0,8535286 | 1 0,0341946 0,3154047 0,7139137 | 22 |
| GOBP_SYNAPTIC_TRANSMISSION_DOPAMINERGIC | 0,8453333 | 1 0,0347157 0,3157124 0,7138758 | 21 |
| GOBP_DNA_DAMAGE_RESPONSE_SIGNAL_TRANSDUCTION_BY_P53_CLASS_MEDIATOR_RESULTING_IN_CELL_CYCLE_ARREST | 0,8442953 | 1 0,0350901 0,3275811 0,7138225 | 17 |
| GOBP_CELLULAR_RESPIRATION | 0,9701493 | 1 0,0144273 0,2415297 0,7137833 | 201 |
| GOBP_NEGATIVE_REGULATION_OF_TRANSFORMING_GROWTH_FACTOR_BETA_RECEPTOR_SIGNALING_PATHWAY | 0,9087635 | 1 0,0259034 0,2698305 0,7136389 | 63 |
| GOBP_POSITIVE_REGULATION_OF_T_CELL_MEDIATED_IMMUNITY | 0,8865069 | 1 0,0297065 0,2857504 0,7127505 | 41 |
| GOBP_POSITIVE_REGULATION_OF_CELL_SUBSTRATE_JUNCTION_ORGANIZATION | 0,8634021 | 1 0,0320704 0,3002565 0,7127021 | 29 |
| GOBP_ANION_HOMEOSTASIS | 0,8671775 | 1 0,0314209 0,2988955 0,7118337 | 30 |
| GOBP_TOLL_LIKE_RECEPTOR_3_SIGNALING_PATHWAY | 0,8409704 | 1 0,0354665 0,3224475 0,71135 | 18 |
| GOBP_AXONAL_TRANSPORT | 0,9126794 | 1 0,0254821 0,2675628 0,7108755 | 64 |
| GOBP_PYRIMIDINE_RIBONUCLEOSIDE_TRIPHOSPHATE_METABOLIC_PROCESS | 0,8483221 | 1 0,0348652 0,3261365 0,7106747 | 17 |
| GOBP_POSITIVE_REGULATION_OF_PEPTIDYL_LYSINE_ACETYLATION | 0,8820577 | 1 0,0297065 0,2921264 0,7106689 | 33 |
| GOBP_REGULATION_OF_INTRACELLULAR_TRANSPORT | 0,9854167 | 1 0,0110006 0,2353538 0,7105856 | 309 |
| GOBP_NCRNA_METABOLIC_PROCESS | 0,9959555 | 1 0,0058146 0,2290952 0,7105563 | 497 |
| GOBP_NCRNA_TRANSCRIPTION | 0,9085366 | 1 0,0267457 0,2722693 0,7099939 | 55 |
| GOBP_CELL_MIGRATION_INVOLVED_IN_SPROUTING_ANGIOGENESIS | 0,9074074 | 1 0,027448 0,2761127 0,7097624 | 50 |
| GOBP_RIBONUCLEOSIDE_MONOPHOSPHATE_METABOLIC_PROCESS | 0,9165644 | 1 0,0266053 0,2734997 0,7086994 | 53 |
| GOBP_LYMPH_VESSEL_MORPHOGENESIS | 0,849734 | 1 0,0343431 0,3149038 0,7075632 | 20 |
| GOBP_VACUOLAR_LOCALIZATION | 0,9151732 | 1 0,0252713 0,2662744 0,7075417 | 65 |
| GOBP_NEUROTRANSMITTER_RECEPTOR_INTERNALIZATION | 0,8485255 | 1 0,0347904 0,3287443 0,7058299 | 16 |
| GOBP_REGULATION_OF_CELL_MORPHOGENESIS | 0,983368 | 1 0,0110006 0,2342508 0,7058133 | 274 |
| GOBP_SNRNA_METABOLIC_PROCESS | 0,9156479 | 1 0,0264649 0,2727367 0,7048561 | 52 |
| GOBP_REGULATION_OF_MYELOID_LEUKOCYTE_MEDIATED_IMMUNITY | 0,8978562 | 1 0,0290688 0,2860365 0,7047969 | 37 |
| GOBP_FOLIC_ACID_METABOLIC_PROCESS | 0,8631436 | 1 0,0344919 0,3339719 0,7044413 | 15 |
| GOBP_GROWTH_PLATE_CARTILAGE_DEVELOPMENT | 0,8604027 | 1 0,0341946 0,3231004 0,7040588 | 17 |
| GOBP_PHOSPHOLIPID_TRANSPORT | 0,9358534 | 1 0,0216519 0,2553348 0,7038918 | 84 |
| GOBP_POSITIVE_REGULATION_OF_RESPONSE_TO_CYTOKINE_STIMULUS | 0,9180929 | 1 0,0263245 0,2721877 0,7034373 | 52 |
| GOBP_MOLTING_CYCLE | 0,9396051 | 1 0,0222268 0,2559737 0,7011137 | 82 |
| GOBP_REGULATION_OF_LEUKOCYTE_MEDIATED_IMMUNITY | 0,9707476 | 1 0,0156181 0,2417295 0,7005883 | 150 |
| GOBP_REGULATION_OF_PROTEIN_EXIT_FROM_ENDOPLASMIC_RETICULUM | 0,8743523 | 1 0,0317091 0,3015846 0,7005276 | 24 |
| GOBP_POSITIVE_REGULATION_OF_HISTONE_ACETYLATION | 0,8841146 | 1 0,0314209 0,2986882 0,7003058 | 27 |
| GOBP_POSITIVE_REGULATION_OF_JUN_KINASE_ACTIVITY | 0,8977273 | 1 0,0291396 0,2876277 0,6997524 | 34 |
| GOBP_REGULATION_OF_PROTEIN_TARGETING | 0,9304245 | 1 0,0236501 0,2599781 0,6994897 | 72 |
| GOBP_POLYSACCHARIDE_CATABOLIC_PROCESS | 0,8672087 | 1 0,0342688 0,3315679 0,6993707 | 15 |
| GOBP_PROTON_TRANSPORTING_TWO_SECTOR_ATPASE_COMPLEX_ASSEMBLY | 0,8672087 | 1 0,0342688 0,3315415 0,6993148 | 15 |
| GOBP_POSITIVE_REGULATION_OF_SUPRAMOLECULAR_FIBER_ORGANIZATION | 0,9707476 | 1 0,0156181 0,2407076 0,6988515 | 152 |
| GOBP_REGULATION_OF_PROTEIN_LOCALIZATION_TO_MEMBRANE | 0,9751082 | 1 0,0151993 0,2403212 0,6980453 | 153 |
| GOBP_NCRNA_PROCESSING | 0,9907121 | 1 0,0094283 0,2283219 0,6971417 | 373 |
| GOBP_BONE_CELL_DEVELOPMENT | 0,8867188 | 1 0,0312771 0,2972643 0,6969673 | 27 |
| GOBP_HISTONE_H3_K4_TRIMETHYLATION | 0,8611482 | 1 0,0338984 0,3133501 0,6968712 | 19 |
| GOBP_OXIDATIVE_PHOSPHORYLATION | 0,9587973 | 1 0,0183973 0,246185 0,6953647 | 113 |
| GOBP_STRESS_GRANULE_ASSEMBLY | 0,8880208 | 1 0,0312053 0,2963969 0,6949334 | 27 |
| GOBP_VACUOLAR_ACIDIFICATION | 0,8867188 | 1 0,0312771 0,3015551 0,6943789 | 23 |
| GOBP_SIGNALING_RECEPTOR_LIGAND_PRECURSOR_PROCESSING | 0,8919271 | 1 0,0309901 0,2940975 0,693277 | 28 |
| GOBP_FATTY_ACID_CATABOLIC_PROCESS | 0,956422 | 1 0,020416 0,2508719 0,6931189 | 94 |
| GOBP_REGULATION_OF_INTRINSIC_APOPTOTIC_SIGNALING_PATHWAY_IN_RESPONSE_TO_DNA_DAMAGE | 0,9079445 | 1 0,0285039 0,2817897 0,6923421 | 36 |
| GOBP_RAB_PROTEIN_SIGNAL_TRANSDUCTION | 0,8656915 | 1 0,0334562 0,3081156 0,6923106 | 20 |
| GOBP_POSITIVE_REGULATION_OF_ESTABLISHMENT_OF_PROTEIN_LOCALIZATION_TO_MITOCHONDRION | 0,8926768 | 1 0,0294228 0,2866892 0,6919631 | 32 |
| GOBP_RESPONSE_TO_INCREASED_OXYGEN_LEVELS | 0,8761651 | 1 0,0329431 0,3055324 0,6915678 | 22 |
| GOBP_NEGATIVE_REGULATION_OF_PROTEIN_TYROSINE_KINASE_ACTIVITY | 0,8878866 | 1 0,0307037 0,2910771 0,6909136 | 29 |
| GOBP_REGULATION_OF_MONOOXYGENASE_ACTIVITY | 0,9169811 | 1 0,0278699 0,2780814 0,690332 | 40 |
| GOBP_REGULATION_OF_CALCIUM_ION_TRANSMEMBRANE_TRANSPORTER_ACTIVITY | 0,9397163 | 1 0,0232248 0,2563985 0,69023 | 73 |
| GOBP_MYELIN_ASSEMBLY | 0,8696809 | 1 0,0332359 0,3067262 0,6891887 | 20 |
| GOBP_DEFENSE_RESPONSE_TO_SYMBIONT | 0,9841438 | 1 0,01251 0,2313618 0,6880532 | 224 |
| GOBP_POSITIVE_REGULATION_OF_DENDRITIC_SPINE_DEVELOPMENT | 0,9058971 | 1 0,0283628 0,2827269 0,6878024 | 33 |
| GOBP_NONRIBOSOMAL_PEPTIDE_BIOSYNTHETIC_PROCESS | 0,8659517 | 1 0,0338245 0,3200767 0,6872201 | 16 |
| GOBP_RNA_5_END_PROCESSING | 0,872 | 1 0,0332359 0,3036529 0,6866073 | 21 |
| GOBP_ACTIN_FILAMENT_POLYMERIZATION | 0,9770492 | 1 0,0157838 0,2374755 0,6861486 | 146 |
| GOBP_REGULATION_OF_MONOCYTE_CHEMOTAXIS | 0,8780488 | 1 0,033677 0,3252125 0,6859653 | 15 |
| GOBP_NEGATIVE_REGULATION_OF_BLOOD_VESSEL_ENDOTHELIAL_CELL_MIGRATION | 0,8984375 | 1 0,0306323 0,2908278 0,6855692 | 28 |
| GOBP_RESPONSE_TO_INTERFERON_GAMMA | 0,9610092 | 1 0,0201218 0,2479607 0,6850757 | 94 |
| GOBP_REGULATION_OF_ADAPTIVE_IMMUNE_RESPONSE | 0,9679204 | 1 0,017313 0,2391436 0,6848904 | 130 |
| GOBP_MITOTIC_G1_S_TRANSITION_CHECKPOINT_SIGNALING | 0,8943299 | 1 0,0303467 0,2884525 0,6846836 | 29 |
| GOBP_EPIDERMIS_MORPHOGENESIS | 0,8937824 | 1 0,0306323 0,2947513 0,684655 | 24 |
| GOBP_REGULATION_OF_LEUKOCYTE_MEDIATED_CYTOTOXICITY | 0,9362245 | 1 0,0275183 0,2715382 0,6843969 | 45 |
| GOBP_NUCLEAR_MEMBRANE_ORGANIZATION | 0,920354 | 1 0,0279403 0,2729639 0,6834922 | 42 |
| GOBP_IMP_METABOLIC_PROCESS | 0,8739946 | 1 0,0333827 0,3179384 0,682629 | 16 |
| GOBP_VESICLE_MEDIATED_TRANSPORT_BETWEEN_ENDOSOMAL_COMPARTMENTS | 0,9224905 | 1 0,0280811 0,2710407 0,6816019 | 43 |
| GOBP_REGULATION_OF_PROTEASOMAL_UBIQUITIN_DEPENDENT_PROTEIN_CATABOLIC_PROCESS | 0,9722838 | 1 0,0171554 0,2381996 0,6810143 | 128 |
| GOBP_SYNAPTIC_VESICLE_RECYCLING | 0,9550827 | 1 0,0222984 0,2534147 0,6805592 | 71 |
| GOBP_CELLULAR_CARBOHYDRATE_CATABOLIC_PROCESS | 0,9015152 | 1 0,0289274 0,2816138 0,6797132 | 32 |
| GOBP_POSITIVE_REGULATION_OF_PROTEIN_CONTAINING_COMPLEX_ASSEMBLY | 0,9771987 | 1 0,0152836 0,2323869 0,678381 | 163 |
| GOBP_TETRAPYRROLE_BIOSYNTHETIC_PROCESS | 0,905972 | 1 0,0289981 0,2834116 0,6778485 | 31 |
| GOBP_NEGATIVE_REGULATION_OF_CARBOHYDRATE_METABOLIC_PROCESS | 0,927044 | 1 0,0273075 0,2679573 0,677464 | 44 |
| GOBP_SULFUR_COMPOUND_CATABOLIC_PROCESS | 0,8989637 | 1 0,0303467 0,2912387 0,6764957 | 24 |
| GOBP_REGULATION_OF_DENDRITIC_SPINE_DEVELOPMENT | 0,9446494 | 1 0,0251307 0,2622924 0,6756138 | 51 |
| GOBP_T_CELL_MEDIATED_IMMUNITY | 0,9540636 | 1 0,0221551 0,2502159 0,6752259 | 74 |
| GOBP_POSITIVE_REGULATION_OF_PROTEIN_LOCALIZATION_TO_CELL_SURFACE | 0,8834688 | 1 0,0333827 0,3200128 0,6749976 | 15 |
| GOBP_PRESYNAPTIC_ENDOCYTOSIS | 0,9421687 | 1 0,0241448 0,2556495 0,674402 | 60 |
| GOBP_REGULATION_OF_VIRAL_TRANSCRIPTION | 0,8894879 | 1 0,0327971 0,3056675 0,6743319 | 18 |
| GOBP_EPITHELIAL_STRUCTURE_MAINTENANCE | 0,8878505 | 1 0,0324331 0,3029633 0,6737715 | 19 |
| GOBP_CARBOHYDRATE_CATABOLIC_PROCESS | 0,9733925 | 1 0,0170763 0,2353779 0,6729468 | 128 |
| GOBP_REGULATION_OF_PROTEIN_LOCALIZATION_TO_PLASMA_MEMBRANE | 0,9587629 | 1 0,0201955 0,2427656 0,6718728 | 90 |
| GOBP_VIRAL_LIFE_CYCLE | 0,9916405 | 1 0,0106795 0,2229664 0,6708862 | 260 |
| GOBP_PURINE_NUCLEOTIDE_TRANSPORT | 0,9028497 | 1 0,030133 0,2888233 0,6708852 | 24 |
| GOBP_POSITIVE_REGULATION_OF_AUTOPHAGY | 0,9767184 | 1 0,0168381 0,2342566 0,6694442 | 127 |
| GOBP_PROTEASOMAL_PROTEIN_CATABOLIC_PROCESS | 0,9979633 | 1 0,0066784 0,2166824 0,6693513 | 464 |
| GOBP_POSITIVE_REGULATION_OF_ATP_METABOLIC_PROCESS | 0,9085052 | 1 0,0295645 0,2819896 0,6693431 | 29 |
| GOBP_MODULATION_BY_HOST_OF_SYMBIONT_PROCESS | 0,9447853 | 1 0,0249901 0,2574544 0,6671225 | 53 |
| GOBP_PINOCYTOSIS | 0,8961385 | 1 0,0318535 0,2945153 0,6666307 | 22 |
| GOBP_AEROBIC_RESPIRATION | 0,9859002 | 1 0,0145143 0,2288689 0,6666045 | 160 |
| GOBP_MRNA_METHYLATION | 0,886059 | 1 0,0327242 0,3100101 0,6656065 | 16 |
| GOBP_MEMBRANE_DOCKING | 0,9674797 | 1 0,0204894 0,242229 0,663467 | 82 |
| GOBP_SUBSTANTIA_NIGRA_DEVELOPMENT | 0,9407314 | 1 0,0266755 0,2659005 0,6632387 | 41 |
| GOBP_NUCLEOSIDE_DIPHOSPHATE_METABOLIC_PROCESS | 0,9800885 | 1 0,0164369 0,2330595 0,6628654 | 119 |
| GOBP_SNARE_COMPLEX_ASSEMBLY | 0,8929539 | 1 0,0328701 0,3140297 0,6623775 | 15 |
| GOBP_REGULATION_OF_T_CELL_MEDIATED_CYTOTOXICITY | 0,9119171 | 1 0,0296355 0,2849171 0,6618118 | 24 |
| GOBP_CIRCADIAN_SLEEP_WAKE_CYCLE | 0,8929539 | 1 0,0328701 0,3133341 0,6609104 | 15 |
| GOBP_REGULATION_OF_DEFENSE_RESPONSE_TO_VIRUS_BY_HOST | 0,9255051 | 1 0,0275886 0,2715041 0,6605262 | 34 |
| GOBP_PORPHYRIN_CONTAINING_COMPOUND_METABOLIC_PROCESS | 0,9432535 | 1 0,0265351 0,2646811 0,6601972 | 41 |
| GOBP_POSITIVE_REGULATION_OF_APOPTOTIC_SIGNALING_PATHWAY | 0,9822024 | 1 0,0166782 0,2328841 0,6599378 | 116 |
| GOBP_DENDRITIC_SPINE_DEVELOPMENT | 0,9703872 | 1 0,0190795 0,2383988 0,6593002 | 86 |
| GOBP_CYTOPLASMIC_MICROTUBULE_ORGANIZATION | 0,9496933 | 1 0,0247086 0,2540505 0,6583021 | 53 |
| GOBP_ANTIMICROBIAL_HUMORAL_RESPONSE | 0,9469027 | 1 0,0264649 0,2624759 0,6572307 | 42 |
| GOBP_AUTOPHAGOSOME_ORGANIZATION | 0,9876126 | 1 0,0171554 0,2334407 0,6557122 | 106 |
| GOBP_CERAMIDE_TRANSPORT | 0,9043127 | 1 0,031998 0,2968899 0,6549676 | 18 |
| GOBP_COLLATERAL_SPROUTING | 0,9153646 | 1 0,0297065 0,2842148 0,65445 | 23 |
| GOBP_REGULATION_OF_GOLGI_ORGANIZATION | 0,9056604 | 1 0,0319257 0,2961712 0,6533821 | 18 |
| GOBP_SULFUR_AMINO_ACID_METABOLIC_PROCESS | 0,9231771 | 1 0,0292811 0,2771242 0,6532657 | 28 |
| GOBP_NADPH_REGENERATION | 0,9033557 | 1 0,0318535 0,2985821 0,6506317 | 17 |
| GOBP_TRNA_5_END_PROCESSING | 0,898374 | 1 0,0325785 0,3079025 0,6494535 | 15 |
| GOBP_REGULATION_OF_AUTOPHAGY | 0,9979296 | 1 0,008938 0,2141516 0,6484218 | 317 |
| GOBP_RESPONSE_TO_VIRUS | 0,9947862 | 1 0,0101264 0,2147719 0,6477965 | 301 |
| GOBP_MONOSACCHARIDE_BIOSYNTHETIC_PROCESS | 0,9717314 | 1 0,021073 0,2400009 0,6476599 | 74 |
| GOBP_GLYCOPROTEIN_CATABOLIC_PROCESS | 0,9106667 | 1 0,0311335 0,2862965 0,6473618 | 21 |
| GOBP_REGULATION_OF_RECEPTOR_RECYCLING | 0,9106667 | 1 0,0311335 0,286067 0,6468429 | 21 |
| GOBP_REGULATION_OF_TRANSCRIPTION_OF_NUCLEOLAR_LARGE_RRNA_BY_RNA_POLYMERASE_I | 0,899729 | 1 0,0325058 0,3059832 0,6454052 | 15 |
| GOBP_POSITIVE_REGULATION_OF_ACTIVATED_T_CELL_PROLIFERATION | 0,9100671 | 1 0,0314929 0,2961792 0,6453956 | 17 |
| GOBP_LATE_ENDOSOME_TO_LYSOSOME_TRANSPORT | 0,9225634 | 1 0,0305608 0,2897983 0,6444934 | 19 |
| GOBP_PROTEIN_N_LINKED_GLYCOSYLATION | 0,9772999 | 1 0,0215798 0,2421461 0,6434283 | 65 |
| GOBP_TETRAHYDROFOLATE_METABOLIC_PROCESS | 0,9225634 | 1 0,0305608 0,2892601 0,6432965 | 19 |
| GOBP_REGULATION_OF_LYSOSOMAL_LUMEN_PH | 0,899729 | 1 0,0325058 0,3048619 0,6430401 | 15 |
| GOBP_REGULATION_OF_DEFENSE_RESPONSE_TO_VIRUS | 0,9700957 | 1 0,0220834 0,2414768 0,6415686 | 64 |
| GOBP_BLASTOCYST_FORMATION | 0,9495586 | 1 0,0261842 0,2603633 0,6415377 | 37 |
| GOBP_POSITIVE_REGULATION_OF_COAGULATION | 0,899729 | 1 0,0325058 0,3041175 0,64147 | 15 |
| GOBP_MATURATION_OF_LSU_RRNA | 0,9296875 | 1 0,0289274 0,2720963 0,6414134 | 28 |
| GOBP_EPITHELIAL_TO_MESENCHYMAL_TRANSITION_INVOLVED_IN_ENDOCARDIAL_CUSHION_FORMATION | 0,9252336 | 1 0,0304181 0,2882777 0,6411117 | 19 |
| GOBP_REGULATION_OF_FIBROBLAST_GROWTH_FACTOR_RECEPTOR_SIGNALING_PATHWAY | 0,9309896 | 1 0,0288568 0,2716763 0,6404234 | 28 |
| GOBP_LIPOPROTEIN_BIOSYNTHETIC_PROCESS | 0,9827982 | 1 0,0187018 0,2317435 0,6402701 | 94 |
| GOBP_REGULATION_OF_UBIQUITIN_DEPENDENT_PROTEIN_CATABOLIC_PROCESS | 0,9935135 | 1 0,013628 0,2202914 0,6400438 | 154 |
| GOBP_NEGATIVE_REGULATION_OF_ERBB_SIGNALING_PATHWAY | 0,9304124 | 1 0,0283628 0,2689872 0,6384799 | 29 |
| GOBP_PROTEIN_MODIFICATION_BY_SMALL_PROTEIN_REMOVAL | 0,9923996 | 1 0,0140757 0,2206149 0,6383528 | 148 |
| GOBP_LACTATION | 0,9495586 | 1 0,0261842 0,2590599 0,6383263 | 37 |
| GOBP_REGULATION_OF_NITRIC_OXIDE_SYNTHASE_ACTIVITY | 0,9304124 | 1 0,0283628 0,2683217 0,6369003 | 29 |
| GOBP_PROTON_TRANSMEMBRANE_TRANSPORT | 0,9911602 | 1 0,0155349 0,2239959 0,6368914 | 117 |
| GOBP_MESODERM_DEVELOPMENT | 0,9782609 | 1 0,0188533 0,2302281 0,6358869 | 92 |
| GOBP_LEUKOCYTE_DEGRANULATION | 0,9679012 | 1 0,0240036 0,2473717 0,6358819 | 50 |
| GOBP_REGULATION_OF_NUCLEASE_ACTIVITY | 0,9279039 | 1 0,0302754 0,2857244 0,6354334 | 19 |
| GOBP_NEUROTRANSMITTER_RECEPTOR_TRANSPORT_TO_PLASMA_MEMBRANE | 0,9061662 | 1 0,031637 0,2958618 0,6352295 | 16 |
| GOBP_NEURON_PROJECTION_ARBORIZATION | 0,9304124 | 1 0,0283628 0,2676078 0,6352056 | 29 |
| GOBP_MATURATION_OF_LSU_RRNA_FROM_TRICISTRONIC_RRNA_TRANSCRIPT_SSU_RRNA_5_8S_RRNA_LSU_RRNA | 0,903794 | 1 0,0322878 0,3004851 0,6338082 | 15 |
| GOBP_POSITIVE_REGULATION_OF_LYMPHOCYTE_DIFFERENTIATION | 0,9824971 | 1 0,0198261 0,2335179 0,6336111 | 77 |
| GOBP_PHOSPHATIDIC_ACID_METABOLIC_PROCESS | 0,9457071 | 1 0,0264649 0,2601056 0,6327955 | 34 |
| GOBP_POSITIVE_REGULATION_OF_CELLULAR_RESPONSE_TO_INSULIN_STIMULUS | 0,924 | 1 0,0304181 0,2794319 0,6318399 | 21 |
| GOBP_PROTEIN_LOCALIZATION_TO_POSTSYNAPSE | 0,9419192 | 1 0,0266755 0,2616373 0,631497 | 32 |
| GOBP_REGULATION_OF_MEMBRANE_PROTEIN_ECTODOMAIN_PROTEOLYSIS | 0,9253333 | 1 0,0303467 0,2790739 0,6310303 | 21 |
| GOBP_NUCLEOSOME_MOBILIZATION | 0,9253333 | 1 0,0303467 0,2788939 0,6306234 | 21 |
| GOBP_POSITIVE_REGULATION_OF_INTRACELLULAR_PROTEIN_TRANSPORT | 0,9934354 | 1 0,0146011 0,2187744 0,6298824 | 142 |
| GOBP_CELLULAR_DEFENSE_RESPONSE | 0,9254328 | 1 0,0302754 0,2781295 0,6295419 | 22 |
| GOBP_REGULATION_OF_MITOCHONDRION_ORGANIZATION | 0,9900332 | 1 0,0157838 0,2198446 0,628772 | 133 |
| GOBP_POSITIVE_REGULATION_OF_LEUKOCYTE_MEDIATED_IMMUNITY | 0,9862857 | 1 0,0182442 0,2274247 0,6282525 | 89 |
| GOBP_CYTOPLASMIC_TRANSLATION | 0,9956236 | 1 0,0144273 0,2181846 0,6277943 | 141 |
| GOBP_FATTY_ACID_BETA_OXIDATION | 0,9833729 | 1 0,0208547 0,2341449 0,62705 | 70 |
| GOBP_VESICLE_TARGETING | 0,965866 | 1 0,0254118 0,2503433 0,626851 | 42 |
| GOBP_ATP_METABOLIC_PROCESS | 0,9968288 | 1 0,0113146 0,21062 0,6265611 | 226 |
| GOBP_ENDOCRINE_HORMONE_SECRETION | 0,9494949 | 1 0,0262543 0,2563886 0,6237525 | 34 |
| GOBP_QUINONE_METABOLIC_PROCESS | 0,9623588 | 1 0,025201 0,255757 0,6221915 | 33 |
| GOBP_ENDOPLASMIC_RETICULUM_TUBULAR_NETWORK_ORGANIZATION | 0,9299191 | 1 0,0306323 0,2817663 0,6216035 | 18 |
| GOBP_RIBOSOME_BIOGENESIS | 0,9979101 | 1 0,0100128 0,2056725 0,6194507 | 296 |
| GOBP_NEGATIVE_REGULATION_OF_POSTTRANSCRIPTIONAL_GENE_SILENCING | 0,9261745 | 1 0,0306323 0,284186 0,6192617 | 17 |
| GOBP_MICROTUBULE_ANCHORING | 0,9469599 | 1 0,0276589 0,2656455 0,6190517 | 25 |
| GOBP_NEGATIVE_REGULATION_OF_JNK_CASCADE | 0,9636136 | 1 0,0251307 0,2544046 0,6189014 | 33 |
| GOBP_POSITIVE_REGULATION_OF_EXOCYTOSIS | 0,9869048 | 1 0,0207818 0,2307259 0,6179059 | 68 |
| GOBP_MONOSACCHARIDE_METABOLIC_PROCESS | 0,9968254 | 1 0,0114178 0,2076945 0,6171531 | 223 |
| GOBP_REGULATION_OF_AUTOPHAGY_OF_MITOCHONDRION | 0,9546028 | 1 0,0259034 0,2510165 0,6167341 | 36 |
| GOBP_POSITIVE_REGULATION_OF_NITRIC_OXIDE_SYNTHASE_ACTIVITY | 0,9262735 | 1 0,0305608 0,2871471 0,6165185 | 16 |
| GOBP_PROTEIN_QUALITY_CONTROL_FOR_MISFOLDED_OR_INCOMPLETELY_SYNTHESIZED_PROTEINS | 0,9505208 | 1 0,0277996 0,2628443 0,616266 | 27 |
| GOBP_PEROXISOME_ORGANIZATION | 0,9546028 | 1 0,0259034 0,2519691 0,6156509 | 35 |
| GOBP_ORGANIC_CATION_TRANSPORT | 0,9476373 | 1 0,0269563 0,2584046 0,6154027 | 30 |
| GOBP_REGULATION_OF_TRANSLATIONAL_ELONGATION | 0,9401596 | 1 0,0294228 0,2735041 0,6145413 | 20 |
| GOBP_ESTABLISHMENT_OF_PROTEIN_LOCALIZATION_TO_MEMBRANE | 0,9979101 | 1 0,0100128 0,2052014 0,6131609 | 238 |
| GOBP_REGULATION_OF_TRANSCRIPTION_REGULATORY_REGION_DNA_BINDING | 0,9685535 | 1 0,0249901 0,2423853 0,6128115 | 44 |
| GOBP_REGULATION_OF_ENDOTHELIAL_CELL_DIFFERENTIATION | 0,9608586 | 1 0,0256225 0,2517196 0,6123937 | 34 |
| GOBP_PROTEIN_LOCALIZATION_TO_LYSOSOME | 0,9685535 | 1 0,0249901 0,2421726 0,6122738 | 44 |
| GOBP_CARBOHYDRATE_PHOSPHORYLATION | 0,9443005 | 1 0,0278699 0,2632857 0,6115658 | 24 |
| GOBP_REGULATION_OF_STEROID_METABOLIC_PROCESS | 0,9894982 | 1 0,0193795 0,2253887 0,611554 | 77 |
| GOBP_REGULATION_OF_STEROID_BIOSYNTHETIC_PROCESS | 0,9756098 | 1 0,0228694 0,2343388 0,611083 | 55 |
| GOBP_NEGATIVE_REGULATION_OF_CATION_CHANNEL_ACTIVITY | 0,9608586 | 1 0,0256225 0,2530879 0,610862 | 32 |
| GOBP_NUCLEAR_TRANSCRIBED_MRNA_CATABOLIC_PROCESS_NONSENSE_MEDIATED_DECAY | 0,964691 | 1 0,0253416 0,2474139 0,6096304 | 37 |
| GOBP_PARAXIAL_MESODERM_DEVELOPMENT | 0,9241192 | 1 0,0312053 0,2887348 0,6090235 | 15 |
| GOBP_PROTEIN_PEPTIDYL_PROLYL_ISOMERIZATION | 0,9646465 | 1 0,0254118 0,2503147 0,6089757 | 34 |
| GOBP_NEGATIVE_REGULATION_OF_LIPID_LOCALIZATION | 0,9698871 | 1 0,024779 0,2502576 0,6088128 | 33 |
| GOBP_TRNA_METABOLIC_PROCESS | 0,9978564 | 1 0,01251 0,205882 0,6049233 | 181 |
| GOBP_PURINE_NUCLEOSIDE_MONOPHOSPHATE_METABOLIC_PROCESS | 0,9771283 | 1 0,0250604 0,2402155 0,6040841 | 43 |
| GOBP_NEGATIVE_REGULATION_OF_PROTEIN_LOCALIZATION_TO_MEMBRANE | 0,9600515 | 1 0,0267457 0,254085 0,6031074 | 29 |
| GOBP_REGULATION_OF_T_CELL_MEDIATED_IMMUNITY | 0,9780488 | 1 0,0227269 0,2306879 0,6030571 | 56 |
| GOBP_NUCLEOTIDE_TRANSMEMBRANE_TRANSPORT | 0,9506008 | 1 0,0290688 0,2709383 0,60255 | 19 |
| GOBP_MATERNAL_PLACENTA_DEVELOPMENT | 0,9600515 | 1 0,0267457 0,2538233 0,6024862 | 29 |
| GOBP_RIBOSOME_ASSEMBLY | 0,9780488 | 1 0,0227269 0,2303394 0,6021461 | 56 |
| GOBP_CARDIOLIPIN_METABOLIC_PROCESS | 0,9356568 | 1 0,0300618 0,2803576 0,6019412 | 16 |
| GOBP_RIBOSOMAL_LARGE_SUBUNIT_BIOGENESIS | 0,9928486 | 1 0,0204894 0,2250984 0,6017159 | 69 |
| GOBP_GOLGI_TO_PLASMA_MEMBRANE_TRANSPORT | 0,9866989 | 1 0,021724 0,2286948 0,6010423 | 59 |
| GOBP_RNA_METHYLATION | 0,9941928 | 1 0,0187776 0,2194119 0,6009706 | 82 |
| GOBP_POSITIVE_REGULATION_OF_MONOOXYGENASE_ACTIVITY | 0,9533679 | 1 0,0273777 0,2579421 0,5991536 | 24 |
| GOBP_RECEPTOR_RECYCLING | 0,9669632 | 1 0,0256225 0,2503588 0,5987947 | 31 |
| GOBP_POLYAMINE_METABOLIC_PROCESS | 0,9380054 | 1 0,0302042 0,270833 0,5974835 | 18 |
| GOBP_PURINE_NUCLEOSIDE_METABOLIC_PROCESS | 0,9546632 | 1 0,0273075 0,257015 0,5970003 | 24 |
| GOBP_NEGATIVE_REGULATION_OF_LEUKOCYTE_MEDIATED_IMMUNITY | 0,9659521 | 1 0,0252713 0,2427239 0,5963598 | 36 |
| GOBP_ESTABLISHMENT_OF_EPITHELIAL_CELL_POLARITY | 0,9720457 | 1 0,0253416 0,2484209 0,5941598 | 31 |
| GOBP_ENDOCYTIC_RECYCLING | 0,9929078 | 1 0,0199742 0,2211174 0,5938229 | 71 |
| GOBP_POSITIVE_REGULATION_OF_MULTICELLULAR_ORGANISM_GROWTH | 0,956 | 1 0,0287155 0,2626118 0,5938069 | 21 |
| GOBP_REGULATION_OF_CELL_MIGRATION_INVOLVED_IN_SPROUTING_ANGIOGENESIS | 0,9735183 | 1 0,0248494 0,2408889 0,5935527 | 37 |
| GOBP_VESICLE_MEDIATED_TRANSPORT_TO_THE_PLASMA_MEMBRANE | 0,9966887 | 1 0,0150299 0,2069028 0,5928227 | 132 |
| GOBP_RIBONUCLEOSIDE_TRIPHOSPHATE_BIOSYNTHETIC_PROCESS | 0,9927798 | 1 0,021073 0,224191 0,5926888 | 61 |
| GOBP_I_KAPPAB_PHOSPHORYLATION | 0,938255 | 1 0,0299907 0,2718942 0,5924768 | 17 |
| GOBP_REGULATION_OF_MITOCHONDRIAL_MEMBRANE_PERMEABILITY_INVOLVED_IN_APOPTOTIC_PROCESS | 0,9760404 | 1 0,0247086 0,2397243 0,5906831 | 37 |
| GOBP_PROTEIN_TARGETING_TO_MITOCHONDRION | 0,995439 | 1 0,0174698 0,2126818 0,5897649 | 96 |
| GOBP_SPHINGOMYELIN_METABOLIC_PROCESS | 0,9433962 | 1 0,0299196 0,2668754 0,5887527 | 18 |
| GOBP_REGULATION_OF_CELLULAR_PROTEIN_CATABOLIC_PROCESS | 0,9989551 | 1 0,0098982 0,1967394 0,5878759 | 238 |
| GOBP_RNA_MODIFICATION | 0,9956757 | 1 0,0134462 0,2022168 0,5878036 | 155 |
| GOBP_REGULATION_OF_CELL_MORPHOGENESIS_INVOLVED_IN_DIFFERENTIATION | 0,9954442 | 1 0,0173915 0,212437 0,5875018 | 86 |
| GOBP_RETROGRADE_TRANSPORT_ENDOSOME_TO_GOLGI | 0,9942792 | 1 0,0177815 0,2122315 0,5861807 | 92 |
| GOBP_RIBOSOMAL_SMALL_SUBUNIT_BIOGENESIS | 0,9964539 | 1 0,0197519 0,2182428 0,5861031 | 71 |
| GOBP_IRON_SULFUR_CLUSTER_ASSEMBLY | 0,9651613 | 1 0,0265351 0,249073 0,5846588 | 26 |
| GOBP_CYTOPLASMIC_SEQUESTERING_OF_PROTEIN | 0,9626667 | 1 0,0283628 0,2584932 0,5844942 | 21 |
| GOBP_RAC_PROTEIN_SIGNAL_TRANSDUCTION | 0,98 | 1 0,0240036 0,2352534 0,5833658 | 39 |
| GOBP_REGULATION_OF_PROTEASOMAL_PROTEIN_CATABOLIC_PROCESS | 0,9989293 | 1 0,0123167 0,1983674 0,5830606 | 176 |
| GOBP_REGULATION_OF_NUCLEOTIDE_METABOLIC_PROCESS | 0,9916667 | 1 0,0204894 0,2171516 0,5815526 | 68 |
| GOBP_GLUTAMINE_FAMILY_AMINO_ACID_BIOSYNTHETIC_PROCESS | 0,9449664 | 1 0,0296355 0,2663911 0,5804851 | 17 |
| GOBP_REGULATION_OF_RAC_PROTEIN_SIGNAL_TRANSDUCTION | 0,9587766 | 1 0,0284333 0,2582282 0,5802178 | 20 |
| GOBP_REGULATION_OF_FATTY_ACID_BETA_OXIDATION | 0,9463087 | 1 0,0295645 0,2659891 0,5796092 | 17 |
| GOBP_CELL_FATE_COMMITMENT_INVOLVED_IN_FORMATION_OF_PRIMARY_GERM_LAYER | 0,9596354 | 1 0,0273075 0,2513757 0,5788326 | 23 |
| GOBP_CELLULAR_RESPONSE_TO_VIRUS | 0,9928401 | 1 0,0205626 0,2156951 0,5765683 | 67 |
| GOBP_REGULATION_OF_TYPE_I_INTERFERON_MEDIATED_SIGNALING_PATHWAY | 0,9810845 | 1 0,0244269 0,2331775 0,5745517 | 37 |
| GOBP_ORGANELLE_TRANSPORT_ALONG_MICROTUBULE | 0,9965517 | 1 0,0179363 0,2087349 0,5739062 | 83 |
| GOBP_REGULATION_OF_NEURON_PROJECTION_ARBORIZATION | 0,9485095 | 1 0,0299196 0,2718473 0,573403 | 15 |
| GOBP_REGULATION_OF_HAIR_FOLLICLE_DEVELOPMENT | 0,9485095 | 1 0,0299196 0,2715944 0,5728695 | 15 |
| GOBP_REGULATION_OF_MEMBRANE_PERMEABILITY | 0,9927971 | 1 0,0209276 0,2161033 0,571543 | 63 |
| GOBP_T_CELL_CYTOKINE_PRODUCTION | 0,9702458 | 1 0,0263947 0,2451299 0,5712428 | 25 |
| GOBP_POSITIVE_REGULATION_OF_MITOCHONDRIAL_FISSION | 0,9654255 | 1 0,0280811 0,2536852 0,5700099 | 20 |
| GOBP_SYNAPTIC_VESICLE_LOCALIZATION | 0,9926199 | 1 0,0223699 0,2207871 0,5687043 | 51 |
| GOBP_REGULATION_OF_FIBROBLAST_MIGRATION | 0,9811794 | 1 0,0241448 0,2335129 0,5680772 | 33 |
| GOBP_TETRAPYRROLE_METABOLIC_PROCESS | 0,9926199 | 1 0,0223699 0,2199813 0,5666287 | 51 |
| GOBP_VIRAL_GENE_EXPRESSION | 0,9965675 | 1 0,017626 0,2051158 0,5665272 | 92 |
| GOBP_POSITIVE_REGULATION_OF_T_CELL_CYTOKINE_PRODUCTION | 0,965287 | 1 0,0282924 0,2543005 0,5655486 | 19 |
| GOBP_REGULATION_OF_CYTOPLASMIC_TRANSLATION | 0,9715395 | 1 0,0263245 0,2425675 0,5652714 | 25 |
| GOBP_POSITIVE_REGULATION_OF_DNA_TEMPLATED_TRANSCRIPTION_INITIATION | 0,9874214 | 1 0,023933 0,2229944 0,5637864 | 44 |
| GOBP_ORGANELLE_FUSION | 0,9988962 | 1 0,0148593 0,1968158 0,5636142 | 135 |
| GOBP_ENDOMEMBRANE_SYSTEM_ORGANIZATION | 1 | 1 0,0065141 0,1821312 0,5633235 | 475 |
| GOBP_CYTOSOLIC_TRANSPORT | 1 | 1 0,0130773 0,1926657 0,56308 | 168 |
| GOBP_ENTRAINMENT_OF_CIRCADIAN_CLOCK | 0,9822109 | 1 0,024779 0,2347051 0,561355 | 31 |
| GOBP_NEGATIVE_REGULATION_OF_PHAGOCYTOSIS | 0,9525745 | 1 0,0297065 0,2655857 0,5601954 | 15 |
| GOBP_TRANSPORT_OF_VIRUS | 0,958445 | 1 0,0288568 0,2602424 0,558753 | 16 |
| GOBP_PROTEIN_FOLDING | 1 | 1 0,0122193 0,1898845 0,558294 | 177 |
| GOBP_MRNA_MODIFICATION | 0,9754204 | 1 0,026114 0,2390013 0,5569607 | 25 |
| GOBP_NEGATIVE_REGULATION_OF_FIBROBLAST_GROWTH_FACTOR_RECEPTOR_SIGNALING_PATHWAY | 0,9651475 | 1 0,0285039 0,2582731 0,5545248 | 16 |
| GOBP_VESICLE_LOCALIZATION | 1 | 1 0,0121213 0,1884185 0,554236 | 178 |
| GOBP_NUCLEOTIDE_TRANSPORT | 0,9847522 | 1 0,0246382 0,2306918 0,5517563 | 31 |
| GOBP_REGULATION_OF_OXIDATIVE_PHOSPHORYLATION | 0,9706275 | 1 0,0280107 0,2472858 0,5499483 | 19 |
| GOBP_REGULATION_OF_CELLULAR_RESPONSE_TO_HEAT | 0,9651007 | 1 0,0285744 0,2518744 0,5488524 | 17 |
| GOBP_POSITIVE_REGULATION_OF_TRANSLATION | 0,9977827 | 1 0,0152836 0,1923149 0,5481073 | 123 |
| GOBP_TRNA_PROCESSING | 0,9977778 | 1 0,0154514 0,1925871 0,5481047 | 122 |
| GOBP_TRANSCRIPTION_BY_RNA_POLYMERASE_I | 0,993865 | 1 0,0221551 0,2106883 0,545941 | 53 |
| GOBP_POSITIVE_REGULATION_OF_EPITHELIAL_CELL_APOPTOTIC_PROCESS | 0,9786951 | 1 0,027448 0,2408889 0,5452484 | 22 |
| GOBP_AMP_METABOLIC_PROCESS | 0,9752604 | 1 0,0264649 0,2367447 0,5451425 | 23 |
| GOBP_NEGATIVE_REGULATION_OF_TRANSCRIPTION_REGULATORY_REGION_DNA_BINDING | 0,967655 | 1 0,028645 0,2467015 0,5442472 | 18 |
| GOBP_REGULATION_OF_CYTOPLASMIC_TRANSPORT | 0,9830729 | 1 0,0260438 0,2308677 0,544225 | 28 |
| GOBP_CELLULAR_RESPONSE_TO_TOPOLOGICALLY_INCORRECT_PROTEIN | 0,9988739 | 1 0,0163561 0,1928212 0,5413269 | 105 |
| GOBP_REGULATION_OF_CALCIUM_ION_DEPENDENT_EXOCYTOSIS | 0,9911728 | 1 0,0238623 0,2178985 0,5369042 | 37 |
| GOBP_MITOCHONDRION_ORGANIZATION | 1 | 1 0,006346 0,173343 0,5362687 | 479 |
| GOBP_NEGATIVE_REGULATION_OF_MIRNA_TRANSCRIPTION | 0,9730458 | 1 0,0283628 0,2412598 0,5322423 | 18 |
| GOBP_POSITIVE_REGULATION_OF_MACROAUTOPHAGY | 0,997619 | 1 0,0201218 0,1976997 0,5294584 | 68 |
| GOBP_SYNAPTIC_VESICLE_TRANSPORT | 0,99875 | 1 0,0229406 0,2131423 0,5285361 | 39 |
| GOBP_HISTONE_LYSINE_DEMETHYLATION | 0,9882813 | 1 0,025763 0,2237087 0,5245085 | 27 |
| GOBP_SYNAPTIC_VESICLE_PRIMING | 0,9757412 | 1 0,0282219 0,2372433 0,5233814 | 18 |
| GOBP_POSITIVE_REGULATION_OF_P38MAPK_CASCADE | 0,9844761 | 1 0,0256225 0,2245872 0,5233707 | 25 |
| GOBP_CYTOPLASMIC_TRANSLATIONAL_INITIATION | 0,9831606 | 1 0,025763 0,2253041 0,5233415 | 24 |
| GOBP_AMINO_ACID_ACTIVATION | 0,9937107 | 1 0,0235793 0,206882 0,5230503 | 44 |
| GOBP_REGULATION_OF_INTRINSIC_APOPTOTIC_SIGNALING_PATHWAY_BY_P53_CLASS_MEDIATOR | 0,9872286 | 1 0,024779 0,219378 0,5224589 | 30 |
| GOBP_PROTEIN_LOCALIZATION_TO_GOLGI_APPARATUS | 0,9895833 | 1 0,0256927 0,2223078 0,5212238 | 27 |
| GOBP_TRNA_MODIFICATION | 0,9977221 | 1 0,0172343 0,188248 0,5211826 | 87 |
| GOBP_PURINE_CONTAINING_COMPOUND_TRANSMEMBRANE_TRANSPORT | 0,9786382 | 1 0,0275886 0,2341931 0,5208311 | 19 |
| GOBP_THYMOCYTE_APOPTOTIC_PROCESS | 0,9661247 | 1 0,0289981 0,2453402 0,5174921 | 15 |
| GOBP_STEROL_BIOSYNTHETIC_PROCESS | 0,9963325 | 1 0,021796 0,1985992 0,5132563 | 52 |
| GOBP_REGULATION_OF_CARDIAC_MUSCLE_CELL_MEMBRANE_REPOLARIZATION | 0,9826435 | 1 0,0273777 0,2300352 0,511584 | 19 |
| GOBP_POSITIVE_REGULATION_OF_TRANSCRIPTION_BY_RNA_POLYMERASE_I | 0,9908854 | 1 0,0256225 0,2219777 0,5111392 | 23 |
| GOBP_ORGANELLE_MEMBRANE_FUSION | 1 | 1 0,0166782 0,1801662 0,5034973 | 102 |
| GOBP_RIBOSOMAL_SMALL_SUBUNIT_ASSEMBLY | 0,9865772 | 1 0,027448 0,2303716 0,5019961 | 17 |
| GOBP_NUCLEOBASE_CONTAINING_SMALL_MOLECULE_BIOSYNTHETIC_PROCESS | 0,9742547 | 1 0,0285744 0,2371986 0,500319 | 15 |
| GOBP_MULTIVESICULAR_BODY_SORTING_PATHWAY | 0,9949622 | 1 0,0235793 0,2018226 0,4986462 | 38 |
| GOBP_MITOCHONDRIAL_TRANSCRIPTION | 0,9865772 | 1 0,027448 0,2287732 0,4985132 | 17 |
| GOBP_UBIQUINONE_METABOLIC_PROCESS | 0,9851752 | 1 0,0277292 0,2246881 0,4956834 | 18 |
| GOBP_NEGATIVE_REGULATION_OF_STEROID_METABOLIC_PROCESS | 0,9893475 | 1 0,0268861 0,2188394 0,4953396 | 22 |
| GOBP_REGULATION_OF_MEMBRANE_REPOLARIZATION | 0,9896507 | 1 0,0253416 0,2124187 0,4950135 | 25 |
| GOBP_MATURATION_OF_SSU_RRNA | 0,9987437 | 1 0,0232248 0,1931564 0,4919174 | 48 |
| GOBP_NEGATIVE_REGULATION_OF_MACROAUTOPHAGY | 0,998739 | 1 0,0234376 0,2006918 0,4903619 | 35 |
| GOBP_RRNA_METHYLATION | 0,9896507 | 1 0,0253416 0,2100511 0,4894962 | 25 |
| GOBP_GLUCOSE_6_PHOSPHATE_METABOLIC_PROCESS | 0,9893617 | 1 0,0268159 0,2167763 0,4870787 | 20 |
| GOBP_POST_GOLGI_VESICLE_MEDIATED_TRANSPORT | 1 | 1 0,016275 0,1718329 0,4817734 | 103 |
| GOBP_PEROXISOMAL_TRANSPORT | 0,9933422 | 1 0,0266755 0,2088787 0,4727938 | 22 |
| GOBP_NEGATIVE_REGULATION_OF_TOR_SIGNALING | 0,9987294 | 1 0,0238623 0,1876049 0,4717812 | 43 |
| GOBP_PROTEIN_INSERTION_INTO_MEMBRANE | 1 | 1 0,0214354 0,1800467 0,4695058 | 55 |
| GOBP_REGULATION_OF_TRANSLATIONAL_FIDELITY | 0,9946595 | 1 0,0267457 0,2101415 0,4673418 | 19 |
| GOBP_GLUCOSE_CATABOLIC_PROCESS | 0,9919137 | 1 0,0273777 0,2116914 0,4670115 | 18 |
| GOBP_PROTEIN_LOCALIZATION_TO_MITOCHONDRION | 1 | 1 0,0150299 0,1640457 0,4665799 | 118 |
| GOBP_ENDOSOME_TO_LYSOSOME_TRANSPORT | 1 | 1 0,0211456 0,1766301 0,463774 | 57 |
| GOBP_SELECTIVE_AUTOPHAGY | 1 | 1 0,0192297 0,1713763 0,4627926 | 75 |
| GOBP_REGULATION_OF_TRANSCRIPTION_BY_RNA_POLYMERASE_I | 0,9974906 | 1 0,0232248 0,1901905 0,4626848 | 33 |
| GOBP_MACROAUTOPHAGY | 1 | 1 0,0097825 0,1534524 0,4621733 | 298 |
| GOBP_MATURATION_OF_SSU_RRNA_FROM_TRICISTRONIC_RRNA_TRANSCRIPT_SSU_RRNA_5_8S_RRNA_LSU_RRNA | 0,998739 | 1 0,0234376 0,1887186 0,4611071 | 35 |
| GOBP_NEGATIVE_REGULATION_OF_ORGAN_GROWTH | 0,9934896 | 1 0,0254821 0,1937267 0,4542126 | 27 |
| GOBP_MITOCHONDRIAL_TRANSPORT | 1 | 1 0,0121213 0,1539382 0,4525931 | 183 |
| GOBP_HISTONE_MRNA_CATABOLIC_PROCESS | 0,9864499 | 1 0,0279403 0,2141084 0,4516152 | 15 |
| GOBP_PROTEIN_TRANSMEMBRANE_TRANSPORT | 1 | 1 0,0207088 0,1710112 0,4511266 | 60 |
| GOBP_RUFFLE_ASSEMBLY | 0,9987406 | 1 0,0233667 0,180957 0,4470932 | 38 |
| GOBP_RESPONSE_TO_MITOCHONDRIAL_DEPOLARISATION | 0,9959947 | 1 0,0266755 0,197208 0,4385783 | 19 |
| GOBP_GPI_ANCHOR_METABOLIC_PROCESS | 0,9974457 | 1 0,0242154 0,1835872 0,4372215 | 30 |
| GOBP_MITOCHONDRIAL_MEMBRANE_ORGANIZATION | 1 | 1 0,0163561 0,1538241 0,4324716 | 107 |
| GOBP_POSITIVE_REGULATION_OF_LIPID_CATABOLIC_PROCESS | 0,9918699 | 1 0,0276589 0,2045184 0,4313871 | 15 |
| GOBP_RIBONUCLEOSIDE_TRIPHOSPHATE_METABOLIC_PROCESS | 1 | 1 0,0192297 0,1573965 0,4250409 | 75 |
| GOBP_DEOXYRIBONUCLEOSIDE_TRIPHOSPHATE_METABOLIC_PROCESS | 0,995935 | 1 0,027448 0,1932748 0,4076712 | 15 |
| GOBP_MITOCHONDRIAL_FUSION | 0,9987229 | 1 0,0241448 0,1706752 0,4064709 | 30 |
| GOBP_PROTEIN_TARGETING_TO_PEROXISOME | 0,9959569 | 1 0,027167 0,1841347 0,4062188 | 18 |
| GOBP_PEPTIDYL_SERINE_DEPHOSPHORYLATION | 0,9973298 | 1 0,0266053 0,1813366 0,4032815 | 19 |
| GOBP_VIRAL_TRANSLATION | 0,9986523 | 1 0,0270265 0,1724156 0,3803653 | 18 |
| GOBP_MITOPHAGY | 0,9987453 | 1 0,0231538 0,1502945 0,3656282 | 33 |
| GOBP_AUTOPHAGOSOME_MATURATION | 1 | 1 0,0207088 0,1150029 0,303377 | 60 |
| GOBP_ENDOSOME_ORGANIZATION | 1 | 1 0,0183973 0,1076048 0,2947303 | 82 |
| GOBP_PROTEIN_TRANSMEMBRANE_IMPORT_INTO_INTRACELLULAR_ORGANELLE | 1 | 1 0,0234376 0,1199154 0,2917351 | 34 |
| GOBP_RRNA_MODIFICATION | 1 | 1 0,0888945 -0,1158981 -0,3220462 | 36 |
| GOBP_REGULATION_OF_RECEPTOR_LOCALIZATION_TO_SYNAPSE | 0,9960938 | 1 0,0780892 -0,1608188 -0,404839 | 16 |
| GOBP_P_BODY_ASSEMBLY | 0,9960317 | 1 0,078921 -0,1582298 -0,4077291 | 21 |
| GOBP_MITOCHONDRIAL_RNA_METABOLIC_PROCESS | 1 | 1 0,0937465 -0,1575272 -0,4625491 | 49 |
| GOBP_GOLGI_TO_ENDOSOME_TRANSPORT | 0,9961089 | 1 0,077884 -0,1829808 -0,4645506 | 17 |
| GOBP_MULTIVESICULAR_BODY_ORGANIZATION | 1 | 1 0,0873098 -0,1737682 -0,4742603 | 31 |
| GOBP_ESTABLISHMENT_OF_PROTEIN_LOCALIZATION_TO_MITOCHONDRIAL_MEMBRANE | 1 | 1 0,0845557 -0,1758962 -0,4789913 | 29 |
| GOBP_PROTEIN_K48_LINKED_DEUBIQUITINATION | 1 | 1 0,0899861 -0,1784565 -0,4829235 | 33 |
| GOBP_REGULATION_OF_AEROBIC_RESPIRATION | 0,9871795 | 1 0,0833634 -0,1787278 -0,4923076 | 28 |
| GOBP_PROTEIN_INSERTION_INTO_ER_MEMBRANE | 0,9871795 | 1 0,0833634 -0,188206 -0,4934146 | 23 |
| GOBP_DE_NOVO_PROTEIN_FOLDING | 0,9952153 | 1 0,0891647 -0,1762815 -0,4971076 | 37 |
| GOBP_MITOCHONDRIAL_TRANSLATION | 0,9935065 | 1 0,1075544 -0,1599368 -0,4978552 | 72 |
| GOBP_INTRACELLULAR_PROTEIN_TRANSMEMBRANE_TRANSPORT | 1 | 1 0,0937465 -0,1709159 -0,5062435 | 50 |
| GOBP_INNER_MITOCHONDRIAL_MEMBRANE_ORGANIZATION | 0,9951691 | 1 0,0897105 -0,1760136 -0,5079283 | 40 |
| GOBP_ATP_BIOSYNTHETIC_PROCESS | 0,9954128 | 1 0,086795 -0,1730591 -0,5086417 | 45 |
| GOBP_LATE_ENDOSOME_TO_VACUOLE_TRANSPORT | 0,9869565 | 1 0,0843144 -0,1933665 -0,5116823 | 24 |
| GOBP_ORGANELLE_DISASSEMBLY | 1 | 1 0,1395997 -0,1534311 -0,5129304 | 117 |
| GOBP_MITOCHONDRIAL_OUTER_MEMBRANE_PERMEABILIZATION_INVOLVED_IN_PROGRAMMED_CELL_DEATH | 1 | 1 0,0886261 -0,1871793 -0,5152377 | 34 |
| GOBP_MAINTENANCE_OF_PROTEIN_LOCALIZATION_IN_ORGANELLE | 0,9951691 | 1 0,0897105 -0,1807179 -0,5215038 | 40 |
| GOBP_MITOCHONDRIAL_GENE_EXPRESSION | 1 | 1 0,1294429 -0,1580551 -0,5225307 | 104 |
| GOBP_ENDOSOME_TRANSPORT_VIA_MULTIVESICULAR_BODY_SORTING_PATHWAY | 0,9953488 | 1 0,0875697 -0,1934486 -0,5279735 | 31 |
| GOBP_NEGATIVE_REGULATION_OF_OXIDATIVE_STRESS_INDUCED_NEURON_DEATH | 0,9807692 | 1 0,0780892 -0,2085859 -0,5298599 | 18 |
| GOBP_REGULATION_OF_TRANSLATIONAL_INITIATION | 0,9934641 | 1 0,1079724 -0,1692609 -0,530143 | 74 |
| GOBP_REGULATION_OF_RRNA_PROCESSING | 0,9882813 | 1 0,0785029 -0,2107795 -0,5306079 | 16 |
| GOBP_POSITIVE_REGULATION_OF_TRANSLATIONAL_INITIATION | 0,9871795 | 1 0,0833634 -0,204441 -0,5359775 | 23 |
| GOBP_CHAPERONE_COFACTOR_DEPENDENT_PROTEIN_REFOLDING | 0,982906 | 1 0,0835991 -0,194801 -0,5365812 | 28 |
| GOBP_CRISTAE_FORMATION | 0,9882813 | 1 0,0785029 -0,2136364 -0,5377997 | 16 |
| GOBP_TRANSLATIONAL_INITIATION | 1 | 1 0,1294429 -0,1635652 -0,5422221 | 109 |
| GOBP_NEGATIVE_REGULATION_OF_INTRINSIC_APOPTOTIC_SIGNALING_PATHWAY_BY_P53_CLASS_MEDIATOR | 0,9761905 | 1 0,0799859 -0,2114401 -0,5448423 | 21 |
| GOBP_ESTABLISHMENT_OF_PROTEIN_LOCALIZATION_TO_PLASMA_MEMBRANE | 0,994186 | 1 0,1006334 -0,1811674 -0,5459357 | 60 |
| GOBP_GLUCOCORTICOID_METABOLIC_PROCESS | 0,9804688 | 1 0,078921 -0,2175791 -0,547725 | 16 |
| GOBP_PYRUVATE_METABOLIC_PROCESS | 1 | 1 0,1198878 -0,1716535 -0,5502398 | 91 |
| GOBP_AUTOPHAGY_OF_MITOCHONDRION | 0,9844961 | 1 0,1198878 -0,1757686 -0,5538987 | 85 |
| GOBP_REGULATION_OF_SYNAPTIC_VESICLE_RECYCLING | 0,964 | 1 0,0810802 -0,2172189 -0,5574678 | 20 |
| GOBP_ER_NUCLEUS_SIGNALING_PATHWAY | 0,9903382 | 1 0,0899861 -0,193524 -0,5593895 | 44 |
| GOBP_NEGATIVE_REGULATION_OF_CARDIAC_MUSCLE_TISSUE_GROWTH | 0,964 | 1 0,0810802 -0,2179827 -0,559428 | 20 |
| GOBP_TERTIARY_ALCOHOL_METABOLIC_PROCESS | 0,9688716 | 1 0,0793435 -0,2213788 -0,5620353 | 17 |
| GOBP_PROTEIN_TARGETING | 1 | 1 0,2249661 -0,1518934 -0,5621795 | 293 |
| GOBP_PHOSPHATIDYLETHANOLAMINE_METABOLIC_PROCESS | 0,9786325 | 1 0,0838361 -0,2145752 -0,5625462 | 23 |
| GOBP_ENDOPLASMIC_RETICULUM_ORGANIZATION | 1 | 1 0,1215433 -0,1788116 -0,5629098 | 87 |
| GOBP_AMYLOID_FIBRIL_FORMATION | 0,9779736 | 1 0,0855357 -0,2100589 -0,5632527 | 26 |
| GOBP_TELOMERASE_RNA_LOCALIZATION | 0,9644269 | 1 0,08042 -0,2182821 -0,5633677 | 19 |
| GOBP_REGULATION_OF_AUTOPHAGOSOME_MATURATION | 0,96875 | 1 0,0795565 -0,2241528 -0,5642733 | 16 |
| GOBP_PEPTIDYL_PROLINE_MODIFICATION | 1 | 1 0,0937465 -0,1918427 -0,5682275 | 50 |
| GOBP_MITOCHONDRIAL_ELECTRON_TRANSPORT_NADH_TO_UBIQUINONE | 0,985782 | 1 0,0891647 -0,1964328 -0,5694048 | 42 |
| GOBP_RNA_INTERFERENCE | 0,9688716 | 1 0,0793435 -0,2252258 -0,5718019 | 17 |
| GOBP_REGULATION_OF_MITOCHONDRIAL_OUTER_MEMBRANE_PERMEABILIZATION_INVOLVED_IN_APOPTOTIC_SIGNALING_PATHWAY | 0,9576923 | 1 0,0793435 -0,2260072 -0,5741143 | 18 |
| GOBP_POSITIVE_REGULATION_OF_DNA_TEMPLATED_TRANSCRIPTION_ELONGATION | 0,9786325 | 1 0,0838361 -0,2092431 -0,5763621 | 28 |
| GOBP_LIPOPROTEIN_CATABOLIC_PROCESS | 0,9583333 | 1 0,0785029 -0,2325398 -0,580217 | 15 |
| GOBP_PURINE_RIBONUCLEOSIDE_METABOLIC_PROCESS | 0,9486166 | 1 0,0813027 -0,2255391 -0,5820972 | 19 |
| GOBP_VESICLE_ORGANIZATION | 1 | 1 0,2192503 -0,1586092 -0,5847109 | 283 |
| GOBP_BEHAVIORAL_RESPONSE_TO_PAIN | 0,9507576 | 1 0,078921 -0,2346058 -0,585372 | 15 |
| GOBP_REGULATION_OF_EARLY_ENDOSOME_TO_LATE_ENDOSOME_TRANSPORT | 0,9346154 | 1 0,0806388 -0,230647 -0,5859006 | 18 |
| GOBP_FIBROBLAST_MIGRATION | 0,9816514 | 1 0,0875697 -0,2004189 -0,5890557 | 45 |
| GOBP_REGULATION_OF_TRIGLYCERIDE_BIOSYNTHETIC_PROCESS | 0,9335938 | 1 0,0815265 -0,2344253 -0,5901331 | 16 |
| GOBP_MODULATION_BY_HOST_OF_VIRAL_GENOME_REPLICATION | 0,9377432 | 1 0,0810802 -0,2330616 -0,5916954 | 17 |
| GOBP_NUCLEOTIDE_PHOSPHORYLATION | 1 | 1 0,1198878 -0,1853737 -0,5930236 | 89 |
| GOBP_REGULATION_OF_ATP_BIOSYNTHETIC_PROCESS | 0,9377432 | 1 0,0810802 -0,2336178 -0,5931076 | 17 |
| GOBP_REGULATION_OF_ATP_METABOLIC_PROCESS | 0,9934211 | 1 0,1083943 -0,1888231 -0,5943687 | 75 |
| GOBP_GOLGI_TO_PLASMA_MEMBRANE_PROTEIN_TRANSPORT | 0,9613527 | 1 0,0916795 -0,2065883 -0,5961587 | 40 |
| GOBP_GLIAL_CELL_APOPTOTIC_PROCESS | 0,9431818 | 1 0,0793435 -0,2391059 -0,5966004 | 15 |
| GOBP_REGULATION_OF_RECEPTOR_BINDING | 0,9328063 | 1 0,0822055 -0,2323072 -0,5995654 | 19 |
| GOBP_REGULATION_OF_DNA_STRAND_ELONGATION | 0,9299611 | 1 0,0815265 -0,2377114 -0,6035003 | 17 |
| GOBP_NECROTIC_CELL_DEATH | 0,9782609 | 1 0,0975449 -0,2050219 -0,603906 | 52 |
| GOBP_POSITIVE_REGULATION_OF_GLYCOPROTEIN_METABOLIC_PROCESS | 0,968254 | 1 0,08042 -0,2357284 -0,6074288 | 21 |
| GOBP_PROTEIN_TARGETING_TO_MEMBRANE | 1 | 1 0,135002 -0,1828601 -0,6080063 | 114 |
| GOBP_PROTEIN_MATURATION_BY_IRON_SULFUR_CLUSTER_TRANSFER | 0,9296875 | 1 0,0817516 -0,241697 -0,6084384 | 16 |
| GOBP_LATE_ENDOSOME_TO_VACUOLE_TRANSPORT_VIA_MULTIVESICULAR_BODY_SORTING_PATHWAY | 0,92607 | 1 0,0817516 -0,2403037 -0,6100816 | 17 |
| GOBP_REGULATION_OF_STEM_CELL_POPULATION_MAINTENANCE | 0,9700855 | 1 0,0843144 -0,2221474 -0,6119071 | 28 |
| GOBP_FOLIC_ACID_CONTAINING_COMPOUND_METABOLIC_PROCESS | 0,965812 | 1 0,0845557 -0,2252846 -0,6137293 | 27 |
| GOBP_MITOCHONDRIAL_RNA_PROCESSING | 0,956 | 1 0,0815265 -0,2393914 -0,614371 | 20 |
| GOBP_POSITIVE_REGULATION_OF_DENDRITE_MORPHOGENESIS | 0,9658537 | 1 0,0919686 -0,2275783 -0,6158524 | 33 |
| GOBP_RESPONSE_TO_INTERFERON_ALPHA | 0,9182879 | 1 0,0822055 -0,2430515 -0,6170577 | 17 |
| GOBP_REGULATION_OF_RUFFLE_ASSEMBLY | 0,9647577 | 1 0,0862866 -0,2304671 -0,6179751 | 26 |
| GOBP_PEPTIDYL_ASPARAGINE_MODIFICATION | 0,957265 | 1 0,0850427 -0,2358246 -0,6182551 | 23 |
| GOBP_CELLULAR_RESPONSE_TO_UNFOLDED_PROTEIN | 0,983871 | 1 0,1226792 -0,1974066 -0,6195466 | 86 |
| GOBP_PROTEIN_LOCALIZATION_TO_PHAGOPHORE_ASSEMBLY_SITE | 0,9143969 | 1 0,0824344 -0,2451841 -0,6224719 | 17 |
| GOBP_SNRNA_TRANSCRIPTION_BY_RNA_POLYMERASE_II | 0,9128788 | 1 0,0810802 -0,2507727 -0,6257105 | 15 |
| GOBP_POSITIVE_REGULATION_OF_CELL_MIGRATION_INVOLVED_IN_SPROUTING_ANGIOGENESIS | 0,948 | 1 0,0819779 -0,2438407 -0,6257897 | 20 |
| GOBP_GTP_METABOLIC_PROCESS | 0,9529915 | 1 0,0852885 -0,2389003 -0,6263186 | 23 |
| GOBP_ARGININE_METABOLIC_PROCESS | 0,9090909 | 1 0,0813027 -0,252242 -0,6293766 | 15 |

GOBP_INFLAMMASOME_COMPLEX_ASSEMBLY	0,9105058	1	0,0826646	-0,2483775 -0,6305794	17

GOBP_REGULATION_OF_NEURON_PROJECTION_REGENERATION	0,9559471	1	0,086795 -0,2361196 -0,6331319	26

GOBP_SNRNA_TRANSCRIPTION	0,9153846	1	0,0817516	-0,2508183 -0,6371406	18

GOBP_REGULATION_OF_TRANSLATION_IN_RESPONSE_TO_STRESS	0,94	1	0,0824344 -0,2483026 -0,6372408	20

GOBP_POSTTRANSLATIONAL_PROTEIN_TARGETING_TO_ENDOPLASMIC_RETICULUM_MEMBRANE	0,8977273	1	0,0819779 -0,2555294 -0,6375793	15

GOBP_POSITIVE_REGULATION_OF_MITOCHONDRION_ORGANIZATION	0,9817073	1	0,104344 -0,2087332 -0,6383037	67

GOBP_POLYAMINE_BIOSYNTHETIC_PROCESS	0,8939394	1	0,0822055	-0,2563964 -0,6397424	15

GOBP_NEGATIVE_REGULATION_OF_LYMPHOCYTE_MEDIATED_IMMUNITY	0,9634703	1	0,0883594 -0,2348739 -0,6406425	30

GOBP_ENDONUCLEOLYTIC_CLEAVAGE_INVOLVED_IN_RRNA_PROCESSING	0,8939394	1	0,0822055	-0,2568329 -0,6408316	15

GOBP_CYTOCHROME_COMPLEX_ASSEMBLY	0,9473684	1	0,0919686	-0,2312238 -0,6409582	35

GOBP_REGULATION_OF_PROTEIN_TARGETING_TO_MEMBRANE	0,972093	1	0,0888945 -0,2352259 -0,6419952	31

GOBP_PEPTIDE_CATABOLIC_PROCESS	0,9484127	1	0,0815265 -0,2492599 -0,6422969	21

GOBP_ENDOSOMAL_TRANSPORT	1	1	0,1882041	-0,179299 -0,6443309	224

GOBP_NEURON_DEATH_IN_RESPONSE_TO_OXIDATIVE_STRESS	0,9358974	1	0,0862866 -0,2339418	-0,644395	28

GOBP_REGULATION_OF_VACUOLE_ORGANIZATION	0,9470899	1	0,0978773	-0,2167671 -0,6458762	51

GOBP_PROTEIN_LOCALIZATION_TO_VACUOLE	0,9698795	1	0,104344 -0,2132182 -0,6502513	66

GOBP_ESTABLISHMENT_OF_PROTEIN_LOCALIZATION_TO_VACUOLE	0,9805825	1	0,0908241 -0,2212132	-0,653671	48

GOBP_NEGATIVE_REGULATION_OF_ADAPTIVE_IMMUNE_RESPONSE	0,9463415	1	0,0931455 -0,2421014 -0,6551538	33

GOBP_PEPTIDYL_ARGININE_MODIFICATION	0,9038462	1	0,0824344	-0,25791 -0,6551555	18

GOBP_RESPIRATORY_CHAIN_COMPLEX_IV_ASSEMBLY	0,9213974	1	0,0883594	-0,2459375 -0,6577947	25

GOBP_POSITIVE_REGULATION_OF_LAMELLIPODIUM_ORGANIZATION	0,9627907	1	0,0894367 -0,2417312 -0,6597499	31

GOBP_MEMBRANE_LIPID_CATABOLIC_PROCESS	0,9543379	1	0,0888945	-0,2421807 -0,6605725	30

GOBP_PROTEIN_DEMANNOSYLATION	0,8832685	1	0,0843144	-0,2610032 -0,6626335	17

GOBP_MODIFIED_AMINO_ACID_TRANSPORT	0,9059829	1	0,0880945 -0,2415401 -0,6653245	28

GOBP_STEM_CELL_DIVISION	0,8951965	1	0,0899861	-0,2499241 -0,6684577	25

GOBP_REGULATION_OF_MITOCHONDRIAL_GENE_EXPRESSION	0,9059829	1	0,0880945 -0,2429787 -0,6692872	28

GOBP_REGULATION_OF_CARBOHYDRATE_CATABOLIC_PROCESS	0,953125	1	0,096563 -0,2260186 -0,6694547	50

GOBP_TRANSCRIPTION_BY_RNA_POLYMERASE_III	0,9465241	1	0,09855 -0,2260472 -0,6702719	53

GOBP_NEGATIVE_REGULATION_OF_CHONDROCYTE_DIFFERENTIATION	0,8793774	1	0,0845557 -0,2641632	-0,670656	17

GOBP_REGULATION_OF_ERAD_PATHWAY	0,912	1	0,0840746	-0,261353	-0,670733	20

GOBP_PTERIDINE_CONTAINING_COMPOUND_METABOLIC_PROCESS	0,9219512	1	0,0946646	-0,2486203 -0,6727944	33

GOBP_REGULATION_OF_GLUCONEOGENESIS	0,9449541	1	0,0897105	-0,2296233 -0,6748909	45

GOBP_UBIQUITIN_DEPENDENT_ERAD_PATHWAY	0,9545455	1	0,1204334 -0,2158986 -0,6756207	83

GOBP_REGULATION_OF_CELL_KILLING	0,9304813	1	0,0995791 -0,2280391	-0,676178	53

GOBP_NEGATIVE_REGULATION_OF_TYPE_I_INTERFERON_PRODUCTION	0,9162996	1	0,0891647	-0,252231 -0,6763331	26

GOBP_POSITIVE_REGULATION_OF_LONG_TERM_SYNAPTIC_POTENTIATION	0,8961538	1	0,0828962 -0,2669417	-0,678098	18

GOBP_REGULATION_OF_MITOCHONDRIAL_MEMBRANE_PERMEABILITY	0,9347826	1	0,1002791 -0,2305721 -0,6791659	52

GOBP_CELLULAR_MODIFIED_AMINO_ACID_CATABOLIC_PROCESS	0,8974359	1	0,0886261	-0,2498353 -0,6806114	27

GOBP_BICARBONATE_TRANSPORT	0,9206349	1	0,0831291	-0,2642751 -0,6809884	21

GOBP_EMBRYO_IMPLANTATION	0,940367	1	0,0899861 -0,2320249 -0,6819495	45

GOBP_POSITIVE_REGULATION_OF_TELOMERASE_RNA_LOCALIZATION_TO_CAJAL_BODY	0,8712121	1	0,0835991	-0,27465 -0,6852876	15

GOBP_ERAD_PATHWAY	0,9824561	1	0,1287887 -0,2071276 -0,6867069	105

GOBP_MITOCHONDRIAL_CYTOCHROME_C_OXIDASE_ASSEMBLY	0,9123506	1	0,0838361 -0,2626936 -0,6888821	22

GOBP_PHOSPHATE_ION_TRANSPORT	0,9123506	1	0,0838361	-0,2629176 -0,6894697	22

GOBP_REGULATION_OF_SUBSTRATE_ADHESION_DEPENDENT_CELL_SPREADING	0,9312169	1	0,0988903 -0,2314251	-0,689551	51

GOBP_TYPE_I_INTERFERON_PRODUCTION	0,9612403	1	0,1215433	-0,2188961 -0,6898061	85

GOBP_MESODERMAL_CELL_DIFFERENTIATION	0,908	1	0,0843144	-0,268853	-0,689981	20

GOBP_MITOCHONDRIAL_RESPIRATORY_CHAIN_COMPLEX_ASSEMBLY	0,9685039	1	0,1221079	-0,2162281 -0,6917289	89

GOBP_GLYCOSYL_COMPOUND_BIOSYNTHETIC_PROCESS	0,8853755	1	0,0850427	-0,2683231 -0,6925192	19

GOBP_REGULATION_OF_NATURAL_KILLER_CELL_MEDIATED_IMMUNITY	0,8646288	1	0,0919686 -0,2601989 -0,6959391	25

GOBP_ANTIMICROBIAL_HUMORAL_IMMUNE_RESPONSE_MEDIATED_BY_ANTIMICROBIAL_PEPTIDE	0,8646288	1	0,0919686	-0,260277 -0,6961478	25

GOBP_CHAPERONE_MEDIATED_PROTEIN_FOLDING	0,9181287	1	0,1059203	-0,2312311 -0,6962308	61

GOBP_MEMBRANE_FUSION	0,9895833	1	0,1412251 -0,2051301 -0,6962719	134

GOBP_MAINTENANCE_OF_PROTEIN_LOCATION_IN_NUCLEUS	0,9003984	1	0,0845557 -0,2664489 -0,6987301	22

GOBP_POSITIVE_REGULATION_OF_CYTOKINE_PRODUCTION_INVOLVED_IN_INFLAMMATORY_RESPONSE	0,8692308	1	0,0845557 -0,2758782 -0,7007989	18

GOBP_MITOCHONDRIAL_TRANSMEMBRANE_TRANSPORT	0,9538462	1	0,1215433	-0,2156328 -0,7022388	94

GOBP_CALCIUM_ION_REGULATED_EXOCYTOSIS	0,9186047	1	0,1055209	-0,2333408 -0,7031569	60

GOBP_REGULATION_OF_CARDIOCYTE_DIFFERENTIATION	0,8924303	1	0,0850427	-0,2681603 -0,7032181	22

GOBP_DNA_DEALKYLATION	0,8846154	1	0,0894367	-0,2691648 -0,7056621	23

GOBP_CELLULAR_RESPONSE_TO_INTERFERON_GAMMA	0,9586207	1	0,1137873	-0,2256698 -0,7065626	77

GOBP_ANTIVIRAL_INNATE_IMMUNE_RESPONSE	0,8409091	1	0,0855357 -0,2845298	-0,709939	15

GOBP_REGULATION_OF_BICELLULAR_TIGHT_JUNCTION_ASSEMBLY	0,8365759	1	0,0873098	-0,2799299 -0,7106844	17

GOBP_MITOCHONDRIAL_OUTER_MEMBRANE_PERMEABILIZATION	0,9026549	1	0,0902635 -0,2610895 -0,7109855	29

GOBP_TRANSLATIONAL_ELONGATION	0,9065934	1	0,1028218	-0,2392285 -0,7114782	55

GOBP_COENZYME_A_METABOLIC_PROCESS	0,8576923	1	0,0852885	-0,2801922 -0,7117576	18

GOBP_GOLGI_VESICLE_TRANSPORT	1	1	0,222056 -0,1942034 -0,7117898	282

GOBP_POSITIVE_REGULATION_OF_CELLULAR_PROTEIN_CATABOLIC_PROCESS	0,9764706	1	0,1521449 -0,2085265 -0,7134739	145

GOBP_DOPAMINE_TRANSPORT	0,8803828	1	0,0962406	-0,2481303 -0,7161478	41

GOBP_RETROGRADE_AXONAL_TRANSPORT	0,8928571	1	0,0847985	-0,2786954 -0,7181469	21

GOBP_MATERNAL_PROCESS_INVOLVED_IN_FEMALE_PREGNANCY	0,9205607	1	0,0922597	-0,2440628 -0,7194525	47

GOBP_PROTEIN_TARGETING_TO_VACUOLE	0,8995215	1	0,0949752 -0,2555031 -0,7205098	37

GOBP_NEGATIVE_REGULATION_OF_BIOMINERALIZATION	0,8287938	1	0,0878313 -0,2843438 -0,7218905	17

GOBP_VACUOLAR_TRANSPORT	0,962963	1	0,1574029	-0,2109703 -0,7230349	148

GOBP_REGULATION_OF_RELEASE_OF_CYTOCHROME_C_FROM_MITOCHONDRIA	0,8930233	1	0,0937465	-0,2500014 -0,7230475	43

GOBP_PSEUDOURIDINE_SYNTHESIS	0,8461538	1	0,0860347	-0,2847618 -0,7233656	18

GOBP_REGULATION_OF_NECROTIC_CELL_DEATH	0,861244	1	0,0975449 -0,2610178 -0,7235478	35

GOBP_PROTEIN_K63_LINKED_UBIQUITINATION	0,8846154	1	0,104344 -0,2446104 -0,7235756	54

GOBP_NUCLEAR_EXPORT	0,9770115	1	0,1501698 -0,2097638 -0,7244057	146

GOBP_VESICLE_TRANSPORT_ALONG_MICROTUBULE	0,8930233	1	0,0937465	-0,250558 -0,7246573	43

GOBP_POSITIVE_REGULATION_OF_NATURAL_KILLER_CELL_MEDIATED_IMMUNITY	0,8257576	1	0,08654 -0,2904786	-0,724782	15

GOBP_RESPONSE_TO_TOPOLOGICALLY_INCORRECT_PROTEIN	0,9642857	1	0,154191 -0,2100718 -0,7252242	147

GOBP_CALCIUM_INDEPENDENT_CELL_CELL_ADHESION_VIA_PLASMA_MEMBRANE_CELL_ADHESION_MOLECULES	0,8219697	1	0,086795 -0,2916094 -0,7276034	15

GOBP_NADH_DEHYDROGENASE_COMPLEX_ASSEMBLY	0,8877005	1	0,1024494	-0,2454598 -0,7278338	53

GOBP_PHOSPHATIDYLINOSITOL_DEPHOSPHORYLATION	0,8849558	1	0,0913924	-0,2673404 -0,7280076	29

GOBP_CARBOHYDRATE_DERIVATIVE_TRANSPORT	0,8963415	1	0,1101223	-0,2384304 -0,7291172	67

GOBP_MYELOID_CELL_ACTIVATION_INVOLVED_IN_IMMUNE_RESPONSE	0,8888889	1	0,1079724 -0,2425198 -0,7302207	61

GOBP_OLIGOSACCHARIDE_LIPID_INTERMEDIATE_BIOSYNTHETIC_PROCESS	0,8418972	1	0,0878313	-0,2830212 -0,7304537	19

GOBP_POSITIVE_REGULATION_OF_CELL_MORPHOGENESIS_INVOLVED_IN_DIFFERENTIATION	0,925	1	0,1096841 -0,2351945 -0,7321642	70

GOBP_REGULATION_OF_JNK_CASCADE	0,9693878	1	0,1412251 -0,2182423 -0,7324436	119

GOBP_PEPTIDYL_TYROSINE_AUTOPHOSPHORYLATION	0,8307692	1	0,0870516 -0,2888777	-0,733821	18

GOBP_VIRAL_RELEASE_FROM_HOST_CELL	0,8632479	1	0,0908241	-0,2699359 -0,7353702	27

GOBP_SALIVARY_GLAND_DEVELOPMENT	0,8493151	1	0,0956031	-0,2700063 -0,7364696	30

GOBP_METANEPHRIC_TUBULE_DEVELOPMENT	0,8307692	1	0,0870516 -0,2900081 -0,7366926	18

GOBP_NEGATIVE_REGULATION_OF_TORC1_SIGNALING	0,8379447	1	0,0880945 -0,2854952	-0,736839	19

GOBP_MODULATION_BY_HOST_OF_VIRAL_PROCESS	0,8589744	1	0,0911073 -0,2706766 -0,7373882	27

GOBP_POSITIVE_REGULATION_OF_TOLL_LIKE_RECEPTOR_SIGNALING_PATHWAY	0,8418803	1	0,0922597 -0,2817948	-0,738774	23

GOBP_COPPER_ION_HOMEOSTASIS	0,8379447	1	0,0880945	-0,2865165 -0,7394749	19

GOBP_POSITIVE_REGULATION_OF_SUBSTRATE_ADHESION_DEPENDENT_CELL_SPREADING	0,8516746	1	0,0982123	-0,2665411 -0,7406384	36

GOBP_INTERFERON_BETA_PRODUCTION	0,8586957	1	0,1055209	-0,2516976 -0,7413925	52

GOBP_NEGATIVE_REGULATION_OF_TRANSLATIONAL_INITIATION	0,8221344	1	0,0891647 -0,2882656 -0,7439892	19

GOBP_NEGATIVE_REGULATION_OF_INTRINSIC_APOPTOTIC_SIGNALING_PATHWAY	0,9212598	1	0,1256399 -0,2325776 -0,7440323	89

GOBP_LONG_TERM_SYNAPTIC_DEPRESSION	0,8333333	1	0,0928481	-0,2703277 -0,7446204	28

GOBP_AMYLOID_BETA_CLEARANCE	0,8461538	1	0,0919686	-0,2734483 -0,7449389	27

GOBP_SUPPRESSION_OF_VIRAL_RELEASE_BY_HOST	0,7954545	1	0,0886261	-0,2987092 -0,7453183	15

GOBP_ISOPRENOID_BIOSYNTHETIC_PROCESS	0,8461538	1	0,0919686	-0,2738375 -0,7459992	27

GOBP_NEGATIVE_REGULATION_OF_VIRAL_GENOME_REPLICATION	0,8785047	1	0,0949752	-0,2535355 -0,7473763	47

GOBP_VESICLE_CYTOSKELETAL_TRAFFICKING	0,8373494	1	0,1137873 -0,2466791	-0,747719	64

GOBP_TOLL_LIKE_RECEPTOR_9_SIGNALING_PATHWAY	0,8007813	1	0,0899861	-0,2980393 -0,7502723	16

GOBP_ANTIGEN_PROCESSING_AND_PRESENTATION_OF_PEPTIDE_ANTIGEN_VIA_MHC_CLASS_I	0,8247863	1	0,0934449	-0,2725392 -0,7507119	28

GOBP_NEGATIVE_REGULATION_OF_PROTEIN_KINASE_B_SIGNALING	0,8373206	1	0,0992333	-0,2701905 -0,7507788	36

GOBP_NEGATIVE_REGULATION_OF_GTPASE_ACTIVITY	0,8274336	1	0,095288 -0,2757962	-0,751034	29

GOBP_GOLGI_ORGANIZATION	0,9438202	1	0,1511488 -0,2212785 -0,7510501	140

GOBP_DOPAMINE_RECEPTOR_SIGNALING_PATHWAY	0,8238095	1	0,0999277	-0,273161 -0,7519142	34

GOBP_POSITIVE_REGULATION_OF_STRESS_ACTIVATED_PROTEIN_KINASE_SIGNALING_CASCADE	0,9565217	1	0,1301056 -0,2255004 -0,7520406	107

GOBP_INTERLEUKIN_8_PRODUCTION	0,8461538	1	0,1071402	-0,2550045 -0,7543221	54

GOBP_REGULATION_OF_LEUKOCYTE_DEGRANULATION	0,8082192	1	0,09855 -0,2772074 -0,7561114	30

GOBP_LENS_MORPHOGENESIS_IN_CAMERA_TYPE_EYE	0,836	1	0,0888945 -0,2955424 -0,7584762	20

GOBP_REGULATION_OF_MACROAUTOPHAGY	0,9240506	1	0,1631801 -0,2208623 -0,7592845	152

GOBP_MITOCHONDRIAL_ATP_SYNTHESIS_COUPLED_PROTON_TRANSPORT	0,78125	1	0,0913924	-0,302 -0,7602429	16

GOBP_REGULATION_OF_GLYCOLYTIC_PROCESS	0,8229665	1	0,1002791 -0,2634163 -0,7602657	41

GOBP_SPHINGOID_METABOLIC_PROCESS	0,8095238	1	0,0902635	-0,295313 -0,7609674	21

GOBP_PROTEIN_LOCALIZATION_TO_CELL_JUNCTION	0,8682171	1	0,1287887 -0,2415861 -0,7613088	85

GOBP_REGULATION_OF_PATTERN_RECOGNITION_RECEPTOR_SIGNALING_PATHWAY	0,8417266	1	0,1256399 -0,2426257 -0,7622778	80

GOBP_DEFENSE_RESPONSE_TO_GRAM_POSITIVE_BACTERIUM	0,8086124	1	0,1013507	-0,2641911 -0,7625022	41

GOBP_RENAL_TUBULAR_SECRETION	0,7884615	1	0,0899861	-0,3005238 -0,7634051	18

GOBP_FEAR_RESPONSE	0,8190476	1	0,1002791 -0,2822681 -0,7641868	32

GOBP_SUBSTRATE_DEPENDENT_CELL_MIGRATION	0,7991453	1	0,095288 -0,2920403 -0,7656342	23

GOBP_POSITIVE_REGULATION_OF_STEM_CELL_DIFFERENTIATION	0,7537879	1	0,0916795	-0,3069405 -0,7658565	15

GOBP_PEPTIDYL_CYSTEINE_MODIFICATION	0,8418605	1	0,0972151	-0,2649842 -0,7663802	43

GOBP_RESPONSE_TO_TYPE_I_INTERFERON	0,8351648	1	0,1079724	-0,2578597 -0,7668885	55

GOBP_B_CELL_RECEPTOR_SIGNALING_PATHWAY	0,7952381	1	0,1020801	-0,2797378 -0,7700179	34

GOBP_CLATHRIN_COAT_ASSEMBLY	0,7769231	1	0,0908241	-0,3032595 -0,7703544	18

GOBP_POSITIVE_REGULATION_OF_ERYTHROCYTE_DIFFERENTIATION	0,8105727	1	0,0962406	-0,2880396 -0,7723502	26

GOBP_DEMETHYLATION	0,8146067	1	0,1110115	-0,2574543 -0,7728851	58

GOBP_REGULATION_OF_CELLULAR_RESPIRATION	0,8186047	1	0,0988903	-0,2677976 -0,7745172	43

GOBP_CELLULAR_COPPER_ION_HOMEOSTASIS	0,7617188	1	0,0928481	-0,3082229 -0,7759082	16

GOBP_PROTEIN_K63_LINKED_DEUBIQUITINATION	0,7857143	1	0,1028218 -0,2822093	-0,776821	34

GOBP_POSITIVE_REGULATION_OF_PROTEOLYSIS_INVOLVED_IN_CELLULAR_PROTEIN_CATABOLIC_PROCESS	0,94	1	0,1420566	-0,230827	-0,779651	123

GOBP_PROTEIN_DEGLYCOSYLATION	0,7816594	1	0,0978773	-0,2916305 -0,7800073	25

GOBP_REGULATION_OF_COLLATERAL_SPROUTING	0,7470817	1	0,0937465	-0,3079262 -0,7817613	17

GOBP_LUNG_MORPHOGENESIS	0,7971014	1	0,1028218	-0,2709908 -0,7820073	40

GOBP_NEGATIVE_REGULATION_OF_STEM_CELL_DIFFERENTIATION	0,7857143	1	0,0919686	-0,303917 -0,7831382	21

GOBP_NEGATIVE_REGULATION_OF_LIPID_BIOSYNTHETIC_PROCESS	0,8058252	1	0,1024494	-0,2650867 -0,7833142	48

GOBP_COPII_COATED_VESICLE_BUDDING	0,7860465	1	0,1013507	-0,287022 -0,7833605	31

GOBP_POSITIVE_REGULATION_OF_INTERLEUKIN_12_PRODUCTION	0,7913043	1	0,0968878 -0,2967939 -0,7853698	24

GOBP_POSITIVE_REGULATION_OF_GLYCOLYTIC_PROCESS	0,7237354	1	0,0956031	-0,3103769 -0,7879831	17

GOBP_REGULATION_OF_NEUTROPHIL_MIGRATION	0,7767442	1	0,1020801 -0,2891923 -0,7892841	31

GOBP_INTESTINAL_ABSORPTION	0,7619048	1	0,0937465 -0,3063317 -0,7893606	21

GOBP_MEMBRANE_PROTEIN_PROTEOLYSIS	0,8235294	1	0,1071402	-0,2662527 -0,7894885	53

GOBP_POSITIVE_REGULATION_OF_PROTEIN_TARGETING_TO_MEMBRANE	0,784141	1	0,0982123 -0,2947198 -0,7902626	26

GOBP_REGULATION_OF_ALPHA_BETA_T_CELL_ACTIVATION	0,8246753	1	0,1198878	-0,2538916 -0,7903201	72

GOBP_NEUTROPHIL_MEDIATED_IMMUNITY	0,7606838	1	0,0982123 -0,3020734 -0,7919376	23

GOBP_ATP_SYNTHESIS_COUPLED_PROTON_TRANSPORT	0,7729084	1	0,0931455	-0,3021099 -0,7922466	22

GOBP_LEUKOCYTE_MEDIATED_CYTOTOXICITY	0,8116883	1	0,1209851	-0,2546781 -0,7927685	72

GOBP_RESPONSE_TO_CATECHOLAMINE	0,8024691	1	0,1182875	-0,2571912 -0,7935168	68

GOBP_PROTEIN_KINASE_A_SIGNALING	0,773913	1	0,0982123 -0,2999465 -0,7937121	24

GOBP_REGULATION_OF_SKELETAL_MUSCLE_CELL_DIFFERENTIATION	0,7307692	1	0,0943564	-0,3132166 -0,7956477	18

GOBP_REGULATION_OF_CARDIAC_MUSCLE_CONTRACTION_BY_REGULATION_OF_THE_RELEASE_OF_SEQUESTERED_CALCIUM_ION	0,7307692	1	0,0943564	-0,3138342 -0,7972168	18

GOBP_RETINAL_METABOLIC_PROCESS	0,7120623	1	0,096563	-0,314113 -0,7974682	17

GOBP_PEPTIDYL_THREONINE_DEPHOSPHORYLATION	0,7382813	1	0,0946646	-0,3168485 -0,7976218	16

GOBP_RIBONUCLEOSIDE_DIPHOSPHATE_METABOLIC_PROCESS	0,9243697	1	0,1301056 -0,2401968 -0,7983719	102

GOBP_POSITIVE_REGULATION_OF_PROTEASOMAL_UBIQUITIN_DEPENDENT_PROTEIN_CATABOLIC_PROCESS	0,8387097	1	0,1342735	-0,253937 -0,7994094	87

GOBP_RESPONSE_TO_ENDOPLASMIC_RETICULUM_STRESS	0,8958333	1	0,2165428	-0,22219 -0,7997417	236

GOBP_REGULATION_OF_T_CELL_MIGRATION	0,7442922	1	0,1035763	-0,2934377 -0,8003811	30

GOBP_HORMONE_BIOSYNTHETIC_PROCESS	0,7757009	1	0,1024494 -0,2744136	-0,801341	46

GOBP_POSITIVE_REGULATION_OF_PROTEIN_AUTOPHOSPHORYLATION	0,7393162	1	0,0999277 -0,3057003 -0,8014463	23

GOBP_RENAL_SODIUM_EXCRETION	0,7234848	1	0,0940503	-0,3214093 -0,8019582	15

GOBP_REGULATION_OF_CD4_POSITIVE_ALPHA_BETA_T_CELL_ACTIVATION	0,7844037	1	0,1006334	-0,2731028 -0,8026825	45

GOBP_REGULATION_OF_ACTIN_FILAMENT_LENGTH	0,9101124	1	0,154191 -0,2369751 -0,8043268	140

GOBP_REGULATION_OF_ENDOCRINE_PROCESS	0,7379913	1	0,1013507 -0,3008788	-0,804743	25

GOBP_DIOL_METABOLIC_PROCESS	0,7564103	1	0,09855 -0,2955611 -0,8051796	27

GOBP_POSITIVE_REGULATION_OF_REGULATED_SECRETORY_PATHWAY	0,7342995	1	0,1079724	-0,2793129 -0,8060226	40

GOBP_ALPHA_BETA_T_CELL_DIFFERENTIATION	0,7985612	1	0,1294429 -0,2567159	-0,806546	80

GOBP_ADP_METABOLIC_PROCESS	0,7985612	1	0,1294429	-0,2567208 -0,8065614	80

GOBP_REGULATION_OF_SKELETAL_MUSCLE_TISSUE_DEVELOPMENT	0,7226563	1	0,0959207	-0,3204082 -0,8065829	16

GOBP_LYSOSOMAL_TRANSPORT	0,8640777	1	0,1464162 -0,2401662 -0,8079491	116

GOBP_REGULATION_OF_LYMPHOCYTE_MIGRATION	0,7177033	1	0,1088201	-0,2799642 -0,8080259	41

GOBP_REGULATION_OF_DNA_DAMAGE_RESPONSE_SIGNAL_TRANSDUCTION_BY_P53_CLASS_MEDIATOR	0,7463415	1	0,1075544 -0,2990009 -0,8091301	33

GOBP_ACTIVATION_OF_NF_KAPPAB_INDUCING_KINASE_ACTIVITY	0,7148438	1	0,096563 -0,3216311 -0,8096615	16

GOBP_POSITIVE_REGULATION_OF_CYTOKINE_PRODUCTION_INVOLVED_IN_IMMUNE_RESPONSE	0,7566138	1	0,1119183 -0,2718524 -0,8100075	51

GOBP_MANNOSYLATION	0,7463415	1	0,1075544	-0,2995795 -0,8106958	33

GOBP_MYELOID_DENDRITIC_CELL_ACTIVATION	0,7193676	1	0,0968878	-0,314319 -0,8112308	19

GOBP_B_CELL_HOMEOSTASIS	0,7393162	1	0,0999277 -0,2979348	-0,811646	27

GOBP_RESPONSE_TO_ANTIBIOTIC	0,7203791	1	0,1079724	-0,280189 -0,8121909	42

GOBP_MIDDLE_EAR_MORPHOGENESIS	0,6848249	0,9963394	0,0988903 -0,3203286 -0,8132486	17

GOBP_POSITIVE_REGULATION_OF_INTRINSIC_APOPTOTIC_SIGNALING_PATHWAY	0,7362637	1	0,1162341 -0,2754945 -0,8149331	54

GOBP_REGULATION_OF_MEMBRANE_DEPOLARIZATION	0,7259615	1	0,1083943 -0,2858165 -0,8150153	38

GOBP_FOREBRAIN_NEURON_DEVELOPMENT	0,7035573	1	0,0982123	-0,3165152 -0,8168989	19

GOBP_REGULATION_OF_KERATINOCYTE_PROLIFERATION	0,7123288	1	0,1063233	-0,2995402 -0,8170263	30

GOBP_POSITIVE_REGULATION_OF_INTERLEUKIN_8_PRODUCTION	0,7128713	1	0,1114627	-0,2863165 -0,8177276	39

GOBP_REGULATION_OF_PROTEIN_LOCALIZATION_TO_CELL_SURFACE	0,7272727	1	0,1079724	-0,2951629 -0,8181989	35

GOBP_POSITIVE_REGULATION_OF_PRODUCTION_OF_MOLECULAR_MEDIATOR_OF_IMMUNE_RESPONSE	0,7903226	1	0,1388051 -0,2599247 -0,8182589	87

GOBP_POSITIVE_REGULATION_OF_INTRACELLULAR_TRANSPORT	0,826087	1	0,1864326	-0,2329562 -0,8182695	181

GOBP_CELL_COMMUNICATION_BY_ELECTRICAL_COUPLING	0,7051282	1	0,1028218	-0,2970831 -0,8183182	28

GOBP_REGULATION_OF_CHOLESTEROL_METABOLIC_PROCESS	0,7077626	1	0,1067299 -0,3013614	-0,821994	30

GOBP_POSITIVE_REGULATION_OF_INFLAMMATORY_RESPONSE	0,7716535	1	0,1388051 -0,2575743 -0,8239984	89

GOBP_VIRION_ASSEMBLY	0,722488	1	0,1083943 -0,2971957 -0,8258182	36

GOBP_VIRAL_PROTEIN_PROCESSING	0,7162791	1	0,1071402 -0,3027355	-0,826247	31

GOBP_ATRIAL_SEPTUM_DEVELOPMENT	0,6653696	0,9872205	0,1006334 -0,3260225 -0,8277041	17

GOBP_PROGRAMMED_NECROTIC_CELL_DEATH	0,6782178	0,9926234	0,1147507 -0,2904305 -0,8294774	39

GOBP_REGULATION_OF_OSTEOBLAST_PROLIFERATION	0,7063492	1	0,0982123	-0,3223333 -0,8305937	21

GOBP_REGULATION_OF_BONE_MINERALIZATION	0,7289157	1	0,1232572 -0,2742698 -0,8313504	64

GOBP_REGULATION_OF_DENDRITIC_SPINE_MORPHOGENESIS	0,6746411	0,9911514	0,1128434 -0,2880941 -0,8314904	41

GOBP_CELLULAR_RESPONSE_TO_AMYLOID_BETA	0,7177033	1	0,1088201	-0,2992471 -0,8315186	36

GOBP_REGULATION_OF_SMOOTH_MUSCLE_CONTRACTION	0,7110092	1	0,1067299	-0,283253 -0,8325152	45

GOBP_PROTEIN_DEMETHYLATION	0,6878049	0,9963394	0,1128434 -0,3079699 -0,8334011	33

GOBP_REGULATION_OF_P38MAPK_CASCADE	0,6985646	1	0,1105647	-0,29588 -0,8343711	37

GOBP_DOPAMINE_SECRETION	0,6976744	1	0,1088201 -0,3057301	-0,83442	31

GOBP_MEMBRANE_PROTEIN_ECTODOMAIN_PROTEOLYSIS	0,6777251	0,9923443	0,1119183 -0,2880779 -0,8350587	42

GOBP_CHONDROCYTE_PROLIFERATION	0,6984127	1	0,0988903	-0,3242906 -0,8356373	21

GOBP_PYRIDINE_CONTAINING_COMPOUND_METABOLIC_PROCESS	0,6883721	0,9963394	0,1096841	-0,306284 -0,8359319	31

GOBP_POSITIVE_REGULATION_OF_TYPE_I_INTERFERON_PRODUCTION	0,701087	1	0,118815 -0,2840062 -0,8365597	52

GOBP_MEMBRANE_DEPOLARIZATION	0,6809816	0,993958	0,1294429 -0,2702315 -0,8369025	69

GOBP_T_CELL_DIFFERENTIATION_IN_THYMUS	0,7228916	1	0,1238422	-0,2761213 -0,8369625	64

GOBP_CARBOHYDRATE_TRANSPORT	0,814433	1	0,1563124 -0,2505385	-0,837567	117

GOBP_POSITIVE_REGULATION_OF_TRANSCRIPTION_ELONGATION_FROM_RNA_POLYMERASE_II_PROMOTER	0,6730769	0,9908895	0,0992333 -0,3299363 -0,8381199	18

GOBP_NEGATIVE_REGULATION_OF_CELL_SUBSTRATE_ADHESION	0,6902174	0,9965975	0,1198878 -0,2848412 -0,8390193	52

GOBP_DECIDUALIZATION	0,6679842	0,9872205	0,1013507 -0,3253568 -0,8397184	19

GOBP_POSITIVE_REGULATION_OF_MONONUCLEAR_CELL_MIGRATION	0,6435644	0,9816504	0,1182875 -0,2953277	-0,843464	39

GOBP_POSITIVE_REGULATION_OF_SODIUM_ION_TRANSMEMBRANE_TRANSPORTER_ACTIVITY	0,6477273	0,9816504	0,1006334 -0,3381918 -0,8438327	15

GOBP_VIRAL_PROCESS	0,969697	1	0,2529611 -0,2198014 -0,8449995	350

GOBP_INTERLEUKIN_2_PRODUCTION	0,7009346	1	0,1088201	-0,2871704 -0,8465258	47

GOBP_NEGATIVE_REGULATION_OF_CARTILAGE_DEVELOPMENT	0,689243	0,9963394	0,0999277	-0,322812 -0,8465356	22

GOBP_REGULATION_OF_INTRINSIC_APOPTOTIC_SIGNALING_PATHWAY	0,8101266	1	0,175204 -0,2460295 -0,8467931	150

GOBP_POSITIVE_REGULATION_OF_MUSCLE_CELL_DIFFERENTIATION	0,6869159	0,9963394	0,1101223	-0,290832 -0,8492857	46

GOBP_GOLGI_TO_VACUOLE_TRANSPORT	0,692	0,9980207	0,0999277 -0,3314137 -0,8505359	20

GOBP_POSITIVE_REGULATION_OF_RELEASE_OF_CYTOCHROME_C_FROM_MITOCHONDRIA	0,6462882	0,9816504	0,1096841 -0,3180769 -0,8507418	25

GOBP_REGULATION_OF_ANTIGEN_RECEPTOR_MEDIATED_SIGNALING_PATHWAY	0,6869159	0,9963394	0,1101223	-0,291432 -0,8510378	46

GOBP_PHYSIOLOGICAL_CARDIAC_MUSCLE_HYPERTROPHY	0,6420233	0,9813188	0,1028218 -0,3354589 -0,8516612	17

GOBP_RESPONSE_TO_LITHIUM_ION	0,6445313	0,9816504	0,1028218	-0,33903 -0,8534608	16

GOBP_RETROGRADE_VESICLE_MEDIATED_TRANSPORT_GOLGI_TO_ENDOPLASMIC_RETICULUM	0,65625	0,9854242	0,1204334 -0,2881596 -0,8535128	50

GOBP_NEGATIVE_REGULATION_OF_ALPHA_BETA_T_CELL_ACTIVATION	0,6637168	0,9872205	0,1088201 -0,3136288 -0,8540575	29

GOBP_HAIR_CELL_DIFFERENTIATION	0,6585366	0,9860988	0,1157344 -0,3158751 -0,8547935	33

GOBP_RECEPTOR_INTERNALIZATION	0,768	1	0,1404062 -0,2647286 -0,8555514	95

GOBP_REGULATION_OF_NEUTROPHIL_CHEMOTAXIS	0,6573705	0,9855311	0,1028218 -0,3265895 -0,8564416	22

GOBP_CELLULAR_RESPONSE_TO_NUTRIENT	0,6333333	0,9808052	0,1167392 -0,3166017 -0,8571384	32

GOBP_INTERFERON_GAMMA_PRODUCTION	0,6441718	0,9816504	0,133555 -0,2768292 -0,8573356	69

GOBP_ANTIGEN_RECEPTOR_MEDIATED_SIGNALING_PATHWAY	0,7647059	1	0,1574029 -0,2531058 -0,8587161	121

GOBP_LYMPHOCYTE_MIGRATION	0,6626506	0,9872205	0,1301056 -0,2833432 -0,8588532	64

GOBP_MITOCHONDRIAL_CALCIUM_ION_TRANSMEMBRANE_TRANSPORT	0,6342412	0,9813188	0,1035763	-0,338553 -0,8595165	17

GOBP_POST_EMBRYONIC_ANIMAL_ORGAN_DEVELOPMENT	0,6367188	0,9813188	0,1035763 -0,3417313 -0,8602609	16

GOBP_CELL_CELL_RECOGNITION	0,6394231	0,9813188	0,1167392 -0,3024019	-0,862309	38

GOBP_BIOLOGICAL_PROCESS_INVOLVED_IN_SYMBIOTIC_INTERACTION	0,8360656	1	0,197822 -0,2412894 -0,8626189	212

GOBP_NEGATIVE_REGULATION_OF_RESPONSE_TO_ENDOPLASMIC_RETICULUM_STRESS	0,6328502	0,9808052	0,1177658 -0,2990036 -0,8628446	40

GOBP_POSITIVE_REGULATION_OF_UBIQUITIN_DEPENDENT_PROTEIN_CATABOLIC_PROCESS	0,7438017	1	0,1455161 -0,2608346 -0,8662876	101

GOBP_REGULATION_OF_POSTSYNAPTIC_MEMBRANE_NEUROTRANSMITTER_RECEPTOR_LEVELS	0,6304348	0,9808052	0,126254 -0,2942024 -0,8665933	52

GOBP_ARTERY_MORPHOGENESIS	0,5934066	0,9706649	0,1314576 -0,2907769 -0,8667812	56

GOBP_POSITIVE_REGULATION_OF_TRANSFORMING_GROWTH_FACTOR_BETA_PRODUCTION	0,6289063	0,9808052	0,104344 -0,3446051 -0,8674952	16

GOBP_POSITIVE_REGULATION_OF_CYTOKINE_PRODUCTION	0,8857143	1	0,2572065 -0,2293291 -0,8678072	325

GOBP_PROTEIN_LOCALIZATION_TO_ENDOPLASMIC_RETICULUM	0,61875	0,9804	0,1380222 -0,2788436 -0,8680443	70

GOBP_CARDIAC_MUSCLE_CELL_ACTION_POTENTIAL	0,6331361	0,9808052	0,1321473 -0,2864269 -0,8681769	63

GOBP_IMMUNE_RESPONSE_REGULATING_CELL_SURFACE_RECEPTOR_SIGNALING_PATHWAY	0,7710843	1	0,175204	-0,247237 -0,8684125	162

GOBP_NEGATIVE_REGULATION_OF_CALCIUM_ION_TRANSMEMBRANE_TRANSPORT	0,6047619	0,9711099	0,1198878 -0,3211872 -0,8695528	32

GOBP_POSITIVE_REGULATION_OF_NATURAL_KILLER_CELL_ACTIVATION	0,66	0,9871099	0,1028218 -0,3392848 -0,8707364	20

GOBP_RESPONSE_TO_NUTRIENT	0,7596154	1	0,1563124 -0,2628095 -0,8707679	113

GOBP_POSITIVE_REGULATION_OF_CALCIUM_ION_DEPENDENT_EXOCYTOSIS	0,6174242	0,9791739	0,1035763 -0,3491345 -0,8711361	15

GOBP_PROTEIN_TARGETING_TO_LYSOSOME	0,6239316	0,9808052	0,1105647 -0,3332787 -0,8737478	23

GOBP_TISSUE_REGENERATION	0,6453488	0,9816504	0,1294429 -0,2899956 -0,8738821	60

GOBP_LYMPHOCYTE_APOPTOTIC_PROCESS	0,5824176	0,9680138	0,1328463 -0,2932799 -0,8742424	56

GOBP_POSITIVE_REGULATION_OF_B_CELL_ACTIVATION	0,6395349	0,9813188	0,1301056 -0,2906168 -0,8757543	60

GOBP_POSITIVE_REGULATION_OF_JNK_CASCADE	0,6369863	0,9813188	0,1429011 -0,2796521 -0,8761733	78

GOBP_RELEASE_OF_SEQUESTERED_CALCIUM_ION_INTO_CYTOSOL_BY_ENDOPLASMIC_RETICULUM	0,6026201	0,9706649	0,1142665 -0,3280828 -0,8775042	25

GOBP_RESPONSE_TO_NICOTINE	0,5980861	0,9706649	0,1209851 -0,3113096 -0,8778822	37

GOBP_GRANULOCYTE_CHEMOTAXIS	0,5864198	0,9690661	0,1412251 -0,2851753 -0,8798565	68

GOBP_RECEPTOR_METABOLIC_PROCESS	0,7529412	1	0,175204 -0,2572307 -0,8801154	145

GOBP_LIPID_HOMEOSTASIS	0,6969697	1	0,1682382 -0,2579092 -0,8801993	125

GOBP_PROTEIN_MANNOSYLATION	0,6031746	0,9706649	0,1079724 -0,3417988 -0,8807528	21

GOBP_REGULATION_OF_LEUKOCYTE_APOPTOTIC_PROCESS	0,619883	0,9805897	0,1328463 -0,2926827	-0,88126	61

GOBP_MAGNESIUM_ION_TRANSPORT	0,5846154	0,9688575	0,1079724 -0,3470836 -0,8816783	18

GOBP_POSITIVE_REGULATION_OF_B_CELL_PROLIFERATION	0,5799087	0,9676999	0,1198878 -0,3237128 -0,8829595	30

GOBP_DEVELOPMENTAL_CELL_GROWTH	0,8125	1	0,195789 -0,2494302 -0,8837657	203

GOBP_DENDRITIC_SPINE_MORPHOGENESIS	0,5989011	0,9706649	0,1307771 -0,2974696 -0,8846904	55

GOBP_CELL_KILLING	0,6434109	0,9816504	0,1521449 -0,2753571 -0,8847654	90

GOBP_PERK_MEDIATED_UNFOLDED_PROTEIN_RESPONSE	0,5914397	0,9706649	0,1079724 -0,3491285 -0,8863654	17

GOBP_NEGATIVE_REGULATION_OF_CYTOKINE_PRODUCTION	0,734375	1	0,2065879	-0,2506939 -0,8865928	191

GOBP_NEGATIVE_REGULATION_OF_TRANSMEMBRANE_TRANSPORT	0,7165354	1	0,1446305	-0,2719872 -0,8888453	97

GOBP_APOPTOTIC_MITOCHONDRIAL_CHANGES	0,6692308	0,9872205	0,1482615 -0,2729513 -0,8889047	94

GOBP_RESPONSE_TO_AMYLOID_BETA	0,5645933	0,9620966	0,1250334 -0,3080881 -0,8891965	41

GOBP_POSITIVE_REGULATION_OF_PEPTIDASE_ACTIVITY	0,6835443	0,9963394	0,1918922 -0,2566392	-0,890485	159

GOBP_T_CELL_RECEPTOR_SIGNALING_PATHWAY	0,712	1	0,1464162 -0,2721576 -0,8920324	98

GOBP_T_CELL_MIGRATION	0,5990338	0,9706649	0,1215433 -0,3086961 -0,8922993	44

GOBP_REGULATION_OF_PEPTIDYL_SERINE_PHOSPHORYLATION	0,7280702	1	0,1521449 -0,2692395 -0,8926313	105

GOBP_CELL_MATRIX_ADHESION	0,734375	1	0,2065879 -0,2527457	-0,893849	191

GOBP_POSITIVE_REGULATION_OF_EXTRINSIC_APOPTOTIC_SIGNALING_PATHWAY	0,5942029	0,9706649	0,1221079 -0,3095223 -0,8946873	44

GOBP_NEUROMUSCULAR_SYNAPTIC_TRANSMISSION	0,5731225	0,9647049	0,1110115 -0,3467769 -0,8950018	19

GOBP_POSITIVE_REGULATION_OF_MITOCHONDRIAL_TRANSLATION	0,5833333	0,9680138	0,1071402 -0,3588267 -0,8953195	15

GOBP_NEGATIVE_REGULATION_OF_RHO_PROTEIN_SIGNAL_TRANSDUCTION	0,5873016	0,9690661	0,1096841 -0,3480742 -0,8969234	21

GOBP_GLUCOSE_IMPORT	0,5898876	0,969894	0,133555 -0,2999229 -0,8975416	57

GOBP_ENDOPLASMIC_RETICULUM_UNFOLDED_PROTEIN_RESPONSE	0,5493827	0,9535954	0,1464162	-0,291078 -0,8980683	68

GOBP_REGULATION_OF_ENDOTHELIAL_CELL_DEVELOPMENT	0,5820313	0,9680138	0,10925 -0,3572988	-0,89945	16

GOBP_SPERM_EGG_RECOGNITION	0,5782609	0,9664677	0,1167392 -0,3399857 -0,8996628	24

GOBP_ACTIVATION_OF_CYSTEINE_TYPE_ENDOPEPTIDASE_ACTIVITY_INVOLVED_IN_APOPTOTIC_PROCESS	0,5454545	0,9525767	0,1455161 -0,2945242 -0,9012019	65

GOBP_CELL_CELL_JUNCTION_ASSEMBLY	0,6868687	0,9963394	0,1695706 -0,2697762 -0,9019154	118

GOBP_REGULATION_OF_EXCRETION	0,5703125	0,9641936	0,1105647 -0,3585925 -0,9027065	16

GOBP_MONONUCLEAR_CELL_MIGRATION	0,627451	0,9808052	0,175204 -0,2707212 -0,9039612	115

GOBP_POSITIVE_REGULATION_OF_CELL_SUBSTRATE_ADHESION	0,664	0,9872205	0,1521449 -0,2785725 -0,9042826	96

GOBP_CELLULAR_RESPONSE_TO_CALCIUM_ION	0,5294118	0,9525709	0,154191 -0,2888281 -0,9046403	74

GOBP_THYROID_GLAND_DEVELOPMENT	0,5494071	0,9535954	0,1137873 -0,3506366 -0,9049635	19

GOBP_REGULATION_OF_ACTION_POTENTIAL	0,5607477	0,9613497	0,1238422 -0,3102395 -0,9059596	46

GOBP_GRANULOCYTE_MIGRATION	0,5658915	0,9621995	0,1631801 -0,2877704	-0,906849	85

GOBP_PROSTANOID_BIOSYNTHETIC_PROCESS	0,5697211	0,9641632	0,1119183 -0,3461107 -0,9076335	22

GOBP_NUCLEOTIDE_SUGAR_BIOSYNTHETIC_PROCESS	0,5564202	0,95855	0,1119183 -0,3579547 -0,9087734	17

GOBP_RHO_PROTEIN_SIGNAL_TRANSDUCTION	0,6568627	0,9855311	0,1709323 -0,2679351 -0,9090277	121

GOBP_POSITIVE_REGULATION_OF_T_CELL_PROLIFERATION	0,5092025	0,9479394	0,1521449 -0,2937609 -0,9097727	69

GOBP_ALPHA_BETA_T_CELL_ACTIVATION	0,5961538	0,9706649	0,1782199 -0,2751528 -0,9103069	112

GOBP_POSITIVE_REGULATION_OF_ACTIN_FILAMENT_POLYMERIZATION	0,5748792	0,9654878	0,1244342 -0,3150389 -0,9106334	44

GOBP_REGULATION_OF_NUCLEOCYTOPLASMIC_TRANSPORT	0,6456693	0,9816504	0,1531588 -0,2786986	-0,910778	97

GOBP_GRANULOCYTE_DIFFERENTIATION	0,5594714	0,9611038	0,1198878 -0,3397088 -0,9108962	26

GOBP_NEGATIVE_REGULATION_OF_SMALL_GTPASE_MEDIATED_SIGNAL_TRANSDUCTION	0,5347594	0,9525709	0,1372508 -0,3074027	-0,911506	53

GOBP_CHEMOKINE_PRODUCTION	0,5266272	0,9520965	0,1464162 -0,3010374 -0,9124621	63

GOBP_INNERVATION	0,5726496	0,9647049	0,1162341 -0,3480764 -0,9125425	23

GOBP_POSITIVE_REGULATION_OF_OXIDATIVE_STRESS_INDUCED_CELL_DEATH	0,5568182	0,95855	0,1101223 -0,3664205 -0,9142668	15

GOBP_REGULATION_OF_EXTENT_OF_CELL_GROWTH	0,6554622	0,9849234	0,1574029 -0,2752245 -0,9147977	102

GOBP_NEGATIVE_REGULATION_OF_LIPID_TRANSPORT	0,56	0,9613497	0,1133129 -0,3570044 -0,9162117	20

GOBP_MACROPHAGE_MIGRATION	0,5215311	0,9520965	0,1307771 -0,3250761 -0,9167032	37

GOBP_RESPONSE_TO_MURAMYL_DIPEPTIDE	0,5423077	0,9525709	0,1128434 -0,3616141 -0,9185895	18

GOBP_INTEGRATED_STRESS_RESPONSE_SIGNALING	0,5396825	0,9525709	0,11524 -0,3568994 -0,9196641	21

GOBP_CELL_SURFACE_RECEPTOR_SIGNALING_PATHWAY_INVOLVED_IN_HEART_DEVELOPMENT	0,5462555	0,9525767	0,1215433 -0,3430991 -0,9199871	26

GOBP_VENTRICULAR_CARDIAC_MUSCLE_TISSUE_DEVELOPMENT	0,5302326	0,9525709	0,1275053 -0,3184852 -0,9211143	43

GOBP_REGULATION_OF_T_HELPER_17_TYPE_IMMUNE_RESPONSE	0,536965	0,9525709	0,1142665	-0,362825	-0,921138	17

GOBP_INTERLEUKIN_12_PRODUCTION	0,5023923	0,9441675	0,133555 -0,3323095 -0,9233892	36

GOBP_NIK_NF_KAPPAB_SIGNALING	0,6315789	0,9808052	0,1644058 -0,2766802 -0,9234952	106

GOBP_REGULATION_OF_RECEPTOR_MEDIATED_ENDOCYTOSIS	0,5625	0,9614707	0,1644058 -0,2861371 -0,9242345	92

GOBP_INTERLEUKIN_10_PRODUCTION	0,5346535	0,9525709	0,1314576 -0,3237724 -0,9247026	39

GOBP_POSITIVE_REGULATION_OF_INTERLEUKIN_6_PRODUCTION	0,5182927	0,9512582	0,1501698 -0,3025836 -0,9252971	67

GOBP_POSITIVE_REGULATION_OF_GTPASE_ACTIVITY	0,6140351	0,9776358	0,24134 -0,2581583 -0,9259876	219

GOBP_POSITIVE_REGULATION_OF_LAMELLIPODIUM_ASSEMBLY	0,526087	0,9520965	0,1232572 -0,3516118 -0,9304276	24

GOBP_NEGATIVE_REGULATION_OF_PROTEOLYSIS	0,6041667	0,9711099	0,2663507 -0,2598544 -0,9316681	234

GOBP_EXTRACELLULAR_VESICLE_BIOGENESIS	0,526087	0,9520965	0,1232572 -0,3528008	-0,933574	24

GOBP_VITAMIN_METABOLIC_PROCESS	0,5426357	0,9525767	0,1669338	-0,296548	-0,93451	85

GOBP_NEGATIVE_REGULATION_OF_SMOOTH_MUSCLE_CELL_PROLIFERATION	0,5167464	0,9502772	0,1314576 -0,3242845 -0,9359421	41

GOBP_REGULATION_OF_BLOOD_CIRCULATION	0,6119403	0,9766609	0,222056 -0,2655451 -0,9365942	183

GOBP_DENDRITIC_CELL_MIGRATION	0,5214008	0,9520965	0,1162341 -0,3692652 -0,9374885	17

GOBP_N_ACETYLGLUCOSAMINE_METABOLIC_PROCESS	0,5115385	0,9479394	0,1167392 -0,3692634 -0,9380207	18

GOBP_MYELOID_LEUKOCYTE_CYTOKINE_PRODUCTION	0,5283843	0,9525709	0,1232572 -0,3513633 -0,9397712	25

GOBP_REGULATION_OF_T_CELL_RECEPTOR_SIGNALING_PATHWAY	0,4904762	0,9401628	0,135002 -0,3415365 -0,9401275	34

GOBP_ACTIN_FILAMENT_BASED_TRANSPORT	0,5115385	0,9479394	0,1167392 -0,3709335 -0,9422631	18

GOBP_REGULATION_OF_LEUKOCYTE_PROLIFERATION	0,5802469	0,9676999	0,2065879 -0,2689414 -0,9424525	163

GOBP_NEGATIVE_REGULATION_OF_DEVELOPMENTAL_GROWTH	0,544	0,9525767	0,1695706 -0,2876105 -0,9426814	98

GOBP_REGULATION_OF_EPITHELIAL_CELL_DIFFERENTIATION	0,5631068	0,9616328	0,1847065 -0,2803302	-0,943066	116

GOBP_CARDIAC_ATRIUM_DEVELOPMENT	0,5110132	0,9479394	0,126254 -0,3518995 -0,9435845	26

GOBP_NEGATIVE_REGULATION_OF_CELL_MATRIX_ADHESION	0,4809524	0,9326074	0,1364904 -0,3491391 -0,9452272	32

GOBP_SUBSTRATE_ADHESION_DEPENDENT_CELL_SPREADING	0,5203252	0,9520965	0,175204 -0,2850403 -0,9452517	100

GOBP_CHEMICAL_HOMEOSTASIS_WITHIN_A_TISSUE	0,4921875	0,9415216	0,1204334 -0,3755235 -0,9453281	16

GOBP_TRIGLYCERIDE_METABOLIC_PROCESS	0,4814815	0,9326074	0,1574029 -0,3065271 -0,9457338	68

GOBP_REGULATION_OF_CELLULAR_RESPONSE_TO_INSULIN_STIMULUS	0,510989	0,9479394	0,1429011 -0,3197438 -0,9458255	54

GOBP_REGULATION_OF_ACTIN_FILAMENT_ORGANIZATION	0,54	0,9525709	0,2765006 -0,2616384 -0,9461658	235

GOBP_SMOOTH_MUSCLE_TISSUE_DEVELOPMENT	0,5019763	0,9439459	0,1198878 -0,3667543 -0,9465618	19

GOBP_LIPID_DROPLET_ORGANIZATION	0,5	0,9430221	0,1204334 -0,3676877 -0,9474637	21

GOBP_ENDOTHELIAL_CELL_APOPTOTIC_PROCESS	0,4951923	0,941623	0,135002 -0,3323748 -0,9477779	38

GOBP_NEGATIVE_REGULATION_OF_NEURON_PROJECTION_DEVELOPMENT	0,54	0,9525709	0,1918922 -0,2794209 -0,9482081	128

GOBP_ACTIVATED_T_CELL_PROLIFERATION	0,5073171	0,9477823	0,1342735 -0,3504022 -0,9482279	33

GOBP_NEGATIVE_REGULATION_OF_LEUKOCYTE_MIGRATION	0,4761905	0,9291869	0,1372508 -0,3503155 -0,9484121	32

GOBP_MULTI_MULTICELLULAR_ORGANISM_PROCESS	0,5454545	0,9525767	0,2192503 -0,2747646 -0,9485851	155

GOBP_CELLULAR_RESPONSE_TO_CAMP	0,492823	0,941623	0,135002	-0,328791 -0,9489488	41

GOBP_REGULATION_OF_CELLULAR_COMPONENT_SIZE	0,686716	0,9963394	0,1439026	-0,251903 -0,9494315	318

GOBP_CYTOPLASMIC_PATTERN_RECOGNITION_RECEPTOR_SIGNALING_PATHWAY	0,5054945	0,9472034	0,143759 -0,3210185 -0,9495964	54

GOBP_MUCOPOLYSACCHARIDE_METABOLIC_PROCESS	0,4605263	0,9232643	0,1669338 -0,3020927 -0,9509133	75

GOBP_BIOLOGICAL_PROCESS_INVOLVED_IN_INTERACTION_WITH_HOST	0,5194805	0,9515789	0,2249661 -0,2773908	-0,950915	151

GOBP_APICAL_JUNCTION_ASSEMBLY	0,4662921	0,9254409	0,1521449 -0,3178214 -0,9511043	57

GOBP_INTERLEUKIN_17_PRODUCTION	0,534188	0,9525709	0,1209851 -0,3628659 -0,9513158	23

GOBP_POSITIVE_REGULATION_OF_LYMPHOCYTE_MIGRATION	0,5099602	0,9479394	0,1193484 -0,3627829 -0,9513543	22

GOBP_PLASMA_MEMBRANE_REPAIR	0,5099602	0,9479394	0,1193484 -0,3629939 -0,9519076	22

GOBP_INTEGRIN_MEDIATED_SIGNALING_PATHWAY	0,4765625	0,9291869	0,1797823 -0,2947374 -0,9520136	92

GOBP_POSITIVE_REGULATION_OF_SIGNAL_TRANSDUCTION_BY_P53_CLASS_MEDIATOR	0,5	0,9430221	0,1256399 -0,3497429 -0,9527837	27

GOBP_MORPHOGENESIS_OF_AN_EPITHELIAL_FOLD	0,4980237	0,9416872	0,1204334 -0,3695146 -0,9536858	19

GOBP_REGULATION_OF_NEUROTRANSMITTER_RECEPTOR_ACTIVITY	0,4739583	0,9291869	0,1446305 -0,3248662 -0,9539087	49

GOBP_PROTEIN_PALMITOYLATION	0,4761905	0,9291869	0,1372508 -0,3525154 -0,9543677	32

GOBP_RENAL_SYSTEM_PROCESS	0,4657534	0,9253885	0,1695706 -0,3049566 -0,9554542	78

GOBP_REGULATION_OF_PHOSPHOLIPID_METABOLIC_PROCESS	0,4513274	0,9170563	0,1357409 -0,3518354 -0,9580999	29

GOBP_VENTRICULAR_CARDIAC_MUSCLE_TISSUE_MORPHOGENESIS	0,4688995	0,9267843	0,1388051 -0,3450727 -0,9588543	36

GOBP_ADAPTIVE_THERMOGENESIS	0,5360825	0,9525709	0,195789 -0,2839244 -0,9599806	131

GOBP_POSITIVE_REGULATION_OF_ACTIN_FILAMENT_BUNDLE_ASSEMBLY	0,4761905	0,9291869	0,1455161 -0,3225343 -0,9610186	51

GOBP_REGULATION_OF_MONONUCLEAR_CELL_MIGRATION	0,4532374	0,917415	0,1766943 -0,3062116 -0,9620509	80

GOBP_CARDIAC_VENTRICLE_MORPHOGENESIS	0,478022	0,9291869	0,1482615 -0,3253518 -0,9624146	54

GOBP_REGULATION_OF_MACROPHAGE_DIFFERENTIATION	0,4769231	0,9291869	0,1215433 -0,3792512 -0,9633922	18

GOBP_PROTEIN_EXIT_FROM_ENDOPLASMIC_RETICULUM	0,4660194	0,9253885	0,1404062 -0,3260555 -0,9634733	48

GOBP_NEGATIVE_REGULATION_OF_HYDROLASE_ACTIVITY	0,5569642	0,95855	0,1439026 -0,2689622 -0,9635194	247

GOBP_MITOCHONDRIAL_DEPOLARIZATION	0,4940711	0,941623	0,1209851 -0,3735169 -0,9640156	19

GOBP_TOLERANCE_INDUCTION	0,4980545	0,9416872	0,1193484 -0,3797659 -0,9641476	17

GOBP_POSITIVE_REGULATION_OF_FAT_CELL_DIFFERENTIATION	0,4382022	0,9105785	0,1574029 -0,3214217 -0,9649172	58

GOBP_POSITIVE_REGULATION_OF_LEUKOCYTE_PROLIFERATION	0,4684685	0,9267843	0,195789 -0,2911724 -0,9652427	109

GOBP_ANDROGEN_RECEPTOR_SIGNALING_PATHWAY	0,480198	0,9318719	0,1395997	-0,338237 -0,9660141	39

GOBP_NEUTRAL_LIPID_BIOSYNTHETIC_PROCESS	0,4641148	0,9252407	0,1395997 -0,3477502 -0,9662942	36

GOBP_POSITIVE_REGULATION_OF_TUMOR_NECROSIS_FACTOR_SUPERFAMILY_CYTOKINE_PRODUCTION	0,4487179	0,9158429	0,1669338 -0,3111867	-0,967156	71

GOBP_TEMPERATURE_HOMEOSTASIS	0,4941176	0,941623	0,2192503 -0,2832301 -0,9690722	145

GOBP_RESPONSE_TO_STARVATION	0,5064935	0,9476574	0,2279872 -0,2728337 -0,9691877	168

GOBP_TRANSEPITHELIAL_TRANSPORT	0,4700855	0,9279845	0,1301056 -0,3559983	-0,969825	27

GOBP_SRP_DEPENDENT_COTRANSLATIONAL_PROTEIN_TARGETING_TO_MEMBRANE	0,486166	0,9382609	0,1221079 -0,3759526 -0,9703018	19

GOBP_CELL_CELL_ADHESION_MEDIATED_BY_CADHERIN	0,46	0,9231849	0,1268757 -0,3783072 -0,9708828	20

GOBP_TIGHT_JUNCTION_ORGANIZATION	0,4619883	0,9242226	0,1563124 -0,3226058 -0,9713576	61

GOBP_INTERLEUKIN_6_PRODUCTION	0,4552846	0,9201205	0,1882041 -0,2932202 -0,9723776	100

GOBP_FLUID_TRANSPORT	0,4782609	0,9291869	0,1232572 -0,3783723 -0,9765469	19

GOBP_ARF_PROTEIN_SIGNAL_TRANSDUCTION	0,4615385	0,9242226	0,1238422 -0,3848181 -0,9775335	18

GOBP_REGULATION_OF_HEART_RATE_BY_CARDIAC_CONDUCTION	0,452381	0,9172128	0,1412251	-0,361177 -0,9778175	32

GOBP_LIPID_STORAGE	0,439759	0,9117962	0,1631801	-0,320688 -0,9780018	66

GOBP_PROTEIN_KINASE_C_ACTIVATING_G_PROTEIN_COUPLED_RECEPTOR_SIGNALING_PATHWAY	0,4782609	0,9291869	0,1232572 -0,3789365 -0,9780031	19

GOBP_MACROPHAGE_CHEMOTAXIS	0,4273504	0,9065813	0,1372508 -0,3555687 -0,9794174	28

GOBP_LEUKOCYTE_CHEMOTAXIS	0,46875	0,9267843	0,2114002 -0,2887044 -0,9799474	134

GOBP_GASTRULATION	0,474359	0,9291869	0,2343926 -0,2851014 -0,9805673	153

GOBP_ENTRY_INTO_HOST	0,4466019	0,9158429	0,208955 -0,2915185 -0,9807048	116

GOBP_REGULATION_OF_LEUKOCYTE_MIGRATION	0,4588235	0,9223114	0,2279872 -0,2867307 -0,9810498	145

GOBP_NEGATIVE_REGULATION_OF_INTERLEUKIN_6_PRODUCTION	0,4230769	0,9043534	0,1380222 -0,3565102 -0,9820108	28

GOBP_ANTIBACTERIAL_HUMORAL_RESPONSE	0,4563492	0,9210223	0,1268757	-0,381826 -0,9838955	21

GOBP_REGULATION_OF_RESPONSE_TO_ENDOPLASMIC_RETICULUM_STRESS	0,4183007	0,9011213	0,175204 -0,3145513 -0,9852082	74

GOBP_NEGATIVE_REGULATION_OF_GLUCONEOGENESIS	0,4431818	0,9132893	0,1256399 -0,3951885 -0,9860468	15

GOBP_REGULATION_OF_CELL_SUBSTRATE_ADHESION	0,5	0,9430221	0,2450418 -0,2827298 -0,9869531	173

GOBP_POSITIVE_REGULATION_OF_PEPTIDYL_TYROSINE_PHOSPHORYLATION	0,45	0,9158429	0,2114002 -0,2909893 -0,9874655	128

GOBP_NEGATIVE_REGULATION_OF_POTASSIUM_ION_TRANSMEMBRANE_TRANSPORT	0,459144	0,922462	0,1250334 -0,3891731 -0,9880305	17

GOBP_REGULATION_OF_NIK_NF_KAPPAB_SIGNALING	0,4444444	0,9145536	0,175204 -0,3103206 -0,9882894	81

GOBP_NEGATIVE_REGULATION_OF_CELLULAR_RESPONSE_TO_INSULIN_STIMULUS	0,452381	0,9172128	0,1412251 -0,3650984 -0,9884339	32

GOBP_POSITIVE_REGULATION_OF_INTERLEUKIN_2_PRODUCTION	0,4145299	0,9	0,1395997 -0,3588565 -0,9884736	28

GOBP_NEGATIVE_REGULATION_OF_TYPE_I_INTERFERON_MEDIATED_SIGNALING_PATHWAY	0,45	0,9158429	0,1256399 -0,3893307 -0,9889966	18

GOBP_NUCLEAR_MEMBRANE_REASSEMBLY	0,4487179	0,9158429	0,133555 -0,3774998	-0,989681	23

GOBP_NEGATIVE_REGULATION_OF_T_CELL_APOPTOTIC_PROCESS	0,4296875	0,9084133	0,1301056 -0,3931544 -0,9897115	16

GOBP_RESPONSE_TO_OSMOTIC_STRESS	0,382716	0,8862233	0,1782199 -0,3214307 -0,9917161	68

GOBP_NEGATIVE_REGULATION_OF_T_CELL_PROLIFERATION	0,4492754	0,9158429	0,1429011	-0,34375 -0,9919708	40

GOBP_REGULATION_OF_PROTEIN_TYROSINE_KINASE_ACTIVITY	0,4078947	0,8971086	0,1782199 -0,3153103 -0,9925189	75

GOBP_NEGATIVE_REGULATION_OF_SMOOTH_MUSCLE_CELL_MIGRATION	0,4342629	0,9090275	0,1307771 -0,3787138 -0,9931313	22

GOBP_TRANSLATIONAL_TERMINATION	0,4461538	0,9158429	0,126254 -0,3909703 -0,9931614	18

GOBP_REGULATION_OF_VIRAL_INDUCED_CYTOPLASMIC_PATTERN_RECOGNITION_RECEPTOR_SIGNALING_PATHWAY	0,4285714	0,9075254	0,1314576 -0,3856688 -0,9937978	21

GOBP_INNER_EAR_AUDITORY_RECEPTOR_CELL_DIFFERENTIATION	0,4059829	0,8971086	0,1412251	-0,360812 -0,9938602	28

GOBP_POSITIVE_REGULATION_OF_BLOOD_PRESSURE	0,4262948	0,9062536	0,1321473 -0,3802929 -0,9972721	22

GOBP_NEGATIVE_REGULATION_OF_CELL_KILLING	0,4179688	0,9009229	0,1321473 -0,3962053 -0,9973916	16

GOBP_RENAL_FILTRATION	0,424	0,9043534	0,1328463 -0,3891119 -0,9986119	20

GOBP_POSITIVE_REGULATION_OF_MYELOID_LEUKOCYTE_CYTOKINE_PRODUCTION_INVOLVED_IN_IMMUNE_RESPONSE	0,4426877	0,9132893	0,1287887 -0,3871732 -0,9992613	19

GOBP_BINDING_OF_SPERM_TO_ZONA_PELLUCIDA	0,4103586	0,8974438	0,135002 -0,3813839 -1,0001333	22

GOBP_NEURON_PROJECTION_EXTENSION	0,4358974	0,9090275	0,2450418 -0,2885303 -1,0001629	161

GOBP_NEGATIVE_REGULATION_OF_PEPTIDYL_SERINE_PHOSPHORYLATION	0,4347826	0,9090275	0,1372508 -0,3779754 -1,0001905	24

GOBP_INTERLEUKIN_1_PRODUCTION	0,3841463	0,8862233	0,1766943	-0,327123 -1,0003384	67

GOBP_N_TERMINAL_PROTEIN_AMINO_ACID_MODIFICATION	0,4017094	0,8935596	0,1420566 -0,3690474 -1,0053737	27

GOBP_COPII_COATED_VESICLE_CARGO_LOADING	0,4280303	0,9075112	0,1281429 -0,4033652 -1,0064489	15

GOBP_ACTIN_FILAMENT_DEPOLYMERIZATION	0,3641304	0,880326	0,1709323 -0,3417478 -1,0066414	52

GOBP_REGULATION_OF_NATURAL_KILLER_CELL_ACTIVATION	0,3931624	0,8896911	0,143759 -0,3698076 -1,0074448	27

GOBP_REGULATION_OF_CELL_MATRIX_ADHESION	0,4214876	0,9043534	0,197822 -0,3035515 -1,0081594	101

GOBP_REGULATION_OF_TRIGLYCERIDE_METABOLIC_PROCESS	0,3931624	0,8896911	0,143759 -0,3702463 -1,0086399	27

GOBP_POSITIVE_REGULATION_OF_MUSCLE_CONTRACTION	0,4142857	0,9	0,1482615 -0,3731169 -1,0101424	32

GOBP_EPITHELIAL_CELL_APOPTOTIC_PROCESS	0,4178082	0,9009229	0,1797823 -0,3224532 -1,0102724	78

GOBP_REGULATION_OF_POTASSIUM_ION_TRANSPORT	0,4041096	0,8958579	0,1830239 -0,3227919 -1,0113338	78

GOBP_RELEASE_OF_CYTOCHROME_C_FROM_MITOCHONDRIA	0,3529412	0,877063	0,1723243 -0,3413053 -1,0120334	53

GOBP_REGULATION_OF_EPITHELIAL_CELL_APOPTOTIC_PROCESS	0,4011628	0,8935596	0,1682382 -0,3360798 -1,0127537	60

GOBP_CELL_COMMUNICATION_INVOLVED_IN_CARDIAC_CONDUCTION	0,3899083	0,8869191	0,1501698 -0,3447402 -1,0132334	45

GOBP_CELL_CELL_JUNCTION_MAINTENANCE	0,4241245	0,9043534	0,1307771 -0,3994483	-1,014117	17

GOBP_REGULATION_OF_RESPONSE_TO_TUMOR_CELL	0,4241245	0,9043534	0,1307771 -0,3996974 -1,0147494	17

GOBP_IRON_ION_TRANSPORT	0,3980583	0,8910522	0,1531588 -0,3438969 -1,0161934	48

GOBP_POSITIVE_REGULATION_OF_MAP_KINASE_ACTIVITY	0,4076923	0,8971086	0,1938133 -0,3120774 -1,0163246	94

GOBP_CERAMIDE_CATABOLIC_PROCESS	0,4241245	0,9043534	0,1307771 -0,4010863 -1,0182756	17

GOBP_ASTROCYTE_DEVELOPMENT	0,3906977	0,8869191	0,1511488 -0,3731152 -1,0183323	31

GOBP_REGULATION_OF_HETEROTYPIC_CELL_CELL_ADHESION	0,4090909	0,8972297	0,1314576	-0,408284 -1,0187219	15

GOBP_AMELOGENESIS	0,3945313	0,8896911	0,1364904 -0,4049495 -1,0194039	16

GOBP_REGULATION_OF_PROTEIN_LOCALIZATION_TO_SYNAPSE	0,4150198	0,9	0,133555 -0,3951901 -1,0199522	19

GOBP_POSITIVE_REGULATION_OF_STRIATED_MUSCLE_CELL_DIFFERENTIATION	0,3835616	0,8862233	0,1511488 -0,3742915	-1,020918	30

GOBP_NEGATIVE_REGULATION_OF_ENDOTHELIAL_CELL_APOPTOTIC_PROCESS	0,3864542	0,8862233	0,1395997 -0,3893539 -1,0210337	22

GOBP_PROTEIN_IMPORT_INTO_MITOCHONDRIAL_MATRIX	0,4153846	0,9	0,1314576 -0,4020051 -1,0211927	18

GOBP_LYMPHOCYTE_CHEMOTAXIS	0,3826087	0,8862233	0,1473312 -0,3861987 -1,0219508	24

GOBP_CYTOPLASMIC_PATTERN_RECOGNITION_RECEPTOR_SIGNALING_PATHWAY_IN_RESPONSE_TO_VIRUS	0,3813953	0,8862233	0,1531588 -0,3746546 -1,0225337	31

GOBP_ACTIN_FILAMENT_BUNDLE_ORGANIZATION	0,3571429	0,877063	0,2616635 -0,3001536 -1,0227086	143

GOBP_CELLULAR_RESPONSE_TO_STARVATION	0,3571429	0,877063	0,2616635 -0,3008358 -1,0250331	143

GOBP_EPITHELIAL_CELL_MORPHOGENESIS	0,373913	0,8832291	0,1492075 -0,3874482 -1,0252573	24

GOBP_REGULATION_OF_LIPID_STORAGE	0,3861386	0,8862233	0,1574029 -0,3590493 -1,0254545	39

GOBP_REGULATION_OF_LIPOPROTEIN_METABOLIC_PROCESS	0,390625	0,8869191	0,1372508 -0,4080686 -1,0272558	16

GOBP_REGULATION_OF_MACROPHAGE_MIGRATION	0,3846154	0,8862233	0,1455161 -0,3772495 -1,0277182	27

GOBP_INTRA_GOLGI_VESICLE_MEDIATED_TRANSPORT	0,4	0,8929924	0,1511488 -0,3798414 -1,0283477	32

GOBP_REGULATION_OF_ACTIN_CYTOSKELETON_REORGANIZATION	0,4047619	0,8961052	0,1501698 -0,3739675 -1,0293984	34

GOBP_NATURAL_KILLER_CELL_ACTIVATION	0,3516484	0,877063	0,175204	-0,348013 -1,0294479	54

GOBP_COTRANSLATIONAL_PROTEIN_TARGETING_TO_MEMBRANE	0,3624454	0,8797754	0,1521449 -0,3850099 -1,0297637	25

GOBP_RENAL_VESICLE_DEVELOPMENT	0,4015152	0,8935596	0,1328463 -0,4135432 -1,0318442	15

GOBP_RESPONSE_TO_EXTRACELLULAR_STIMULUS	0,2028984	0,7744131	0,2492466 -0,2687293 -1,0327594	375

GOBP_INOSITOL_PHOSPHATE_MEDIATED_SIGNALING	0,3691589	0,8814579	0,1563124 -0,3504552 -1,0330778	47

GOBP_EXTRACELLULAR_MATRIX_ASSEMBLY	0,3663366	0,8810414	0,1619789 -0,3621732 -1,0343765	39

GOBP_EPITHELIAL_CELL_DIFFERENTIATION	0,2586661	0,8179127	0,203509 -0,2619658 -1,0347453	480

GOBP_CELL_ADHESION_MEDIATED_BY_INTEGRIN	0,3353659	0,8693282	0,1900233 -0,3387583 -1,0359189	67

GOBP_POSITIVE_REGULATION_OF_LEUKOCYTE_MIGRATION	0,3461538	0,8738784	0,2114002 -0,3181031	-1,035948	94

GOBP_REGULATION_OF_SIGNALING_RECEPTOR_ACTIVITY	0,3877551	0,8862233	0,2311267 -0,3088106 -1,0363998	119

GOBP_OLIGOPEPTIDE_TRANSPORT	0,4015152	0,8935596	0,1328463 -0,4155174 -1,0367701	15

GOBP_MYOBLAST_FUSION	0,3744292	0,8832291	0,1531588 -0,3801352 -1,0368575	30

GOBP_MITOCHONDRIAL_CALCIUM_ION_HOMEOSTASIS	0,3665339	0,8810414	0,143759 -0,3954479 -1,0370144	22

GOBP_APOPTOTIC_PROCESS_INVOLVED_IN_MORPHOGENESIS	0,3760684	0,8849204	0,1473312 -0,3964401 -1,0393363	23

GOBP_REGULATION_OF_ANATOMICAL_STRUCTURE_SIZE	0,201317	0,7744131	0,2492466 -0,2711217 -1,0404331	412

GOBP_HEMOSTASIS	0,373494	0,8832291	0,2572065 -0,2965288 -1,0415488	162

GOBP_ENDOTHELIUM_DEVELOPMENT	0,375	0,8838614	0,2279872 -0,3148934 -1,0417838	112

GOBP_HISTONE_H3_ACETYLATION	0,3461538	0,8738784	0,1918922 -0,3352322 -1,0418884	71

GOBP_GLYCEROLIPID_CATABOLIC_PROCESS	0,352657	0,877063	0,1631801 -0,3611275 -1,0438544	44

GOBP_POSITIVE_REGULATION_OF_INTERFERON_BETA_PRODUCTION	0,3636364	0,8799278	0,1596467 -0,3760292	-1,044873	36

GOBP_O_GLYCAN_PROCESSING	0,354067	0,877063	0,1619789	-0,370695 -1,0453469	37

GOBP_CELLULAR_RESPONSE_TO_EXTRACELLULAR_STIMULUS	0,3770981	0,8849204	0,203509 -0,2930801 -1,0477723	212

GOBP_NEUROMUSCULAR_JUNCTION_DEVELOPMENT	0,3429952	0,8738784	0,1656567 -0,3630897	-1,04778	40

GOBP_AXON_EXTENSION	0,3557692	0,877063	0,2343926 -0,3173649 -1,0515264	113

GOBP_GRANULOCYTE_ACTIVATION	0,3632479	0,8798983	0,1501698 -0,3860901 -1,0518021	27

| GOBP_CARDIAC_MUSCLE_CELL_ACTION_POTENTIAL_INVOLVED_IN_CONTRACTION | 0,3488372 0,8764774 0,1608014 -0,3641254 -1,0531139 | 43 |
| --- | --- | --- |
| GOBP_ENDOPLASMIC_RETICULUM_TO_GOLGI_VESICLE_MEDIATED_TRANSPORT | 0,31 0,8581768 0,2572065 -0,311954 -1,0536688 | 123 |
| GOBP_HEART_MORPHOGENESIS | 0,3265339 0,8660673 0,203509 -0,2941094 -1,0554248 | 208 |
| GOBP_POSITIVE_REGULATION_OF_ANTIGEN_RECEPTOR_MEDIATED_SIGNALING_PATHWAY | 0,3675889 0,881322 0,1429011 -0,4089856 -1,0555572 | 19 |
| GOBP_REGULATION_OF_ACTIN_FILAMENT_BASED_PROCESS | 0,2742816 0,8289344 0,203509 -0,2790796 -1,0559768 | 339 |
| GOBP_NEGATIVE_REGULATION_OF_RECEPTOR_INTERNALIZATION | 0,3712121 0,8831254 0,1388051 -0,4240154 -1,0579738 | 15 |
| GOBP_DETECTION_OF_STIMULUS_INVOLVED_IN_SENSORY_PERCEPTION_OF_PAIN | 0,3333333 0,8660673 0,1511488 -0,4109702 -1,0589946 | 21 |
| GOBP_REGULATION_OF_RHO_PROTEIN_SIGNAL_TRANSDUCTION | 0,3269231 0,8660673 0,197822 -0,338891 -1,0598465 | 73 |
| GOBP_PLASMA_MEMBRANE_ORGANIZATION | 0,3069307 0,8571747 0,2572065 -0,3134856 -1,0601569 | 120 |
| GOBP_RESPONSE_TO_COLD | 0,3333333 0,8660673 0,1682382 -0,3668361 -1,0603554 | 44 |
| GOBP_NEGATIVE_REGULATION_OF_PROGRAMMED_NECROTIC_CELL_DEATH | 0,3671875 0,881322 0,1420566 -0,4213065 -1,0605804 | 16 |
| GOBP_RESPONSE_TO_DOPAMINE | 0,2967033 0,8470473 0,1918922 -0,3568859 -1,0613978 | 55 |
| GOBP_NEGATIVE_REGULATION_OF_CD4_POSITIVE_ALPHA_BETA_T_CELL_ACTIVATION | 0,3247863 0,8660673 0,1596467 -0,4053915 -1,0628042 | 23 |
| GOBP_REGULATION_OF_WOUND_HEALING | 0,296 0,8469703 0,2343926 -0,3292669 -1,0641267 | 95 |
| GOBP_REGULATION_OF_PEPTIDYL_TYROSINE_PHOSPHORYLATION | 0,2984493 0,8492127 0,203509 -0,3023884 -1,0650877 | 185 |
| GOBP_POSITIVE_REGULATION_OF_PROTEIN_ACETYLATION | 0,3459716 0,8738784 0,1631801 -0,3678634 -1,0663352 | 42 |
| GOBP_HYPOTHALAMUS_DEVELOPMENT | 0,3674242 0,881322 0,1395997 -0,4279177 -1,0677103 | 15 |
| GOBP_NEURON_PROJECTION_ORGANIZATION | 0,3546099 0,877063 0,1999152 -0,3349139 -1,0681461 | 82 |
| GOBP_RIG_I_SIGNALING_PATHWAY | 0,3537118 0,877063 0,154191 -0,3994922 -1,0684987 | 25 |
| GOBP_MESODERM_MORPHOGENESIS | 0,2857143 0,8386161 0,195789 -0,3616473 -1,0697793 | 54 |
| GOBP_REGULATION_OF_ACTIN_FILAMENT_BUNDLE_ASSEMBLY | 0,3178295 0,8622448 0,222056 -0,3333262 -1,0710291 | 90 |
| GOBP_SKELETAL_MUSCLE_TISSUE_REGENERATION | 0,3463415 0,8738784 0,1656567 -0,3959183 -1,0713994 | 33 |
| GOBP_TUMOR_NECROSIS_FACTOR_SUPERFAMILY_CYTOKINE_PRODUCTION | 0,3365385 0,8693282 0,24134 -0,3238809 -1,0715177 | 112 |
| GOBP_REGULATION_OF_MITOCHONDRIAL_MEMBRANE_POTENTIAL | 0,2909091 0,8451947 0,2042948 -0,3502145 -1,0716062 | 65 |
| GOBP_POSITIVE_REGULATION_OF_PROTEIN_TYROSINE_KINASE_ACTIVITY | 0,3364929 0,8693282 0,1656567 -0,3698641 -1,0721346 | 42 |
| GOBP_PEPTIDYL_PROLINE_HYDROXYLATION | 0,3598485 0,8774738 0,1412251 -0,4298995 -1,0726554 | 15 |
| GOBP_COPPER_ION_TRANSPORT | 0,3598485 0,8774738 0,1412251 -0,4300203 -1,0729567 | 15 |
| GOBP_MYELOID_CELL_DEVELOPMENT | 0,2802198 0,8352792 0,197822 -0,3631856 -1,0743297 | 54 |
| GOBP_CARDIAC_MUSCLE_CELL_CONTRACTION | 0,2923977 0,8455459 0,1999152 -0,3567476 -1,0744225 | 62 |
| GOBP_ENDOCRINE_PROCESS | 0,2967033 0,8470473 0,1918922 -0,3606325 -1,0750149 | 56 |
| GOBP_REGULATION_OF_EXOSOMAL_SECRETION | 0,35 0,8764774 0,1446305 -0,4232876 -1,0752556 | 18 |
| GOBP_SARCOPLASMIC_RETICULUM_CALCIUM_ION_TRANSPORT | 0,3318584 0,8660673 0,1608014 -0,3951515 -1,0760561 | 29 |
| GOBP_RECEPTOR_MEDIATED_ENDOCYTOSIS | 0,1415523 0,6791978 0,2492466 -0,3005089 -1,0771312 | 206 |
| GOBP_WOUND_HEALING_SPREADING_OF_EPIDERMAL_CELLS | 0,35 0,8764774 0,1446305 -0,4243378 -1,0779231 | 18 |
| GOBP_PROTEIN_LOCALIZATION_TO_ENDOSOME | 0,3217391 0,8660673 0,1619789 -0,4088576 -1,0819102 | 24 |
| GOBP_REGULATION_OF_MACROPHAGE_CHEMOTAXIS | 0,3174603 0,8622448 0,155242 -0,4200722 -1,082449 | 21 |
| GOBP_POSITIVE_REGULATION_OF_CHEMOKINE_PRODUCTION | 0,3119266 0,8585408 0,1695706 -0,3683192 -1,0825351 | 45 |
| GOBP_NEGATIVE_REGULATION_OF_BLOOD_CIRCULATION | 0,3478261 0,8764774 0,1473312 -0,4194666 -1,0826077 | 19 |
| GOBP_REGULATION_OF_SYSTEMIC_ARTERIAL_BLOOD_PRESSURE_BY_HORMONE | 0,3076923 0,857788 0,1644058 -0,4131119 -1,0830443 | 23 |
| GOBP_REGULATION_OF_MEMBRANE_LIPID_DISTRIBUTION | 0,2802198 0,8352792 0,197822 -0,3644704 -1,0839542 | 55 |
| GOBP_POSITIVE_REGULATION_OF_NIK_NF_KAPPAB_SIGNALING | 0,2804233 0,8352792 0,1938133 -0,3638181 -1,0840273 | 51 |
| GOBP_INTERLEUKIN_4_PRODUCTION | 0,2948718 0,8468136 0,1682382 -0,4137557 -1,0847323 | 23 |
| GOBP_REGULATION_OF_EPIDERMIS_DEVELOPMENT | 0,28125 0,8359478 0,1918922 -0,366328 -1,0850431 | 50 |
| GOBP_REGULATION_OF_RESPONSE_TO_WOUNDING | 0,2745098 0,8289344 0,2712886 -0,3198384 -1,085121 | 121 |
| GOBP_REGULATION_OF_EPIDERMAL_GROWTH_FACTOR_ACTIVATED_RECEPTOR_ACTIVITY | 0,3318777 0,8660673 0,1596467 -0,4058835 -1,0855931 | 25 |
| GOBP_VESICLE_BUDDING_FROM_MEMBRANE | 0,2923977 0,8455459 0,1999152 -0,3610097 -1,0872585 | 62 |
| GOBP_NEGATIVE_REGULATION_OF_ACTIN_FILAMENT_BUNDLE_ASSEMBLY | 0,3274336 0,8660673 0,1619789 -0,4001193 -1,0895842 | 29 |
| GOBP_MULTICELLULAR_ORGANISM_AGING | 0,3253589 0,8660673 0,1695706 -0,393117 -1,0897302 | 35 |
| GOBP_NEURAL_CREST_CELL_MIGRATION | 0,2698413 0,8262377 0,197822 -0,3661114 -1,0908603 | 51 |
| GOBP_SINGLE_FERTILIZATION | 0,3100775 0,8581768 0,2249661 -0,3462103 -1,0910104 | 85 |
| GOBP_STRIATED_MUSCLE_CONTRACTION | 0,2857143 0,8386161 0,2820134 -0,3218924 -1,0921144 | 138 |
| GOBP_REGULATION_OF_ENDOPLASMIC_RETICULUM_UNFOLDED_PROTEIN_RESPONSE | 0,3247863 0,8660673 0,1596467 -0,401417 -1,0935564 | 27 |
| GOBP_POSITIVE_REGULATION_OF_RESPONSE_TO_ENDOPLASMIC_RETICULUM_STRESS | 0,3047619 0,8565046 0,175204 -0,4040611 -1,0939179 | 32 |
| GOBP_ENDODERM_DEVELOPMENT | 0,2771084 0,8322113 0,208955 -0,3612973 -1,0951429 | 64 |
| GOBP_RESPONSE_TO_CALCIUM_ION | 0,26 0,8179308 0,2820134 -0,3248032 -1,0970687 | 123 |
| GOBP_NEGATIVE_REGULATION_OF_LEUKOCYTE_PROLIFERATION | 0,2604167 0,8179308 0,1999152 -0,3746281 -1,1000255 | 49 |
| GOBP_PROTEIN_CONTAINING_COMPLEX_REMODELING | 0,3257576 0,8660673 0,1492075 -0,4412399 -1,1009511 | 15 |
| GOBP_FERTILIZATION | 0,2692308 0,8262377 0,2712886 -0,3325889 -1,1019681 | 113 |
| GOBP_POSITIVE_REGULATION_OF_LOCOMOTION | 0,1371985 0,6680012 0,2492466 -0,2798485 -1,1028388 | 455 |
| GOBP_REGULATION_OF_T_CELL_APOPTOTIC_PROCESS | 0,3144105 0,8598592 0,1644058 -0,4124611 -1,1031857 | 25 |
| GOBP_POSITIVE_REGULATION_OF_T_CELL_MIGRATION | 0,284 0,8386161 0,1656567 -0,4301707 -1,1039848 | 20 |
| GOBP_MEMBRANE_RAFT_ORGANIZATION | 0,2828685 0,8376329 0,1656567 -0,4212687 -1,1047264 | 22 |
| GOBP_VIRAL_BUDDING | 0,3162393 0,861089 0,1619789 -0,4058701 -1,1056876 | 27 |
| GOBP_UBIQUITIN_DEPENDENT_PROTEIN_CATABOLIC_PROCESS_VIA_THE_MULTIVESICULAR_BODY_SORTING_PATHWAY | 0,3047619 0,8565046 0,175204 -0,401739 -1,1058435 | 34 |
| GOBP_DETECTION_OF_TEMPERATURE_STIMULUS | 0,3115385 0,8582281 0,154191 -0,4360757 -1,1077405 | 18 |
| GOBP_REGULATION_OF_MUSCLE_SYSTEM_PROCESS | 0,2179059 0,7873562 0,2492466 -0,3154511 -1,1080364 | 181 |
| GOBP_ESTABLISHMENT_OF_PROTEIN_LOCALIZATION_TO_ENDOPLASMIC_RETICULUM | 0,2744186 0,8289344 0,1830239 -0,3842739 -1,1113868 | 43 |
| GOBP_PROTEIN_O_LINKED_GLYCOSYLATION | 0,2805755 0,8352792 0,2279872 -0,3539981 -1,1121858 | 80 |
| GOBP_NEGATIVE_REGULATION_OF_ION_TRANSMEMBRANE_TRANSPORT | 0,2857143 0,8386161 0,2139279 -0,357304 -1,1122249 | 72 |
| GOBP_FOAM_CELL_DIFFERENTIATION | 0,2925764 0,8455459 0,1709323 -0,4159454 -1,1125052 | 25 |
| GOBP_POSITIVE_REGULATION_OF_INTERLEUKIN_1_PRODUCTION | 0,2560386 0,8138217 0,1938133 -0,3849476 -1,1127075 | 44 |
| GOBP_MULTICELLULAR_ORGANISMAL_MOVEMENT | 0,2614679 0,8179308 0,1864326 -0,3796431 -1,1158174 | 45 |
| GOBP_CELLULAR_RESPONSE_TO_GLUCOSE_STARVATION | 0,2383178 0,8012686 0,197822 -0,3791953 -1,1177984 | 47 |
| GOBP_CELL_CELL_JUNCTION_ORGANIZATION | 0,3372052 0,8693282 0,203509 -0,3152236 -1,1186324 | 172 |
| GOBP_LIPID_TRANSLOCATION | 0,234375 0,7983096 0,2114002 -0,3812978 -1,1196096 | 49 |
| GOBP_POSITIVE_REGULATION_OF_SMOOTH_MUSCLE_CONTRACTION | 0,2698413 0,8262377 0,1695706 -0,4349607 -1,120814 | 21 |
| GOBP_CARDIAC_MUSCLE_CONTRACTION | 0,2608696 0,8179308 0,2616635 -0,3364867 -1,1221781 | 107 |
| GOBP_IMMUNE_RESPONSE_TO_TUMOR_CELL | 0,2846154 0,8386161 0,1619789 -0,4419256 -1,1226007 | 18 |
| GOBP_POSITIVE_REGULATION_OF_LEUKOCYTE_DEGRANULATION | 0,2957198 0,8468136 0,1596467 -0,4424862 -1,1233815 | 17 |
| GOBP_RESPONSE_TO_GROWTH_HORMONE | 0,2731278 0,8289344 0,1782199 -0,4197364 -1,1254826 | 26 |
| GOBP_MAINTENANCE_OF_SYNAPSE_STRUCTURE | 0,268 0,8262377 0,1709323 -0,4391581 -1,1270498 | 20 |
| GOBP_REGULATION_OF_PEPTIDASE_ACTIVITY | 0,1106213 0,623409 0,2878051 -0,2969592 -1,1278737 | 328 |
| GOBP_RESPONSE_TO_MECHANICAL_STIMULUS | 0,199829 0,7744131 0,2492466 -0,3207227 -1,1286497 | 166 |
| GOBP_POSITIVE_REGULATION_OF_STRESS_FIBER_ASSEMBLY | 0,2511628 0,8106689 0,1918922 -0,3903656 -1,1290049 | 43 |
| GOBP_POSITIVE_REGULATION_OF_SIGNALING_RECEPTOR_ACTIVITY | 0,2829268 0,8376329 0,1847065 -0,4172334 -1,1290805 | 33 |
| GOBP_POSITIVE_REGULATION_OF_DENDRITIC_SPINE_MORPHOGENESIS | 0,2918288 0,8455459 0,1608014 -0,4448423 -1,1293632 | 17 |
| GOBP_REGULATION_OF_CELL_ADHESION_MEDIATED_BY_INTEGRIN | 0,3 0,8499289 0,1766943 -0,4110955 -1,1315985 | 34 |
| GOBP_AUTOPHAGY_OF_NUCLEUS | 0,2890625 0,8430641 0,1619789 -0,4497559 -1,1321976 | 16 |
| GOBP_POSITIVE_REGULATION_OF_EPITHELIAL_CELL_DIFFERENTIATION | 0,2336449 0,7983096 0,1999152 -0,3877561 -1,132323 | 46 |
| GOBP_REGULATION_OF_SYSTEMIC_ARTERIAL_BLOOD_PRESSURE | 0,2544379 0,8131684 0,2165428 -0,3739755 -1,133542 | 63 |
| GOBP_NEGATIVE_CHEMOTAXIS | 0,2372093 0,7998324 0,197822 -0,3931907 -1,1371756 | 43 |
| GOBP_EMBRYONIC_PATTERN_SPECIFICATION | 0,2245989 0,7901992 0,2192503 -0,3840117 -1,1386659 | 53 |
| GOBP_LUNG_ALVEOLUS_DEVELOPMENT | 0,2583732 0,8179127 0,1918922 -0,4039505 -1,1391261 | 37 |
| GOBP_TOXIN_TRANSPORT | 0,2904762 0,8446742 0,1797823 -0,4139081 -1,1393408 | 34 |
| GOBP_NEGATIVE_REGULATION_OF_ACTIN_FILAMENT_DEPOLYMERIZATION | 0,2644231 0,8203448 0,1900233 -0,4003112 -1,1415007 | 38 |
| GOBP_NEGATIVE_REGULATION_OF_PEPTIDASE_ACTIVITY | 0,1716647 0,7265238 0,2492466 -0,3298424 -1,1433672 | 161 |
| GOBP_STRESS_FIBER_ASSEMBLY | 0,224 0,7901992 0,2712886 -0,3541181 -1,1444408 | 95 |
| GOBP_PROTEIN_O_LINKED_MANNOSYLATION | 0,2695313 0,8262377 0,1682382 -0,4549839 -1,1453584 | 16 |
| GOBP_RESPONSE_TO_AMINO_ACID_STARVATION | 0,2083333 0,7744131 0,2249661 -0,3904903 -1,1466019 | 49 |
| GOBP_DETECTION_OF_MECHANICAL_STIMULUS | 0,2548077 0,8131684 0,1938133 -0,4022135 -1,1469251 | 38 |
| GOBP_NEGATIVE_REGULATION_OF_PEPTIDYL_TYROSINE_PHOSPHORYLATION | 0,2149533 0,7825382 0,208955 -0,3929787 -1,1475741 | 46 |
| GOBP_SKELETAL_MUSCLE_CONTRACTION | 0,2731707 0,8289344 0,1882041 -0,4241983 -1,1479283 | 33 |
| GOBP_NEGATIVE_REGULATION_OF_EPITHELIAL_CELL_APOPTOTIC_PROCESS | 0,2679426 0,8262377 0,1882041 -0,4147759 -1,1497691 | 35 |
| GOBP_NEGATIVE_REGULATION_OF_SODIUM_ION_TRANSPORT | 0,2607004 0,8179308 0,1709323 -0,4530883 -1,1502981 | 17 |
| GOBP_REGULATION_OF_RIG_I_SIGNALING_PATHWAY | 0,25 0,8103969 0,1737478 -0,4532396 -1,151341 | 18 |
| GOBP_TRIGLYCERIDE_BIOSYNTHETIC_PROCESS | 0,2557078 0,8134581 0,1882041 -0,4224222 -1,1521996 | 30 |
| GOBP_NEGATIVE_REGULATION_OF_ION_TRANSPORT | 0,239741 0,8013406 0,2492466 -0,3485397 -1,1522733 | 104 |
| GOBP_INTERFERON_ALPHA_PRODUCTION | 0,247012 0,8075292 0,1782199 -0,4394873 -1,1525024 | 22 |
| GOBP_DENDRITE_EXTENSION | 0,2585366 0,8179127 0,1938133 -0,4262704 -1,1535356 | 33 |
| GOBP_REGULATION_OF_TUBE_SIZE | 0,203252 0,7744131 0,2878571 -0,3478995 -1,1537054 | 100 |
| GOBP_SENSORY_PERCEPTION_OF_TEMPERATURE_STIMULUS | 0,24 0,8014933 0,1813831 -0,4501011 -1,1551339 | 20 |
| GOBP_NEGATIVE_REGULATION_OF_ERK1_AND_ERK2_CASCADE | 0,245614 0,8075292 0,2192503 -0,3838748 -1,1561218 | 62 |
| GOBP_RESPONSE_TO_CHEMOKINE | 0,2155963 0,7830255 0,2065879 -0,393866 -1,1576203 | 45 |
| GOBP_CELLULAR_RESPONSE_TO_EXTERNAL_STIMULUS | 0,0772318 0,561238 0,2878051 -0,321959 -1,1579769 | 266 |
| GOBP_THYROID_HORMONE_METABOLIC_PROCESS | 0,2689394 0,8262377 0,1656567 -0,4647144 -1,1595231 | 15 |
| GOBP_EPITHELIAL_CELL_DEVELOPMENT | 0,2458376 0,8075292 0,2492466 -0,3254976 -1,1614253 | 171 |
| GOBP_ENDOPLASMIC_RETICULUM_CALCIUM_ION_HOMEOSTASIS | 0,2478261 0,8080806 0,1864326 -0,440278 -1,1650543 | 24 |
| GOBP_DIOL_BIOSYNTHETIC_PROCESS | 0,2301587 0,7963341 0,1847065 -0,453141 -1,1676612 | 21 |
| GOBP_KERATINOCYTE_MIGRATION | 0,233463 0,7983096 0,1813831 -0,4600518 -1,1679768 | 17 |
| GOBP_NEGATIVE_REGULATION_OF_POTASSIUM_ION_TRANSPORT | 0,228 0,7916784 0,1864326 -0,456145 -1,1706449 | 20 |
| GOBP_MYD88_DEPENDENT_TOLL_LIKE_RECEPTOR_SIGNALING_PATHWAY | 0,2539063 0,8131684 0,1737478 -0,4650883 -1,1707948 | 16 |
| GOBP_TRANSFORMING_GROWTH_FACTOR_BETA_PRODUCTION | 0,2372093 0,7998324 0,197822 -0,4290185 -1,1709073 | 31 |
| GOBP_REGULATION_OF_GRANULOCYTE_CHEMOTAXIS | 0,2392344 0,8012686 0,1999152 -0,4224537 -1,1710523 | 35 |
| GOBP_REGULATION_OF_BODY_FLUID_LEVELS | 0,0510366 0,4576243 0,3217759 -0,3238971 -1,17197 | 275 |
| GOBP_POSITIVE_REGULATION_OF_WOUND_HEALING | 0,2415459 0,8030878 0,1999152 -0,4063721 -1,1726815 | 40 |
| GOBP_POSITIVE_REGULATION_OF_NUCLEOCYTOPLASMIC_TRANSPORT | 0,2032967 0,7744131 0,2343926 -0,394741 -1,1739807 | 55 |
| GOBP_PROTEIN_EXPORT_FROM_NUCLEUS | 0,2134831 0,7803554 0,2311267 -0,3911378 -1,1742069 | 58 |
| GOBP_SKELETAL_MUSCLE_ORGAN_DEVELOPMENT | 0,1566199 0,6993542 0,2492466 -0,3463377 -1,175572 | 134 |
| GOBP_PROTEIN_HYDROXYLATION | 0,2521368 0,8117205 0,1830239 -0,4322722 -1,1776133 | 27 |
| GOBP_POSTSYNAPSE_ORGANIZATION | 0,2125594 0,7803554 0,2492466 -0,3411827 -1,1782526 | 146 |
| GOBP_CELL_DIFFERENTIATION_INVOLVED_IN_METANEPHROS_DEVELOPMENT | 0,2230769 0,7901992 0,1847065 -0,464912 -1,1809919 | 18 |
| GOBP_POSITIVE_REGULATION_OF_INTERFERON_ALPHA_PRODUCTION | 0,2230769 0,7901992 0,1847065 -0,4661829 -1,1842201 | 18 |
| GOBP_POSITIVE_REGULATION_OF_COLD_INDUCED_THERMOGENESIS | 0,2206897 0,7888973 0,2529611 -0,3811534 -1,1847052 | 76 |
| GOBP_NERVE_DEVELOPMENT | 0,2242424 0,7901992 0,2343926 -0,3878363 -1,1867236 | 65 |
| GOBP_INTRINSIC_APOPTOTIC_SIGNALING_PATHWAY_IN_RESPONSE_TO_ENDOPLASMIC_RETICULUM_STRESS | 0,1966292 0,7683417 0,24134 -0,396931 -1,1878458 | 57 |
| GOBP_COLLAGEN_CATABOLIC_PROCESS | 0,2435897 0,8052169 0,1864326 -0,431452 -1,1884389 | 28 |
| GOBP_RESPONSE_TO_WOUNDING | 0,0591777 0,4845135 0,3217759 -0,3018215 -1,1886079 | 423 |
| GOBP_POSITIVE_REGULATION_OF_EPIDERMIS_DEVELOPMENT | 0,2391304 0,8012686 0,1900233 -0,4492584 -1,1888181 | 24 |
| GOBP_POSITIVE_REGULATION_OF_LEUKOCYTE_APOPTOTIC_PROCESS | 0,2094862 0,7750512 0,1938133 -0,4613239 -1,1906379 | 19 |
| GOBP_SEGMENTATION | 0,2206897 0,7888973 0,2529611 -0,3830963 -1,1907442 | 76 |
| GOBP_CELL_JUNCTION_MAINTENANCE | 0,2211538 0,7889462 0,208955 -0,4178391 -1,1914821 | 38 |
| GOBP_POSITIVE_REGULATION_OF_INTERLEUKIN_1_BETA_PRODUCTION | 0,2248804 0,7901992 0,2065879 -0,4232901 -1,1936631 | 37 |
| GOBP_CELL_VOLUME_HOMEOSTASIS | 0,2350427 0,7984227 0,1900233 -0,4345145 -1,1968745 | 28 |
| GOBP_CRANIAL_NERVE_DEVELOPMENT | 0,2173913 0,7873562 0,2114002 -0,4148626 -1,1971829 | 40 |
| GOBP_ESTABLISHMENT_OF_ENDOTHELIAL_BARRIER | 0,1699029 0,7245462 0,24134 -0,405689 -1,1987851 | 48 |
| GOBP_REGULATION_OF_VASOCONSTRICTION | 0,1860465 0,7540202 0,2249661 -0,4146548 -1,1992537 | 43 |
| GOBP_MUSCLE_SYSTEM_PROCESS | 0,0270873 0,3314885 0,3524879 -0,3175757 -1,1998478 | 338 |
| GOBP_NEGATIVE_REGULATION_OF_RECEPTOR_MEDIATED_ENDOCYTOSIS | 0,2222222 0,7901992 0,195789 -0,4360067 -1,2009847 | 28 |
| GOBP_BASEMENT_MEMBRANE_ASSEMBLY | 0,2310606 0,7978812 0,1797823 -0,4813774 -1,2010994 | 15 |
| GOBP_MORPHOGENESIS_OF_AN_EPITHELIAL_SHEET | 0,1923077 0,7680255 0,24134 -0,4039063 -1,2012387 | 55 |
| GOBP_PHARYNGEAL_SYSTEM_DEVELOPMENT | 0,2063492 0,7744131 0,195789 -0,4675665 -1,2048329 | 21 |
| GOBP_RESPONSE_TO_FUNGUS | 0,2063492 0,7744131 0,195789 -0,4677148 -1,2052151 | 21 |
| GOBP_NEURON_PROJECTION_EXTENSION_INVOLVED_IN_NEURON_PROJECTION_GUIDANCE | 0,2057416 0,7744131 0,2165428 -0,4338147 -1,2054417 | 36 |
| GOBP_RELAXATION_OF_CARDIAC_MUSCLE | 0,2272727 0,7916784 0,1813831 -0,4855425 -1,211492 | 15 |
| GOBP_NEGATIVE_REGULATION_OF_CHEMOTAXIS | 0,1635514 0,7148626 0,24134 -0,4110577 -1,2117229 | 47 |
| GOBP_REGULATION_OF_RYANODINE_SENSITIVE_CALCIUM_RELEASE_CHANNEL_ACTIVITY | 0,2063492 0,7744131 0,195789 -0,4712836 -1,2144112 | 21 |
| GOBP_INSULIN_LIKE_GROWTH_FACTOR_RECEPTOR_SIGNALING_PATHWAY | 0,2222222 0,7901992 0,195789 -0,4435261 -1,2216968 | 28 |
| GOBP_TRANSCYTOSIS | 0,21875 0,7879614 0,1882041 -0,4856113 -1,2224586 | 16 |
| GOBP_SUPEROXIDE_ANION_GENERATION | 0,2070485 0,7744131 0,2065879 -0,4560625 -1,2228875 | 26 |
| GOBP_REGULATION_OF_PROGRAMMED_NECROTIC_CELL_DEATH | 0,2086957 0,7744131 0,2042948 -0,4624155 -1,223634 | 24 |
| GOBP_POSITIVE_REGULATION_OF_EPIDERMAL_CELL_DIFFERENTIATION | 0,196 0,7683417 0,2020717 -0,4773898 -1,2251672 | 20 |
| GOBP_POSITIVE_REGULATION_OF_RESPONSE_TO_WOUNDING | 0,171875 0,7265238 0,2489111 -0,414623 -1,2280904 | 50 |
| GOBP_CHOLESTEROL_STORAGE | 0,2083333 0,7744131 0,1900233 -0,4925114 -1,2288802 | 15 |
| GOBP_REGULATION_OF_ACTOMYOSIN_STRUCTURE_ORGANIZATION | 0,104957 0,6209246 0,2878051 -0,3942699 -1,229476 | 84 |
| GOBP_CALCIUM_ION_IMPORT | 0,1975309 0,7692965 0,2529611 -0,3987503 -1,2302715 | 68 |
| GOBP_POSITIVE_REGULATION_OF_KERATINOCYTE_DIFFERENTIATION | 0,2045455 0,7744131 0,1918922 -0,4932349 -1,2306853 | 15 |
| GOBP_REGULATION_OF_KERATINOCYTE_DIFFERENTIATION | 0,2136752 0,7803554 0,1999152 -0,4471078 -1,2315628 | 28 |
| GOBP_T_CELL_APOPTOTIC_PROCESS | 0,1730769 0,7265238 0,2377938 -0,4319494 -1,231718 | 38 |
| GOBP_MAINTENANCE_OF_CELL_POLARITY | 0,2038462 0,7744131 0,1938133 -0,4850019 -1,2320251 | 18 |
| GOBP_REGULATION_OF_DENDRITE_EXTENSION | 0,2094017 0,7750512 0,2020717 -0,4709731 -1,2347377 | 23 |
| GOBP_NEGATIVE_REGULATION_OF_TRANSPORTER_ACTIVITY | 0,1952663 0,7683417 0,2489111 -0,4080027 -1,2366803 | 63 |
| GOBP_NEGATIVE_REGULATION_OF_AXONOGENESIS | 0,1929825 0,7680255 0,2489111 -0,411098 -1,2381103 | 62 |
| GOBP_PHOSPHOLIPASE_C_ACTIVATING_G_PROTEIN_COUPLED_RECEPTOR_SIGNALING_PATHWAY | 0,1741573 0,7286292 0,2572065 -0,4141936 -1,2395053 | 57 |
| GOBP_ACTIN_FILAMENT_ORGANIZATION | 0,022429 0,2884173 0,3524879 -0,3219711 -1,2438732 | 389 |
| GOBP_REGULATION_OF_B_CELL_PROLIFERATION | 0,1732673 0,7265238 0,24134 -0,4358756 -1,2448726 | 39 |
| GOBP_NEGATIVE_REGULATION_OF_AXON_EXTENSION | 0,1753555 0,7308245 0,2343926 -0,4295239 -1,2450718 | 42 |
| GOBP_POSITIVE_REGULATION_OF_MYOBLAST_DIFFERENTIATION | 0,1969697 0,7684477 0,195789 -0,4998267 -1,2471329 | 15 |
| GOBP_REGULATION_OF_MACROPHAGE_DERIVED_FOAM_CELL_DIFFERENTIATION | 0,1778656 0,7367598 0,2114002 -0,4838998 -1,2489045 | 19 |
| GOBP_VASCULAR_ASSOCIATED_SMOOTH_MUSCLE_CONTRACTION | 0,1912351 0,7667771 0,2042948 -0,4764471 -1,2494251 | 22 |
| GOBP_NEGATIVE_REGULATION_OF_ATP_DEPENDENT_ACTIVITY | 0,1884615 0,7580835 0,2020717 -0,4918896 -1,2495216 | 18 |
| GOBP_POSITIVE_REGULATION_OF_INTERLEUKIN_10_PRODUCTION | 0,1894273 0,7603402 0,2165428 -0,4663635 -1,2505086 | 26 |
| GOBP_REGULATION_OF_NMDA_RECEPTOR_ACTIVITY | 0,1884615 0,7580835 0,2020717 -0,4929161 -1,252129 | 18 |
| GOBP_CELLULAR_RESPONSE_TO_FATTY_ACID | 0,1834061 0,7473501 0,2192503 -0,4682076 -1,2522879 | 25 |
| GOBP_HAIR_FOLLICLE_MATURATION | 0,1931818 0,7680255 0,197822 -0,5019961 -1,2525458 | 15 |
| GOBP_POSITIVE_REGULATION_OF_LIPID_STORAGE | 0,203125 0,7744131 0,195789 -0,4998643 -1,2583386 | 16 |
| GOBP_XENOBIOTIC_TRANSPORT | 0,1658537 0,7178712 0,2450418 -0,4652125 -1,2589175 | 33 |
| GOBP_POSITIVE_REGULATION_OF_PATTERN_RECOGNITION_RECEPTOR_SIGNALING_PATHWAY | 0,15311 0,6935605 0,2529611 -0,4550958 -1,2615369 | 35 |
| GOBP_POSITIVE_REGULATION_OF_ALPHA_BETA_T_CELL_PROLIFERATION | 0,1945525 0,7683417 0,1999152 -0,4969602 -1,2616798 | 17 |
| GOBP_PROTEIN_LOCALIZATION_TO_CELL_CELL_JUNCTION | 0,1692308 0,723322 0,2139279 -0,5011288 -1,2729914 | 18 |
| GOBP_REGULATION_OF_STRIATED_MUSCLE_CONTRACTION | 0,1686574 0,7227339 0,2492466 -0,4077547 -1,2752105 | 73 |
| GOBP_SYNCYTIUM_FORMATION | 0,1497585 0,6878113 0,2572065 -0,4421986 -1,2760671 | 40 |
| GOBP_NATURAL_KILLER_CELL_MEDIATED_IMMUNITY | 0,144186 0,684728 0,2572065 -0,4412334 -1,2761234 | 43 |
| GOBP_ENDOPLASMIC_RETICULUM_TO_CYTOSOL_TRANSPORT | 0,1666667 0,7197318 0,2279872 -0,4636147 -1,2770311 | 28 |
| GOBP_CELLULAR_COMPONENT_MAINTENANCE | 0,1629213 0,7137481 0,2663507 -0,4271801 -1,2783682 | 57 |
| GOBP_BASEMENT_MEMBRANE_ORGANIZATION | 0,159292 0,7040708 0,2377938 -0,4697782 -1,2792757 | 29 |
| GOBP_REGULATION_OF_SYSTEMIC_ARTERIAL_BLOOD_PRESSURE_MEDIATED_BY_A_CHEMICAL_SIGNAL | 0,152381 0,6931829 0,2529611 -0,4729523 -1,2804276 | 32 |
| GOBP_SKIN_DEVELOPMENT | 0,0467085 0,4458081 0,3217759 -0,3653569 -1,2832335 | 186 |
| GOBP_NEGATIVE_REGULATION_OF_CALCIUM_ION_TRANSPORT | 0,1516588 0,691483 0,2529611 -0,4427707 -1,2834708 | 42 |
| GOBP_REGULATION_OF_THE_FORCE_OF_HEART_CONTRACTION | 0,1712062 0,7265238 0,2139279 -0,507281 -1,2878823 | 17 |
| GOBP_CHONDROCYTE_DEVELOPMENT | 0,1581197 0,7013643 0,2343926 -0,467631 -1,2880941 | 28 |
| GOBP_AMINOGLYCAN_CATABOLIC_PROCESS | 0,1314741 0,6552847 0,2489111 -0,4919336 -1,2900366 | 22 |
| GOBP_DIGESTION | 0,1656442 0,7178712 0,2765006 -0,4168151 -1,2908697 | 69 |
| GOBP_PLASMA_MEMBRANE_TUBULATION | 0,15 0,6878113 0,2279872 -0,5083556 -1,2913492 | 18 |
| GOBP_EPIDERMIS_DEVELOPMENT | 0,0172953 0,247066 0,3524879 -0,3598278 -1,2930297 | 223 |
| GOBP_CARDIAC_MUSCLE_TISSUE_DEVELOPMENT | 0,0334424 0,3758353 0,3217759 -0,3683488 -1,2933894 | 178 |
| GOBP_ENDOTHELIAL_CELL_DEVELOPMENT | 0,1578947 0,7011945 0,2765006 -0,4313515 -1,2987881 | 61 |
| GOBP_REGULATION_OF_LEUKOCYTE_CHEMOTAXIS | 0,1868884 0,7558018 0,2492466 -0,4079412 -1,2991852 | 81 |
| GOBP_POSITIVE_REGULATION_OF_NEUTROPHIL_MIGRATION | 0,1452991 0,684728 0,2450418 -0,4959121 -1,3001195 | 23 |
| GOBP_REGULATION_OF_CARDIAC_MUSCLE_CONTRACTION | 0,1485714 0,6878113 0,2820134 -0,4335712 -1,3029571 | 59 |
| GOBP_NEGATIVE_REGULATION_OF_INTERFERON_GAMMA_PRODUCTION | 0,14 0,6759789 0,24134 -0,5077549 -1,3030958 | 20 |
| GOBP_CALCIUM_ION_REGULATED_EXOCYTOSIS_OF_NEUROTRANSMITTER | 0,155642 0,6961275 0,2249661 -0,5133584 -1,3033114 | 17 |
| GOBP_DENDRITIC_SPINE_MAINTENANCE | 0,171875 0,7265238 0,2139279 -0,5177474 -1,3033567 | 16 |
| GOBP_VENTRICULAR_CARDIAC_MUSCLE_CELL_ACTION_POTENTIAL | 0,1452991 0,684728 0,2450418 -0,4732192 -1,3034868 | 28 |
| GOBP_MESENCHYME_DEVELOPMENT | 0,0176068 0,250563 0,3524879 -0,3657101 -1,3089584 | 244 |
| GOBP_IRE1_MEDIATED_UNFOLDED_PROTEIN_RESPONSE | 0,1439689 0,684728 0,2343926 -0,5164328 -1,3111166 | 17 |
| GOBP_POSITIVE_REGULATION_OF_VASOCONSTRICTION | 0,1309524 0,6552847 0,2489111 -0,5089761 -1,3115381 | 21 |
| GOBP_WATER_HOMEOSTASIS | 0,1374408 0,6680012 0,2663507 -0,4525361 -1,311778 | 42 |
| GOBP_MUSCLE_CELL_DIFFERENTIATION | 0,0182319 0,2575078 0,3524879 -0,3498897 -1,3120908 | 300 |
| GOBP_REGULATION_OF_HEART_RATE | 0,1170561 0,6334603 0,2878051 -0,4221318 -1,3164979 | 79 |
| GOBP_SECONDARY_PALATE_DEVELOPMENT | 0,1282051 0,6540073 0,2616635 -0,5024866 -1,3173555 | 23 |
| GOBP_POSITIVE_REGULATION_OF_CYTOSOLIC_CALCIUM_ION_CONCENTRATION_INVOLVED_IN_PHOSPHOLIPASE_C_ACTIVATING_G_PROTEIN_COUPLED_SIGNALING_PATHWAY | 0,132 0,6559841 0,2489111 -0,514083 -1,3193362 | 20 |
| GOBP_REGULATION_OF_CARDIAC_MUSCLE_CELL_ACTION_POTENTIAL | 0,1347826 0,6619324 0,2572065 -0,5008524 -1,325345 | 24 |
| GOBP_STORE_OPERATED_CALCIUM_ENTRY | 0,1423077 0,6810828 0,2343926 -0,5217888 -1,3254729 | 18 |
| GOBP_DETECTION_OF_TEMPERATURE_STIMULUS_INVOLVED_IN_SENSORY_PERCEPTION | 0,155303 0,6960014 0,222056 -0,5323164 -1,3281988 | 15 |
| GOBP_REGULATION_OF_MUSCLE_CONTRACTION | 0,1009853 0,6180254 0,2878051 -0,3894776 -1,3282686 | 126 |
| GOBP_MUSCLE_TISSUE_DEVELOPMENT | 0,0079518 0,1422617 0,3807304 -0,3541618 -1,3348484 | 318 |
| GOBP_CARDIOCYTE_DIFFERENTIATION | 0,0849717 0,5824129 0,2878051 -0,4022116 -1,3373457 | 114 |
| GOBP_MACROPHAGE_DIFFERENTIATION | 0,1244019 0,6437713 0,2820134 -0,4830734 -1,3423171 | 36 |
| GOBP_RESPONSE_TO_TESTOSTERONE | 0,1162791 0,6334603 0,2878571 -0,4935177 -1,3469432 | 31 |
| GOBP_FORMATION_OF_PRIMARY_GERM_LAYER | 0,0559735 0,4725528 0,3217759 -0,4121906 -1,3470255 | 97 |
| GOBP_ENDODERMAL_CELL_DIFFERENTIATION | 0,1287617 0,6540073 0,2492466 -0,47409 -1,3540138 | 39 |
| GOBP_NEGATIVE_REGULATION_OF_ION_TRANSMEMBRANE_TRANSPORTER_ACTIVITY | 0,0678165 0,5171182 0,2878051 -0,4544059 -1,3545454 | 56 |
| GOBP_MUSCLE_CONTRACTION | 0,005146 0,1033661 0,4070179 -0,3742939 -1,3650002 | 268 |
| GOBP_VESICLE_CARGO_LOADING | 0,108 0,6209246 0,2765006 -0,5334938 -1,3691518 | 20 |
| GOBP_REGULATION_OF_METALLOPEPTIDASE_ACTIVITY | 0,1174242 0,6334603 0,2572065 -0,5489058 -1,3695917 | 15 |
| GOBP_HYPEROSMOTIC_RESPONSE | 0,108 0,6209246 0,2765006 -0,5342264 -1,3710319 | 20 |
| GOBP_POSITIVE_REGULATION_OF_LEUKOCYTE_CHEMOTAXIS | 0,0677217 0,5171182 0,2878051 -0,452727 -1,3722423 | 63 |
| GOBP_LENS_FIBER_CELL_DIFFERENTIATION | 0,1162791 0,6334603 0,2878571 -0,5028304 -1,37236 | 31 |
| GOBP_PLATELET_MORPHOGENESIS | 0,109375 0,6226089 0,2712886 -0,5460464 -1,3745957 | 16 |
| GOBP_ALPHA_BETA_T_CELL_PROLIFERATION | 0,1162791 0,6334603 0,2878571 -0,5042184 -1,3761483 | 31 |
| GOBP_HEART_PROCESS | 0,0300212 0,3578315 0,3524879 -0,3856854 -1,3768822 | 194 |
| GOBP_WOUND_HEALING | 0,0040032 0,0845138 0,4070179 -0,3627652 -1,3778098 | 328 |
| GOBP_NEGATIVE_REGULATION_OF_ENDOPLASMIC_RETICULUM_STRESS_INDUCED_INTRINSIC_APOPTOTIC_SIGNALING_PATHWAY | 0,1167315 0,6334603 0,2616635 -0,5443259 -1,3819315 | 17 |
| GOBP_INTERMEDIATE_FILAMENT_BASED_PROCESS | 0,1213189 0,6395918 0,2878051 -0,4842586 -1,3830555 | 39 |
| GOBP_RESPONSE_TO_FATTY_ACID | 0,1068976 0,6209246 0,2878051 -0,4712612 -1,3850941 | 45 |
| GOBP_VASOCONSTRICTION | 0,0579524 0,4816976 0,3217759 -0,4671972 -1,3926753 | 56 |
| GOBP_TISSUE_MIGRATION | 0,0047712 0,0974202 0,4070179 -0,385571 -1,396077 | 263 |
| GOBP_ECTODERM_DEVELOPMENT | 0,1115385 0,6263281 0,2663507 -0,549647 -1,3962397 | 18 |
| GOBP_REGULATION_OF_PROTEIN_EXPORT_FROM_NUCLEUS | 0,0678451 0,5171182 0,2878051 -0,5119348 -1,3972085 | 31 |
| GOBP_NEUTRAL_LIPID_CATABOLIC_PROCESS | 0,1175194 0,6334603 0,2878051 -0,5241066 -1,4017977 | 25 |
| GOBP_WATER_SOLUBLE_VITAMIN_METABOLIC_PROCESS | 0,0554864 0,4725528 0,3217759 -0,4703137 -1,4019652 | 56 |
| GOBP_MUSCLE_CELL_CELLULAR_HOMEOSTASIS | 0,1011673 0,6180254 0,2820134 -0,5556171 -1,4105975 | 17 |
| GOBP_ANATOMICAL_STRUCTURE_ARRANGEMENT | 0,094697 0,6050565 0,2878571 -0,5665833 -1,4136992 | 15 |
| GOBP_SYMPATHETIC_NERVOUS_SYSTEM_DEVELOPMENT | 0,1115385 0,6263281 0,2663507 -0,55834 -1,418322 | 18 |
| GOBP_LIVER_MORPHOGENESIS | 0,1076923 0,6209246 0,2712886 -0,5595513 -1,4213991 | 18 |
| GOBP_POSITIVE_REGULATION_OF_MACROPHAGE_MIGRATION | 0,1038462 0,6209246 0,2765006 -0,5616254 -1,4266679 | 18 |
| GOBP_STRIATED_MUSCLE_CELL_DIFFERENTIATION | 0,0104889 0,1759227 0,3807304 -0,3994145 -1,427329 | 214 |
| GOBP_BRANCHING_INVOLVED_IN_BLOOD_VESSEL_MORPHOGENESIS | 0,1026277 0,6197349 0,2878051 -0,5398937 -1,4286553 | 24 |
| GOBP_RENAL_ABSORPTION | 0,0976563 0,6130256 0,2878571 -0,5693333 -1,4332172 | 16 |
| GOBP_NEGATIVE_REGULATION_OF_AXON_EXTENSION_INVOLVED_IN_AXON_GUIDANCE | 0,0947116 0,6050565 0,2878051 -0,5345718 -1,4334025 | 26 |
| GOBP_REGULATION_OF_CALCIUM_ION_IMPORT | 0,1066743 0,6209246 0,2878051 -0,5295452 -1,433642 | 32 |
| GOBP_REGULATION_OF_INSULIN_LIKE_GROWTH_FACTOR_RECEPTOR_SIGNALING_PATHWAY | 0,1133289 0,6289167 0,2878051 -0,574665 -1,4338642 | 15 |
| GOBP_MUSCLE_ORGAN_DEVELOPMENT | 0,0016112 0,0438636 0,4550599 -0,3997705 -1,4341706 | 265 |
| GOBP_B_CELL_PROLIFERATION | 0,0759724 0,5563902 0,2878051 -0,4852088 -1,4371616 | 50 |
| GOBP_BODY_FLUID_SECRETION | 0,0812285 0,5719984 0,2878051 -0,4640666 -1,4372067 | 69 |
| GOBP_PHOSPHATIDYLSERINE_METABOLIC_PROCESS | 0,1504873 0,6878113 0,2492466 -0,5493002 -1,4404735 | 22 |
| GOBP_EPIDERMAL_CELL_DIFFERENTIATION | 0,0413123 0,4246236 0,3217759 -0,4244529 -1,4431974 | 127 |
| GOBP_SMOOTH_MUSCLE_CONTRACTION | 0,0620715 0,4981249 0,3217759 -0,4526296 -1,4435789 | 82 |
| GOBP_ENDODERM_FORMATION | 0,0836042 0,5801999 0,2878051 -0,4959435 -1,4482512 | 46 |
| GOBP_CARDIAC_MUSCLE_CELL_DIFFERENTIATION | 0,0922982 0,5997765 0,2878051 -0,4617203 -1,453524 | 87 |
| GOBP_INTERMEDIATE_FILAMENT_ORGANIZATION | 0,1027668 0,6197349 0,2820134 -0,5671039 -1,4636471 | 19 |
| GOBP_ANGIOGENESIS_INVOLVED_IN_WOUND_HEALING | 0,0978236 0,6130256 0,2878051 -0,5476292 -1,4647122 | 25 |
| GOBP_REGULATION_OF_POSTSYNAPSE_ORGANIZATION | 0,0819963 0,5747391 0,2878051 -0,4688072 -1,4661458 | 73 |
| GOBP_RESPONSE_TO_MUSCLE_STRETCH | 0,1432641 0,6837026 0,2492466 -0,5625063 -1,4751049 | 22 |
| GOBP_REGULATION_OF_SYNCYTIUM_FORMATION_BY_PLASMA_MEMBRANE_FUSION | 0,1082257 0,6209246 0,2878051 -0,5817747 -1,4770064 | 17 |
| GOBP_POSITIVE_REGULATION_OF_GRANULOCYTE_CHEMOTAXIS | 0,083847 0,5801999 0,2878051 -0,5821269 -1,4787467 | 18 |
| GOBP_TRIGLYCERIDE_CATABOLIC_PROCESS | 0,0816335 0,5732653 0,2878051 -0,5878831 -1,4799137 | 16 |
| GOBP_ADRENERGIC_RECEPTOR_SIGNALING_PATHWAY | 0,1240032 0,6435974 0,2878051 -0,565105 -1,4815207 | 23 |
| GOBP_LIPID_IMPORT_INTO_CELL | 0,106495 0,6209246 0,2878051 -0,5938214 -1,4816619 | 15 |
| GOBP_NITRIC_OXIDE_MEDIATED_SIGNAL_TRANSDUCTION | 0,1349654 0,661965 0,2492466 -0,5757975 -1,4860845 | 19 |
| GOBP_POSITIVE_REGULATION_OF_VACUOLE_ORGANIZATION | 0,1211119 0,6395918 0,2878051 -0,5807119 -1,4963879 | 21 |
| GOBP_ADENYLATE_CYCLASE_ACTIVATING_ADRENERGIC_RECEPTOR_SIGNALING_PATHWAY | 0,130188 0,6552847 0,2492466 -0,5799985 -1,4969271 | 19 |
| GOBP_CRANIAL_NERVE_MORPHOGENESIS | 0,1214333 0,6395918 0,2878051 -0,5733963 -1,5032578 | 23 |
| GOBP_REGULATION_OF_ENDOPLASMIC_RETICULUM_STRESS_INDUCED_INTRINSIC_APOPTOTIC_SIGNALING_PATHWAY | 0,0726036 0,5412925 0,2878051 -0,5466046 -1,5056277 | 28 |
| GOBP_CELLULAR_RESPONSE_TO_LIPOPROTEIN_PARTICLE_STIMULUS | 0,1124383 0,6263281 0,2878051 -0,5527089 -1,5057118 | 27 |
| GOBP_POSITIVE_REGULATION_OF_MACROPHAGE_CHEMOTAXIS | 0,1019392 0,6192943 0,2878051 -0,6034642 -1,5057219 | 15 |
| GOBP_PLASMINOGEN_ACTIVATION | 0,0918458 0,5997765 0,2878051 -0,5965772 -1,5145869 | 17 |
| GOBP_POSITIVE_REGULATION_OF_INTERLEUKIN_4_PRODUCTION | 0,0918458 0,5997765 0,2878051 -0,5970536 -1,5157962 | 17 |
| GOBP_RECEPTOR_CATABOLIC_PROCESS | 0,1085833 0,6209246 0,2878051 -0,5608046 -1,5277665 | 27 |
| GOBP_VASCULAR_WOUND_HEALING | 0,0848258 0,5824129 0,2878051 -0,6043047 -1,5342055 | 17 |
| GOBP_EPIBOLY | 0,0899204 0,5958215 0,2878051 -0,5543401 -1,5366445 | 35 |
| GOBP_SEMAPHORIN_PLEXIN_SIGNALING_PATHWAY | 0,0428971 0,4326394 0,3217759 -0,5319651 -1,5420209 | 42 |
| GOBP_VITAMIN_TRANSPORT | 0,0726142 0,5412925 0,2878051 -0,5702542 -1,543172 | 33 |
| GOBP_HETEROTYPIC_CELL_CELL_ADHESION | 0,0414791 0,4246236 0,3217759 -0,534716 -1,5499951 | 42 |
| GOBP_CARDIAC_CELL_DEVELOPMENT | 0,0581808 0,4821669 0,3217759 -0,5191506 -1,5635341 | 62 |
| GOBP_HYALURONAN_METABOLIC_PROCESS | 0,1021583 0,6192943 0,2878051 -0,5998268 -1,5725498 | 23 |
| GOBP_CELLULAR_RESPONSE_TO_MECHANICAL_STIMULUS | 0,0319945 0,3710617 0,3217759 -0,523656 -1,5767145 | 61 |
| GOBP_NEGATIVE_REGULATION_OF_TUMOR_NECROSIS_FACTOR_SUPERFAMILY_CYTOKINE_PRODUCTION | 0,0547606 0,4717694 0,3217759 -0,5609158 -1,5817627 | 37 |
| GOBP_ACYLGLYCEROL_HOMEOSTASIS | 0,060142 0,4890772 0,3217759 -0,6188668 -1,5882521 | 20 |
| GOBP_DIGESTIVE_SYSTEM_PROCESS | 0,0500095 0,4549289 0,3217759 -0,529931 -1,5908681 | 58 |
| GOBP_GLYCOSAMINOGLYCAN_CATABOLIC_PROCESS | 0,063766 0,5002749 0,2878051 -0,6335915 -1,6085586 | 17 |
| GOBP_MUSCLE_CELL_DEVELOPMENT | 0,0025145 0,0634035 0,4317077 -0,4721072 -1,6170519 | 142 |
| GOBP_PARASYMPATHETIC_NERVOUS_SYSTEM_DEVELOPMENT | 0,0476289 0,4476616 0,3217759 -0,6424582 -1,6172988 | 16 |
| GOBP_GANGLION_DEVELOPMENT | 0,0802987 0,5678014 0,2878051 -0,6578889 -1,6415185 | 15 |
| GOBP_ACTOMYOSIN_STRUCTURE_ORGANIZATION | 0,0008176 0,0253854 0,4772708 -0,471327 -1,6447874 | 175 |
| GOBP_POSITIVE_REGULATION_OF_BLOOD_CIRCULATION | 0,03064 0,3597322 0,3524879 -0,6089147 -1,6618927 | 31 |
| GOBP_LENS_DEVELOPMENT_IN_CAMERA_TYPE_EYE | 0,0387875 0,4070517 0,3217759 -0,5458735 -1,6647498 | 66 |
| GOBP_MOTOR_NEURON_AXON_GUIDANCE | 0,0497593 0,4544318 0,3217759 -0,6294853 -1,6879039 | 26 |
| GOBP_SKELETAL_MUSCLE_ADAPTATION | 0,0462793 0,4458081 0,3217759 -0,665422 -1,6893695 | 17 |
| GOBP_POSITIVE_REGULATION_OF_PROTEIN_EXPORT_FROM_NUCLEUS | 0,0462793 0,4458081 0,3217759 -0,6655585 -1,6897161 | 17 |
| GOBP_EPITHELIAL_CELL_CELL_ADHESION | 0,0300971 0,3578315 0,3524879 -0,672417 -1,6927159 | 16 |
| GOBP_T_CELL_CHEMOTAXIS | 0,070048 0,5284543 0,2878051 -0,6828727 -1,7038564 | 15 |
| GOBP_REGULATION_OF_ACTIN_FILAMENT_BASED_MOVEMENT | 0,0454551 0,4458081 0,3217759 -0,6159959 -1,707556 | 35 |
| GOBP_KERATINOCYTE_DIFFERENTIATION | 0,0158172 0,2339109 0,3524879 -0,5380056 -1,7134058 | 81 |
| GOBP_SARCOMERE_ORGANIZATION | 0,0425918 0,4324796 0,3217759 -0,618508 -1,7145195 | 35 |
| GOBP_REGULATION_OF_DIGESTIVE_SYSTEM_PROCESS | 0,0892494 0,5945215 0,2878051 -0,6568968 -1,7226326 | 22 |
| GOBP_CARDIAC_MYOFIBRIL_ASSEMBLY | 0,0404581 0,4184363 0,3217759 -0,6802893 -1,7271146 | 17 |
| GOBP_CELLULAR_RESPONSE_TO_OSMOTIC_STRESS | 0,0194105 0,2661509 0,3524879 -0,6229128 -1,7308891 | 36 |
| GOBP_FATTY_ACID_TRANSMEMBRANE_TRANSPORT | 0,0375475 0,3973034 0,3217759 -0,6871194 -1,7444548 | 17 |
| GOBP_ACROSOME_REACTION | 0,0372373 0,3973034 0,3217759 -0,6529391 -1,750793 | 26 |
| GOBP_POSITIVE_REGULATION_OF_HEART_RATE | 0,0513533 0,4583496 0,3217759 -0,699977 -1,8037122 | 21 |
| GOBP_ACTIN_FILAMENT_BASED_MOVEMENT | 0,0005051 0,0167934 0,4772708 -0,5452125 -1,8037644 | 112 |
| GOBP_REGULATION_OF_WATER_LOSS_VIA_SKIN | 0,0635857 0,5002749 0,2878051 -0,7005312 -1,8080118 | 19 |
| GOBP_SECRETION_BY_TISSUE | 0,0238789 0,3028159 0,3524879 -0,6644746 -1,8094627 | 29 |
| GOBP_PEPTIDE_CROSS_LINKING | 0,0196268 0,2681373 0,3524879 -0,7188253 -1,8095424 | 16 |
| GOBP_CARDIAC_MUSCLE_TISSUE_MORPHOGENESIS | 0,0217396 0,2850425 0,3524879 -0,6267662 -1,8302789 | 46 |
| GOBP_ACTIN_MEDIATED_CELL_CONTRACTION | 0,0012403 0,0356311 0,4550599 -0,581818 -1,8334795 | 85 |
| GOBP_CELLULAR_COMPONENT_ASSEMBLY_INVOLVED_IN_MORPHOGENESIS | 0,0030469 0,0699439 0,4317077 -0,5681211 -1,835054 | 92 |
| GOBP_KERATINIZATION | 0,0529859 0,465004 0,3217759 -0,7390224 -1,8439573 | 15 |
| GOBP_MESENCHYME_MORPHOGENESIS | 0,0102972 0,1734828 0,3807304 -0,6402647 -1,8873831 | 47 |
| GOBP_BLASTODERM_SEGMENTATION | 0,0467523 0,4458081 0,3217759 -0,756558 -1,8877108 | 15 |
| GOBP_MUSCLE_ORGAN_MORPHOGENESIS | 0,0009273 0,02787 0,4772708 -0,6427226 -1,9314936 | 59 |
| GOBP_CD8_POSITIVE_ALPHA_BETA_T_CELL_ACTIVATION | 0,0168227 0,2421563 0,3524879 -0,7581691 -1,9457556 | 20 |
| GOBP_STRIATED_MUSCLE_CELL_DEVELOPMENT | 0,0015248 0,0424339 0,4550599 -0,6698871 -2,0046873 | 57 |
| GOBP_SEMAPHORIN_PLEXIN_SIGNALING_PATHWAY_INVOLVED_IN_NEURON_PROJECTION_GUIDANCE | 0,0187501 0,2624788 0,3524879 -0,8332772 -2,0791354 | 15 |
